# Pond Water Microbiome Taxa Profiles and Antibiotic Resistance Genes Associated with Acidity and Tannins

**DOI:** 10.1101/2024.07.10.602992

**Authors:** Maya Vaccaro, Andrew M. Pilat, Logan Gusmano, Minh T. N. Pham, Daniel Barich, Audrey Gibson, Mwï Epalle, Dominick J. Frost, Elianajoy Volin, Zachary C. Slimak, Chelsea C. Menke, M. Siobhan Fennessy, Joan L. Slonczewski

## Abstract

Microbial communities of small freshwater bodies are poorly understood. Four ponds in Knox County, Ohio, were sampled over two years to investigate the relationship between the microbial taxa profiles, antibiotic resistance genes (ARGs), and environmental factors such as pH and tannin concentrations. For each site, microbial communities were collected by filtration and metagenomes were analyzed by short-read sequencing. Taxa profiles were predicted by the Kraken2/Bracken pipelines. Bacterial taxa with high abundance in these ponds included Betaproteobacteria (*Polynucleobacter* and *Methylopumilus*) and Actinobacteria (*Planktophila*, *Nanopelagicus*, and *Mycolicibacterium*). One pond, a former quarry with elevated pH, showed high prevalence of Cyanobacteria with a seasonal shift from *Synechococcus* to *Planktothrix* in the fall. *Planktothrix* increase was associated with acidification. ARGs were quantified using the ShortBRED pipeline to detect and quantify hits to a marker set derived from the Comprehensive Antibiotic Resistance Database (CARD). The top two ARGs with the largest marker hits encode components of a *Stenotrophomonas* drug efflux pump powered by proton-motive force (*smeABC*) and a mycobacterial global regulator that activates a drug pump and other cell defenses (*mtrA*). Pump function and global activation of transcription incur large energy expenditures, whose fitness cost may increase at high external pH where the cell’s proton-motive force is diminished. The *smeABC* and *mtrA* prevalence showed a modest correlation with acidifying conditions (low pH and high tannins) which contribute a large transmembrane pH difference to the proton-motive force, thus increasing the cell’s energy available for pump function and global gene expression.

**IMPORTANCE:** Compared to rivers and lakes, pond microbial ecosystems are understudied despite close contact with agriculture and recreation. Environmental microbes offer health benefits as well as hazards for human contact. Small water bodies may act as reservoirs for drug-resistant organisms and transfer of antibiotic resistance genes. Yet, the public is rarely aware of the potential for exposure to ARG-carrying organisms in recreational water bodies. Little is known about the capacity for freshwater microbial communities to remediate drug pollution and which biochemical factors may select against antibiotic resistance genes. This study analyzes the bacterial taxa composition and ARG prevalence including possible influence of factors such as pH and tannic acid levels.

## INTRODUCTION

Ponds are defined as small, shallow bodies of water with less than 30% emergent vegetation (1–3). Studies estimate that ponds comprise 30 percent of earth’s standing fresh water. Freshwater microbes play important roles as producers, consumers and scavengers (4), and they mediate production and consumption of greenhouse gases such as methane (5, 6).

Aquatic microbial communities are used as indicators of ecosystem quality (7). Yet the microbiomes of small water bodies are understudied when compared to lakes and rivers. Pond systems are complex and variable, showing a wide range of nutrient, pH, and mineral concentrations. Their small size means that they are more susceptible to eutrophication and changing pollutant levels from agricultural runoff. Their variable chemistry provides the opportunity of a “natural experiment” offering a range of characteristics to test which environmental variables influence the taxa profiles and prevalence of antibiotic resistance genes (ARGs). Chemical factors of interest include for example pH (8), tannin levels, nitrates and phosphates (9, 10).

Pond microbiomes are a potential source of antibiotic-resistant organisms that increasingly threaten global public health (11, 12). Antibiotic resistance genes (ARGs) in environmental microbial communities have the potential for transfer to pathogenic bacteria (13, 14). A modest level of antibiotic resistance is an ancient, widespread phenomenon naturally and historically occurring in all environments (15). But inputs of human origin can add substantially to the native ARG pool as well as add additional types of ARGs (16, 17). Yet, the public is rarely aware of the potential for exposure to ARG-carrying organisms in recreational water bodies such as rivers and ponds (18, 19). For example, in our region of Knox County, Ohio, previous studies report evidence of microcystins in local ponds (20) and drug-resistant *Acinetobacter baumannii* in a recreational river (21).

Little is known about the capacity for aquatic microbial communities to remediate pharmaceutical drug pollution and what chemical factors may select for or against ARGs (22). For example, a frontline mechanism of bacterial drug resistance is multidrug efflux pumps that confer resistance to multiple antibiotics (MDR). These efflux pumps are membrane protein complexes that export many substrates, including a wide array of antibiotics, metals, and metabolic byproducts from the cytoplasm (23, 24). Drug efflux spends energy, in most cases from the proton motive force (PMF) although some pump families use hydrolysis of ATP. pH plays an important role in powering MDR pumps because the transmembrane pH difference (ΔpH) is a component of PMF. At low external pH, the ΔpH is relatively large and can enhance contributions of efflux pumps (25). Pond water may be acidified by plant-derived polyphenols such as tannic acids (tannins). By contrast at high external pH (often a consequence of microbial photosynthesis) the proton-motive force is composed solely of electrical potential which is partly spent maintaining an inverted ΔpH (26, 27). Thus at high pH we predict bacteria will select against PMF-driven efflux pumps, and against large global regulons involving energy-expensive gene expression.

Our study was designed to reveal the prevalence patterns of microbial taxa and ARGs in pond water microbiomes, including the roles of pH and other chemical factors. We took water samples from four ponds on or near agricultural land in Knox County, Ohio (**Figure 1**). Burtnett pond is a restored wetland surrounded by adjacent agricultural fields. Ariel-Foundation Park Lake 2 (Foundation Pond) was formerly a silica gravel quarry. McManis pond receives fecal input from livestock. Porter pond is surrounded by pine and oak trees.

**Figure 1.**
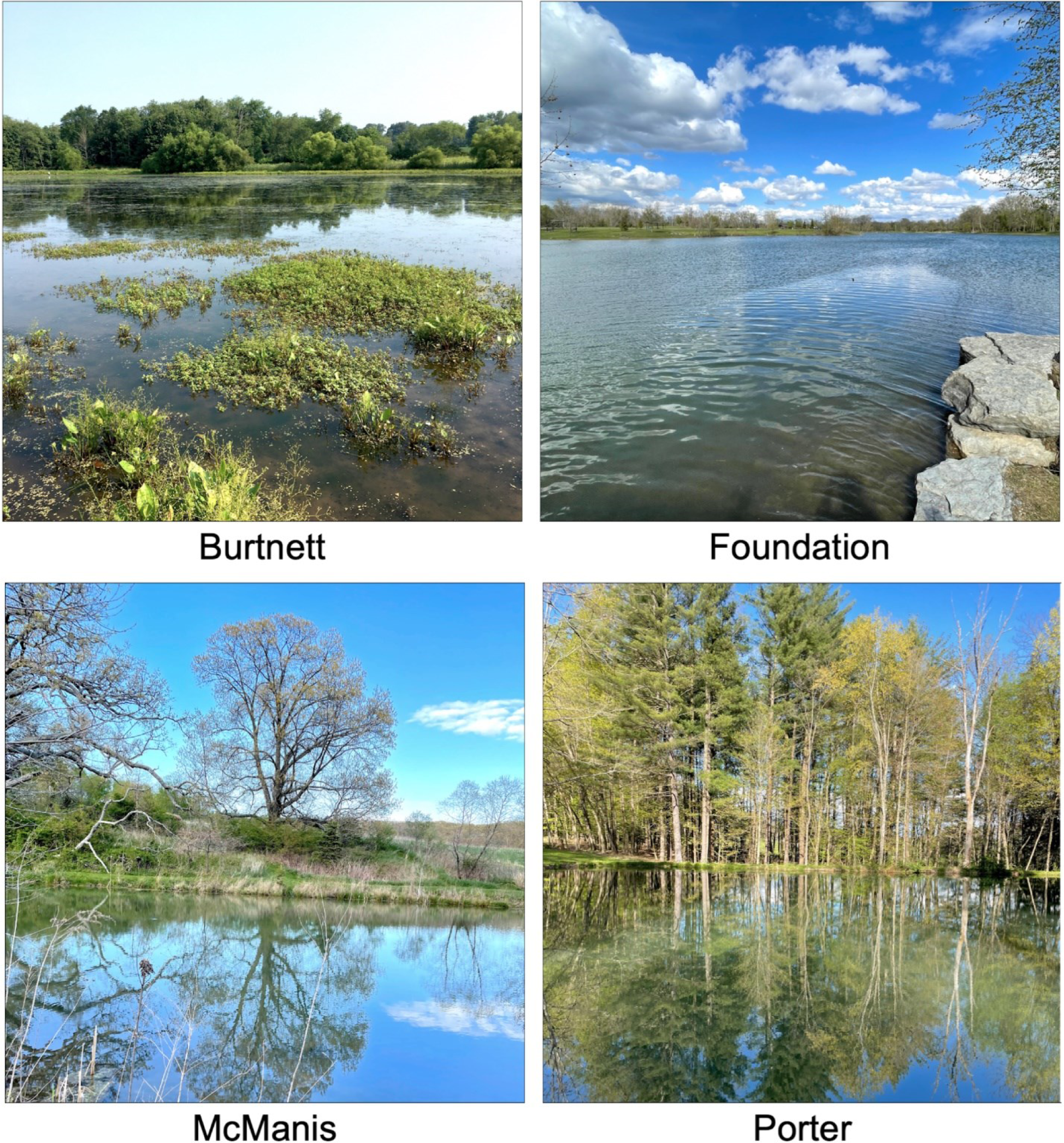
Ponds sampled during 2021 and 2022. Each small water body is located in Knox County, Ohio.

We sampled water from these pond sites in two periods, the summer and fall of 2021, and in the fall of 2022; these collections were treated as distinct datasets. We surveyed the pond microbial metagenomes for taxa markers, using Kraken2/Bracken pipelines to classify marker gene hits and estimate relative abundance of taxa (28–30). We also quantified ARG abundance, using ShortBRED pipeline (31) to match marker sequences from the Comprehensive Antibiotic Resistance Database (CARD) (32). We characterized the relationships between pond chemistry, taxonomic profiles, and ARG prevalence with a particular focus on the role of pH and tannin concentration.

## RESULTS

### Ponds physical traits and chemistry

Water samples were obtained from the four ponds during two seasons, in 2021 and in 2022 (**Figure 1**; see Methods). All metadata are presented in **Supplemental Tables S1, S2.** The ponds varied in their water chemistry, including pH, tannin concentration and phosphate levels (**Figure 2**). Burtnett pond had low pH values from September through November (pH 5.9 – 6.3) with generally higher levels of tannins, dissolved organic carbon (DOC) and phosphate than the other ponds. By contrast, Foundation pond maintained higher pH, always above 7 (pH 7.1 – 8.9) with relatively low levels of phosphate and ammonia. In Foundation pond, the conductivity (150 – 670 μS/cm) was consistently higher than in the other three ponds (100 – 300 μS/cm), suggesting slightly higher salinity. McManis and Porter ponds had variable levels of pH, with low levels of tannins, phosphate and ammonia.

**Figure 2.**
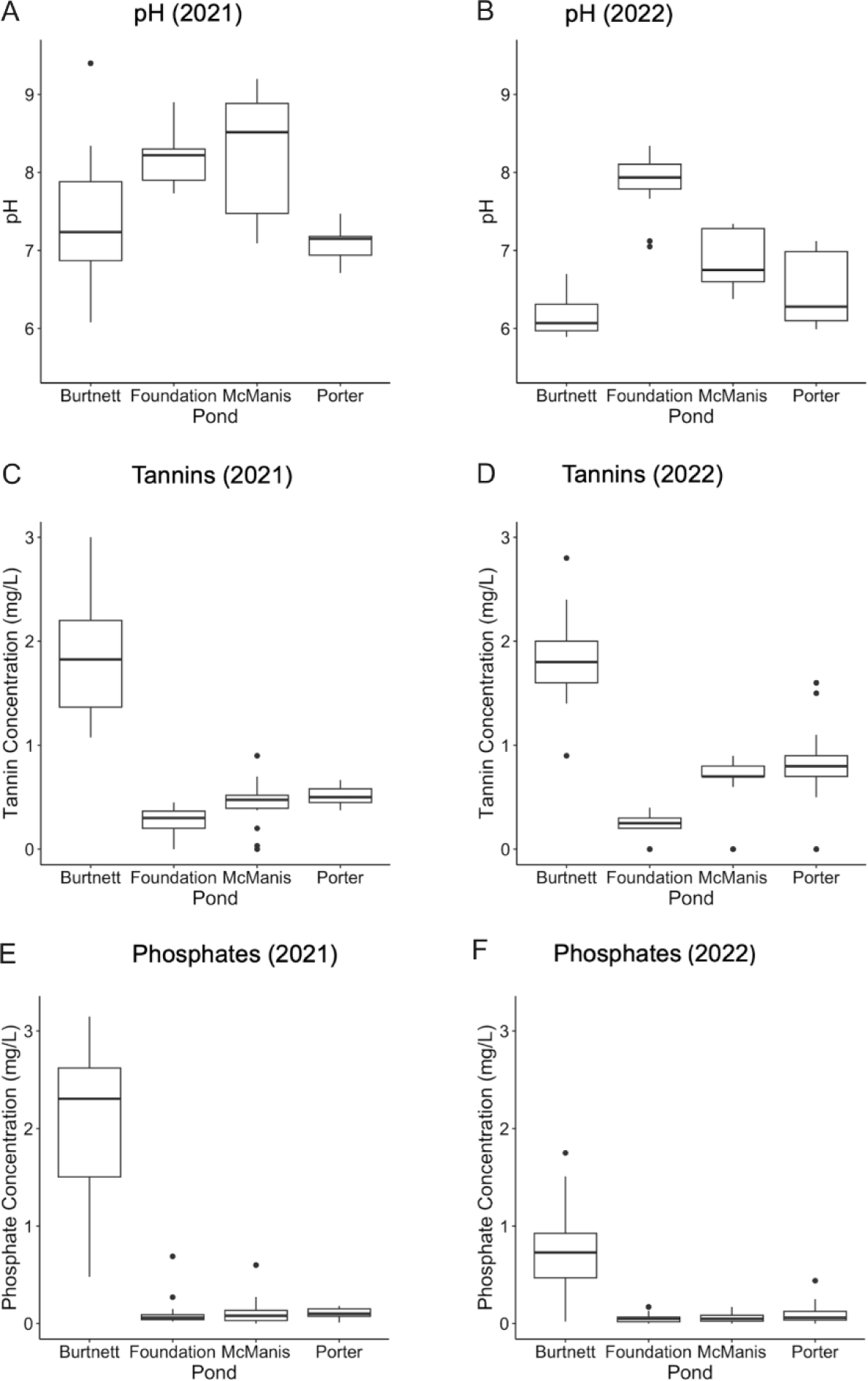
pH, tannin concentration (mg/L) and phosphate concentration (mg/L) in each pond were compiled across each sampling year, 2021 and 2022.

Tannic acids dissociate with a pKa of approximately 6 and thus can lower the pH of freshwater (33). In our data we saw strong correlations between pH and tannin levels (Spearman correlation R = -0.46, P < 0.001, for 2021; R = -0.73, P < 0.001, for 2022). The pH and tannin levels are plotted in **Figure 3**. The relationship appears especially strong for Burtnett pond, where tannin levels were consistently high and pH was low, especially in the fall months.

**Figure 3.**
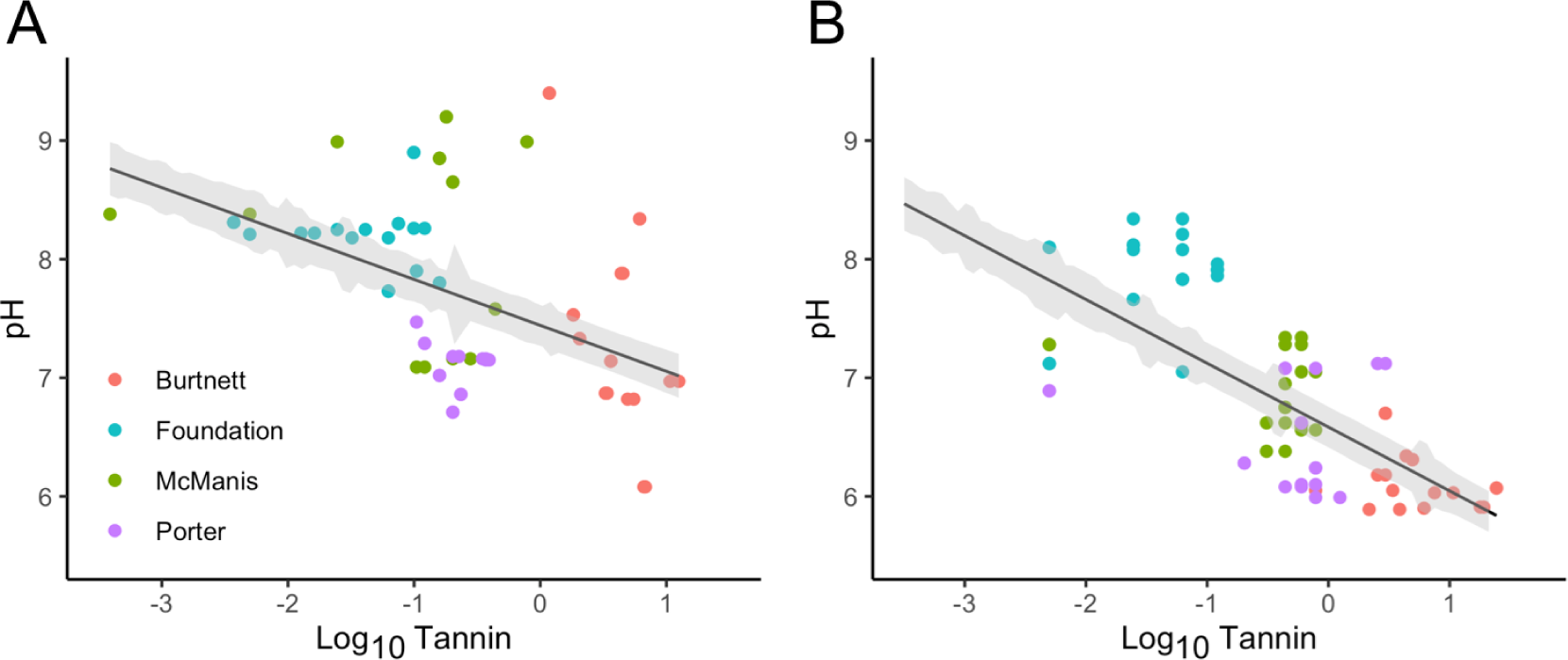
**Plots of pH against log10 tannin concentration**. Symbols indicate: red (Burtnett Pond), cyan (Foundation Pond), green (McManis Pond), purple (Porter Pond).

### All ponds showed high prevalence of Actinobacteria and Betaproteobacteria, with high Cyanobacteria in Foundation pond

To predict the relative abundance of microbial taxa in each sample, we applied the Kraken2/Bracken pipeline (28) to our 150-bp read metagenomes (**Figure 4**). Kraken2 assigns taxa to short sequences by k-mer alignment to a reference database; Bracken then uses taxa assignments to estimate relative abundance of taxa in the sample. Using these pipelines, we obtained consistent estimates of bacterial taxa, though not for archaea or eukaryotes, whose marker representation was limited in the database (see Methods).

**Figure 4.**
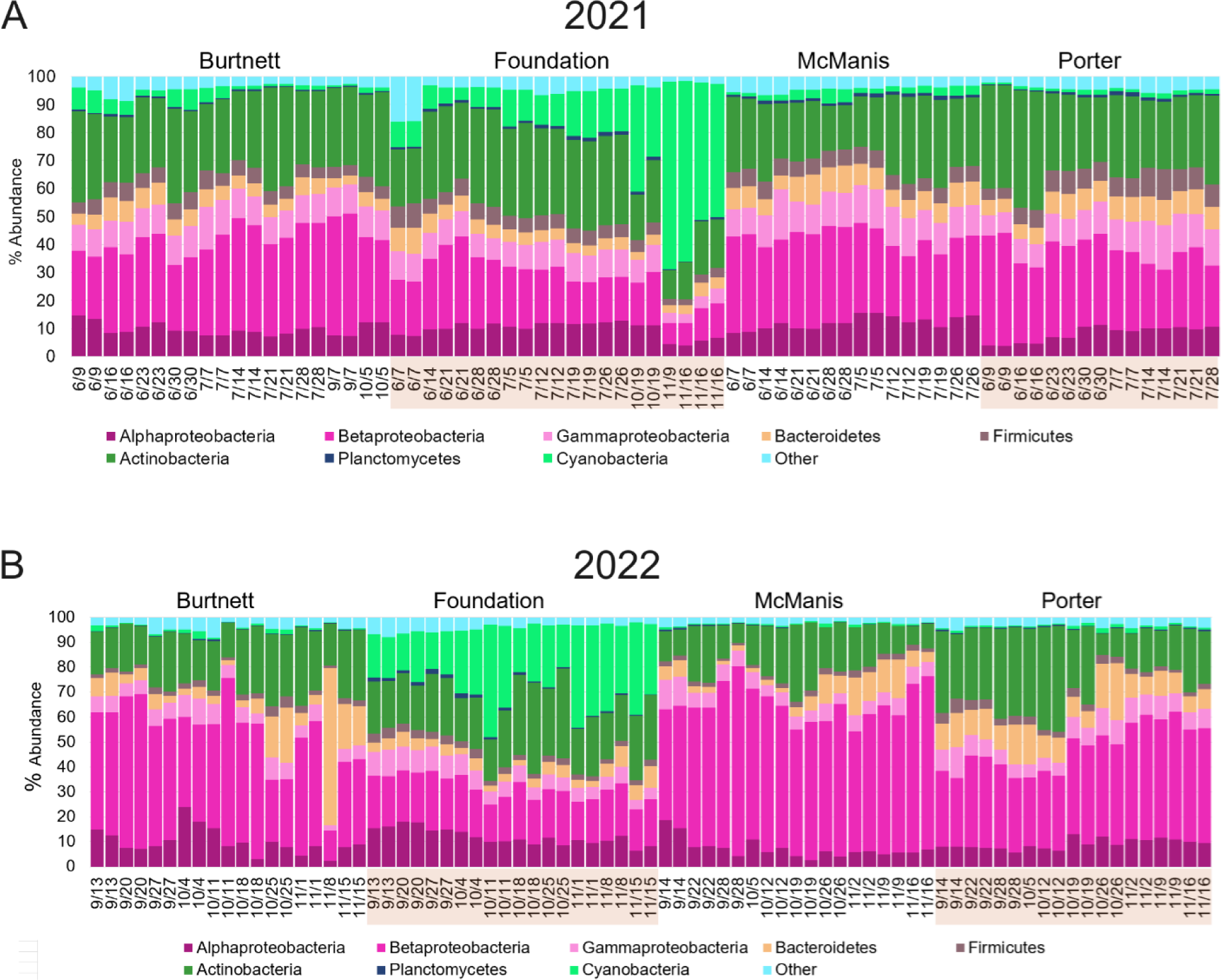
Taxa abundance: major bacterial phyla and classes predicted by Kraken2/Bracken in each pond across sampling dates. Taxa that make up less than one percent of the overall sample are grouped together and labeled as “Other.” **A,** samples from 2021; **B**, samples from 2022.

In both the 2021 and 2022 metagenomes, the taxa Actinobacteria and Betaproteobacteria showed high abundance across all ponds and seasons (**Figure 4**). At the genus level (**Figure 5**) the major predicted Actinobacteria were *Planktophila* and *Nanopelagicus*. These genera are chemoheterotrophs that are commonly found in freshwater communities (34, 35). Major Betaproteobacteria predicted were *Polynucleobacter* and *Limnohabitans* (36, 37) and *Methylopumilus* (38). These organisms are typical of high-quality freshwater systems.

**Figure 5.**
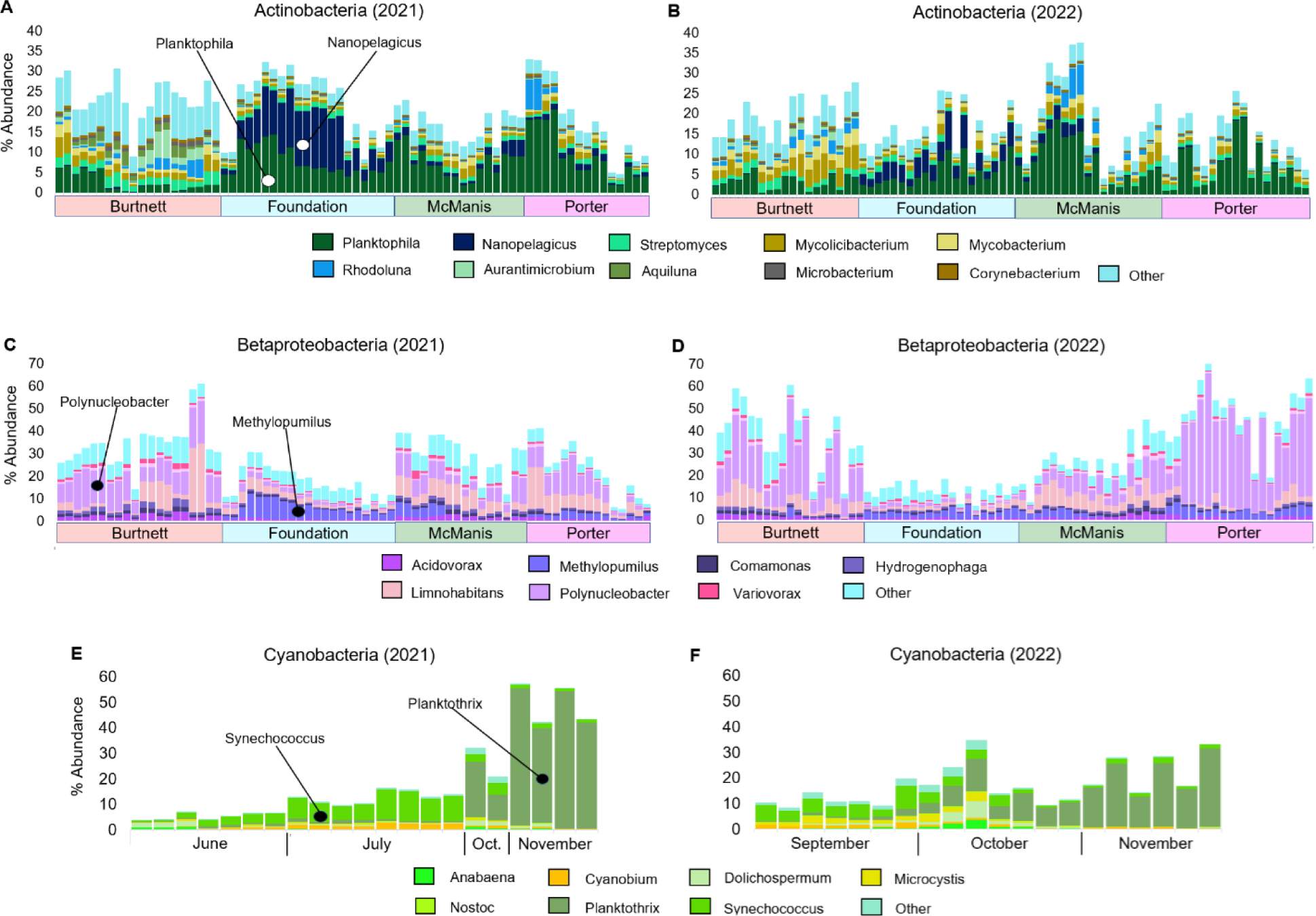
Actinobacteria, Betaproteobacteria, and Cyanobacteria. Genera predicted by Kraken2/Bracken. Genera predicted for: **A**, Actinobacteria in 2021; **B**, Actinobacteria in 2022; **C**, Betaproteobacteria in 2021; **D**, Betaproteobacteria in 2022**; E**, Cyanobacteria in Foundation Pond, 2021; **F**, Cyanobacteria in Foundation Pond, 2022.

*Polynucleobacter* species commonly feed on algal products (37). *Methylopumilus* and related methylotrophs oxidize single-carbon compounds such as methylamine, an important niche in water and soil environmental ecosystems (38).

Foundation pond showed taxa patterns that were distinct from those of the other three ponds. Cyanobacteria were highly abundant in the fall months (September through November) (**Figure 4**). The 2021 dataset showed a steady increase in proportion of Cyanobacteria across the summer and fall months. The genera of Cyanobacteria shifted from the picocyanobacteria *Synechococcus* in the summer to the filamentous *Planktothrix* in the fall. *Synechococcus* species are important oxygenic phototrophs (39) although blooms may produce toxins (40) Previously, cyanobacteria are reported to shift from *Synechococcus* in the spring to filamentous species in the fall (41) *Planktothrix* can also contribute to toxin-producing blooms (42, 43). Besides phototrophs, Foundation samples in the summer months of 2021 showed distinctive genera of Actinobacteria (*Nanopelagicus*) and of Betaproteobacteria (*Methylopumilus*). Thus, Foundation pond appeared to show succession of prevalent taxa from *Nanopelagicus*, *Methylopumilus* and *Synechococcus* in the summer to *Planktothrix* in the fall. The *Planktothrix* abundance showed a negative correlation with pH; that is, increased levels with acidity (Spearman R = -0.76, P < 0.01). No effect was seen of phosphate level (R = -0.05, P = 0.74), which is previously associated with *Planktothrix* blooms (44).

### Ponds showed distinctive patterns of ARG abundance

Environmental bacteria possess a wide range of defenses against antibiotics, including broad-substrate efflux transporters as well as highly specific drug pumps and enzymes that modify the drug or its substrate (11, 15). The genes encoding many of these are compiled in the CARD database (32). We detected ARGs from CARD in our metagenomes using a marker set constructed by the ShortBRED pipeline (31) (Table S3). ShortBRED converts the CARD database into homologous protein families and searches for unique motifs to use as unique and selective “markers” for each protein family. The unique protein motifs are then screened against all the proteins in the universal UNIPROT database that are not listed in CARD. Nonspecific markers are thus eliminated. All markers in Table S3 were derived from the CARD database accessed 06/14/2021 (see Methods). Because of the UniProt sequence filtration, we cannot assume that all possible ARGs would be found; but those identified are highly specific.

The read hits for each ARG were compiled from all samples for each year, 2021 and 2022. The number of FARG hits for each pond, and the hits summed for all ponds, are shown for the top 30 ranked ARGs each year (**Figure 6**). In both years 2021 and 2022 the three top-ranked ARGs were *smeB*, *mtrA*, and OXA-156. The *smeB* gene encodes a subunit of the SmeABC multidrug efflux pump of the *Stenotrophomonas maltophilia*, an opportunistic pathogen (45, 46).

**Figure 6.**
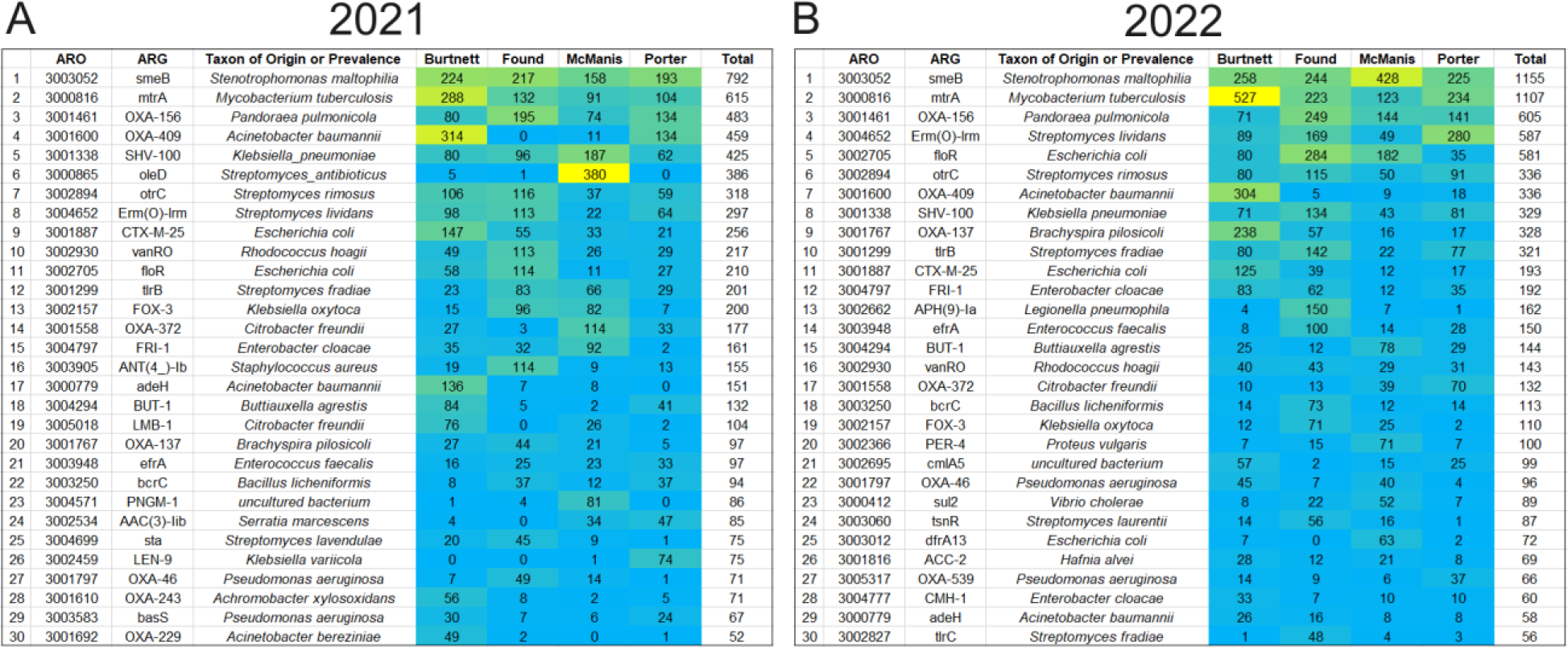
Antibiotic resistance genes (ARGs) predicted by ShortBRED markers. Relative abundance of top 30 ARG marker hits is shown. Read hit numbers are ranked in descending order by total hits across all samples. Heat map shows relative abundance of ARG hits in each pond sampling location. Yellow, high values; cyan, low values.

*S. maltophilia* with *smeB* has been found as a contaminant of drinking water (47); it is a member of a genus widespread in environmental sources such as soil and plants (48). The *mtrA* gene is a global response regulator found in environmental species of *Mycobacterium* and *Corynebacterium* as well as *Mycobacterium tuberculosis* (49, 50). The MtrAB two-component regulatory system governs cell morphology; its loss increases sensitivity to vancomycin and rifampicin but decreases sensitivity to isoniazid. In *Streptomyces* spp., MtrA regulates production of antibiotics (51). The third-ranked ARG in our ponds, OXA-156, encodes a beta-lactamase, one of many found in environmental microbiomes (52, 53).

Other ARGs showed variable ranking depending on the pond. The beta-lactamase OXA- 409 was highly prevalent in Burtnett pond and absent or barely present in other ponds. The macrolide glycotransferase gene *oleD* showed high prevalence in McManis pond in 2021 but did not show up in 2022. Such findings would be expected for the patchy nature of pond water systems with high amounts of suspended particulates.

### Abundance of certain ARGs showed correlations with acidity and tannin concentration

We tested the hypothesis that ARG abundance in the pond microbial communities is associated with various chemical factors such as pH and tannin concentration. Using ShortBRED, we counted ARG marker hits from all sample dates; for nearly all dates, two independent samples were analyzed. **Figure 6** lists the top 30 ranked ARGs from the total of all samples, for 2021 and separately for 2022. For meaningful comparison, we compiled a list of ARGs that were found in the top 30 ranked ARGs for both 2021 and 2022. This list of top-ranked ARGs shared by both years (20 in all) were tested using Spearman correlations between marker hit numbers and measures of pond chemistry (**Figure 7**; P-values in Table S4). Ten of the twenty top-ranked ARGs (*smeB*, *mtrA*, OXA-409, *otrC*, *Erm(O)-lrm*, *CX-M-25*, *floR*, OXA-372, *adeH*, BUT-1) showed a positive correlation with acidity and tannin concentration. Of these ARGs, *smeB*, *mtrA*, OXA-409, CTX-M-25, and OXA-372 also showed positive correlation with acidity and tannins in 2022. The differences between the two years may be associated with the high variance of many pond water factors and with the seasonal difference of the sample ranges (June through November of 2021, September through November of 2022). For example, the summer season of samples in 2021 showed generally higher pH values (**Figures 3 and 8**).

**Figure 7.**
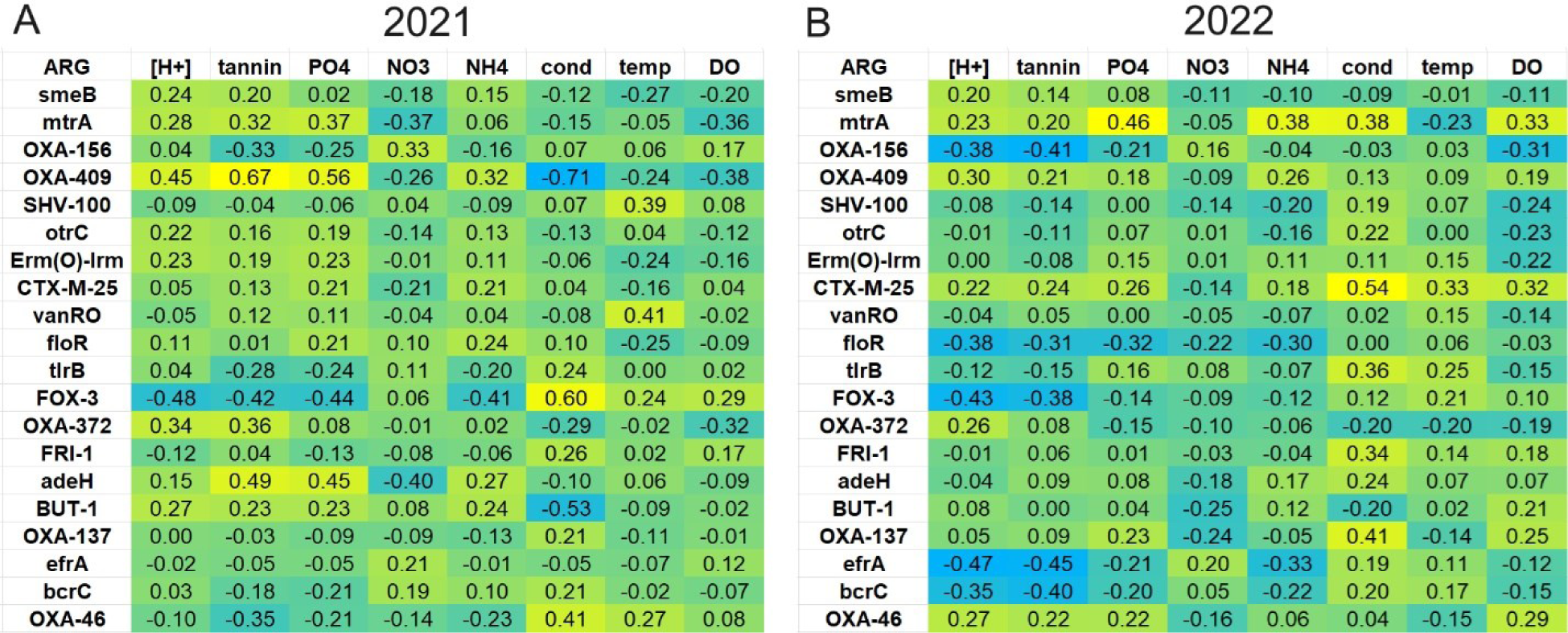
Heat map of Spearman correlations between top-ranked ARGs and chemical measures. The 20 ARGs shown are those that appeared in both 2021 and 2022 among the top 30 ranked markers for each year. Cell color indicates the range of Spearman R values from positive (yellow) to negative (cyan). Chemical and physical measures include acidity, tannin concentration (mg/L), phosphate concentration (mg/L), nitrate concentration (mg/L), ammonia concentration (mg/L) conductivity (μS/cm), temperature (°C), and dissolved oxygen (mg/L). P-values are provided in Supplemental Table S4.

**Figure 8.**
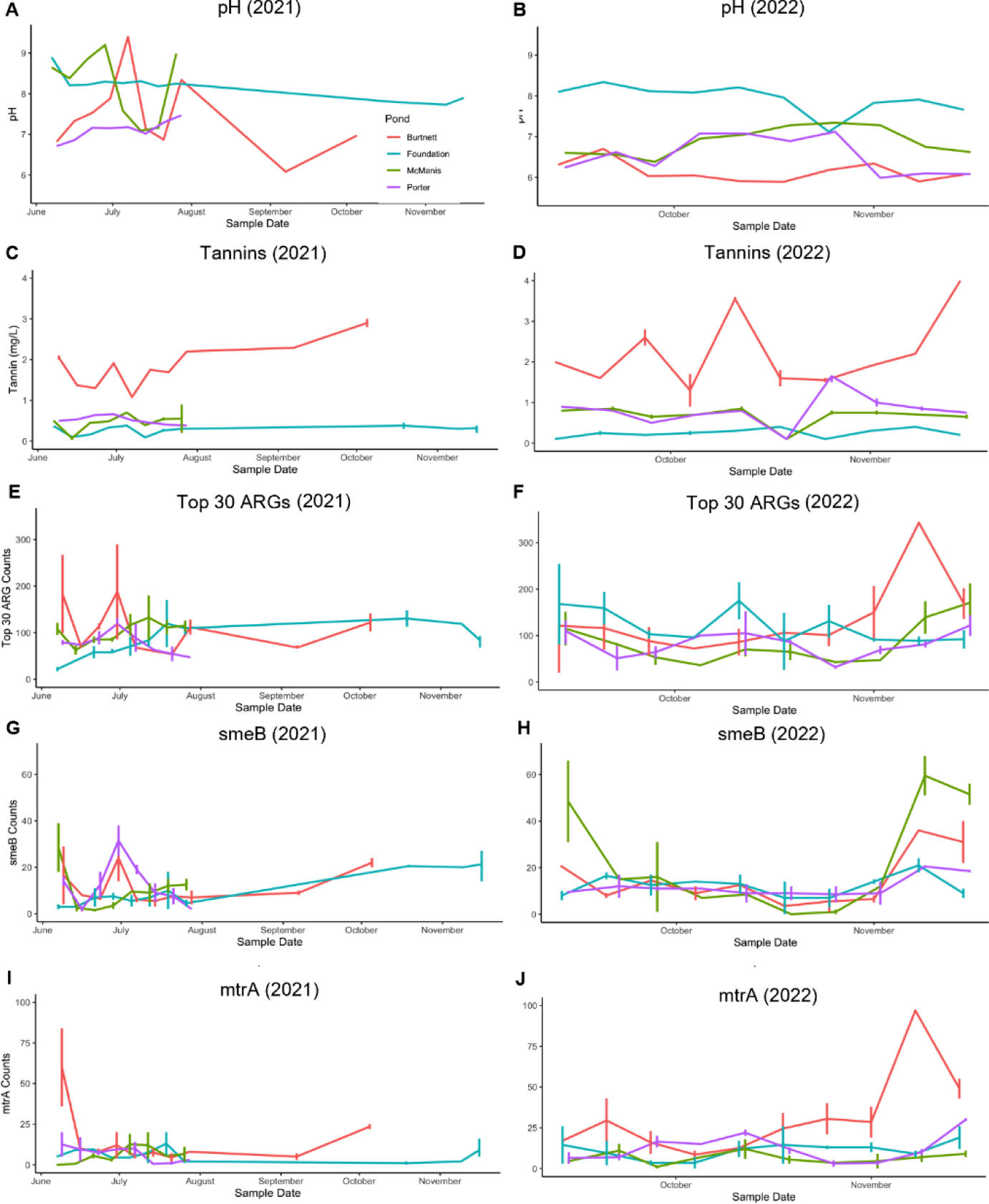
Time course of pond water measurement of pH and tannin levels; and of ARG counts for sum of top 30 ranked ARGs, *smeB*, and *mtrA*. Line colors: orange, Burtnett; cyan, Foundation; green, McManis; violet, Porter.

To investigate the roles of pH and tannins in ARG abundance, we separated the ARG counts among the four ponds. We then plotted pond ARG abundance over the course of time, in parallel with the time course of pH levels and tannin concentrations (**Figure 8**). During the fall of 2022, three of the four ponds had pH values generally below pH 7, and in Burtnett pond the tannins reached a high concentration. In Burtnett, the same period in the fall saw an increase in levels of *smeB*, *mtrA*, and the top 30 ranked ARGs. The Burtnett *smeB* and *mtrA* counts showed weak correlations with pH (*smeB*: R = -0.18, P = 0.27; *mtrA*: R = -0.39, P = 0.02) and strong correlations with tannin concentration (*smeB*: R = 0.55, P < 0.001; *mtrA*: R = 0.33, P = 0.04). These correlations suggest a connection between acidity (conferring high proton motive force) and drug resistance mechanisms that are energy-expensive.

Tannins are a component of overall dissolved organic carbon (DOC) which might affect ARGs by providing substrate that increases respiration (54, 55). During the 2022 sampling, dissolved organic carbon (DOC) was measured for alternate samples. DOC levels were generally higher in Burtnett pond than in the other three ponds (**Supplemental Table S5**). DOC levels showed a significant correlation with pH overall (R = -0.64, P < 0.001) but not with pH in Burtnett pond, the source of most pH variation (R = -0.13, P = 0.74). Only OXA-409 and OXA- 46 showed positive correlations with DOC.

Phosphate was also investigated as a factor in ARG abundance. Two of the highly ranked ARGs, *mtrA* and *OXA-409*, showed positive correlation with phosphate levels in both 2021 and 2022 (**Figure 7**). These two ARGs were found predominantly in Burtnett pond, where phosphate was high (**Figure 6**). Nevertheless, within the Burtnett dataset combining 2021 and 2022, the two ARGs showed no positive correlation with phosphate level (*mtrA*: R = -0.44, p = 0.005; OXA-409: R = 0.25, p = 0.12).

## DISCUSSION

For our investigation of microbial community composition and ARG abundance, the four ponds offered a good natural experiment in their shared location, with similar geology and agricultural proximity, while showing some variety in chemical and physical factors (**Figures 2 and 8**). The Burtnett pond had significantly higher levels of tannic acids and phosphate than the other three ponds, whereas Foundation pond had higher pH values. In addition, Foundation pond had higher conductivity values (**Tables S1, S2**) which may indicate higher salt content. The ponds also showed interesting variation over the seasons. Burtnett showed a rise in tannins during the fall months during tree leaf senescence and leaf inputs to the pond (September through November) whereas McManis and Porter ponds had a decline in pH during October through November. In general, the late fall was associated with acidification, likely associated with influx of tannic acids from fallen leaves.

The microbiomes of all four ponds had high proportions of Actinobacteria and Betaproteobacteria (**Figure 4**) which commonly occur in high-quality environments with minimal human disruption (34, 37). Within these categories the ponds showed interesting differences in genera; Foundation had a higher proportion of *Nanopelagicus* species, whereas McManis had a higher amount of *Planktophila*. The presence of the Betaproteobacterium, *Methylopumilus*, shows a capacity for turnover of reduced one-carbon compounds such as methanol and methylamine, an important role in aquatic carbon cycles (56). In Foundation, the succession of Cyanobacteria from *Synechococcus* in the summer to *Planktothrix* in the fall is important for relevance to oxygen production but also the potential for microcystin-producing blooms (41, 42). The increase of *Planktothrix* parallels a decrease in pH, a relationship not reported previously, although late bloom decomposition lowers the water pH (57) Our pond microbiomes showed many ARGs that are found in environmental taxa such as *Stentrophomonas*, *Mycobacterium*, *Pandorea*, *Streptomyces*, and *Acinetobacter* (**Figure 6**). While harmless for most people, many environmental microbes are now showing up as opportunistic pathogens of immune-compromised patients. This is concerning, for example, for *S. maltophilia* (46) and for non-tuberculosis *Mycobacterium* species (58), which are sources of the top two ARGs found in our samples (*smeB* and *mtrA* respectively).

The prevalence of ARGs in freshwater bodies is subject to numerous factors, most importantly the influx of human and agricultural sources of drug-resistant bacteria (13, 17). At the same time, the environmental communities that receive such inputs possess some resilience and ability to outcompete drug-resistant organisms. For example, our study of river ARGs associated with wastewater showed that the water downstream of the plant had largely restored its ARG counts to levels comparable to those upstream (21). Thus, it is of interest to consider which environmental factors might affect the resilience of water bodies receiving ARG inputs.

In our present study, we show modest evidence that abundance of certain ARGs depends upon water pH, with possible enhancement by acidification associated with tannin inputs. It would be informative to follow up this evidence with controlled experiments in microcosms, to assess the magnitude of the pH effect, and to determine whether it is mediated by energetics.

Energy cost is an important tradeoff of ARGs, so it would be good to know how this factor influences the environmental prevalence of drug resistant bacteria.

## METHODS

### Water sampling and metadata

Water was sampled from four ponds around Knox County, Ohio, a historically agricultural county with a scenic river and one small city (Mount Vernon, pop. 17,000) (**Figure 1**). While none of the four sampling sites was used directly in agriculture, each was located near agricultural land. Burtnett Pond (40°20’58” N, 82°19’31” W) was a restored wetland with a high abundance of duckweed and cattails and high traffic by geese and other waterfowl. It was surrounded by soybean fields and sheep pastures. The water had high concentrations of tannin and phosphate. Foundation Pond (40°23’10” N, 82°29’49” W) was a restored quarry for silica gravel formerly used by a glass factory. The pH was generally high (pH 7.1-9.0); the water was clear with a large population of geese and minimal input from surrounding vegetation. Porter Pond (40° 22’ 21.1188’’ N, 82° 24’ 56.97’’ W) had heavy input of leaf litter from oak, maple, and pine trees. McManis Pond (40°23’53” N, 82°24’24” W) generally had neutral to high pH (pH pH 6.7 – 9.0). Vegetation included cattails, weeping willows and duckweed; there was fecal input from poultry and goats.

Water microbiomes were obtained and analyzed using methods based on (21). Pond water was sampled weekly from June to July of 2021, and between September and November of 2021. A second dataset was collected from each pond weekly from September to November of 2022. All analysis was performed separately on each of the two datasets, using the same methods for both.

The values of pH, conductivity (μS/cm), temperature (°C) and dissolved oxygen (DO) concentration (mg/L) were measured in the field using a Hannah pH/conductivity combination meter and a YSI Pro20 DO meter (Yellow Springs Instruments, Yellow Springs, Ohio). Four 1- liter samples were collected from each site between 9:00-11:00 am and kept on ice in Whirl-Pak bags. Once in the lab, 100 ml of each sample was filtered using a sterile 0.22-micron Microfil V filter to collect microbial samples. During our sampling period in 2021, we switched to acid washing the outer plastic filter from the Microfil filters and using sterile 0.22 μm MF-Millipore® filters in the plastic casings. Filter samples were stored in sterile 2-ml microfuge tubes at -80 °C until processing. All colorimetric analysis was done within 48 h of sample collection using a Hach DR900 multiparameter portable colorimeter. Test ‘N Tube kits were used to measure orthophosphate, low range ammonia, low range nitrate, and tannin concentration in mg/L using the protocol outlined in the kits (Hach, Loveland, CO). Low range nitrate was analyzed using the cadmium reduction method, and tannin concentration was evaluated using the tyrosine method. Reusable glassware for colorimetry was placed in 10% HCl acid wash overnight and rinsed with deionized water between uses. For the 2022 dataset, dissolved organic carbon (DOC) was measured by combustion analysis (UC Davis Analytical Lab). 50-ml water samples were acidified to remove inorganic carbon, then injected into the high temperature combustion reactor with an oxidative catalyst. Following oxidation to completion, the CO2 was measured at 4.2 um by infrared detection.

### DNA isolation and sequencing

From each pond and sample date, two filter samples were processed for DNA sequencing. Metagenomic DNA was isolated using the ZymoBIOMICS DNA mini-prep kit, as described (21). For each set of preps, 75 μl of the ZymoBIOMICS microbial community standard was prepared under the same conditions to serve as a control. This mock community contained defined proportions of 10 microbes (5 Gram-positive bacteria, 3 Gram-negative bacteria, 2 fungal microbes).

Purity of DNA samples was determined using Nanodrop analysis (Thermo Fisher Scientific). Admera Health performed library construction with NexteraXT library kit (Illumina, San Diego, CA, USA) using Illumina® 8-nt dual-indices following the manufacturer’s recommendations. Amplifiable molar concentration of each library was measured by KAPA SYBR® FAST qPCR with QuantStudio ® 5 System (Applied Biosystems, California, USA).

Sequencing was performed on an Illumina® NovaSeq S4 (Illumina, California, USA) with a read length configuration of 150 PE for 40 M PE reads (20M in each direction) for each sample (Admera Health, New Jersey, USA).

### Taxa profiles analysis

The DNA sequence FASTQ files were trimmed using Trimmomatic to remove Nextera adapters and poor-quality sequences (59). From each filtered sequence, we used Kraken2 with the RefSeq reference database to identify the sequences in our pond samples.

Kraken2 assigns identity to the sequences in samples of interest by matching k-mers in the sample of interest to k-mers of the lowest common ancestor (LCA) in a reference database (28). The reference database used was the Kraken2 Standard Refseq Collection containing archaea, bacteria, viral, plasmid and human sequences, dated June 2023 (https://benlangmead.github.io/aws-indexes/k2 - accessed 09/13/2023). We omitted “human” sequences results because the values appeared inconsistent and increased the variability of overall abundance predictions. We used Bracken (Bayesian Re-estimation of Abundance with Kraken2) to compute the relative abundance of identified organisms in our samples (28–30).

### ARG marker analysis

Metagenomic short reads were matched to ARG markers generated by ShortBRED-Identify (31) using the updated CARD 3.1.2 database of ARGs (32) (https://card.mcmaster.ca - accessed 06/14/2021). ShortBRED-Identify was run with true markers (minimum length 8 amino acid residues) filtered against the reference database UniRef90 (https://www.uniprot.org/uniref/ - UniProt release 2021_03 on 06/01/21). The marker list is presented in **Supplemental Table S3**. Marker sets were evaluated by the compositions of fully specific, partly specific, and best non-specific markers they contained, numbers of gene families with fully specific markers, appearance of markers with abnormally high hits, and non-specific blastp hits.

The marker set included markers for one ARG, *mtrA* (ARO_3000816) which has since been reassigned to an unrelated gene by the CARD curation. Nevertheless, our inspection confirms that our marker sequences do match sequences of the *Mycobacterium* gene *mtrA* (50). The marker sequences labeled *mtrA* were: MDTMRQRILVVDDDASLAEMLTIVLRGEGFDTAV, VIGDGTQALTAVRELRPDLVLLDLMLPGMNGIDV, VCRVLRADSGVPIVMLTAKTDTVDVVLGLESGAD.

To match the target sequences, we ran ShortBRED-Quantify (31) against the trimmed FASTQ files for each sample. The count of target matches for each ARG (counts per ARG) was obtained for each sequenced sample.

### Statistical analysis

All statistical analysis was performed using R statistical software (v4.1.2; R Core Team 2021). Spearman correlations by Rfit (60). Correlations were performed within each pond, and across all four ponds. Heat maps of each correlation table were made with blue representing lowest values and yellow representing highest values.

## ACKNOWLEDGEMENTS

This project was funded by National Science Foundation grant MCB-1923077 and by the Kenyon Philip and Sheila Jordan Fund. We thank the property owners of the three private ponds, and the manager of Ariel Foundation Park, for their gracious permission to obtain water samples.

## REFERENCES

1. Richardson DC, Holgerson MA, Farragher MJ, Hoffman KK, King KBS, Alfonso MB, Andersen MR, Cheruveil KS, Coleman KA, Farruggia MJ, Fernandez RL, Hondula KL, López Moreira Mazacotte GA, Paul K, Peierls BL, Rabaey JS, Sadro S, Sánchez ML, Smyth RL, Sweetman JN. 2022. A functional definition to distinguish ponds from lakes and wetlands. Sci Rep 12.

2. Chopyk J, Allard S, Nasko DJ, Bui A, Mongodin EF, Sapkota AR. 2018. Agricultural freshwater pond supports diverse and dynamic bacterial and viral populations. Front Microbiol 9.

3. Harper LR, Buxton AS, Rees HC, Bruce K, Brys R, Halfmaerten D, Read DS, Watson H V., Sayer CD, Jones EP, Priestley V, Mächler E, Múrria C, Garcés-Pastor S, Medupin C, Burgess K, Benson G, Boonham N, Griffiths RA, Lawson Handley L, Hänfling B. 2019. Prospects and challenges of environmental DNA (eDNA) monitoring in freshwater ponds. Hydrobiologia. Springer International Publishing 10.1007/s10750-018-3750-5.

4. Boyd CE. 2020. Microorganisms and Water Quality, p. 233–267. In Water Quality. Springer International Publishing.

5. Perez-Coronel E, Michael Beman J. 2022. Multiple sources of aerobic methane production in aquatic ecosystems include bacterial photosynthesis. Nat Commun 13.

6. Reis PCJ, Tsuji JM, Weiblen C, Schiff SL, Scott M, Stein LY, Neufeld JD. 2024. Enigmatic persistence of aerobic methanotrophs in oxygen-limiting freshwater habitats. ISME J 10.1093/ismejo/wrae041.

7. Sagova-Mareckova M, Boenigk J, Bouchez A, Cermakova K, Chonova T, Cordier T, Eisendle U, Elersek T, Fazi S, Fleituch T, Frühe L, Gajdosova M, Graupner N, Haegerbaeumer A, Kelly AM, Kopecky J, Leese F, Nõges P, Orlic S, Panksep K, Pawlowski J, Petrusek A, Piggott JJ, Rusch JC, Salis R, Schenk J, Simek K, Stovicek A, Strand DA, Vasquez MI, Vrålstad T, Zlatkovic S, Zupancic M, Stoeck T. 2021. Expanding ecological assessment by integrating microorganisms into routine freshwater biomonitoring. Water Res. Elsevier Ltd 10.1016/j.watres.2020.116767.

8. Pu H, Yuan Y, Qin L, Liu X. 2023. pH Drives Differences in Bacterial Community β- Diversity in Hydrologically Connected Lake Sediments. Microorganisms 11:676.

9. Wu Y, Wen Y, Zhou J, Wu Y. 2014. Phosphorus release from lake sediments: Effects of pH, temperature and dissolved oxygen. KSCE Journal of Civil Engineering 18:323–329.

10. Ni X, Yuan Y, Liu W. 2020. Impact factors and mechanisms of dissolved reactive phosphorus (DRP) losses from agricultural fields: A review and synthesis study in the Lake Erie basin. Science of the Total Environment 714.

11. Bengtsson-Palme J, Kristiansson E, Larsson DGJ. 2018. Environmental factors influencing the development and spread of antibiotic resistance. FEMS Microbiol Rev 42:68–80.

12. Zhang Q, Zhang Z, Lu T, Peijnenburg WJGM, Gillings M, Yang X, Chen J, Penuelas J, Zhu YG, Zhou NY, Su J, Qian H. 2020. Cyanobacterial blooms contribute to the diversity of antibiotic-resistance genes in aquatic ecosystems. Commun Biol 3:1–10.

13. M Pärnänen KM, Narciso-da-Rocha C, Kneis D, Berendonk TU, Cacace D, Thuy Do T, Elpers C, Fatta-Kassinos D, Henriques I, Jaeger T, Karkman A, Luis Martinez J, Michael SG, Michael-Kordatou I, Rodriguez-Mozaz S, Schwartz T, Sheng H, Sørum H, Stedtfeld RD, Tiedje JM, Varela Della Giustina S, Walsh F, Vaz-Moreira I, Virta M, Manaia CM. 2019. H E A L T H A N D M E D I C I N E Antibiotic resistance in European wastewater treatment plants mirrors the pattern of clinical antibiotic resistance prevalence.

14. Nnadozie CF, Odume ON. 2019. Freshwater environments as reservoirs of antibiotic resistant bacteria and their role in the dissemination of antibiotic resistance genes. Environmental Pollution. Elsevier Ltd 10.1016/j.envpol.2019.113067.

15. D’Costa VM, King CE, Kalan L, Morar M, Sung WWL, Schwarz C, Froese D, Zazula G, Calmels F, Debruyne R, Golding GB, Poinar HN, Wright GD. 2011. Antibiotic resistance is ancient. Nature 477:457–461.

16. Jara D, Bello-Toledo H, Domínguez M, Cigarroa C, Fernández P, Vergara L, Quezada-Aguiluz M, Opazo-Capurro A, Lima CA, González-Rocha G. 2020. Antibiotic resistance in bacterial isolates from freshwater samples in Fildes Peninsula, King George Island, Antarctica. Sci Rep 10:3145.

17. Hultman J, Tamminen M, Pärnänen K, Cairns J, Karkman A, Virta M. 2018. Host range of antibiotic resistance genes in wastewater treatment plant influent and effluent. FEMS Microbiol Ecol 94.

18. Nappier SP, Liguori K, Ichida AM, Stewart JR, Jones KR. 2020. Antibiotic resistance in recreational waters: State of the science. Int J Environ Res Public Health. MDPI AG 10.3390/ijerph17218034.

19. Rajasekar A, Murava RT, Norgbey E, Vadde KK, Qiu M, Guo S, Yu T, Wang R, Zhao C. 2023. Distribution of Antibiotic Resistance Genes and Their Associations with Bacterial Communities and Water Quality in Freshwater Lakes. Water Air Soil Pollut 234.

20. Mrdjen I, Fennessy S, Schaal A, Dennis R, Slonczewski JL, Lee S, Lee J. 2018. Tile Drainage and Anthropogenic Land Use Contribute to Harmful Algal Blooms and Microbiota Shifts in Inland Water Bodies. Environ Sci Technol 52:8215–8223.

21. Murphy A, Barich D, Fennessy MS, Slonczewski JL. 2021. An Ohio State Scenic River Shows Elevated Antibiotic Resistance Genes, Including Acinetobacter Tetracycline and Macrolide Resistance, Downstream of Wastewater Treatment Plant Effluent. Microbiol Spectr 9:e00941.

22. Yang Y, Song W, Lin H, Wang W, Du L, Xing W. 2018. Antibiotics and antibiotic resistance genes in global lakes: A review and meta-analysis. Environ Int. Elsevier Ltd 10.1016/j.envint.2018.04.011.

23. Alvarez-Ortega C, Olivares J, Martínez JL. 2013. RND multidrug efflux pumps: What are they good for? Front Microbiol. Frontiers Research Foundation 10.3389/fmicb.2013.00007.

24. Du D, Wang-Kan X, Neuberger A, van Veen HW, Pos KM, Piddock LJV, Luisi BF. 2018. Multidrug efflux pumps: Structure, function and regulation. Nat Rev Microbiol 16:523– 539.

25. Liu Y, Van Horn AM, Pham MTN, Dinh BNN, Chen R, Raphael SDR, Paulino A, Thaker K, Somadder A, Frost DJ, Menke CC, Slimak ZC, Slonczewski JL. 2024. Fitness trade- offsof multidrug effluxpumps in Escherichia coli K-12 in acid or base, and with aromatic phytochemicals. Appl Environ Microbiol 90.

26. Slonczewski JL, Rosen BP, Alger JR, Macnab RM. 1981. pH homeostasis in *Escherichia coli*: Measurement by P^31^ nuclear magnetic resonance of methylphosphonate and phosphate. Proceedings of the National Academy of Sciences 78:6271–6275.

27. Slonczewski JL, Fujisawa M, Dopson M, Krulwich TA. 2009. Cytoplasmic pH measurement and homeostasis in bacteria and archaea., p. 1–79, 317. In Advances in Microbial Physiology.

28. Wood DE, Salzberg SL. 2014. Kraken: Ultrafast metagenomic sequence classification using exact alignments. Genome Biol 15.

29. Lu J, Breitwieser FP, Thielen P, Salzberg SL. 2017. Bracken: Estimating species abundance in metagenomics data. PeerJ Comput Sci 2017:1–17.

30. Lu J, Rincon N, Wood DE, Breitwieser FP, Pockrandt C, Langmead B, Salzberg SL, Steinegger M. 2022. Metagenome analysis using the Kraken software suite. Nat Protoc 17:2815–2839.

31. Kaminski J, Gibson MK, Franzosa EA, Segata N, Dantas G, Huttenhower C. 2015. High-specificity targeted functional profiling in microbial communities with ShortBRED. PLoS Comput Biol 11.

32. Alcock BP, Raphenya AR, Lau TTY, Tsang KK, Bouchard M, Edalatmand A, Huynh W, Nguyen AL V., Cheng AA, Liu S, Min SY, Miroshnichenko A, Tran HK, Werfalli RE, Nasir JA, Oloni M, Speicher DJ, Florescu A, Singh B, Faltyn M, Hernandez-Koutoucheva A, Sharma AN, Bordeleau E, Pawlowski AC, Zubyk HL, Dooley D, Griffiths E, Maguire F, Winsor GL, Beiko RG, Brinkman FSL, Hsiao WWL, Domselaar G V., McArthur AG. 2020. CARD 2020: antibiotic resistome surveillance with the comprehensive antibiotic resistance database. Nucleic Acids Res 48:D517–D525.

33. Boyd CE. 2020. Acidity, p. 215–231. In Water Quality. Springer International Publishing, Cham.

34. Neuenschwander SM, Ghai R, Pernthaler J, Salcher MM. 2018. Microdiversification in genome-streamlined ubiquitous freshwater Actinobacteria. ISME Journal 12:185–198.

35. Lipko IA. 2020. Phylogeny of the freshwater lineages within the phyla Actinobacteria (Overview). Limnol Freshw Biol 358–363.

36. Hahn MW, Jezberová J, Koll U, Saueressig-Beck T, Schmidt J. 2016. Complete ecological isolation and cryptic diversity in Polynucleobacter bacteria not resolved by 16S rRNA gene sequences. ISME Journal 10:1642–1655.

37. Horňák K, Kasalický V, Šimek K, Grossart HP. 2017. Strain-specific consumption and transformation of alga-derived dissolved organic matter by members of the Limnohabitans-C and Polynucleobacter-B clusters of Betaproteobacteria. Environ Microbiol 19:4519–4535.

38. Salcher MM, Neuenschwander SM, Posch T, Pernthaler J. 2015. The ecology of pelagic freshwater methylotrophs assessed by a high-resolution monitoring and isolation campaign. ISME Journal 9:2442–2453.

39. Liang Y, Zhang M, Wang M, Zhang W, Qiao C, Luo Q, Lu X, Kelly RM. 2020. Freshwater Cyanobacterium Synechococcus elongatus PCC 7942 Adapts to an Environment with Salt Stress via Ion-Induced Enzymatic Balance of Compatible Solutes 10.1128/AEM.

40. Bubak I, Śliwińska-Wilczewska S, Głowacka P, Szczerba A, Możdżeń K. 2020. The importance of allelopathic picocyanobacterium Synechococcus sp. On the abundance, biomass formation, and structure of phytoplankton assemblages in three freshwater lakes. Toxins (Basel) 12.

41. Te SH, Kok JWK, Luo R, You L, Sukarji NH, Goh KC, Sim ZY, Zhang D, He Y, Gin KYH. 2023. Coexistence of Synechococcus and Microcystis Blooms in a Tropical Urban Reservoir and Their Links with Microbiomes. Environ Sci Technol 57:1613–1624.

42. Pancrace C, Barny MA, Ueoka R, Calteau A, Scalvenzi T, Pédron J, Barbe V, Piel J, Humbert JF, Gugger M. 2017. Insights into the Planktothrix genus: Genomic and metabolic comparison of benthic and planktic strains. Sci Rep 7.

43. Hendrayanti D, Prihantini NB, Ningsih F, Maulana F. 2023. Next Generation Sequencing (NGS) for Cyanobacterial study in Agung and Sunter Barat Lakes, North Jakarta, Indonesia. Biodiversitas 24:1117–1124.

44. Moiron M, Rimet F, Girel C, Jacquet S. 2021. Die hard in Lake Bourget! The case of Planktothrix rubescens reborn. Ann Limnol 57.

45. Li XZ, Zhang L, Poole K. 2002. SmeC, an outer membrane multidrug efflux protein of Stenotrophomonas maltophilia. Antimicrob Agents Chemother 46:333–343.

46. Cho HH, Sung JY, Kwon KC, Koo SH. 2012. Expression of sme efflux pumps and multilocus sequence typing in clinical isolates of Stenotrophomonas maltophilia. Ann Lab Med 32:38–43.

47. Gu Q, Sun M, Lin T, Zhang Y, Wei X, Wu S, Zhang S, Pang R, Wang J, Ding Y, Liu Z, Chen L, Chen W, Lin X, Zhang J, Chen M, Xue L, Wu Q. 2022. Characteristics of Antibiotic Resistance Genes and Antibiotic-Resistant Bacteria in Full-Scale Drinking Water Treatment System Using Metagenomics and Culturing. Front Microbiol 12.

48. Ryan RP, Monchy S, Cardinale M, Taghavi S, Crossman L, Avison MB, Berg G, van der Lelie D, Dow JM. 2009. The versatility and adaptation of bacteria from the genus Stenotrophomonas. Nat Rev Microbiol 10.1038/nrmicro2163.

49. Möker N, Brocker M, Schaffer S, Krämer R, Morbach S, Bott M. 2004. Deletion of the genes encoding the MtrA-MtrB two-component system of Corynebacterium glutamicum has a strong influence on cell morphology, antibiotics susceptibility and expression of genes involved in osmoprotection. Mol Microbiol 54:420–438.

50. Gorla P, Plocinska R, Sarva K, Satsangi AT, Pandeeti E, Donnelly R, Dziadek J, Rajagopalan M, Madiraju M V. 2018. MtrA response regulator controls cell division and cell wall metabolism and affects susceptibility of mycobacteria to the first line antituberculosis drugs. Front Microbiol 9.

51. Zhu Y, Zhang P, Zhang J, Wang J, Lu Y, Pang X. 2020. Impact on Multiple Antibiotic Pathways Reveals MtrA as a Master Regulator of Antibiotic Production in Streptomyces spp. and Potentially in Other Actinobacteria. Appl Environ Microbiol 86.

52. Schneider I, Bauernfeind A. 2015. Intrinsic carbapenem-hydrolyzing oxacillinases from members of the genus pandoraea. Antimicrob Agents Chemother 59:7136–7141.

53. Yoon EJ, Jeong SH. 2021. Class D β-lactamases. Journal of Antimicrobial Chemotherapy. Oxford University Press 10.1093/jac/dkaa513.

54. Madritch MD, Jordan LM, Lindroth RL. 2007. Interactive effects of condensed tannin and cellulose additions on soil respiration. Canadian Journal of Forest Research 37:2063– 2067.

55. Siniscalchi D, Cardoso A da S, Corrêa DC da C, Ferreira MR, Andrade MEB, da Cruz LHG, Ruggieri AC, Reis RA. 2022. Effects of condensed tannins on greenhouse gas emissions and nitrogen dynamics from urine-treated grassland soil. Environmental Science and Pollution Research 29:85026–85035.

56. Salcher MM, Schaefle D, Kaspar M, Neuenschwander SM, Ghai R. 2019. Evolution in action: habitat transition from sediment to the pelagial leads to genome streamlining in Methylophilaceae. ISME Journal 13:2764–2777.

57. Yan X, Xu X, Wang M, Wang G, Wu S, Li Z, Sun H, Shi A, Yang Y. 2017. Climate warming and cyanobacteria blooms: Looks at their relationships from a new perspective. Water Res 125:449–457.

58. Honda JR, Virdi R, Chan ED. 2018. Global environmental nontuberculous mycobacteria and their contemporaneous man-made and natural niches. Front Microbiol. Frontiers Media S.A. 10.3389/fmicb.2018.02029.

59. Bolger AM, Lohse M, Usadel B. 2014. Trimmomatic: A flexible trimmer for Illumina sequence data. Bioinformatics 30:2114–2120.

60. Kloke JD, Mckean JW. Rfit: Rank-based Estimation for Linear Models.

