## Supplemental Tables S1,S2,S4,S5 for "Pond Water Microbiome Taxa Profiles and Antibiotic Resistance Genes Associated with Acidity and Tannins"

Vaccaro et al. 2024. Table S1. Metadata for 2021

| Date | Pond | Latitude | Longitude | Field pH | Conductivity (µS/cm) | Water Temperature | DO mg/L | Tannins mg/L | PO4, mg/L | Nitrate, NO3 - mg/L | Ammonia, NH4 mg/L | Water Vegetation | Species | Leaf debris input | Species | Air Temperature (°C) | Weather Conditions | Wind Direction | Wind Speed (mph) | Humidity (%) | Notes |
|---|---|---|---|---|---|---|---|---|---|---|---|---|---|---|---|---|---|---|---|---|---|
| 6/9/2021 | Burtnett | 40°20'58" N | 82°19'31" W | 6.82 | 145 | 23.7 | 4.05 | 2 | 2.51 | 0 | 0.02 | yes | duckweed, cattails, | yes | oak | 22.8 | cloudy | northwest | 6 | 80 | has been raining for past few days, dogs, sheep |
| 6/9/2021 | Burtnett | 40°20'58" N | 82°19'31" W | 6.82 | 145 | 23.7 | 4.05 | 2.3 | 2.59 | 0 | 0.01 | yes | duckweed, cattails, | yes | oak | 22.8 | cloudy | northwest | 6 | 80 | has been raining for past few days, dogs, sheep |
| 6/16/2021 | Burtnett | 40°20'58" N | 82°19'31" W | 7.33 | 107 | 19.3 | 7.17 | 1.5 | 1.9 | 0 | 0.01 | yes | duckweed, cattails, | yes | oak | 23.3 | sunny | southeast | 9 | 36 | minnows, sheep, dogs |
| 6/16/2021 | Burtnett | 40°20'58" N | 82°19'31" W | 7.33 | 107 | 19.3 | 7.17 | 1.3 | 1.9 | 0.2 | 0.01 | yes | duckweed, cattails, | yes | oak | 23.3 | sunny | southeast | 9 | 36 | minnows, sheep, dogs |
| 6/23/2021 | Burtnett | 40°20'58" N | 82°19'31" W | 7.53 | 97 | 21.6 | 9.04 | 1.4 | 2.25 | 0 | 0.03 | yes | duckweed, cattails, | yes | oak | 23 | sunny | northeast | 5 | 39 | minnows, sheep, dogs |
| 6/23/2021 | Burtnett | 40°20'58" N | 82°19'31" W | 7.53 | 97 | 21.6 | 9.04 | 1.3 | 2.43 | 0.2 | 0.05 | yes | duckweed, cattails, | yes | oak | 23 | sunny | northeast | 5 | 39 | minnows, sheep, dogs |
| 6/30/2021 | Burtnett | 40°20'58" N | 82°19'31" W | 7.88 | 122 | 26.4 | 8.97 | 1.9 | 2.89 | 0 | 0 | yes | duckweed, cattails, | yes | oak | 27 | partially cloudy | east | 5 | 71 | minnows, sheep, dogs, mineral analysis on Friday due to illness |
| 6/30/2021 | Burtnett | 40°20'58" N | 82°19'31" W | 7.88 | 122 | 26.4 | 8.97 | 1.9 | 3.04 | 0 | 0.01 | yes | duckweed, cattails, | yes | oak | 27 | partially cloudy | east | 5 | 71 | minnows, sheep, dogs, mineral analysis on Friday due to illness |
| 7/7/2021 | Burtnett | 40°20'58" N | 82°19'31" W | 9.4 | 156 | 28.7 | 12.58 | 1 | 2.97 | 0.1 | 0.04 | yes | cattails, low viny vegetation | yes | oak | 28 | partially cloudy | northeast | 3 | 66 | minnows, snails, dogs, evidence of water birds and raccoons |
| 7/7/2021 | Burtnett | 40°20'58" N | 82°19'31" W | 9.4 | 156 | 28.7 | 12.58 | 1.1 | 3.15 | 0.1 | 0 | yes | cattails, low viny vegetation | yes | oak | 28 | partially cloudy | northeast | 3 | 66 | minnows, snails, dogs, evidence of water birds and raccoons |
| 7/14/2021 | Burtnett | 40°20'58" N | 82°19'31" W | 7.14 | 112 | 26.5 | 8.87 | 1.8 | 1.51 | 0 | 0.06 | yes | cattails, low viny vegetation | yes | oak | 26 | cloudy | northeast | 10 | 73 | minnows, snails, dogs |
| 7/14/2021 | Burtnett | 40°20'58" N | 82°19'31" W | 7.14 | 112 | 26.5 | 8.87 | 1.7 | 1.49 | 0 | 0.03 | yes | cattails, low viny vegetation | yes | oak | 26 | cloudy | northeast | 10 | 73 | minnows, snails, dogs |
| 7/21/2021 | Burtnett | 40°20'58" N | 82°19'31" W | 6.87 | 106 | 22.6 | 7.18 | 1.7 | 2.71 | 0.1 | 0.04 | yes | cattails, low viny vegetation, white flowers | yes | oak | 24 | cloudy | south | 9 | 61 | minnows, snails, dogs, geese, ducks, water more full, white flowers |
| 7/21/2021 | Burtnett | 40°20'58" N | 82°19'31" W | 6.87 | 106 | 22.6 | 7.18 | 1.8 | 2.18 | 0 | 0.01 | yes | cattails, low viny vegetation, white flowers | yes | oak | 24 | cloudy | south | 9 | 61 | minnows, snails, dogs, geese, ducks, water more full, white flowers |
| 7/28/2021 | Burtnett | 40°20'58" N | 82°19'31" W | 8.34 | 118 | 25 | 10.65 | 2.2 | 2.36 | 0 | 0.05 | yes | cattails, low viny vegetation, white flowers,algae | yes | oak | 30 | sunny | southeast | 5 | 51 | minnows, snails, dogs, geese, ducks, water up to cattails, green algae |
| 7/28/2021 | Burtnett | 40°20'58" N | 82°19'31" W | 8.34 | 118 | 25 | 10.65 | 2.2 | 2.36 | 0 | 0.05 | yes | cattails, low viny vegetation, white flowers,algae | yes | oak | 30 | sunny | southeast | 5 | 51 | minnows, snails, dogs, geese, ducks, water up to cattails, green algae |
| 9/7/2021 | Burtnett | 40°20'58" N | 82°19'31" W | 6.08 | 91 | 19.8 | 6.06 | 2.3 | 1.45 | 0 | 0.05 | yes | cattails, low viny vegetation, white flowers,algae | yes | oak | 28 | sunny | northeast | 8 | 58 | minnows, snails, vultures, sheep, dogs |
| 9/7/2021 | Burtnett | 40°20'58" N | 82°19'31" W | 6.08 | 91 | 19.8 | 6.06 | 2.2 | 1 | 0 | 0.05 | yes | cattails, low viny vegetation, white flowers,algae | yes | oak | 28 | sunny | northeast | 8 | 58 | minnows, snails, vultures, sheep, dogs |
| 10/5/2021 | Burtnett | 40°20'58" N | 82°19'31" W | 6.97 | 100 | 20 | 9.07 | 3 | 0.48 | 0 | 0.07 | yes | cattails, low viny vegetation, white flowers,algae | yes | oak | 14 | foggy | southwest | 0 | 100 | minnows, snails, dogs, sheep, ducks |
| 10/5/2021 | Burtnett | 40°20'58" N | 82°19'31" W | 6.97 | 100 | 20 | 9.07 | 2.3 | 0.53 | 0 | 0.07 | yes | cattails, low viny vegetation, white flowers,algae | yes | oak | 14 | foggy | southwest | 0 | 100 | minnows, snails, dogs, sheep, ducks |
| 6/7/2021 | Foundation | 40°23'10" N | 82°29'49" W | 8.9 | 590 | 24.6 | 9 | 0.3 | 0.06 | 0.3 | 0 | yes | algae | yes | willow | 25 | raining/thunder | northeast? | 8 | 70 | large amount of geese and goose poop, large fish in water |
| 6/7/2021 | Foundation | 40°23'10" N | 82°29'49" W | 8.9 | 590 | 24.6 | 9 | 0.4 | 0.15 | 0 | 0 | yes | algae | yes | willow | 25 | raining/thunder | northeast? | 8 | 70 | large amount of geese and goose poop, large fish in water |
| 6/14/2021 | Foundation | 40°23'10" N | 82°29'49" W | 8.21 | 584 | 26.5 | 10.23 | 0 | 0.05 | 0 | 0 | yes | algae | yes | willow | 28.3 | partially cloudy | southeast | 13 | 49 | large amount of geese and goose poop, large fish in water |
| 6/21/2021 | Foundation | 40°23'10" N | 82°29'49" W | 8.22 | 561 | 26 | 8.41 | 0.2 | 0.03 | 0.1 | 0 | yes | duckweed, cattails, | yes | willow | 26 | partially cloudy | southeast | 13 | 65 | turtles, evidence of geese |
| 6/21/2021 | Foundation | 40°23'10" N | 82°29'49" W | 8.22 | 561 | 26 | 8.41 | 0 | 0.04 | 0 | 0 | yes | duckweed, cattails, | yes | willow | 26 | partially cloudy | southeast | 13 | 65 | turtles, evidence of geese |
| 6/28/2021 | Foundation | 40°23'10" N | 82°29'49" W | 8.3 | 560 | 28.7 | 9.6 | 0.3 | 0.03 | 0.2 | 0 | yes | algae | yes | willow | 31 | sunny | northeast | 4 | 57 | turtles, bass, bluegill, geese |
| 6/28/2021 | Foundation | 40°23'10" N | 82°29'49" W | 8.3 | 560 | 28.7 | 9.6 | 0.3 | 0.09 | 0.2 | 0 | yes | algae | yes | willow | 31 | sunny | northeast | 4 | 57 | turtles, bass, bluegill, geese |
| 7/5/2021 | Foundation | 40°23'10" N | 82°29'49" W | 8.26 | 544 | 27.6 | 10.12 | 0.4 | 0.06 | 0.2 | 0 | yes | duckweed, cattails, | yes | willow | 31 | sunny | northeast | 8 | 57 | turtles, bass, bluegill, geese |
| 7/5/2021 | Foundation | 40°23'10" N | 82°29'49" W | 8.26 | 544 | 27.6 | 10.12 | 0.4 | 0.09 | 0.2 | 0 | yes | duckweed, cattails, | yes | willow | 31 | sunny | northeast | 8 | 57 | turtles, bass, bluegill, geese |
| 7/12/2021 | Foundation | 40°23'10" N | 82°29'49" W | 8.31 | 525 | 26 | 9.75 | 0.2 | 0.02 | 0 | 0 | yes | algae, submerged tree | yes | willow | 28 | cloudy | south | 8 | 74 | geese - swam through sampling site |
| 7/12/2021 | Foundation | 40°23'10" N | 82°29'49" W | 8.31 | 525 | 26 | 9.75 | 0.4 | 0.27 | 0.2 | 0 | yes | algae, submerged tree | yes | willow | 28 | cloudy | south | 8 | 74 | geese - swam through sampling site |
| 7/19/2021 | Foundation | 40°23'10" N | 82°29'49" W | 8.18 | 531 | 26.7 | 8.77 | 0.3 | 0.06 | 0.3 | 0 | yes | algae | yes | willow | 28 | partially cloudy | south | 6 | 51 | geese, osprey |
| 7/19/2021 | Foundation | 40°23'10" N | 82°29'49" W | 8.18 | 531 | 26.7 | 8.77 | 0.2 | 0.09 | 0.1 | 0 | yes | algae | yes | willow | 28 | partially cloudy | south | 6 | 51 | geese, osprey |
| 7/26/2021 | Foundation | 40°23'10" N | 82°29'49" W | 8.25 | 529 | 28 | 9.56 | 0.2 | 0.06 | 0.3 | 0 | yes | algae, submerged tree | yes | willow | 30 | sunny | west | 5 | 67 | geese, osprey |
| 7/26/2021 | Foundation | 40°23'10" N | 82°29'49" W | 8.25 | 529 | 28 | 9.56 | 0.2 | 0.08 | 0.2 | 0 | yes | algae, submerged tree | yes | willow | 30 | sunny | west | 5 | 67 | geese, osprey |
| 10/19/2021 | Foundation | 40°23'10" N | 82°29'49" W | 7.8 | 561 | 17.9 | 7.92 | 0.4 | 0.69 | 0.1 | 0.03 | yes | milfoil? Algae mats | yes | willow | 14 | clear, sunny | northwest | 5 | 61 | carp, turtles, bass, geese |
| 10/19/2021 | Foundation | 40°23'10" N | 82°29'49" W | 7.8 | 561 | 17.9 | 7.92 | 0.8 | 0.05 | 0.3 | 0.03 | yes | milfoil? Algae mats | yes | willow | 14 | clear, sunny | northwest | 5 | 61 | carp, turtles, bass, geese |
| 11/9/2021 | Foundation | 40°23'10" N | 82°29'49" W | 7.73 | 559 | 11.6 | 9.6 | 0.3 | 0.02 | 0.3 | 0.02 | yes | milfoil? Algae mats | yes | willow | 18 | clear, sunny | northeast | 8 | 46 | minnows |
| 11/16/2021 | Foundation | 40°23'10" N | 82°29'49" W | 7.73 | 559 | 11.6 | 9.6 | 0.3 | 0.15 | 0.2 | 0 | yes | milfoil? Algae mats | yes | willow | 18 | clear, sunny | northeast | 8 | 46 | minnows |
| 11/16/2021 | Foundation | 40°23'10" N | 82°29'49" W | 7.9 | 568 | 9.5 | 10.11 | 0.3 | 0.04 | 0.2 | 0 | yes | milfoil? Algae mats | yes | willow | 9 | cloudy | northwest | 3 | 64 | geese |

| | | | | | | | | | | | | | | | | | | | | | |
|---|---|---|---|---|---|---|---|---|---|---|---|---|---|---|---|---|---|---|---|---|---|
| **11/16/2021** | **Foundation** | 40°23'10" N | 82°29'49" W | 7.9 | 568 | 9.5 | 10.11 | 0.5 | 0.04 | 0.2 | 0 | yes | milfoil? Algae mats | yes | willow | 9 | cloudy | northwest | 3 | 64 | geese |
| **6/7/2021** | **McManis** | 40°23'53" N | 82°24'24" W | 8.65 | 200 | 25.5 | 10.1 | 0.4 | 0.03 | 0 | 0.03 | yes | duckweed, algae | yes | willow, oa | 25 | sunny | northeast? | 8 | 70 | peacock, duck,goat, around pond |
| **6/7/2021** | **McManis** | 40°23'53" N | 82°24'24" W | 8.65 | 200 | 25.5 | 10.1 | 0.5 | 0.6 | 0.3 | 0 | yes | duckweed, algae | yes | willow, oa | 25 | sunny | northeast? | 8 | 70 | peacock, duck,goat, around pond |
| **6/14/2021** | **McManis** | 40°23'53" N | 82°24'24" W | 8.38 | 196 | 26 | 10.81 | 0 | 0.07 | 0.3 | 0 | yes | duckweed, algae | yes | willow, oa | 28.3 | partially cloudy | southeast | 13 | 49 | peacock, duck,goat, around pond |
| **6/14/2021** | **McManis** | 40°23'53" N | 82°24'24" W | 8.38 | 196 | 26 | 10.81 | 0 | 0 | 0.1 | 0.02 | yes | duckweed, algae | yes | willow, oa | 28.3 | partially cloudy | southeast | 13 | 49 | peacock, duck,goat, around pond |
| **6/21/2021** | **McManis** | 40°23'53" N | 82°24'24" W | 8.85 | 202 | 25.8 | 9.65 | 0.4 | 0.07 | 0.1 | 0.01 | yes | duckweed, algae | yes | willow, oa | 26 | partially cloudy | southeast | 13 | 65 | peacock, duck,goat, cat around pond. Minnows in pond |
| **6/21/2021** | **McManis** | 40°23'53" N | 82°24'24" W | 8.85 | 202 | 25.8 | 9.65 | 0.6 | 0.6 | 0.2 | 0 | yes | duckweed, algae | yes | willow, oa | 26 | partially cloudy | southeast | 13 | 65 | peacock, duck,goat, cat around pond. Minnows in pond |
| **6/28/2021** | **McManis** | 40°23'53" N | 82°24'24" W | 9.2 | 266 | 28.9 | 9.9 | 0.4 | 0.02 | 0.3 | 0.02 | yes | duckweed, algae | yes | willow, oa | 31 | sunny | northeast | 4 | 57 | peacock, duck,goat, cat around pond. Minnows in pond |
| **6/28/2021** | **McManis** | 40°23'53" N | 82°24'24" W | 9.2 | 266 | 28.9 | 9.9 | 0.6 | 0.15 | 0.4 | 0 | yes | duckweed, algae | yes | willow, oa | 31 | sunny | northeast | 4 | 57 | peacock, duck,goat, cat around pond. Minnows in pond |
| **7/5/2021** | **McManis** | 40°23'53" N | 82°24'24" W | 7.58 | 211 | 26.5 | 8.08 | 0.7 | 0.11 | 0.3 | 0 | yes | duckweed, algae | yes | willow, oa | 31 | sunny | northeast | 8 | 57 | peacock, duck,goat, around pond, more duckweed than before |
| **7/5/2021** | **McManis** | 40°23'53" N | 82°24'24" W | 7.58 | 211 | 26.5 | 8.08 | 0.8 | 0.03 | 0.3 | 0 | yes | duckweed, algae | yes | willow, oa | 31 | sunny | northeast | 8 | 57 | peacock, duck,goat, around pond, more duckweed than before |
| **7/12/2021** | **McManis** | 40°23'53" N | 82°24'24" W | 7.09 | 203 | 24.2 | 5.8 | 0.4 | 0.12 | 0.2 | 0.02 | yes | duckweed, algae | yes | willow, oa | 28 | cloudy | south | 8 | 74 | peacock, duck,goat, around pond, duckweed |
| **7/12/2021** | **McManis** | 40°23'53" N | 82°24'24" W | 7.09 | 203 | 24.2 | 5.8 | 0.4 | 0.27 | 0 | 0.02 | yes | duckweed, algae | yes | willow, oa | 28 | cloudy | south | 8 | 74 | peacock, duck,goat, around pond, duckweed |
| **7/19/2021** | **McManis** | 40°23'53" N | 82°24'24" W | 7.16 | 211 | 24.4 | 10.05 | 0.5 | 0.06 | 0.1 | 0 | yes | duckweed, algae | yes | willow, oa | 28 | partially cloudy | south | 6 | 51 | peacock, duck,goat, around pond, duckweed |
| **7/19/2021** | **McManis** | 40°23'53" N | 82°24'24" W | 7.16 | 211 | 24.4 | 10.05 | 0.8 | 0.13 | 0 | 0 | yes | duckweed, algae | yes | willow, oa | 28 | partially cloudy | south | 6 | 51 | peacock, duck,goat, around pond, duckweed |
| **7/26/2021** | **McManis** | 40°23'53" N | 82°24'24" W | 8.99 | 221 | 26.2 | 11.78 | 0.2 | 0.09 | 0.2 | 0 | yes | duckweed, algae | yes | willow, oa | 30 | sunny | west | 5 | 67 | peacock, duck,goat, around pond, duckweed |
| **7/26/2021** | **McManis** | 40°23'53" N | 82°24'24" W | 8.99 | 221 | 26.2 | 11.78 | 0.2 | 0.01 | 0.2 | 0 | yes | duckweed, algae | yes | willow, oa | 30 | sunny | west | 5 | 67 | peacock, duck,goat, around pond, duckweed |
| **6/9/2021** | **Porter** | 40° 22' 21.118 | 82° 24' 56.97" | 6.71 | 96 | 23.4 | 6.06 | 0.5 | 0.07 | 0.4 | 0.09 | no | n/a | yes | oak, map | 22.8 | cloudy | northwest | 6 | 80 | has been raining for past few days |
| **6/9/2021** | **Porter** | 40° 22' 21.118 | 82° 24' 56.97" | 6.71 | 96 | 23.4 | 6.06 | 0.6 | 0.11 | 0.4 | 0.1 | no | n/a | yes | oak, map | 22.8 | cloudy | northwest | 6 | 80 | has been raining for past few days |
| **6/16/2021** | **Porter** | 40° 22' 21.118 | 82° 24' 56.97" | 6.86 | 90 | 22.8 | 7.72 | 0.5 | 0.15 | 0.4 | 0 | yes | algae | yes | oak, map | 23.3 | sunny | southeast | 9 | 36 | fish in water |
| **6/16/2021** | **Porter** | 40° 22' 21.118 | 82° 24' 56.97" | 6.86 | 90 | 22.8 | 7.72 | 0.6 | 0.09 | 0.2 | 0 | yes | algae | yes | oak, map | 23.3 | sunny | southeast | 9 | 36 | fish in water |
| **6/23/2021** | **Porter** | 40° 22' 21.118 | 82° 24' 56.97" | 7.16 | 89 | 22.5 | 8.42 | 0.6 | 0.16 | 0.2 | 0.01 | yes | algae | yes | oak, map | 23 | sunny | northeast | 5 | 39 | fish in water |
| **6/23/2021** | **Porter** | 40° 22' 21.118 | 82° 24' 56.97" | 7.16 | 89 | 22.5 | 8.42 | 0.6 | 0.11 | 0.2 | 0 | yes | algae | yes | oak, map | 23 | sunny | northeast | 5 | 39 | fish in water |
| **6/30/2021** | **Porter** | 40° 22' 21.118 | 82° 24' 56.97" | 7.15 | 90 | 26.3 | 7.25 | 0.7 | 0.18 | 0 | 0 | yes | algae | yes | oak, map | 27 | partially cloudy | east | 5 | 71 | fish in water, mineral analysis on Friday due to illness |
| **6/30/2021** | **Porter** | 40° 22' 21.118 | 82° 24' 56.97" | 7.15 | 90 | 26.3 | 7.25 | 0.7 | 0.01 | 0.2 | 0 | yes | algae | yes | oak, map | 27 | partially cloudy | east | 5 | 71 | fish in water, mineral analysis on Friday due to illness |
| **7/7/2021** | **Porter** | 40° 22' 21.118 | 82° 24' 56.97" | 7.18 | 90 | 26.8 | 7.49 | 0.5 | 0.1 | 0.2 | 0.02 | yes | algae | yes | oak, map | 28 | partially cloudy | northeast | 3 | 66 | carp, bass, sunfish, freshwater mussels |
| **7/7/2021** | **Porter** | 40° 22' 21.118 | 82° 24' 56.97" | 7.18 | 90 | 26.8 | 7.49 | 0.5 | 0.1 | 0.2 | 0.03 | yes | algae | yes | oak, map | 28 | partially cloudy | northeast | 3 | 66 | carp, bass, sunfish, freshwater mussels |
| **7/14/2021** | **Porter** | 40° 22' 21.118 | 82° 24' 56.97" | 7.02 | 93 | 23.7 | 8.62 | 0.5 | 0.15 | 0.6 | 0.06 | yes | algae | yes | oak, map | 26 | cloudy | northeast | 10 | 73 | carp, bass, sunfish, freshwater mussels |
| **7/14/2021** | **Porter** | 40° 22' 21.118 | 82° 24' 56.97" | 7.02 | 93 | 23.7 | 8.62 | 0.4 | 0.05 | 0.7 | 0.03 | yes | algae | yes | oak, map | 26 | cloudy | northeast | 10 | 73 | carp, bass, sunfish, freshwater mussels |
| **7/21/2021** | **Porter** | 40° 22' 21.118 | 82° 24' 56.97" | 7.29 | 103 | 22.1 | 11.16 | 0.4 | 0.16 | 1.4 | 0 | yes | algae | yes | oak, map | 24 | cloudy | south | 9 | 61 | carp, bass, sunfish, freshwater mussels |
| **7/21/2021** | **Porter** | 40° 22' 21.118 | 82° 24' 56.97" | 7.29 | 103 | 22.1 | 11.16 | 0.5 | 0.05 | 1.2 | 0 | yes | algae | yes | oak, map | 24 | cloudy | south | 9 | 61 | carp, bass, sunfish, freshwater mussels |
| **7/28/2021** | **Porter** | 40° 22' 21.118 | 82° 24' 56.97" | 7.47 | 108 | 25 | 10.8 | 0.4 | 0.08 | 1.5 | 0.05 | yes | algae | yes | oak, map | 30 | sunny | southeast | 5 | 51 | carp, bass, sunfish, freshwater mussels |

Vaccaro et al. 2024. Table S2. Metadata for 2022

| Date | Pond | Latitude | Longitude | Field pH | Conductivity (uS/cm) | Water Temperature (°C) | DO mg/L | Tannins mg/L | PO4,-P mg/L | Nitrate, NO3- mg/L | Ammonia NH4-N, mg/L | Water vegetation | Species | Leaf debris input | species | Air Temp (°C) | Weather Conditions | Wind Direction | Wind Speed (mph) | Humidity (%) | Notes | DOC |
|---|---|---|---|---|---|---|---|---|---|---|---|---|---|---|---|---|---|---|---|---|---|---|
| 9/13/2022 | Burtnett | 40°20'58" N | 82°19'31" W | 6.31 | 137 | 18.4 | n/a | 2 | 1.44 | 0 | 0 | yes | duckweed, cattails, grasses | yes | willow | 18.9 | partially cloudy | east | 16 | 75 | | |
| 9/13/2022 | Burtnett | 40°20'58" N | 82°19'31" W | 6.31 | 137 | 18.4 | n/a | 2 | 1.44 | 0 | 0 | yes | duckweed, cattails, grasses | yes | willow | 18.9 | partially cloudy | east | 16 | 75 | | |
| 9/20/2022 | Burtnett | 40°20'58" N | 82°19'31" W | 6.7 | 184 | 16.2 | 2.41 | 1.6 | 0.73 | 0.1 | 0.05 | yes | cattails, duckweed | yes | willow | 21.1 | sunny | east | 1 | 84 | | 9.6 |
| 9/20/2022 | Burtnett | 40°20'58" N | 82°19'31" W | 6.7 | 184 | 16.2 | 2.41 | 1.6 | 0.82 | 0.1 | 0.04 | yes | cattails, duckweed | yes | willow | 21.1 | sunny | east | 1 | 84 | | |
| 9/27/2022 | Burtnett | 40°20'58" N | 82°19'31" W | 6.03 | 122 | 10.72 | 2.93 | 2.4 | 0.77 | 0 | 0.06 | yes | Duckweed (light) | yes | willow | 10 | Partly Cloudy | west | 28 | 74.5 | Ducks | 6.9 |
| 9/27/2022 | Burtnett | 40°20'58" N | 82°19'31" W | 6.03 | 122 | 10.72 | 2.93 | 2.8 | 0.41 | 0 | 0.08 | yes | Duckweed (light) | yes | willow | 10 | Partly Cloudy | west | 28 | 74.5 | Ducks | |
| 10/4/2022 | Burtnett | 40°20'58" N | 82°19'31" W | 6.05 | 144 | 8.11 | 3.95 | 1.7 | 0.81 | 0 | 0.03 | yes, ducl | duckweed | Yes | Ducks | 16.1 | Scattered Clouds | NNW | 6 | 60 | | 11.0 |
| 10/4/2022 | Burtnett | 40°20'58" N | 82°19'31" W | 6.05 | 144 | 8.11 | 3.95 | 0.9 | 0.24 | 0.7 | 0 | yes, ducl | duckweed | Yes | Ducks | 16.1 | Scattered Clouds | NNW | 6 | 60 | | |
| 10/11/2022 | Burtnett | 40°20'58" N | 82°19'31" W | 5.91 | 196 | 6.17 | 4.72 | 3.5 | 1.51 | 0 | 0.13 | N/A | Geese | N/A | N/A | 10 | Sunny | SSE | 1.864 | 86 | More fine sediments | 12.0 |
| 10/11/2022 | Burtnett | 40°20'58" N | 82°19'31" W | 5.91 | 196 | 6.17 | 4.72 | 3.6 | 1.75 | 0 | 0.07 | N/A | Geese | N/A | N/A | 10 | Sunny | SSE | 1.864 | 86 | More fine sediments | |
| 10/18/2022 | Burtnett | 40°20'58" N | 82°19'31" W | 5.89 | 195 | 6.61 | 4.21 | 1.4 | 0.39 | 0.2 | 0.16 | Duckwee | Duckweed less abundant | | | 2.78 | Cloudy | W | 10.56 | 88 | | 8.2 |
| 10/18/2022 | Burtnett | 40°20'58" N | 82°19'31" W | 5.89 | 195 | 6.61 | 4.21 | 1.8 | 0.41 | 1.3 | 0.19 | Duckwee | Duckweed less abundant | | | 2.78 | Cloudy | W | 10.56 | 88 | | |
| 10/25/2022 | Burtnett | 40°20'58" N | 82°19'31" W | 6.18 | 197 | 9 | 3.1 | 1.5 | 0.68 | 1.7 | 0.1 | yes | | | | 9.44 | partly cloudy | SE | 10 | 44% | | 11.6 |
| 10/25/2022 | Burtnett | 40°20'58" N | 82°19'31" W | 6.18 | 197 | 9 | 3.1 | 1.6 | 1.03 | 2 | 0.33 | yes | | | | 9.44 | partly cloudy | SE | 10 | 44% | | |
| 11/1/2022 | Burtnett | 40°20'58" N | 82°19'31" W | 6.34 | 226 | 13.1 | 2.42 | 1.9 | 0.81 | 2.4 | 0.8 | Loss of D | Large Flock of Geese | | | 13.9 | Cloudy | WSW | 9.32 | 95% | | 8.8 |
| 11/1/2022 | Burtnett | 40°20'58" N | 82°19'31" W | 6.34 | 226 | 13.1 | 2.42 | 1.9 | 0.82 | 2 | 0.68 | Loss of D | Large Flock of Geese | | | 13.9 | Cloudy | WSW | 9.32 | 95% | | |
| 11/8/2022 | Burtnett | 40°20'58" N | 82°19'31" W | 5.9 | 270 | 5.39 | 4.33 | 2.2 | 0.63 | 0 | 0.04 | getation | low tide, low sediment | | | 6.67 | Sunny, Slight Wind | NNE | 11 | 68% | | 12.9 |
| 11/15/2022 | Burtnett | 40°20'58" N | 82°19'31" W | 6.07 | 180 | 3.1 | 3.4 | 3.6 | 0.53 | 0 | 0.17 | Pond mo | Ducks | | | 2.78 | Cloudy, Light Rain | ESE | 7 | 75% | | 8.0 |
| 11/15/2022 | Burtnett | 40°20'58" N | 82°19'31" W | 6.07 | 180 | 3.1 | 3.4 | 4 | 0.66 | 0 | 0.22 | Pond mo | Ducks | | | 2.78 | Cloudy, Light Rain | ESE | 7 | 75% | | |
| 9/13/2022 | Foundation | 40°23'10" N | 82°29'49" W | 8.1 | 512 | 22.3 | n/a | 0 | 0.01 | 0 | 0.03 | yes | milfoil(?) some sort of fanned aquatic plant | no | | 18.9 | partially cloudy | east | 16 | 75 | | |
| 9/13/2022 | Foundation | 40°23'10" N | 82°29'49" W | 8.1 | 512 | 22.3 | n/a | 0 | 0.01 | 0 | 0.03 | yes | milfoil(?) some sort of fanned aquatic plant | no | | 18.9 | partially cloudy | east | 16 | 75 | | |
| 9/20/2022 | Foundation | 40°23'10" N | 82°29'49" W | 8.34 | 561 | 23.1 | 8.4 | 0.3 | 0.05 | 0.3 | 0 | yes | green algae-like plant | no | | 21.1 | sunny | east | 1 | 84 | | 2.5 |
| 9/20/2022 | Foundation | 40°23'10" N | 82°29'49" W | 8.34 | 561 | 23.1 | 8.4 | 0.2 | 0.13 | 0.4 | 0.03 | yes | green algae-like plant | no | | 21.1 | sunny | east | 1 | 84 | | |
| 9/27/2022 | Foundation | 40°23'10" N | 82°29'49" W | 8.12 | 524 | 18.67 | 7.97 | 0.2 | 0.08 | 0 | 0 | yes | Algae like plant | no | | 10 | Partly Cloudy | west | 28 | 74.5 | osprey, ducks | 2.3 |
| 9/27/2022 | Foundation | 40°23'10" N | 82°29'49" W | 8.12 | 524 | 18.67 | 7.97 | 0.2 | 0.04 | 0 | 0.01 | yes | Algae like plant | no | | 10 | Partly Cloudy | west | 28 | 74.5 | osprey, ducks | |
| 10/4/2022 | Foundation | 40°23'10" N | 82°29'49" W | 8.08 | 638 | 15.7 | 8.8 | 0.2 | 0.02 | 0.4 | 0 | Leafy Wa | leaves | Yes | N/A | 16.1 | Scattered Clouds | NNW | 6 | 60 | | 2.4 |
| 10/4/2022 | Foundation | 40°23'10" N | 82°29'49" W | 8.08 | 638 | 15.7 | 8.8 | 0.3 | 0.02 | 1 | 0 | Leafy Wa | leaves | Yes | N/A | 16.1 | Scattered Clouds | NNW | 6 | 60 | | |
| 10/11/2022 | Foundation | 40°23'10" N | 82°29'49" W | 8.21 | 642 | 14.5 | 9.62 | 0.3 | 0.06 | 0.3 | 0.05 | N/A | N/A | N/A | N/A | 10 | Sunny | SSE | 1.864 | 86 | | 2.4 |
| 10/11/2022 | Foundation | 40°23'10" N | 82°29'49" W | 8.21 | 642 | 14.5 | 9.62 | 0.3 | 0 | 0.4 | 0.04 | N/A | N/A | N/A | N/A | 10 | Sunny | SSE | 1.864 | 86 | | |
| 10/18/2022 | Foundation | 40°23'10" N | 82°29'49" W | 7.96 | 148 | 11.4 | N/A | 0.4 | 0.05 | 0.5 | 0.14 | | DO Machine Broken | | | 2.78 | Cloudy | W | 10.56 | 88 | | 2.2 |
| 10/18/2022 | Foundation | 40°23'10" N | 82°29'49" W | 7.96 | 148 | 11.4 | N/A | 0.4 | 0.06 | 0.4 | 0.1 | | DO Machine Broken | | | 2.78 | Cloudy | W | 10.56 | 88 | | |
| 10/25/2022 | Foundation | 40°23'10" N | 82°29'49" W | 7.12 | 659 | 12.8 | N/A | 0 | 0 | 0.3 | 0.03 | | | | | 9.44 | partly cloudy | SE | 10 | 44% | | 2.2 |
| 10/25/2022 | Foundation | 40°23'10" N | 82°29'49" W | 7.12 | 659 | 12.8 | N/A | 0 | 0.08 | 0.3 | 0 | | | | | 9.44 | partly cloudy | SE | 10 | 44% | | |
| 11/1/2022 | Foundation | 40°23'10" N | 82°29'49" W | 7.83 | 666 | 13.4 | 9.85 | 0.3 | 0.17 | 0.2 | 0 | Clear Water, Low Sediment | | | | 13.9 | Cloudy | WSW | 9.32 | 95% | | 2.0 |
| 11/1/2022 | Foundation | 40°23'10" N | 82°29'49" W | 7.83 | 666 | 13.4 | 9.85 | 0.3 | 0.17 | 0.2 | 0 | Clear Water, Low Sediment | | | | 13.9 | Cloudy | WSW | 9.32 | 95% | | |
| 11/8/2022 | Foundation | 40°23'10" N | 82°29'49" W | 7.91 | 671 | 11.83 | 10.3 | 0.4 | 0.03 | 0.3 | 0 | | | not a lot of leaf litter | | 6.67 | Sunny, Slight Wind | NNE | 11 | 68% | | 2.1 |
| 11/8/2022 | Foundation | 40°23'10" N | 82°29'49" W | 7.91 | 671 | 11.83 | 10.3 | 0.4 | 0.06 | 0.4 | 0 | | | not a lot of leaf litter | | 6.67 | Sunny, Slight Wind | NNE | 11 | 68% | | |
| 11/15/2022 | Foundation | 40°23'10" N | 82°29'49" W | 7.66 | 668 | 8.56 | 9.88 | 0.2 | 0.05 | 0.1 | 0.03 | | | | | 2.78 | Cloudy, Light Rain | ESE | 7 | 75% | | 1.8 |
| 11/15/2022 | Foundation | 40°23'10" N | 82°29'49" W | 7.66 | 668 | 8.56 | 9.88 | 0.2 | 0 | 0.3 | 0.03 | | | | | 2.78 | Cloudy, Light Rain | ESE | 7 | 75% | | |
| 9/14/2022 | McManis | 40°23'53" N | 82°24'24" W | 6.6 | 226 | 20.8 | 5.7 | 0.8 | 0.08 | 0.2 | 0.03 | yes | duckweed! covering whole pond | yes | willow, oak, mullberry | 18.9 | cloudy | south | 0 | 84 | | 4.3 |
| 9/14/2022 | McManis | 40°23'53" N | 82°24'24" W | 6.6 | 226 | 20.8 | 5.7 | 0.8 | 0.08 | 0.2 | 0.03 | yes | duckweed! covering whole pond | yes | willow, oak, mullberry | 18.9 | cloudy | south | 0 | 84 | | |
| 9/22/2022 | McManis | 40°23'53" N | 82°24'24" W | 6.56 | 228 | 21.4 | 4.31 | 0.9 | 0.03 | 0.1 | 0.08 | yes | lots of duckweed | yes | willow, oak, mullberry | 16.1 | sunny | est northwe | 13 | 72 | | 4.3 |
| 9/22/2022 | McManis | 40°23'53" N | 82°24'24" W | 6.56 | 228 | 21.4 | 4.31 | 0.8 | 0 | 0.1 | 0.05 | yes | lots of duckweed | yes | willow, oak, mullberry | 16.1 | sunny | est northwe | 13 | 72 | | |
| 9/28/2022 | McManis | 40°23'53" N | 82°24'24" W | 6.38 | 228 | 16.1 | 1.66 | 0.6 | 0.09 | 0.4 | 0.01 | yes | lots of duckweed | yes | willow, oak, mullberry | 12.2 | Cloudy | northwest | 17 | 77 | Ducks, Pigs, Peacocks | 4.0 |
| 9/28/2022 | McManis | 40°23'53" N | 82°24'24" W | 6.38 | 228 | 16.1 | 1.66 | 0.7 | 0.07 | 0.4 | 0.06 | yes | lots of duckweed | yes | willow, oak, mullberry | 12.2 | Cloudy | northwest | 17 | 77 | Ducks, Pigs, Peacocks | |

| | | | | | | | | | | | | | | | | | | | | | | |
|---|---|---|---|---|---|---|---|---|---|---|---|---|---|---|---|---|---|---|---|---|---|---|
| **10/5/2022** | **McManis** | 40°23'53" N | 82°24'24" W | 6.95 | 286 | 15.4 | 2.95 | 0.7 | 0 | 0.4 | 0.06 | Lots of d | duckweed | Yes | Ducks, Pigs, Goats, Dog, Cats, F | 17.8 | Sunny | NW | 3 | 51 | | 4.1 |
| **10/12/2022** | **McManis** | 40°23'53" N | 82°24'24" W | 7.05 | 289 | 14.3 | 3.2 | 0.9 | 0.11 | 0.2 | 0.13 | | | | | 17.8 | Sunny | SSE | 6.214 | 83 | | 3.8 |
| **10/12/2022** | **McManis** | 40°23'53" N | 82°24'24" W | 7.05 | 289 | 14.3 | 3.2 | 0.8 | 0.17 | 0.2 | 0.13 | | | | | 17.8 | Sunny | SSE | 6.214 | 83 | | |
| **10/19/2022** | **McManis** | 40°23'53" N | 82°24'24" W | 7.28 | 295 | 10.8 | 5.56 | 0 | 0.04 | 0.4 | 0 | Duckwee | Ducks | Leaf Debris at Bottom | Willow, Pine, Oak | 2.78 | Rainy, Cold | W | 12 | 94 | | 3.9 |
| **10/19/2022** | **McManis** | 40°23'53" N | 82°24'24" W | 7.28 | 295 | 10.8 | 5.56 | 0 | 0.09 | 0.1 | 0 | Duckwee | Ducks | Leaf Debris at Bottom | Willow, Pine, Oak | 2.78 | Rainy, Cold | W | 12 | 94 | | |
| **10/26/2022** | **McManis** | 40°23'53" N | 82°24'24" W | 7.34 | 299 | 14.1 | 7.32 | 0.7 | 0.07 | 0 | 0.03 | yes | lots of duckweed (more than last week) | Yes | Lots of leaves and ducks! | 12.2 | rainy | SSW | 9 | 91% | | 4.1 |
| **10/26/2022** | **McManis** | 40°23'53" N | 82°24'24" W | 7.34 | 299 | 14.1 | 7.32 | 0.8 | 0 | 0 | 0.05 | yes | lots of duckweed (more than last week) | Yes | Lots of leaves and ducks! | 12.2 | rainy | SSW | 9 | 91% | | |
| **11/2/2022** | **McManis** | 40°23'53" N | 82°24'24" W | 7.28 | 151 | 12 | 7.37 | 0.7 | 0.02 | 0.2 | 0.03 | Duckwee | Ducks, Pigs nearby | Leaf Debris | | 6.6 | Cloudy, Foggy | SSE | 1 | 99% | | 4.1 |
| **11/2/2022** | **McManis** | 40°23'53" N | 82°24'24" W | 7.28 | 151 | 12 | 7.37 | 0.8 | 0.15 | 0.1 | 0 | Duckwee | Ducks, Pigs nearby | Leaf Debris | | 6.6 | Cloudy, Foggy | SSE | 1 | 99% | | |
| **11/9/2022** | **McManis** | 40°23'53" N | 82°24'24" W | 6.75 | 305 | 11.9 | 6.73 | 0.7 | 0.03 | 0 | 0.02 | less duckweed | | | | 6.11 | Sunny, Slight Wind | SSE | 5 | 62% | | 4.0 |
| **11/9/2022** | **McManis** | 40°23'53" N | 82°24'24" W | 6.75 | 305 | 11.9 | 6.73 | 0.7 | 0 | 0.2 | 0.2 | less duckweed | | | | 6.11 | Sunny, Slight Wind | SSE | 5 | 62% | | |
| **11/16/2022** | **McManis** | 40°23'53" N | 82°24'24" W | 6.62 | 303 | 7.6 | 7.63 | 0.7 | 0.05 | 0.3 | 0 | Light Du | Ducks and Wild Birds | | | 0.56 | Cloudy, Light Snow, W | W | 11 | 84% | | 3.8 |
| **11/16/2022** | **McManis** | 40°23'53" N | 82°24'24" W | 6.62 | 303 | 7.6 | 7.63 | 0.6 | 0.04 | 0 | 0.03 | Light Du | Ducks and Wild Birds | | | 0.56 | Cloudy, Light Snow, W | W | 11 | 84% | | |
| **9/14/2022** | **Porter** | 40° 22' 21.11 | 82° 24' 56.97" | 6.24 | 94 | 18.6 | 4.59 | 0.9 | 0.04 | 1.1 | 0.21 | no | | yes | oak, pine | 18.9 | cloudy | south | 0 | 84 | | 2.7 |
| **9/14/2022** | **Porter** | 40° 22' 21.11 | 82° 24' 56.97" | 6.24 | 94 | 18.6 | 4.59 | 0.9 | 0.06 | 0.8 | 0.24 | no | | yes | oak, pine | 18.9 | cloudy | south | 0 | 84 | | |
| **9/22/2022** | **Porter** | 40° 22' 21.11 | 82° 24' 56.97" | 6.62 | 96 | 20.6 | 6.62 | 0.8 | 0.03 | 0.8 | 0.04 | no | some side vegetation, but nothing on water | yes | oak, pine | 16.1 | sunny | est northwe | 13 | 72 | | 2.7 |
| **9/22/2022** | **Porter** | 40° 22' 21.11 | 82° 24' 56.97" | 6.62 | 96 | 20.6 | 6.62 | 0.8 | 0.03 | 0.6 | 0.16 | no | some side vegetation, but nothing on water | yes | oak, pine | 16.1 | sunny | est northwe | 13 | 72 | | |
| **9/28/2022** | **Porter** | 40° 22' 21.11 | 82° 24' 56.97" | 6.28 | 81 | 14.9 | 4.97 | 0.5 | 0.2 | 1.1 | 0.13 | yes | Clear, no duckweed but pine needles | yes | oak, pine | 12.2 | Cloudy | northwest | 17 | 77 | Oak/Pine Trees | 2.3 |
| **9/28/2022** | **Porter** | 40° 22' 21.11 | 82° 24' 56.97" | 6.28 | 81 | 14.9 | 4.97 | 0.5 | 0.06 | 0.5 | 0.15 | yes | Clear, no duckweed but pine needles | yes | oak, pine | 12.2 | Cloudy | northwest | 17 | 77 | Oak/Pine Trees | |
| **10/5/2022** | **Porter** | 40° 22' 21.11 | 82° 24' 56.97" | 7.08 | 112 | 13.7 | 6.55 | 0.7 | 0.06 | 0.3 | 0.24 | clear | N/A | Yes | Blue Heron | 18.3 | Sunny | NW | 3 | 51 | | 2.4 |
| **10/12/2022** | **Porter** | 40° 22' 21.11 | 82° 24' 56.97" | 7.08 | 110 | 12.2 | 6.28 | 0.7 | 0.44 | 0.3 | 0.25 | | | | | 17.8 | Sunny | SSE | 6.214 | 83 | | 2.3 |
| **10/12/2022** | **Porter** | 40° 22' 21.11 | 82° 24' 56.97" | 7.08 | 110 | 12.2 | 6.28 | 0.9 | 0.25 | 0.4 | 0.26 | | | | | 17.8 | Sunny | SSE | 6.214 | 83 | | |
| **10/19/2022** | **Porter** | 40° 22' 21.11 | 82° 24' 56.97" | 6.89 | 109 | 8.8 | 6.68 | 0 | 0.13 | 0.4 | 0 | | | Leaf Debris at Bottom | Maple, Pine, Oak | 3.33 | Rainy, Cold | W | 12 | 93 | | 2.4 |
| **10/19/2022** | **Porter** | 40° 22' 21.11 | 82° 24' 56.97" | 6.89 | 109 | 8.8 | 6.68 | 0 | 0.05 | 0.3 | 0 | | | Leaf Debris at Bottom | Maple, Pine, Oak | 3.33 | Rainy, Cold | W | 12 | 93 | | |
| **10/26/2022** | **Porter** | 40° 22' 21.11 | 82° 24' 56.97" | 7.12 | 110 | 12.6 | 7.64 | 1.6 | 0 | 0.2 | 0.01 | yes | Plants at the bottom | Yes | Lots of leaves - Pines, Oak, Ma | 12.2 | rainy | SSW | 9 | 91% | | 3.6 |
| **10/26/2022** | **Porter** | 40° 22' 21.11 | 82° 24' 56.97" | 7.12 | 110 | 12.6 | 7.64 | 1.5 | 0.06 | 0.2 | 0 | yes | Plants at the bottom | Yes | Lots of leaves - Pines, Oak, Ma | 12.2 | rainy | SSW | 9 | 91% | | |
| **11/2/2022** | **Porter** | 40° 22' 21.11 | 82° 24' 56.97" | 5.99 | 107 | 10.1 | 6.51 | 1.1 | 0.02 | 0.2 | 0 | Lots of le | Blue Heron | Lots of Leaf Debris | | 6.6 | Cloudy, Foggy | SSE | 1 | 99% | | 3.0 |
| **11/2/2022** | **Porter** | 40° 22' 21.11 | 82° 24' 56.97" | 5.99 | 107 | 10.1 | 6.51 | 0.9 | 0.03 | 0.1 | 0 | Lots of le | Blue Heron | Lots of Leaf Debris | | 6.6 | Cloudy, Foggy | SSE | 1 | 99% | | |
| **11/9/2022** | **Porter** | 40° 22' 21.11 | 82° 24' 56.97" | 6.1 | 109 | 9.7 | 4.71 | 0.9 | 0.04 | 0.4 | 0 | more leaves in pond | | | | 7.22 | Sunny, Slight Wind | SSE | 6 | 58% | | 2.7 |
| **11/9/2022** | **Porter** | 40° 22' 21.11 | 82° 24' 56.97" | 6.1 | 109 | 9.7 | 4.71 | 0.8 | 0.13 | 0.4 | 0 | more leaves in pond | | | | 7.22 | Sunny, Slight Wind | SSE | 6 | 58% | | |
| **11/16/2022** | **Porter** | 40° 22' 21.11 | 82° 24' 56.97" | 6.08 | 103 | 5.8 | 4.56 | 0.7 | 0.1 | 0.1 | 0.05 | Pond = Blue-ish Green in Color | | Lots of Leaves | | 0.56 | Cloudy, Light Snow, W | W | 11 | 81% | | 2.4 |
| **11/16/2022** | **Porter** | 40° 22' 21.11 | 82° 24' 56.97" | 6.08 | 103 | 5.8 | 4.56 | 0.7 | 0.12 | 0.1 | 0.05 | Pond = Blue-ish Green in Color | | Lots of Leaves | | 0.56 | Cloudy, Light Snow, W | W | 11 | 81% | | |
| **NA** | **Zymo Standard** | | | | | | | | | | | | | | | | | | | | | |
| **NA** | **Zymo Standard** | | | | | | | | | | | | | | | | | | | | | |
| **NA** | **Zymo Standard** | | | | | | | | | | | | | | | | | | | | | |
| **NA** | **Zymo Standard** | | | | | | | | | | | | | | | | | | | | | |
| **NA** | **Zymo Standard** | | | | | | | | | | | | | | | | | | | | | |
| | **Zymo Standard** | | | | | | | | | | | | | | | | | | | | | |

**Vaccaro et al. 2024. Table S4. P-values for Figure 7. Heat map of Spearman correlations between top-ranked ARGs and chemical measures.**

| 2021 | acidity | tannin | PO4 | NO3 | NH4 | cond | temp | DO |
|---|---|---|---|---|---|---|---|---|
| smeB | 0.04 | 0.09 | 0.89 | 0.13 | 0.22 | 0.32 | 0.02 | 0.08 |
| mtrA | 0.02 | 0.01 | 0.00 | 0.00 | 0.62 | 0.20 | 0.68 | 0.00 |
| OXA-156 | 0.77 | 0.00 | 0.03 | 0.00 | 0.19 | 0.58 | 0.62 | 0.14 |
| OXA-409 | 0.00 | 0.00 | 0.00 | 0.03 | 0.01 | 0.00 | 0.04 | 0.00 |
| SHV-100 | 0.45 | 0.76 | 0.61 | 0.71 | 0.45 | 0.56 | 0.00 | 0.50 |
| otrC | 0.07 | 0.18 | 0.11 | 0.25 | 0.27 | 0.26 | 0.71 | 0.32 |
| Erm(O)-lrm | 0.05 | 0.12 | 0.05 | 0.93 | 0.35 | 0.62 | 0.04 | 0.18 |
| CTX-M-25 | 0.65 | 0.29 | 0.07 | 0.08 | 0.08 | 0.77 | 0.17 | 0.74 |
| vanRO | 0.70 | 0.31 | 0.37 | 0.72 | 0.74 | 0.49 | 0.00 | 0.89 |
| floR | 0.34 | 0.93 | 0.07 | 0.39 | 0.04 | 0.41 | 0.03 | 0.44 |
| tlrB | 0.75 | 0.02 | 0.04 | 0.38 | 0.10 | 0.04 | 0.98 | 0.86 |
| FOX-3 | 0.00 | 0.00 | 0.00 | 0.60 | 0.00 | 0.00 | 0.04 | 0.02 |
| OXA-372 | 0.00 | 0.00 | 0.50 | 0.97 | 0.89 | 0.01 | 0.84 | 0.01 |
| FRI-1 | 0.31 | 0.75 | 0.27 | 0.51 | 0.65 | 0.03 | 0.88 | 0.16 |
| adeH | 0.20 | 0.00 | 0.00 | 0.00 | 0.02 | 0.39 | 0.61 | 0.44 |
| BUT-1 | 0.02 | 0.05 | 0.05 | 0.49 | 0.05 | 0.00 | 0.44 | 0.85 |
| OXA-137 | 0.99 | 0.80 | 0.43 | 0.46 | 0.27 | 0.08 | 0.38 | 0.95 |
| efrA | 0.87 | 0.69 | 0.70 | 0.08 | 0.94 | 0.65 | 0.58 | 0.33 |
| bcrC | 0.77 | 0.12 | 0.08 | 0.11 | 0.40 | 0.08 | 0.89 | 0.54 |
| OXA-46 | 0.38 | 0.00 | 0.08 | 0.26 | 0.05 | 0.00 | 0.02 | 0.53 |

| 2022 | acidity | tannin | PO4 | NO3 | NH4 | cond | temp | DO |
|---|---|---|---|---|---|---|---|---|
| smeB | 0.35 | 0.48 | 0.69 | 0.06 | 0.07 | 0.69 | 0.71 | 0.36 |
| mtrA | 0.01 | 0.01 | 0.00 | 0.99 | 0.00 | 0.00 | 0.10 | 0.01 |
| OXA-156 | 0.00 | 0.00 | 0.01 | 0.36 | 0.54 | 0.50 | 0.60 | 0.01 |
| OXA-409 | 0.00 | 0.00 | 0.00 | 0.47 | 0.00 | 0.13 | 0.18 | 0.12 |
| SHV-100 | 0.45 | 0.34 | 0.65 | 0.11 | 0.05 | 0.21 | 0.16 | 0.05 |
| otrC | 0.93 | 0.48 | 0.73 | 0.39 | 0.07 | 0.09 | 0.41 | 0.06 |
| Erm(O)-lrm | 0.89 | 0.62 | 0.40 | 0.73 | 0.54 | 0.46 | 0.06 | 0.08 |
| CTX-M-25 | 0.04 | 0.01 | 0.00 | 0.09 | 0.38 | 0.00 | 0.01 | 0.01 |
| vanRO | 0.62 | 0.84 | 0.98 | 0.41 | 0.27 | 0.71 | 0.39 | 0.25 |
| floR | 0.00 | 0.01 | 0.01 | 0.08 | 0.02 | 0.67 | 0.92 | 0.80 |
| tlrB | 0.20 | 0.24 | 0.33 | 0.63 | 0.13 | 0.00 | 0.01 | 0.23 |
| FOX-3 | 0.00 | 0.00 | 0.10 | 0.34 | 0.23 | 0.59 | 0.29 | 0.40 |
| OXA-372 | 0.16 | 0.98 | 0.07 | 0.24 | 0.38 | 0.07 | 0.19 | 0.11 |
| FRI-1 | 0.77 | 0.89 | 0.81 | 0.81 | 0.58 | 0.01 | 0.45 | 0.13 |
| adeH | 0.89 | 0.21 | 0.26 | 0.06 | 0.54 | 0.05 | 0.43 | 0.56 |
| BUT-1 | 0.42 | 0.76 | 0.80 | 0.04 | 0.40 | 0.06 | 0.45 | 0.08 |
| OXA-137 | 0.48 | 0.18 | 0.01 | 0.04 | 0.51 | 0.00 | 0.44 | 0.04 |
| efrA | 0.00 | 0.00 | 0.10 | 0.21 | 0.01 | 0.15 | 0.21 | 0.31 |
| bcrC | 0.04 | 0.01 | 0.23 | 0.19 | 0.49 | 0.09 | 0.12 | 0.21 |
| OXA-46 | 0.02 | 0.02 | 0.01 | 0.25 | 0.80 | 0.32 | 0.41 | 0.02 |

**Vaccaro et al. 2024. Table S5. Spearman correlations between ARGs and DOC values in 2022.**

| ARG | Spearman F | P-value |
|---|---|---|
| smeB | -0.02 | 0.47 |
| mtrA | 0.05 | 0.43 |
| OXA-156 | -0.41 | 0.00 |
| Erm(O)-lrm | -0.15 | 0.28 |
| floR | -0.19 | 0.21 |
| otrC | -0.24 | 0.09 |
| OXA-409 | 0.35 | 0.01 |
| SHV-100 | -0.1 | 0.27 |
| OXA-137 | -0.1 | 0.91 |
| tlrB | -0.27 | 0.05 |
| CTX-M-25 | 0.27 | 0.03 |
| FRI-1 | 0.06 | 0.91 |
| efrA | -0.54 | 0.00 |
| BUT-1 | 0.13 | 0.55 |
| vanRO | 0.03 | 0.79 |
| OXA-372 | 0 | 0.61 |
| bcrC | -0.35 | 0.14 |
| FOX-3 | -0.27 | 0.06 |
| OXA-46 | 0.53 | 0.00 |
| adeH | 0.28 | 0.21 |

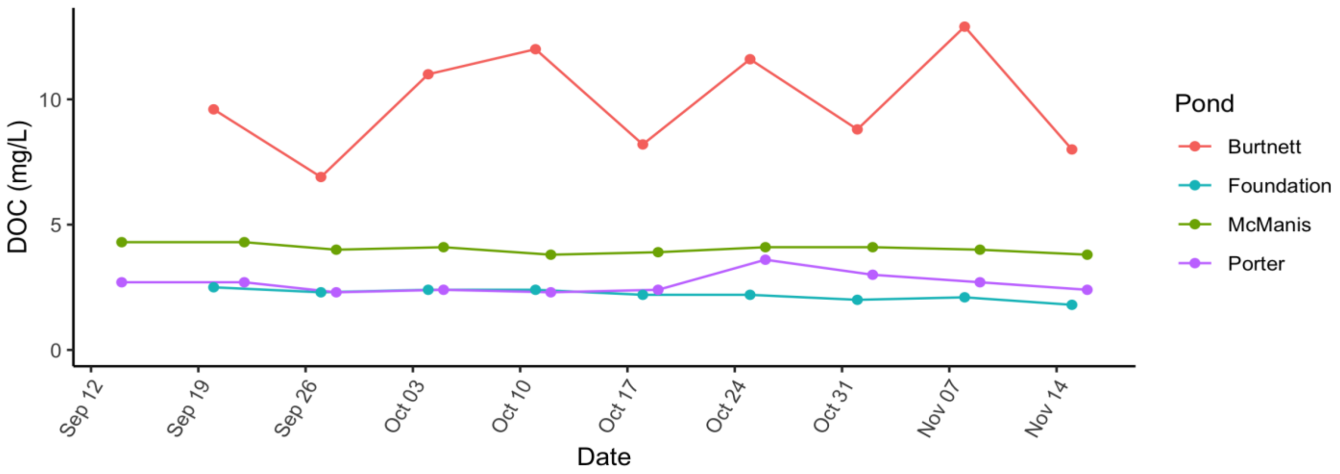
