## Supplemental Table S3 for "Pond Water Microbiome Taxa Profiles and Antibiotic Resistance Genes Associated with Acidity and Tannins"

**Vaccaro et al. 2024. Table S3. ShortBRED Markers**

>gb|AAG07064_1|ARO_3003693|mexK__Pseudomonas_aeruginosa_PAO1__TM_#01
ASLRSTGEEVQR
>gb|AAG07064_1|ARO_3003693|mexK__Pseudomonas_aeruginosa_PAO1__TM_#02
LESREALRKWLIERMNEDFPH
>gb|AAG07064_1|ARO_3003693|mexK__Pseudomonas_aeruginosa_PAO1__TM_#03
QFLQSALTGSH
>gb|CAC87665_1|ARO_3000855|CAU-1_beta-lactamase__Caulobacter_vibrioides__TM_#01
MKRLILAAAASLLALASAAHAD
>gb|CAC87665_1|ARO_3000855|CAU-1_beta-lactamase__Caulobacter_vibrioides__TM_#02
TSSEGHVVLDGGPNAETGKL
>gb|CAC87665_1|ARO_3000855|CAU-1_beta-lactamase__Caulobacter_vibrioides__TM_#03
ISRDDAPAMAAGHHIGDNIYGPTPMPAAKPDRSFGDQTKLKLGEIA
>gb|CAA27276_1|ARO_3002661|APH(7__)-Ia__Streptomyces_hygroscopicus__TM_#01
MTQESLLLLDRIDSDDSYASLRNDQEFWEPLARRALEELGLPVPPVLRVPGESTNPVLVG
EPDPVIKLFGEHWCGPESLASESEAYAVLADAPVPVPRLLGRGELRPGTGAWPWPYLVMS
RMTGTTWRSAMDGTTDRNALLALARELGRVLGRLHRVPLTGNTVLTPHSEVFPELLR
>gb|CAA27276_1|ARO_3002661|APH(7__)-Ia__Streptomyces_hygroscopicus__TM_#02
RLEDWLPDVDT
>gb|CAA27276_1|ARO_3002661|APH(7__)-Ia__Streptomyces_hygroscopicus__TM_#03
LAATEVTGIVDFTDVYAGDSRYSLVQLHLNAFRGDREILAALLDGAQWKRTEDFARELLA
FTFLHDFEVFEETPLDLSGFTDPEELAQFLWGPPDTAPGA
>gb|AAP82271|ARO_3004123|Burkholderia_pseudomallei_Omp38__Burkholderia_pseudomallei__TM_#01
ANTTGATDPLTGFNIGGTNAASI
>gb|ASE57902_1|ARO_3003731|fusD__Staphylococcus_saprophyticus__TM_#01
AHLVNAYNSVNDPNTIASIKDVTRDEILSTFNSRNTTIRSNVEKLMNVQLTKEQAQKILT
TIQMYVKPFEHPSNKQVTNL
>gb|ASE57902_1|ARO_3003731|fusD__Staphylococcus_saprophyticus__TM_#02
HFHHFLNKIQ
>gb|AVK94777_1|ARO_3004694|MCR-4_2__Escherichia_coli__TM_#01
AIFNLPLFGIVRK
>gb|AVK94777_1|ARO_3004694|MCR-4_2__Escherichia_coli__TM_#02
LASITNLLLTGLLPSYLIYKADIHYQPFFKELLHKLAFMLLMFVGIGIVAFFYYQDYAAF
>gb|AVK94777_1|ARO_3004694|MCR-4_2__Escherichia_coli__TM_#03
RNNSELRRYIVPTYFVSSASKYLNE
>gb|AVK94777_1|ARO_3004694|MCR-4_2__Escherichia_coli__TM_#04
YLQTPMEYQQLGLDAKNASRNPNT
>gb|AVK94777_1|ARO_3004694|MCR-4_2__Escherichia_coli__TM_#05
NAQDTVIDVLS
>gb|AVK94777_1|ARO_3004694|MCR-4_2__Escherichia_coli__TM_#06
KILAVAPSQDTVIF
>gb|AVK94777_1|ARO_3004694|MCR-4_2__Escherichia_coli__TM_#07
LAWVSNDFSQDNQLNMTCVAQR
>gb|AVK94777_1|ARO_3004694|MCR-4_2__Escherichia_coli__TM_#08
QSQLDIFAPCRY
>gb|WP_001572373_1|ARO_3004684|MCR-9__Enterobacteriales__TM_#01
FVFTGILPAILLFSIKIQYPEKWYKGIAYRLLSVL
>gb|WP_001572373_1|ARO_3004684|MCR-9__Enterobacteriales__TM_#02
MSCLENNAAK
>gb|WP_001572373_1|ARO_3004684|MCR-9__Enterobacteriales__TM_#03
STSVYNPDRDLFRECRG
>gb|ENV74419_1|ARO_3001735|OXA-280__Acinetobacter_johnsonii_ANC_3681__TM_#01
LITTFGSACT
>gb|ENV74419_1|ARO_3001735|OXA-280__Acinetobacter_johnsonii_ANC_3681__TM_#02
WEKDMTLG
>gb|ENV74419_1|ARO_3001735|OXA-280__Acinetobacter_johnsonii_ANC_3681__TM_#03
QRIGYGNQQIGT
>gb|ENV74419_1|ARO_3001735|OXA-280__Acinetobacter_johnsonii_ANC_3681__TM_#04
LFVEKLAN
>gb|ENV74419_1|ARO_3001735|OXA-280__Acinetobacter_johnsonii_ANC_3681__TM_#05
VQDMLLIEQK
>gb|ENV74419_1|ARO_3001735|OXA-280__Acinetobacter_johnsonii_ANC_3681__TM_#06
MKTGISASVREQLVKQSL
>gb|ABX54688_1|ARO_3002968|vanXYL__Enterococcus_faecalis__TM_#01
MDNDYKYYLQLVNKQYPWQINNGSKKMVRVPYTDKEIYLDAVVVEHLIQLIETIQLQEKI
EIVDGYRTIDEQKELWEFSLKDRGKRYTHDYVAYPGCSEHHTGLALDIGLKKTAHDIIAP
KFNGEEAKK
>gb|ABX54688_1|ARO_3002968|vanXYL__Enterococcus_faecalis__TM_#02
HTVGEKVS
>gb|BAB41258_1|ARO_3000124|mecI__Staphylococcus_aureus_subsp__aureus_N315__TM_#01
MDNKTYEISSAEWEVMNIIWMKKYASANNIIEEIQMQKDWSPKTIRTLITRLYKKGFIDR
KKDNKIFQYYSLVEESDIKYKTSKNFINKVYKGGFNSLVLNFVEKEDLSQDEIEELRNIL
NKK
>gb|APB03219_1|ARO_3003986|TaeA__Paenibacillus_sp__LC231__TM_#01
HEELQQKLLKLNQTLET
>gb|APB03219_1|ARO_3003986|TaeA__Paenibacillus_sp__LC231__TM_#02
QRHGGEASGSKAEASIAKSASGSSDGTDSDS
>gb|AAB21326_1|ARO_3002651|APH(3_)-Vc__Micromonospora_chalcea__TM_#01
MYAMLRRKYQHYEWTSVNEGDSGASVYRLAGQQPELYVKFAPREPENSAFDLAGEADRLT
WLTRHGIPVPCIVECGGDDTSVFLVTEAVT
>gb|AAB21326_1|ARO_3002651|APH(3_)-Vc__Micromonospora_chalcea__TM_#02
GGCPFDRSLAVTVA
>gb|ADA63498_1|ARO_3004678|rmtE__Escherichia_coli__TM_#01
AEVLSSKKYTSVDPAVVRRVCMETAPKYPKKKEAIKAVKNELHIIHEVFLQNECYKNALS
FLSQLSLDFNNAQLIDITMQIMQSHTSTKERLGDIEAVCSFLSTHISKEGSVMDIGCGFN
PFALPLLHEFPATYYAYDICSEGINILNKYFSILKKGEYRAELLDAVSVTPKEKVD
>gb|ADA63498_1|ARO_3004678|rmtE__Escherichia_coli__TM_#02
QQQKKGRGFSILEELDFDKAIVSFPIKSLGGKQKGMETFYSNLFEENLPSSLEIIEKQTF
SNEMFYVIQNKTKNGGNQS
>gb|APB03217_1|ARO_3003983|CatU__Paenibacillus_sp__LC231__TM_#01
RSNKCTFSITVDIDITRLLYSLKAN
>gb|APB03217_1|ARO_3003983|CatU__Paenibacillus_sp__LC231__TM_#02
VEFKTSFSPEGE
>gb|APB03217_1|ARO_3003983|CatU__Paenibacillus_sp__LC231__TM_#03
CLWTAFSNDFYRFHDH
>gb|APB03217_1|ARO_3003983|CatU__Paenibacillus_sp__LC231__TM_#04
KELQSLADSFEDWLT
>gb|BAE78083_1|ARO_3003549|mdtO__Escherichia_coli_str__K-12_substr__W3110__TM_#01
AGTLGMLSKVMRIPRQQEVTAL
>gb|ABA71730_1|ARO_3002965|vanWG__Enterococcus_faecalis__TM_#01
MIEVYKLTQ
>gb|ABA71730_1|ARO_3002965|vanWG__Enterococcus_faecalis__TM_#02

YFYFKMKFDGNRYAKKTSEKLLPN
>gb|ABA71730_1|ARO_3002965|vanWG__Enterococcus_faecalis__TM_#03
TINKVIIE
>gb|ABA71730_1|ARO_3002965|vanWG__Enterococcus_faecalis__TM_#04
LRNDTDNTFQIEISFDDNFMYGRILSQSSVNIEYTVFNSSVSYFKRE
>gb|AEZ36150_1|ARO_3000448|qepA__Klebsiella_aerogenes__TM_#01
LASMAALLA
>gb|AEZ36150_1|ARO_3000448|qepA__Klebsiella_aerogenes__TM_#02
RAALAAYALAA
>gb|AEZ36150_1|ARO_3000448|qepA__Klebsiella_aerogenes__TM_#03
VIGSLLSPQLAARWPAARILVVGLSAAAFGFAVLGLGQGLWWLVPATIV
>gb|AEZ36150_1|ARO_3000448|qepA__Klebsiella_aerogenes__TM_#04
FGSVGLVVYRQALTSAALPGLPADALQ
>gb|AKQ05898_1|ARO_3004590|tet(54)__uncultured_bacterium__TM_#01
YPVLIEKSAC
>gb|AKQ05898_1|ARO_3004590|tet(54)__uncultured_bacterium__TM_#02
VIYEQICKMRTQVECGRYVDVVGKT
>gb|AKQ05898_1|ARO_3004590|tet(54)__uncultured_bacterium__TM_#03
DEDVEILRGDLIEILMKTIDDVPCHFNQSIDSIEQNDDDVIVYFKDARVEHYDLIIGADG
IHSSIRR
>gb|AKQ05898_1|ARO_3004590|tet(54)__uncultured_bacterium__TM_#04
TNEKLVSITSDKDPKTAQVAFMFRSQHVWNNLRDEHEQMQFLRDIFHDFGWEAQKILELM
PNSH
>gb|AKQ05898_1|ARO_3004590|tet(54)__uncultured_bacterium__TM_#05
KVQIISNAIKLPEYE
>gb|AAB53259_1|ARO_3002671|Lactobacillus_reuteri_cat-TC__Lactobacillus_reuteri__TM_#01
FLVTRVINSNT
>gb|AAB53259_1|ARO_3002671|Lactobacillus_reuteri_cat-TC__Lactobacillus_reuteri__TM_#02
VSKTFSGIWT
>gb|AAB53259_1|ARO_3002671|Lactobacillus_reuteri_cat-TC__Lactobacillus_reuteri__TM_#03
IGLMTGFYNID
>gb|AAL85333_1|ARO_3004773|CKO-1__Citrobacter_koseri__TM_#01
MQEEQRLHARIGIAVLDTATNSITH
>gb|AAL85333_1|ARO_3004773|CKO-1__Citrobacter_koseri__TM_#02
AALLREVDRKALALSASMQFEPSQLVEYSPITEKHVAPDAMSW
>gb|AAL85333_1|ARO_3004773|CKO-1__Citrobacter_koseri__TM_#03
LNGPQAVTQFLRD
>gb|AAL85333_1|ARO_3004773|CKO-1__Citrobacter_koseri__TM_#04
QTLNTLLLGNVLQPSSRE
>gb|AAL85333_1|ARO_3004773|CKO-1__Citrobacter_koseri__TM_#05
DNGSRSIVSVVWPTSQKPLLVVIYITQTPATMAQRDAAIVRIGESLFSTLAVYD
>gb|AAB05628_1|ARO_3002950|vanXB__Enterococcus_faecalis__TM_#01
AVALALREAQIH
>gb|AAB05628_1|ARO_3002950|vanXB__Enterococcus_faecalis__TM_#02
GNAEAQNRRCLRKIMESSGFQSYRFEWWHYK
>gb|BAE85478_1|ARO_3003728|vanRI__Desulfitobacterium_hafniense_Y51__TM_#01
SAEPNQEEHV
>gb|WP_071593233_1|ARO_3005317|OXA-539__Pseudomonas_aeruginosa__TM_#01
MAIRIFAILFS
>gb|WP_071593233_1|ARO_3005317|OXA-539__Pseudomonas_aeruginosa__TM_#02
TFAHAQEG
>gb|WP_071593233_1|ARO_3005317|OXA-539__Pseudomonas_aeruginosa__TM_#03
ERSDWRKFFSEFQAKGTIVVADERQ
>gb|WP_071593233_1|ARO_3005317|OXA-539__Pseudomonas_aeruginosa__TM_#04
DKARRYLK
>gb|WP_071593233_1|ARO_3005317|OXA-539__Pseudomonas_aeruginosa__TM_#05
PSTSNGDY
>gb|WP_071593233_1|ARO_3005317|OXA-539__Pseudomonas_aeruginosa__TM_#06
SIEALPPNPAV
>gb|AVI44920_1|ARO_3004470|poxtA__Staphylococcus_aureus__TM_#01
MKGKNMNLAFGLEEIYEDAEFQIGDL
>gb|AVI44920_1|ARO_3004470|poxtA__Staphylococcus_aureus__TM_#02
LDNGSLTSGNARIGY
>gb|AVI44920_1|ARO_3004470|poxtA__Staphylococcus_aureus__TM_#03
KKYEQELEEIYKKLETAVNAEQEALLARMGTLQERLEYFDFYEAETILLEFADKMSIDAE
LYHRPMRE
>gb|AVI44920_1|ARO_3004470|poxtA__Staphylococcus_aureus__TM_#04
RIINKIMYINKATHKISVYDGDYYIYKKKYAEEQRIREMA
>gb|AVI44920_1|ARO_3004470|poxtA__Staphylococcus_aureus__TM_#05
GELQKRNRL
>gb|AVI44920_1|ARO_3004470|poxtA__Staphylococcus_aureus__TM_#06
QVPLEVENITFHYSGYPTLYQ
>gb|AVI44920_1|ARO_3004470|poxtA__Staphylococcus_aureus__TM_#07
DEGCIRFNQ
>gb|AVI44920_1|ARO_3004470|poxtA__Staphylococcus_aureus__TM_#08
ENKTVIDNVESEGYTPWQIRAV
>gb|AVI44920_1|ARO_3004470|poxtA__Staphylococcus_aureus__TM_#09
PYSRELLEYYYEINGSVAKF
>gb|AIA09786_1|ARO_3000861|rmtC__Pseudomonas_aeruginosa__TM_#01
MKTNDNYIEEVTAKVLTSGKYSTLYPPTVRRVTERLFDRYPPKQLEKEVRKKLHQAYGAY
IGGIDGKRLEKKIEKIIHEIPNPTTDEATRTEWEKEICLKILNLHTSTNERTVAYDELYQ
KIFEVTGVPTSITDAGCALNPFSFPFFTEAGMLGQYIGFDLDKGMIEAIEHSLRTLNAPE
GIVVKQGDILSDPSGESDLLLMFKLYTLLDRQEEASGLKILQEWKYKNAVISFPIKTISG
RDVGMEENYTVKFENDLVGSDLRIMQKLKLGNEMYFIVSRL
>gb|AAF61417_1|ARO_3002244|CARB-5__Acinetobacter_calcoaceticus_subsp__anitratus__TM_#01
HKASFFSVVITFLCLTLSLNANATDSVLEAVTNAETELGARIGLA
>gb|AAF61417_1|ARO_3002244|CARB-5__Acinetobacter_calcoaceticus_subsp__anitratus__TM_#02
HDLETGKRWEHKSNERFPL
>gb|AAF61417_1|ARO_3002244|CARB-5__Acinetobacter_calcoaceticus_subsp__anitratus__TM_#03
STFKTLACANVLQRVDLGKERIDRVVRFSESNLVTYSPVTEKHVGKKGMSLAELCQATLS
TSDNSAANFILQAIGGPKALTK
>gb|AAF61417_1|ARO_3002244|CARB-5__Acinetobacter_calcoaceticus_subsp__anitratus__TM_#04
IAMVTTLEKLLIDETLSIKSRQQLESWLKGNEVGDALFRKGVPSDWIVADRTGAGGYGSR
AITAVMWPPNRKPIVAALYITETDASFEERNAVIAKIGEQIAK
>gb|ACH58999_1|ARO_3002489|LRA-10__uncultured_bacterium_BLR10__TM_#01
MAIAILSSCFAPLASRAA
>gb|ACH58999_1|ARO_3002489|LRA-10__uncultured_bacterium_BLR10__TM_#02
RFTDALETNVLTQMGMKSTYVHVPQSAMANYAWGYDQANKPGRMNPGVLADGI
>gb|ACH58999_1|ARO_3002489|LRA-10__uncultured_bacterium_BLR10__TM_#03
TMAWEANPAQKLTPPSVPSG
>gb|ACH58999_1|ARO_3002489|LRA-10__uncultured_bacterium_BLR10__TM_#04
GPDRIKAAHAILEQL
>gb|AAC77110_1|ARO_3004290|Escherichia_coli_ampC__Escherichia_coli_str__K-12_substr__MG1655__TM_#01

WQILNALQ
>gb|CAI99385_1|ARO_3002856|dfrA24__Escherichia_coli__TM_#01
MTYQLDVSKILSFDLEAIVAATE
>gb|CAI99385_1|ARO_3002856|dfrA24__Escherichia_coli__TM_#02
QGDLKRFREITQGGIVIMGAGTYKSLPSPLKDRINIVITKKSEISWTACYDVRVVNSPED
ALRMVGRIIDEKEEQGRDRPRVFVIGGASIYQALMPFVSTLHWTEVHVEQLPEEIGLDTY
IEDFLSLRGTSTPKRKSNLVLPPTPTTP
>gb|ADG84870_1|ARO_3002858|dfrA12__Klebsiella_pneumoniae__TM_#01
MNSESVRIY
>gb|ADG84870_1|ARO_3002858|dfrA12__Klebsiella_pneumoniae__TM_#02
TLVISRQANYRATGCVVVSTLSHAIALASEL
>gb|ADG84870_1|ARO_3002858|dfrA12__Klebsiella_pneumoniae__TM_#03
ELVSTETIQAV
>gb|ADT70779_1|ARO_3001805|OXA-198__Pseudomonas_aeruginosa__TM_#01
MHKHMSKLFIAFLAFLLSVPAAAEDQTLAELFAQQGIDGTIVISSLHNGKTFIHNDPRAK
>gb|ADT70779_1|ARO_3001805|OXA-198__Pseudomonas_aeruginosa__TM_#02
KDDVLKWDGHIY
>gb|ADT70779_1|ARO_3001805|OXA-198__Pseudomonas_aeruginosa__TM_#03
NFLKKVHLRTLPFSASSYETLRQIML
>gb|ADT70779_1|ARO_3001805|OXA-198__Pseudomonas_aeruginosa__TM_#04
EKDLPLRQKLTRKALQAKGIIE
>gb|WP_058652112_1|ARO_3005346|DfrA34__Gammaproteobacteria__TM_#01
MITACVAIDSDGGFGAQGTLPWAIPEEFAFYQEHVRGGICIIGGRSFNDLVHLSLSPKGG
LYKKCLLRTTPHIVVSSSHELVYDPSIMALIEADRRHLDLYFVNTVDAAVKLAKGLGGMH
ANKDIHFIGGKRIYDAGLDYCDEVYTSILPAVYLNCDTFFPVEKLSRMFTPELYKTIPNQ
VHADIPVIKWTRKRA
>gb|ABU39979_1|ARO_3000217|blaR1__Bacillus_clausii__TM_#01
TVLTLPFIPIHLL
>gb|ABU39979_1|ARO_3000217|blaR1__Bacillus_clausii__TM_#02
TNKLTNLESPAIQTPMTFGWLKTYILLPKNIELYLSDDEIRH
>gb|ABU39979_1|ARO_3000217|blaR1__Bacillus_clausii__TM_#03
LTLGQREYKAYGQTIMRFLERNSRFLYLTNPLHSSKKAFKNTKSYNIAFFYWRVKSGAQL
KKPWVVFAGLTRFCIAQFPFLTATAVSTERYQFDESQAVVEDYSTYFAGNEGSFVLYSLT
SDQFEIYNKEKSVR
>gb|ABU39979_1|ARO_3000217|blaR1__Bacillus_clausii__TM_#04
GRDDSWLEWDGVEYEDEAWNSG
>gb|ABU39979_1|ARO_3000217|blaR1__Bacillus_clausii__TM_#05
ERIKQRNIQSFVNQLDYGNKDLSGGLN
>gb|ABU39979_1|ARO_3000217|blaR1__Bacillus_clausii__TM_#06
QLDFKEEHVQFVKEVMKLEENQKGTLYGKTGTGIVNGHA
>gb|ABU39979_1|ARO_3000217|blaR1__Bacillus_clausii__TM_#07
QKIMHMEARLLKSLYPFCQAKEF
>gb|CBY88906_1|ARO_3004105|TMB-1__Achromobacter_sp__sallbros1__TM_#01
MRPFLFLIIFISHFAFANEEIPGLEVEEIDNGVFLHKSYSRVEGW
>gb|CBY88906_1|ARO_3004105|TMB-1__Achromobacter_sp__sallbros1__TM_#02
EKLVDWIRSKKYEL
>gb|CBY88906_1|ARO_3004105|TMB-1__Achromobacter_sp__sallbros1__TM_#03
KSITTYASALTNEILKREGKEQARSSFKGNEFSLMDGFLEVYYPGGGHTIDNLVVWIPSS
K
>gb|CBY88906_1|ARO_3004105|TMB-1__Achromobacter_sp__sallbros1__TM_#04
SGLGYTGEAKIDQWPQSARNTISK
>gb|CBY88906_1|ARO_3004105|TMB-1__Achromobacter_sp__sallbros1__TM_#05
FELLKHTKVLAEKASNKANHGDR
>gb|AJF36617_1|ARO_3004596|erm(46)__Rhodococcus_hoagii__TM_#01
QKLQHRVD
>gb|AJF36617_1|ARO_3004596|erm(46)__Rhodococcus_hoagii__TM_#02
GRVSASAFT
>gb|AJF36617_1|ARO_3004596|erm(46)__Rhodococcus_hoagii__TM_#03
GQWSELFAIIDRRSPATDRKRAEYRR
>gb|ACN32294_1|ARO_3000237|tolC__Escherichia_coli__TM_#01
PQTPEQNAIADGYAPDSPAPVVQQTSARTTTSNGHNPFRN
>gb|AFQ93498_1|ARO_3002676|catB3__Enterobacter_cloacae__TM_#01
SFPFFYMQEEPAFS
>gb|AFQ93498_1|ARO_3002676|catB3__Enterobacter_cloacae__TM_#02
GNDVWIGSEAM
>gb|AFQ93498_1|ARO_3002676|catB3__Enterobacter_cloacae__TM_#03
GAVIGSRSLVT
>gb|BAA32494_1|ARO_3000449|FomB__Streptomyces_wedmorensis__TM_#01
MLENLTIRSSRVVDLNLVKVRLSTNLEDFAAYSYFSAFAEDES
>gb|BAA32494_1|ARO_3000449|FomB__Streptomyces_wedmorensis__TM_#02
DIPAELYADRTDRTFRGKRFKGGYYLVHYFGEPAHLITVGRT
>gb|BAA32494_1|ARO_3000449|FomB__Streptomyces_wedmorensis__TM_#03
DLTAHMESGDYVTDSSTLFESPQISTARVRNVVIVDYDPARPQGLMPISPAAAGT
>gb|BAA32494_1|ARO_3000449|FomB__Streptomyces_wedmorensis__TM_#04
DFDTYVDSLARMRAQLTELVEGARCYRANADMLAKEVRDSTLKQLAE
>gb|AAP08996_1|ARO_3000172|FosB__Bacillus_cereus_ATCC_14579__TM_#01
QEDFKCLIQRLEE
>gb|AAP08996_1|ARO_3000172|FosB__Bacillus_cereus_ATCC_14579__TM_#02
RDEKPHMTFY
>gb|AAG07594_1|ARO_3000808|mexI__Pseudomonas_aeruginosa_PAO1__TM_#01
DVRVDLAYETSRFIQA
>gb|AAG07594_1|ARO_3000808|mexI__Pseudomonas_aeruginosa_PAO1__TM_#02
HQNEGRMG
>gb|AAG07594_1|ARO_3000808|mexI__Pseudomonas_aeruginosa_PAO1__TM_#03
GQVLEFSLGHRWLTGGLALLVCISLPLLYSMPK
>gb|AAG07594_1|ARO_3000808|mexI__Pseudomonas_aeruginosa_PAO1__TM_#04
FTPQALARQFVRTQDGNLVPLSTVVRVA
>gb|AFV91534_1|ARO_3004786|FIM-1__Pseudomonas_aeruginosa__Partial_TM_#01
MRPLPHSYLKSLVICLLTAFAALTPVVNSGVQAAQPKDVPVTFTAITQGVWMHTSMKHME
NWGHVPSNGLIVEKGDFSILVDTAWDDPQTAQIIEWSKDTLKKPIRW
>gb|AFV91534_1|ARO_3004786|FIM-1__Pseudomonas_aeruginosa__Partial_TM_#02
AALRQQGIVTYAAADSNRMAPQNGLTPAEHDLIFDSEHSTSVLHPLVIFDPGPGHTRDNI
VVGLPEQGIVFGGCLIRPSGSTSLGNTADADLAHWKTAVLAVAQRFAEAQQIIPSHGPMA
GRELFELTAQLAEKASIPSTP
>gb|AAN06707_1|ARO_3000165|tet(A)__Shigella_sonnei__TM_#01
RNSSNSRCT
>gb|AHA41500_1|ARO_3002913|vanO__Rhodococcus_hoagii__TM_#01
KSAGIATPSFWVVAENEKVDADH
>gb|AHA41500_1|ARO_3002913|vanO__Rhodococcus_hoagii__TM_#02
MTMHGKKR
>gb|AEX49906_1|ARO_3003583|basS__Pseudomonas_aeruginosa__TM_#01
MSRAAVPSVRRRLLVNLLVGFVLCWLSVAALTYHLSLKQVNRLFDDDMVDFGEAALRLLD
LATEDQAGEDGSITEIIERSREAIQGLPLLRRESALGYALWRDGQPLLSSLNLPPEITAQ

```
GPGFSTV
>gb|AEX49906_1|ARO_3003583|basS__Pseudomonas_aeruginosa__TM_#02
LLGGLVWFGVARGLAPLREVQAEVQQRSARHLQPIAVEAVPLEIRGLIDELNLLLERLR
>gb|AEX49906_1|ARO_3003583|basS__Pseudomonas_aeruginosa__TM_#03
KAHARGLL
>gb|AEX49906_1|ARO_3003583|basS__Pseudomonas_aeruginosa__TM_#04
ETVHVMGIDLWLKA
>gb|AEX49906_1|ARO_3003583|basS__Pseudomonas_aeruginosa__TM_#05
RVENRAQHAVLRVRDNGPGVALEEQQAIFTRFYRSPATS
>gb|AEX49906_1|ARO_3003583|basS__Pseudomonas_aeruginosa__TM_#06
FGSIGLGKGLEGKGLEVQVFLPKTQPDATRPPARGPDSGRSHI
>gb|BAM62793_1|ARO_3002233|IMP-42__Acinetobacter_soli__TM_#01
KPYGLGNL
>gb|AAD01868_1|ARO_3002867|dfrF__Enterococcus_faecalis__TM_#01
KGEQKQFRELTT
>gb|AAD01868_1|ARO_3002867|dfrF__Enterococcus_faecalis__TM_#02
PNRMNIVVSTTTEYQGDNLVSVKSLEDALLLAKGR
>gb|AAD01868_1|ARO_3002867|dfrF__Enterococcus_faecalis__TM_#03
TFFPEFDINDFEVLIGETLGEEVKYTRTFYVRKNELSRFWI
>gb|AAA65958_1|ARO_3002955|vanYA__Enterococcus_faecium__TM_#01
VEFQNYDQNPKEHLENSGTSENTQEKT
>gb|AAA65958_1|ARO_3002955|vanYA__Enterococcus_faecium__TM_#02
GSTHTNSRR
>gb|AAL27445_1|ARO_3002924|vanRE__Enterococcus_faecalis__TM_#01
VSLLSTLLSNEGYEVYEAMSGKESLEI
>gb|AAL27445_1|ARO_3002924|vanRE__Enterococcus_faecalis__TM_#02
SIREKQFFPVLMLTARG
>gb|AAL27445_1|ARO_3002924|vanRE__Enterococcus_faecalis__TM_#03
SQSIDETNEYAKNGLNISVNSRKVFL
>gb|AHA41499_1|ARO_3002948|vanHO__Rhodococcus_hoagii__TM_#01
MSYRDLGLIDSEVIAERRVRALDDSSPSAVPTTGVRVFGCGHDEAVLFREMGTRLGITPS
ITEEAIS
>gb|CAL48457_1|ARO_3002857|dfrA26__Escherichia_coli__TM_#01
MADEEYDPLLDDDMEDAKVAVIAARAQNGCIGRHGK
>gb|CAL48457_1|ARO_3002857|dfrA26__Escherichia_coli__TM_#02
NGALPGRTNIVVTRQQGYEAEGARVVDSIEEAISLAQSIALIEAVDEIMVLGGGEIYTQA
LPQADILYLTEVHASVDGDAFFPDVDLSQYQETQRQDFEPSGGNPYPFSFVVYQRT
>gb|AEM66528_1|ARO_3001809|OXA-209__Riemerella_anatipestifer__TM_#01
MKKTFILLNLILLVNLNGYCQTKSLKSNEIVKPEFR
>gb|AEM66528_1|ARO_3001809|OXA-209__Riemerella_anatipestifer__TM_#02
RYLKKLNYRGMVFDTLTIDQFWLEGE
>gb|AAP40270_1|ARO_3001671|OXA-49__Acinetobacter_baumannii__TM_#01
LSGCTVQHNLINET
>gb|AAP40270_1|ARO_3001671|OXA-49__Acinetobacter_baumannii__TM_#02
SQLAHTQLPFSEKVQAN
>gb|AAP40270_1|ARO_3001671|OXA-49__Acinetobacter_baumannii__TM_#03
GWVEQPDGKIVAFAL
>gb|AAP40270_1|ARO_3001671|OXA-49__Acinetobacter_baumannii__TM_#04
MEMRSEMPASIRNELLMKSLKQLNII
>gb|AAB05626_1|ARO_3002943|vanHB__Enterococcus_faecalis__TM_#01
NAFRTLSPDFHIIPTLISDAISAD
>gb|AAB05626_1|ARO_3002943|vanHB__Enterococcus_faecalis__TM_#02
ATILALRK
>gb|AAB05626_1|ARO_3002943|vanHB__Enterococcus_faecalis__TM_#03
STIHAVAQQNFRLDC
>gb|AAB05626_1|ARO_3002943|vanHB__Enterococcus_faecalis__TM_#04
SRKIEADYVQ
>gb|AAB05626_1|ARO_3002943|vanHB__Enterococcus_faecalis__TM_#05
CADTRHLIGQSEIGE
>gb|AAB05626_1|ARO_3002943|vanHB__Enterococcus_faecalis__TM_#06
GSLVEALGS
>gb|AAB05626_1|ARO_3002943|vanHB__Enterococcus_faecalis__TM_#07
QFVYTDCSQKVLDHPFLSQLL
>gb|APB03221_1|ARO_3003988|AAC(2_)-IIb__Paenibacillus_sp__LC231__TM_#01
MNHRKGNEPTAAALMELHVLAMFTHDGNMQIRTINEPWPGEEL
>gb|APB03221_1|ARO_3003988|AAC(2_)-IIb__Paenibacillus_sp__LC231__TM_#02
GIAGQLRALVEDEPIVTEEVLTRPKHFAAYMNLLRA
>gb|APB03221_1|ARO_3003988|AAC(2_)-IIb__Paenibacillus_sp__LC231__TM_#03
QTTQAKQTVRITPGNIREYSLTGFEWLTTEIDYD
>gb|APB03221_1|ARO_3003988|AAC(2_)-IIb__Paenibacillus_sp__LC231__TM_#04
RAHEAGLETS
>gb|APB03221_1|ARO_3003988|AAC(2_)-IIb__Paenibacillus_sp__LC231__TM_#05
GNSSSRRVANKLGLSYYGVNFTIS
>gb|AHE40557_1|ARO_3002626|ANT(6)-Ia__Exiguobacterium_sp__S3-2__TM_#01
GKNITRYTEDMYKKYVENDYF
>gb|ALH22601_1|ARO_3000620|adeL__Acinetobacter_baumannii__TM_#01
TARILADVADIESSFHDAERGPRGQ
>gb|ALH22601_1|ARO_3000620|adeL__Acinetobacter_baumannii__TM_#02
IYLEKYGEPTSIEDLQKNHK
>gb|CDO61516_1|ARO_3003949|efrB__Enterococcus_faecium__Partial_TM_#01
LDTCRIFRRNAKSPVNIAKHAKFIRRLKWLCSENMTGFSVLKLYVRKKKPLKGFKQVNHR
FKWF
>gb|CAA52904_1|ARO_3002688|catS__Streptococcus_pyogenes__Partial_TM_#01
MAYANDMQRYGSNYGMIGKPDI
>gb|AAX55643_1|ARO_3003558|y56_beta-lactamase__Yersinia_enterocolitica__TM_#01
SLPTWAAAIPGSLDKQLAALEHS
>gb|AAX55643_1|ARO_3003558|y56_beta-lactamase__Yersinia_enterocolitica__TM_#02
SATKIILSQIS
>gb|CCI79240_1|ARO_3005047|eptB__Klebsiella_pneumoniae_subsp__rhinoscleromatis_SB3432__TM_#01
MPQRQAVFYSSFSRFAC
>gb|AFM55000_1|ARO_3001692|OXA-229__Acinetobacter_bereziniae__TM_#01
MKFKMKGLF
>gb|AFM55000_1|ARO_3001692|OXA-229__Acinetobacter_bereziniae__TM_#02
ILSSLAFSGCVYDSKLQRPVISER
>gb|AFM55000_1|ARO_3001692|OXA-229__Acinetobacter_bereziniae__TM_#03
ERKQLTLS
>gb|AET10444_1|ARO_3004032|tetA(46)__Streptococcus_australis__TM_#01
FSTVLLLLFGGQSLASGQ
>gb|AET10444_1|ARO_3004032|tetA(46)__Streptococcus_australis__TM_#02
EFLVNQQPIVDYN
>gb|AET10444_1|ARO_3004032|tetA(46)__Streptococcus_australis__TM_#03
KKGASQEDLMEAVAQAAFADDLERMSH
>gb|CAC47933_1|ARO_3005166|Tet(X1)__Bacteroides_thetaiotaomicron__TM_#01
```

SIGLEQENGKWLLHFENKPTALADFIIVSNGGMSKI
>gb|CAC47933_1|ARO_3005166|Tet(X1)__Bacteroides_thetaiotaomicron__TM_#02
TNCPEFYKLCNN
>gb|CAC47933_1|ARO_3005166|Tet(X1)__Bacteroides_thetaiotaomicron__TM_#03
NNGKGLDFKPTKSVSEFLTNRF
>gb|CAC47933_1|ARO_3005166|Tet(X1)__Bacteroides_thetaiotaomicron__TM_#04
TIKIFPLDK
>gb|CAC47933_1|ARO_3005166|Tet(X1)__Bacteroides_thetaiotaomicron__TM_#05
ADSTKNEIEMRNPSFTFQQLMNV
>gb|AAC36944_1|ARO_3003006|blt__Bacillus_subtilis_subsp__subtilis_str__168__TM_#01
MKKSINEQKTIF
>gb|AAC36944_1|ARO_3003006|blt__Bacillus_subtilis_subsp__subtilis_str__168__TM_#02
VTSVFILKESLSIEERHQLSSHTKESNFIK
>gb|AAC36944_1|ARO_3003006|blt__Bacillus_subtilis_subsp__subtilis_str__168__TM_#03
RMIQLCLITGAILAFVSTVMSG
>gb|AAC36944_1|ARO_3003006|blt__Bacillus_subtilis_subsp__subtilis_str__168__TM_#04
KNDAAALN
>gb|AAC43550_1|ARO_3003967|lrfA__Mycolicibacterium_smegmatis_MC2_155__TM_#01
MSTCIEGTPSTTRTPT
>gb|AAC43550_1|ARO_3003967|lrfA__Mycolicibacterium_smegmatis_MC2_155__TM_#02
LAAFAPSTEL
>gb|AAC43550_1|ARO_3003967|lrfA__Mycolicibacterium_smegmatis_MC2_155__TM_#03
TGLVFVAVGFLMILLFRHNLTVAAIIASFV
>gb|AAC43550_1|ARO_3003967|lrfA__Mycolicibacterium_smegmatis_MC2_155__TM_#04
MLVLAAAAVVGVAFRR
>gb|ACU86041_1|ARO_3004539|mphH__Brachybacterium_faecium_DSM_4810__TM_#01
MPEDLDALLDLAARHGLDLDGGT
>gb|ACU86041_1|ARO_3004539|mphH__Brachybacterium_faecium_DSM_4810__TM_#02
LERAAVEGRLLAMLAPHLDVAVPDWRIS
>gb|ACU86041_1|ARO_3004539|mphH__Brachybacterium_faecium_DSM_4810__TM_#03
AADGEVSWHVDMASTVYARSLGSVVAQLHAVDAEAAAATGIEVRSPAQVRGAWRQDLARV
GAEFE
>gb|ACU86041_1|ARO_3004539|mphH__Brachybacterium_faecium_DSM_4810__TM_#04
LMFHQVSAPSAIFEVALQAYAEGGGRPWPGLARHCTEMFSAAPLGYGLYALATGEAAHRE
AAAAALNPPEER
>gb|AAC78336_1|ARO_3002826|EreA2__Providencia_stuartii__TM_#01
MTWRTTRTLLQPQKL
>gb|AAC78336_1|ARO_3002826|EreA2__Providencia_stuartii__TM_#02
LARASLIRY
>gb|AAC78336_1|ARO_3002826|EreA2__Providencia_stuartii__TM_#03
FNAIGLECGAIQASRLSEWLNSTAGAHELERFSDTLTFS
>gb|AAC78336_1|ARO_3002826|EreA2__Providencia_stuartii__TM_#04
IDHLMKPHVD
>gb|AAC78336_1|ARO_3002826|EreA2__Providencia_stuartii__TM_#05
LKLRLASLAPVLK
>gb|AAC78336_1|ARO_3002826|EreA2__Providencia_stuartii__TM_#06
FRKASDRIESIEYTLETLR
>gb|AAC78336_1|ARO_3002826|EreA2__Providencia_stuartii__TM_#07
FFDGTSLEGDTSVRDSYMAGVVDG
>gb|AAC78336_1|ARO_3002826|EreA2__Providencia_stuartii__TM_#08
QYVIDACG
>gb|AAF09244_1|ARO_3000849|SFH-1__Serratia_fonticola__TM_#01
TLGAKIVATQMTYDLQKSQWGSIVNFTRQGNNKYPNLEKSLPDTVFPGDFNLQNGSIRAM
YLGEAHTKDGIFVYFPAERVLYGNCILKENLGNMSFANRTEYPKTLEKLKGLIEQGELKV
DSIIAGHDTPIHDVGLIDHYLTLLEKAPK
>gb|AAG05187_1|ARO_3005067|ParS__Pseudomonas_aeruginosa_PAO1__TM_#01
AAAIFVVDHVIDAFYDSIVENYHRDAVRGQAYSLVEKLAPLDQAGRQRQLEDWRPHYGLE
LSLTDARQAKLTQEEQALLDKNLLVVREDFTEFISRIDAGPQLLDIKLPPEPSLTPLFTV
LAYILLGVLVGIALLVWVRPHWRDLETLRLAAQRFGDGDLSSRTRIFRRSDIRT
>gb|AAG05187_1|ARO_3005067|ParS__Pseudomonas_aeruginosa_PAO1__TM_#02
NKQVDAEVRHDLIE
>gb|AAG05187_1|ARO_3005067|ParS__Pseudomonas_aeruginosa_PAO1__TM_#03
RLEHGNVGSHREI
>gb|AAG05187_1|ARO_3005067|ParS__Pseudomonas_aeruginosa_PAO1__TM_#04
HSRVEIALLDQGDSCQIRVNDDGPGIPADA
>gb|AAG05187_1|ARO_3005067|ParS__Pseudomonas_aeruginosa_PAO1__TM_#05
GGYAEALETPQGGASFRLTWERPR
>gb|BAL40892_1|ARO_3004768|CIA-1__Chryseobacterium_indologenes_NBRC_14944__TM_#01
LMVSAFATAQKSV
>gb|BAL40892_1|ARO_3004768|CIA-1__Chryseobacterium_indologenes_NBRC_14944__TM_#02
MADKGKFST
>gb|BAL40892_1|ARO_3004768|CIA-1__Chryseobacterium_indologenes_NBRC_14944__TM_#03
VVQHFMDSKGAKDLQIK
>gb|BAL40892_1|ARO_3004768|CIA-1__Chryseobacterium_indologenes_NBRC_14944__TM_#04
ESSTNATVS
>gb|BAL40892_1|ARO_3004768|CIA-1__Chryseobacterium_indologenes_NBRC_14944__TM_#05
TRMVSDISKIVWDNFNK
>gb|CAA11707_1|ARO_3000463|gimA__Streptomyces_ambofaciens__TM_#01
MRRGDLHETYRLDYAPHMH
>gb|ACD56151_1|ARO_3002848|arr-3__Escherichia_coli__TM_#01
EPLRIVGVVEDWEGHPVELIR
>gb|AVX52225_1|ARO_3004516|MCR-8__Klebsiella_pneumoniae__TM_#01
MFKYLLSFKLNPVQRT
>gb|AVX52225_1|ARO_3004516|MCR-8__Klebsiella_pneumoniae__TM_#02
INVDVHNIHNLLFFASLPIFLFCFLSILLTPVMVIPYLCRPLLVVLILISAC
>gb|AVX52225_1|ARO_3004516|MCR-8__Klebsiella_pneumoniae__TM_#03
FLSTLFLLGIVPAIILALPSTDNKRGAFRIELWWLAHICIAVVLLAM
>gb|AVX52225_1|ARO_3004516|MCR-8__Klebsiella_pneumoniae__TM_#04
QASSTILQSVGEDAVRPIYSNAP
>gb|AVX52225_1|ARO_3004516|MCR-8__Klebsiella_pneumoniae__TM_#05
NEYNEVRAASEENLLDILK
>gb|AVX52225_1|ARO_3004516|MCR-8__Klebsiella_pneumoniae__TM_#06
KRVPTDDMPAMKVIGECVNKDGTCFDEVLLNQLSSRIN
>gb|AVX52225_1|ARO_3004516|MCR-8__Klebsiella_pneumoniae__TM_#07
TSKVFSPTCDSNLIEKCSNK
>gb|AVX52225_1|ARO_3004516|MCR-8__Klebsiella_pneumoniae__TM_#08
NEQTHIPMFMWFSSSFAQHSKLNLECLTGNADKQYSHDNFYHSILGLFNVKTSVYKPELD
MFTLCRQSDHTPLSSAVVREKTDGN
>gb|ACH58991_1|ARO_3002484|LRA-13__uncultured_bacterium_BLR13__TM_#01
MNFRHIVMAALCGLAWTPAIHATEVCIAIAEAGTGA
>gb|ACH58991_1|ARO_3002484|LRA-13__uncultured_bacterium_BLR13__TM_#02
KLPFRPGYV
>gb|ACH58991_1|ARO_3002484|LRA-13__uncultured_bacterium_BLR13__TM_#03

KSLGMQRFADYTRNFKYGNADVSGDAENDGLSMS
>gb|ACH58991_1|ARO_3002484|LRA-13__uncultured_bacterium_BLR13__TM_#04
DKIVNRRLGVSAHAYDMTAQLTKFD
>gb|ACH58991_1|ARO_3002484|LRA-13__uncultured_bacterium_BLR13__TM_#05
DNGKNYFFNYGLASRETGQAVTSH
>gb|ACH58991_1|ARO_3002484|LRA-13__uncultured_bacterium_BLR13__TM_#06
MAATLTSYAQVNGQLALTDTVSRHMPKLRGGG
>gb|ACH58991_1|ARO_3002484|LRA-13__uncultured_bacterium_BLR13__TM_#07
VAAGGARTYSNLTVGLLGIITAQSMGMPFAEAMENRLFPQ
>gb|ACH58991_1|ARO_3002484|LRA-13__uncultured_bacterium_BLR13__TM_#08
QANAPVRINPAVLATEAYGVKTDAA
>gb|ACH58991_1|ARO_3002484|LRA-13__uncultured_bacterium_BLR13__TM_#09
EKLQRAVTGTHTAYFKTGELTQDLIWEQYPAASKLDRMLAGVSEKMVFESNPATRLA
>gb|ACH58991_1|ARO_3002484|LRA-13__uncultured_bacterium_BLR13__TM_#10
GAERVTAAWHILDQLDQR
>gb|CAA64891_1|ARO_3000812|mtrE__Neisseria_gonorrhoeae__TM_#01
MNTTLKTTLTSVAAAFALSACTMIPQYEQPKVEVAETFQNDTSVSSIRAVDLGWH
>gb|CAA64891_1|ARO_3000812|mtrE__Neisseria_gonorrhoeae__TM_#02
SLAQRVLKTREETYNAVRIAVQGRRDFRRRPAPA
>gb|AYV52072_1|ARO_3002882|lmrD__Lactococcus_lactis__TM_#01
TDFFTHQHAGQSQDALQQIAQQLSQQMHQTVDWHNVPEVVKSLPQAAQDQITANLPKGTT
LETLKTVAT
>gb|AYV52072_1|ARO_3002882|lmrD__Lactococcus_lactis__TM_#02
VTFAWLTVAASPVAILSAVIIIRQSKKATDK
>gb|CAB13167_1|ARO_3003064|ykkD__Bacillus_subtilis_subsp__subtilis_str__168__TM_#01
MLHWISLLCAGCLEMAGVALMNQYAKEKSVKWVLLIIVGFAASFSLLSYAME
>gb|CAB13167_1|ARO_3003064|ykkD__Bacillus_subtilis_subsp__subtilis_str__168__TM_#02
GALIGILFYKEQKDAKRIFFIALILCSAVGLKILS
>gb|ACH59002_1|ARO_3002483|LRA-5__uncultured_bacterium_BLR5__TM_#01
MKTIFGKRRQSAVVLITLIAILLASGQPYQSSQVRGAACLPDIIFDEPSQGPEKNEAISM
LTERLSSIINAAGGDIGIAVIHVETGHTTAIQGTTQ
>gb|ACH59002_1|ARO_3002483|LRA-5__uncultured_bacterium_BLR5__TM_#02
WTANAAMWRRPIDRTVAQLIEVSII
>gb|ACH59002_1|ARO_3002483|LRA-5__uncultured_bacterium_BLR5__TM_#03
AAVTHRMRALGFPNIEIVSTVREFSENRTRPNTGSAEDLARLLVQLQKGELLQPQHSALL
LGFMHRATTGTERLRGSLPVGTPVADKTGTGDAGVVTNDVGIITLPKGQGHLAIAVLISG
SKLSPA
>gb|ABV26707_1|ARO_3002850|arr-5__Klebsiella_pneumoniae__TM_#01
ELAIGDLISTGFISHFERDRALKH
>gb|ABV26707_1|ARO_3002850|arr-5__Klebsiella_pneumoniae__TM_#02
VALSGSDGP
>gb|ABV26707_1|ARO_3002850|arr-5__Klebsiella_pneumoniae__TM_#03
HPLKIVGILREWERHSPEALKT
>gb|AEO25219_1|ARO_3004651|lin__Listeria_monocytogenes_FSL_R2-561__TM_#01
TQGKLAYIKQLSTDT
>gb|AEO25219_1|ARO_3004651|lin__Listeria_monocytogenes_FSL_R2-561__TM_#02
PQAKIAYFDQELNGLNQTKSLLENISE
>gb|AEO25219_1|ARO_3004651|lin__Listeria_monocytogenes_FSL_R2-561__TM_#03
TFAEYEESKA
>gb|CAB12108_2|ARO_3003059|tmrB__Bacillus_subtilis_subsp__subtilis_str__168__TM_#01
KDDFQSYPLWRAFNYSLLASLTDTYRGILIVPMTIVHPEYFNEIIGRLRQEGRIVHHFTL
MASKETLL
>gb|CAB12108_2|ARO_3003059|tmrB__Bacillus_subtilis_subsp__subtilis_str__168__TM_#02
SPIFEDHIQTDNLSIQDVAENIAARAELPLD
>gb|ADB54781_1|ARO_3004556|dfrB7__Aeromonas_hydrophila__Partial_TM_#01
MGDRVRKKSGAAWQGQ
>gb|AHJ02283_1|ARO_3003186|CARB-23__Vibrio_parahaemolyticus_UCM-V493__TM_#01
MVRVFTRYSLLNIAKVRIKTK
>gb|AHJ02283_1|ARO_3003186|CARB-23__Vibrio_parahaemolyticus_UCM-V493__TM_#02
LNEDISLIEK
>gb|AHJ02283_1|ARO_3003186|CARB-23__Vibrio_parahaemolyticus_UCM-V493__TM_#03
IDERNIVVWSPVMDKL
>gb|AHJ02283_1|ARO_3003186|CARB-23__Vibrio_parahaemolyticus_UCM-V493__TM_#04
NYGSRGISAMIWKDNYKPVYISIYVTDTDLSLQARDQLIAQISQLILEHYKES
>gb|CAD32565_1|ARO_3001770|OXA-43__Burkholderia_pseudomallei__TM_#01
MKFRHALSSAFVLLGCIAASAHAKTICTAIADAGTGKLLVQDGDCGR
>gb|CAD32565_1|ARO_3001770|OXA-43__Burkholderia_pseudomallei__TM_#02
DPVLPYRDSYIA
>gb|CAD32565_1|ARO_3001770|OXA-43__Burkholderia_pseudomallei__TM_#03
RKLPVSPTAVDMTERIVESTTLADGTVVHGKTGVSYPLLADGTRDWARGSGWFVGWIVRG
>gb|CAD32565_1|ARO_3001770|OXA-43__Burkholderia_pseudomallei__TM_#04
QTLVFARLT
>gb|ACH58994_1|ARO_3002512|LRA-17__uncultured_bacterium_BLR17__TM_#01
MSIFRTILFVSILLLTSLANSPHATAQVTNTDR
>gb|ACH58994_1|ARO_3002512|LRA-17__uncultured_bacterium_BLR17__TM_#02
AALAGTVDQVKAN
>gb|ACH58994_1|ARO_3002512|LRA-17__uncultured_bacterium_BLR17__TM_#03
MTGAKVMIDDQDAPVVEDGGNSDYIYGGKGVGSLFAPVHVDRKLHDHDNITLGGT
>gb|ACH58994_1|ARO_3002512|LRA-17__uncultured_bacterium_BLR17__TM_#04
YMLSEVTLPGMPTYPNVGKDFMYTYGAMRKLQFDIWVAAHSSQFGLQDVRKETDGYNPGA
FGDKKKYLTTIDKTEDIYKEHFKGGK
>gb|AAA25334_1|ARO_3002543|AAC(3)-IXa__Micromonospora_chalcea__TM_#01
MEEMSLLNHSGGPVTRSRIKHDLADLGLK
>gb|AAA25334_1|ARO_3002543|AAC(3)-IXa__Micromonospora_chalcea__TM_#02
YNNGRLPEALP
>gb|AAA25334_1|ARO_3002543|AAC(3)-IXa__Micromonospora_chalcea__TM_#03
LVNGQRVWRQFRDIDSEEGAFDYSTVRRGV
>gb|AAA25334_1|ARO_3002543|AAC(3)-IXa__Micromonospora_chalcea__TM_#04
GPVFNFAINWIEAKLKR
>gb|AAL27446_1|ARO_3002935|vanSE__Enterococcus_faecalis__TM_#01
ITTILVILPLVAKMFLSLRVWQGTEFFYPILYILNRSLGVWLIVTPLFIWLIVTYM
>gb|AAL27446_1|ARO_3002935|vanSE__Enterococcus_faecalis__TM_#02
TPNEKIVLRVELAEFENEINHIRIDSLENKKMAEEAG
>gb|AAL27446_1|ARO_3002935|vanSE__Enterococcus_faecalis__TM_#03
QTYLELDSTKRKNYIDIISDKANRLEHLLNDFFEIAKTGKSREEVY
>gb|AAL27446_1|ARO_3002935|vanSE__Enterococcus_faecalis__TM_#04
LNDTTIKLTLEKVDEKVVVSVGNITDKVSEKDIDQLFEPFYRGDK
>gb|AAL27446_1|ARO_3002935|vanSE__Enterococcus_faecalis__TM_#05
DFKVSIIL
>gb|AAK99679_1|ARO_3000822|pmrA__Streptococcus_pneumoniae_R6__TM_#01
FYSVIIYLLCANASSPLQLG
>gb|AML99881_1|ARO_3001327|mdtK__Salmonella_enterica_subsp__enterica_serovar_Typhimurium__TM_#01
SEIINIVAGEATP

>gb|ATL63235_1|ARO_3004357|catV__Brevibacillus_brevis_Vm4__TM_#01
MKFQRIDLDNWSRRSYFEHYLNRVN
>gb|ATL63235_1|ARO_3004357|catV__Brevibacillus_brevis_Vm4__TM_#02
TSFHADGELGYWESMIPSYTFFHQDDQTFSTMWTEFA
>gb|ATL63235_1|ARO_3004357|catV__Brevibacillus_brevis_Vm4__TM_#03
KYGDNKGLVAKELEPPYTFP
>gb|ATL63235_1|ARO_3004357|catV__Brevibacillus_brevis_Vm4__TM_#04
WATNYKEWLGVE
>gb|AAG08359_1|ARO_3003682|OpmH__Pseudomonas_aeruginosa_PAO1__TM_#01
NPLVHYGKYVDER
>gb|AAG08359_1|ARO_3003682|OpmH__Pseudomonas_aeruginosa_PAO1__TM_#02
QSKPRQQY
>gb|AAA23018_1|ARO_3004454|Campylobacter_coli_chloramphenicol_acetyltransferase__Campylobacter_coli__TM_#01
MQFTKIDINNWTRKEYFDHYFG
>gb|AAA23018_1|ARO_3004454|Campylobacter_coli_chloramphenicol_acetyltransferase__Campylobacter_coli__TM_#02
GVTTIINRHEEFRTALDENGQVGVFSEMLPC
>gb|AAA23018_1|ARO_3004454|Campylobacter_coli_chloramphenicol_acetyltransferase__Campylobacter_coli__TM_#03
SIWTEFTADYTEFLQNYQKDIDAFGERMGMS
>gb|ACC85616_1|ARO_3002804|FosA2__Enterobacter_cloacae__TM_#01
KSVTFWHELLGLTLHAR
>gb|ACC85616_1|ARO_3002804|FosA2__Enterobacter_cloacae__TM_#02
AEDFEPFSHKLEQAGVT
>gb|AAL19304_1|ARO_3000791|mdsC__Salmonella_enterica_subsp__enterica_serovar_Typhimurium_str__LT2__TM_#01
DASRTPADLSVMDESEINRRYEAA
>gb|AAL19304_1|ARO_3000791|mdsC__Salmonella_enterica_subsp__enterica_serovar_Typhimurium_str__LT2__TM_#02
RAHVQKQIEIARYENVIQQAFRDVADGLAGQR
>gb|AAG07066_1|ARO_3003710|mexL__Pseudomonas_aeruginosa_PAO1__TM_#01
KAKCAEQLPALYFQLAEGAPLEK
>gb|AAG07066_1|ARO_3003710|mexL__Pseudomonas_aeruginosa_PAO1__TM_#02
HEAIALTRLMAAQAGQNPKLSELFFEAGPKQVIDEMERLLEQARRSGKLAFPDARHAAEH
FFMLVKGCANYRLLIGCAEPLDEAEGERHVEEVVALFLRAFAAGG
>gb|CAM96571_1|ARO_3004605|ermZ__Streptomyces_ambofaciens__TM_#01
MTLKSPLPPQSVSAPADSRST
>gb|CAM96571_1|ARO_3004605|ermZ__Streptomyces_ambofaciens__TM_#02
GSDTIPPD
>gb|CAM96571_1|ARO_3004605|ermZ__Streptomyces_ambofaciens__TM_#03
PGTPLLAVEIDPR
>gb|CAM96571_1|ARO_3004605|ermZ__Streptomyces_ambofaciens__TM_#04
GQPVRLIGNLPFV
>gb|CAM96571_1|ARO_3004605|ermZ__Streptomyces_ambofaciens__TM_#05
DMGPARMR
>gb|CAM96571_1|ARO_3004605|ermZ__Streptomyces_ambofaciens__TM_#06
RGLAFSRQDFTPVPRADTQTLMVAPHRRPSVPWREKAAYQRFVQRV
>gb|AAF86691_1|ARO_3001816|ACC-2__Hafnia_alvei__TM_#01
MRKKMQNTLK
>gb|AAF86691_1|ARO_3001816|ACC-2__Hafnia_alvei__TM_#02
LSVITCLA
>gb|AAF86691_1|ARO_3001816|ACC-2__Hafnia_alvei__TM_#03
LIQPLMQKNNIPGMSVAVT
>gb|AAF86691_1|ARO_3001816|ACC-2__Hafnia_alvei__TM_#04
NYIYNYGLA
>gb|AAF86691_1|ARO_3001816|ACC-2__Hafnia_alvei__TM_#05
KQPQQPVT
>gb|AAF86691_1|ARO_3001816|ACC-2__Hafnia_alvei__TM_#06
SHYVPELRGSSFDH
>gb|AAF86691_1|ARO_3001816|ACC-2__Hafnia_alvei__TM_#07
YLKAWKPAD
>gb|AAF86691_1|ARO_3001816|ACC-2__Hafnia_alvei__TM_#08
AGTHRVYSNIGTGLLGMIAA
>gb|AAF86691_1|ARO_3001816|ACC-2__Hafnia_alvei__TM_#09
ENYAWGYNKK
>gb|AAF86691_1|ARO_3001816|ACC-2__Hafnia_alvei__TM_#10
LGNEAYGI
>gb|AAF86691_1|ARO_3001816|ACC-2__Hafnia_alvei__TM_#11
RYVQANMGQLKLD
>gb|AAA26615_1|ARO_3004457|Staphylococcus_intermedius_chloramphenicol_acetyltransferase__Staphylococcus_intermedius__TM_#01
NQQTTYSITKEIDITLFKDMIKKKGYEIYPSLIYAIMEVVNKNKVFRTGINSENKLGYWD
KLNPLYTVFNKQTEKFTNIWTESD
>gb|AAA26615_1|ARO_3004457|Staphylococcus_intermedius_chloramphenicol_acetyltransferase__Staphylococcus_intermedius__TM_#02
SFYNNYKNDL
>gb|AAA26615_1|ARO_3004457|Staphylococcus_intermedius_chloramphenicol_acetyltransferase__Staphylococcus_intermedius__TM_#03
EYKDKEEMFPKK
>gb|AAA26615_1|ARO_3004457|Staphylococcus_intermedius_chloramphenicol_acetyltransferase__Staphylococcus_intermedius__TM_#04
ENNKIYIPVALQLHH
>gb|AAA26615_1|ARO_3004457|Staphylococcus_intermedius_chloramphenicol_acetyltransferase__Staphylococcus_intermedius__TM_#05
VCDGYHASLF
>gb|ALF06101_1|ARO_3004764|CBP-1__Clostridium_botulinum__TM_#01
MKKIVNSKLKVNKFKMCIFISILIFSLTGCGNVENKTSENTKPEIQ
>gb|ALF06101_1|ARO_3004764|CBP-1__Clostridium_botulinum__TM_#02
NADKRFAYCSTFKSLISGAILQKYSSDQLKQVIKYSPKDVLSYAPVTKNHVDKGMTIE
>gb|ALF06101_1|ARO_3004764|CBP-1__Clostridium_botulinum__TM_#03
SIDLKEYTTGNILSDDKKKILINWMS
>gb|CAA27061_1|ARO_3002648|APH(3_)-IVa__Bacillus_circulans__TM_#01
MNESTRNWPEELLELLGQTELTVNKIGYSGDHVYHVKEYRGTPAFLKIAPSVWWRTLRPE
IEALAWLDGKLPVPKILYTAEHGGMDYLLMEALGGKDGSHETIQAKRKLFVKLYAEGLRS
VHGLDIRECPLSNGLEKKLRDAKRIVDESLVDPADIKEEYDCTPEELYGLLLESKPVTED
LVFAHGDYCAPNLIIDGE
>gb|CAA27061_1|ARO_3002648|APH(3_)-IVa__Bacillus_circulans__TM_#02
RHDYGDDRYKALFLELYGLDGLDEDKVRYYIRLDEFF
>gb|NP_249397_1|ARO_3002679|Pseudomonas_aeruginosa_catB7__Pseudomonas_aeruginosa_PAO1__TM_#01
PLLCTGDIPALYRHWKQRQATA
>gb|CAA26199_1|ARO_3004089|ANT(3__)-IIa__Escherichia_coli__TM_#01
MVTAQWRFSWLLVMT
>gb|CAA26199_1|ARO_3004089|ANT(3__)-IIa__Escherichia_coli__TM_#02
FDPVPEQDL
>gb|AAA99503_1|ARO_3003250|bcrC__Bacillus_licheniformis__TM_#01
MSFSELNIDAFRF
>gb|AAA99503_1|ARO_3003250|bcrC__Bacillus_licheniformis__TM_#02
LTRTTKNRLMVIYAVIAFVVAEILGKIMGSLHSNYQPFAT
>gb|AAA99503_1|ARO_3003250|bcrC__Bacillus_licheniformis__TM_#03
GFLIFLFH
>gb|AAA99503_1|ARO_3003250|bcrC__Bacillus_licheniformis__TM_#04
HQMLSLYEKVEQRIVPSKNKSNDKSKNF

>gb|APB03220_1|ARO_3003987|VatI__Paenibacillus_sp__LC231__TM_#01
MTGPNPNERYPIPGDN
>gb|APB03220_1|ARO_3003987|VatI__Paenibacillus_sp__LC231__TM_#02
DIEVINQYIGAIVSGDMDLLRKMRQN
>gb|ACL82961_1|ARO_3002911|vanM__Enterococcus_faecium__TM_#01
AEIANNIDIG
>gb|ACL82961_1|ARO_3002911|vanM__Enterococcus_faecium__TM_#02
TCEKPCIDWDNEHCR
>gb|ACL82961_1|ARO_3002911|vanM__Enterococcus_faecium__TM_#03
MQDKGYQIQR
>gb|ACL82961_1|ARO_3002911|vanM__Enterococcus_faecium__TM_#04
QVIYKDDKPAADS
>gb|ACL82961_1|ARO_3002911|vanM__Enterococcus_faecium__TM_#05
ERIKEAAKN
>gb|ACL82961_1|ARO_3002911|vanM__Enterococcus_faecium__TM_#06
VSAGITIPELIDHLIVLAVKE
>gb|CAA44667_1|ARO_3001305|ErmU__Streptomyces_lincolnensis__TM_#01
MPSRYGSRQDLGQNFLVDPDIIKLIRRAPNERKVPSLIWRRRGH
>gb|CAA44667_1|ARO_3001305|ErmU__Streptomyces_lincolnensis__TM_#02
SARAPENVKVVGEDILRFRLPTVPHTVVGNIPFHVTTATMRRILVAPAWVS
>gb|CAA44667_1|ARO_3001305|ErmU__Streptomyces_lincolnensis__TM_#03
CSLVTAESWPWFDFSVLKRVPRFAFR
>gb|CAA44667_1|ARO_3001305|ErmU__Streptomyces_lincolnensis__TM_#04
RERREYQA
>gb|CAA44667_1|ARO_3001305|ErmU__Streptomyces_lincolnensis__TM_#05
QRIGRVQDSDLSAWFRAHG
>gb|CAA44667_1|ARO_3001305|ErmU__Streptomyces_lincolnensis__TM_#06
GMARGGRSVPRTRRPRGLPPRTSRGPRRNSG
>gb|AAZ04784_1|ARO_3002711|QnrA5__Shewanella_algae__TM_#01
MDIIDKVFQQEDFSRQDL
>gb|AAZ04784_1|ARO_3002711|QnrA5__Shewanella_algae__TM_#02
DASFEDCSFIESGA
>gb|AAZ04784_1|ARO_3002711|QnrA5__Shewanella_algae__TM_#03
AYISGCNLAY
>gb|AAZ04784_1|ARO_3002711|QnrA5__Shewanella_algae__TM_#04
DLTFADLDGLDPRRVNLE
>gb|BAA14147_1|ARO_3004667|Staphylococcus_aureus_norA__Staphylococcus_aureus__TM_#01
HDPKKSTTSGFQKLEPQLLTKINWKVFITPVILTLVLSFGLSAFETL
>gb|BAA14147_1|ARO_3004667|Staphylococcus_aureus_norA__Staphylococcus_aureus__TM_#02
AWSLLYSVVVLILLVF
>gb|BAA14147_1|ARO_3004667|Staphylococcus_aureus_norA__Staphylococcus_aureus__TM_#03
APIYMAIGVSLAGVVIVLIEKQHRAKLKEQNM
>gb|AAV85982_1|ARO_3000535|macB__Neisseria_gonorrhoeae__TM_#01
IIAKQSYVASATPM
>gb|AAV85982_1|ARO_3000535|macB__Neisseria_gonorrhoeae__TM_#02
VKDKLFADS
>gb|ACS83748_1|ARO_3000573|tet(43)__uncultured_bacterium_AOTet43__TM_#01
MPPSHHMLRPIEQCSILWNDVRYSNSVRLKEAGMTATTQASAPA
>gb|ACS83748_1|ARO_3000573|tet(43)__uncultured_bacterium_AOTet43__TM_#02
LAFGLISLTVIAVA
>gb|ACS83748_1|ARO_3000573|tet(43)__uncultured_bacterium_AOTet43__TM_#03
LGVPADDAEGLTKDRLDVK
>gb|ACS83748_1|ARO_3000573|tet(43)__uncultured_bacterium_AOTet43__TM_#04
AVTWACGIGAAAALMGFV
>gb|ACS83748_1|ARO_3000573|tet(43)__uncultured_bacterium_AOTet43__TM_#05
MAVIGTLIAVL
>gb|ACS83748_1|ARO_3000573|tet(43)__uncultured_bacterium_AOTet43__TM_#06
TTTLPNGD
>gb|ACS83748_1|ARO_3000573|tet(43)__uncultured_bacterium_AOTet43__TM_#07
SLDLATSYFAGERIAYLFLAVVVGVIAGWGALTLSNSKEMEDVH
>gb|BAA03776_1|ARO_3000316|mphA__Escherichia_coli__TM_#01
AKVEPEARVLAMLKNRLPFAVPDWRVANAELVAYPMLEDS
>gb|BAA03776_1|ARO_3000316|mphA__Escherichia_coli__TM_#02
KVADDVDRVRREFVVNDKRL
>gb|BAA03776_1|ARO_3000316|mphA__Escherichia_coli__TM_#03
DLYVGHVLIDNTER
>gb|AJP77085|ARO_3003717|ESP-1__Chryseobacterium_sp__Stok-2__TM_#01
MKKLTLILSLIIAFFGYAQTVTEPTNHSAE
>gb|ACH58988_1|ARO_3002487|LRA-8__uncultured_bacterium_BLR8__TM_#01
MSKSSLKGLVLLALVAAIAAPSWAARKEKPAAKAPPCEQCAVWNADQEPFKIW
>gb|ACH58988_1|ARO_3002487|LRA-8__uncultured_bacterium_BLR8__TM_#02
VYQRRPSDQVLRTGKPDPGDPQLARAGPIPPVENVWVVHDEELLGLGPTRFT
>gb|ACH58988_1|ARO_3002487|LRA-8__uncultured_bacterium_BLR8__TM_#03
AQCLKIVYADSLNAVSAEGFRFTASTTYPNVLQDLEQSFKRVESLPCDVIVSVHPEQSDF
FPRMAKRVDGKPESIKDPEGCKRYVAGARERLALRVASEKQGS
>gb|ACL82962_1|ARO_3002953|vanXM__Enterococcus_faecium__TM_#01
MEKGFTFLDEILND
>gb|ACL82962_1|ARO_3002953|vanXM__Enterococcus_faecium__TM_#02
YELADALLKVQELAFN
>gb|ACL82962_1|ARO_3002953|vanXM__Enterococcus_faecium__TM_#03
YSCFDFPVK
>gb|AAA22597_1|ARO_3000495|ErmD__Bacillus_anthracis__TM_#01
MKKKNHKYR
>gb|AAA22597_1|ARO_3000495|ErmD__Bacillus_anthracis__TM_#02
EEIVDRANISI
>gb|AAA22597_1|ARO_3000495|ErmD__Bacillus_anthracis__TM_#03
NDSKFVDILTRKTAQHS
>gb|AAA22597_1|ARO_3000495|ErmD__Bacillus_anthracis__TM_#04
GIVREISKEH
>gb|AAA22597_1|ARO_3000495|ErmD__Bacillus_anthracis__TM_#05
DAPLSHKHYIAFRGLAEYALKEPNIPLCVRLRGIFTPR
>gb|AAA22597_1|ARO_3000495|ErmD__Bacillus_anthracis__TM_#06
TVGTLTENQWAVIFNTMTQYVMHHKWPRANKRKPGEI
>gb|BAA11236_1|ARO_3000206|emrK__Escherichia_coli__TM_#01
GSVTVVNHKDTNYVRQGDILVSLDKTDATIALNKAKNNLANIVRQTNKLYLQDKQYSAEV
ASARIQYQQSLEDYNRRVPLAKQGVISKETLEHTKDTLISSKAALNAAIQAYKANKALVM
NTPLNRQPQVVEAADATKEAWLALKR
>gb|BAA11236_1|ARO_3000206|emrK__Escherichia_coli__TM_#02
IGQSVNIISDLYGE
>gb|BAA11236_1|ARO_3000206|emrK__Escherichia_coli__TM_#03
KNEDIAEMPELASTVTSMPAYT
>gb|BAA11236_1|ARO_3000206|emrK__Escherichia_coli__TM_#04
SNIISHNGQL

>gb|AAG07762_1|ARO_3003030|mexV__Pseudomonas_aeruginosa_PAO1__TM_#01
MLLRRMLIMLAAVIAVVAILAGYKVYSIRQQIALFSAPKPPISVTASLAEK
>gb|AAG07762_1|ARO_3003030|mexV__Pseudomonas_aeruginosa_PAO1__TM_#02
DQVKLDQPLIQLESD
>gb|AAG07762_1|ARO_3003030|mexV__Pseudomonas_aeruginosa_PAO1__TM_#03
RAEYQRGRELIGSKAISKSEFDRLAAQWAKT
>gb|AAG07762_1|ARO_3003030|mexV__Pseudomonas_aeruginosa_PAO1__TM_#04
HLPEQDFPLLSRGQLVKVRVAAYPAQVFDAE
>gb|AAG07762_1|ARO_3003030|mexV__Pseudomonas_aeruginosa_PAO1__TM_#05
EQGQVSKDDKGQPQQVVERRFVRIGERREGLAVVL
>gb|AAD25538_1|ARO_3000174|tet(G)__Pseudomonas_sp___TM_#01
CAPLLGQFSDG
>gb|AAD25538_1|ARO_3000174|tet(G)__Pseudomonas_sp___TM_#02
ETGKLVRIE
>gb|AAD25538_1|ARO_3000174|tet(G)__Pseudomonas_sp___TM_#03
AGPLGFTALYSATI
>gb|AAM09851_1|ARO_3002923|vanRD__Enterococcus_faecium__TM_#01
GDKIMGLSV
>gb|AAM09851_1|ARO_3002923|vanRD__Enterococcus_faecium__TM_#02
PCIKQEAERTEYDIRGMTISKS
>gb|CAA24743_1|ARO_3002655|APH(4)-Ia__Escherichia_coli__TM_#01
MKKPELTATSVEKFLIEKFDSVSD
>gb|CAA24743_1|ARO_3002655|APH(4)-Ia__Escherichia_coli__TM_#02
SCADGFYKDRYVYRHFASAALPIPEVLDIGEFSESL
>gb|CAA24743_1|ARO_3002655|APH(4)-Ia__Escherichia_coli__TM_#03
VAEAMDAIAAADLSQTSGFGPFGPQGIGQYTTWRDFICAIADPHVYHWQTVMDDTVSASV
AQALDEL
>gb|CAA24743_1|ARO_3002655|APH(4)-Ia__Escherichia_coli__TM_#04
YFERRHPELAGSPRLRAYMLRIGLDQLYQSLVDGNFDDAAWAQGRCDAIVRSGAGTVGRT
QIARRSAAVWTDGCVEVLADSGNRRPSTRPRAKE
>gb|AAA19882_1|ARO_3003562|blaF__Mycolicibacterium_fortuitum__TM_#01
VMMTRSQAH
>gb|CAC99773_1|ARO_3003770|Listeria_monocytogenes_mprF__Listeria_monocytogenes_EGD-e__TM_#01
MKEKLMQAYAWFQKNSTVVKIVFITFVM
>gb|CAC99773_1|ARO_3003770|Listeria_monocytogenes_mprF__Listeria_monocytogenes_EGD-e__TM_#02
PSLKENLTSQSPEQIF
>gb|CAC99773_1|ARO_3003770|Listeria_monocytogenes_mprF__Listeria_monocytogenes_EGD-e__TM_#03
GSGLLLILSSAVPNAIYHVPF
>gb|CAC99773_1|ARO_3003770|Listeria_monocytogenes_mprF__Listeria_monocytogenes_EGD-e__TM_#04
PIAIAKNGEGTIVGFASMMPSYTDEM
>gb|CAC99773_1|ARO_3003770|Listeria_monocytogenes_mprF__Listeria_monocytogenes_EGD-e__TM_#05
EKAKEDGFQTFNA
>gb|CAC99773_1|ARO_3003770|Listeria_monocytogenes_mprF__Listeria_monocytogenes_EGD-e__TM_#06
ANSQVVLDFPLEEETKKPDSE
>gb|CAF05908_1|ARO_3000845|GIM-1__Pseudomonas_aeruginosa__TM_#01
MKNVLVFLILLVALPALAQGHKPLEVIKIEDGVYLHTSFKNI
>gb|CAF05908_1|ARO_3000845|GIM-1__Pseudomonas_aeruginosa__TM_#02
LLLSWATDRGYQVM
>gb|CAF05908_1|ARO_3000845|GIM-1__Pseudomonas_aeruginosa__TM_#03
KKLLAREGKPVPTHYFKDDEFTLGNGLIELYYPGA
>gb|CAF05908_1|ARO_3000845|GIM-1__Pseudomonas_aeruginosa__TM_#04
HEWEGLGYVGDASISSWADSIKNIVSKKYPIQ
>gb|CAF05908_1|ARO_3000845|GIM-1__Pseudomonas_aeruginosa__TM_#05
SSDILDHTIDLAESASNKLMQPTAEASAD
>gb|AAC60780_1|ARO_3003049|rosB__Yersinia_enterocolitica_(type_O_8)__TM_#01
NETTSLSQLF
>gb|AAC60780_1|ARO_3003049|rosB__Yersinia_enterocolitica_(type_O_8)__TM_#02
NVYSTHHFCPWRKSVNLP
>gb|AAC60780_1|ARO_3003049|rosB__Yersinia_enterocolitica_(type_O_8)__TM_#03
GRFVQSCTVSRLWSG
>gb|AAC60780_1|ARO_3003049|rosB__Yersinia_enterocolitica_(type_O_8)__TM_#04
SADIMSLARLDCALVI
>gb|AAB41958_1|ARO_3000802|OprJ__Pseudomonas_aeruginosa__TM_#01
GAAAQRQGAAIDT
>gb|AAB41958_1|ARO_3000802|OprJ__Pseudomonas_aeruginosa__TM_#02
QAIPRSPGQR
>gb|AAB41958_1|ARO_3000802|OprJ__Pseudomonas_aeruginosa__TM_#03
EGRSLVVHRGGRS
>gb|AAA22277_1|ARO_3003007|bmr__Bacillus_subtilis__TM_#01
FLLTTVHSYVA
>gb|AAA22277_1|ARO_3003007|bmr__Bacillus_subtilis__TM_#02
TLAIGIALTIAWKAPAHLKAST
>gb|AEL31272_1|ARO_3002756|QnrB41__Citrobacter_freundii__TM_#01
THCDLTNSELGDLD
>gb|AEL31272_1|ARO_3002756|QnrB41__Citrobacter_freundii__TM_#02
VDLQGVKLD
>gb|BAA03674_1|ARO_3001300|myrA__Micromonospora_griseorubida__TM_#01
MHPDLLPHLRCPVCGQPLHQADAAPP
>gb|BAA03674_1|ARO_3001300|myrA__Micromonospora_griseorubida__TM_#02
TAAARAVPRRVRPGDGVGEPV
>gb|BAA03674_1|ARO_3001300|myrA__Micromonospora_griseorubida__TM_#03
DSLTRHFEPAGQSTHRHRLQLTR
>gb|BAA03674_1|ARO_3001300|myrA__Micromonospora_griseorubida__TM_#04
RVAALSEPVTVTAAVRLARYRPI
>gb|ABG77965_1|ARO_3004042|Enterobacter_cloacae_acrA__Enterobacter_cloacae__TM_#01
NRYQKLLGT
>gb|AAN63648_1|ARO_3000844|TUS-1_beta-lactamase__Myroides_odoratus_DSM_2801__TM_#01
MYHYFSSLFVLIFSTLVYPQSDKLKIEP
>gb|AAN63648_1|ARO_3000844|TUS-1_beta-lactamase__Myroides_odoratus_DSM_2801__TM_#02
TFTFGSTKQFNLGKEKIE
>gb|AAN63648_1|ARO_3000844|TUS-1_beta-lactamase__Myroides_odoratus_DSM_2801__TM_#03
VEAWPTTIKAVKRKFKK
>gb|WP_188331861_1|ARO_3005345|Trimethoprim-resistant_dihydrofolate_reductase_DfrA43__Proteus_penneri__TM_#01
MHMKMNIIVAMHEASRG
>gb|WP_188331861_1|ARO_3005345|Trimethoprim-resistant_dihydrofolate_reductase_DfrA43__Proteus_penneri__TM_#02
ETRASIVSNTPGCMAFASLELCLQYLRQLHPSTIVFAIGGSSLYKEILAMQMLCERIYMT
LVSGGPKTHSFDTFFPEIDETVYSKRICGGSGEHDDWKYKFVIYERPTSESVQSIETISQ
GH
>gb|CAA90432_1|ARO_3003026|fusH__Streptomyces_lividans__TM_#01
LHTGEFRVDSV
>gb|CAA90432_1|ARO_3003026|fusH__Streptomyces_lividans__TM_#02
TQSADVDGDGRA
>gb|AAG05914|ARO_3004075|MuxC__Pseudomonas_aeruginosa_PAO1__TM_#01

```
SQYGLSLDSVRTAIAAANSNGPKGAVEKDDKHWQVDANDQLR
>gb|AAG05914|ARO_3004075|MuxC__Pseudomonas_aeruginosa_PAO1__TM_#02
LKRPEGASLARRSDRFFAAFMLRYRASLGWALEHSRLMVVIMLACIAMNLW
>gb|AAG05914|ARO_3004075|MuxC__Pseudomonas_aeruginosa_PAO1__TM_#03
SAKMGEYRKILSS
>gb|AAG05914|ARO_3004075|MuxC__Pseudomonas_aeruginosa_PAO1__TM_#04
EKVLTRLRERI
>gb|AAG05914|ARO_3004075|MuxC__Pseudomonas_aeruginosa_PAO1__TM_#05
KLPQLVDVNSDSQDKGVQTRLVIDRDRAATLGINVEM
>gb|CAD12227_2|ARO_3000478|tet(33)__Corynebacterium_glutamicum__TM_#01
MSSLTSARGSLATVLITASLDAAGMGLVMPILPALLHEAGVTADAVPLNVGVLI
>gb|CAD12227_2|ARO_3000478|tet(33)__Corynebacterium_glutamicum__TM_#02
HQRAKRFGLLSACYGGGMIAGPAMGGLFGAISPHLPFLLAALLSASNLALTFILLRETRP
DSPARSASLAQHRGRPGLSAVPGITFLLIAFGLVQF
>gb|CAD12227_2|ARO_3000478|tet(33)__Corynebacterium_glutamicum__TM_#03
DWSPVEVGISLSVFGIVQVLVQALLTGRIVEWIGEAKTVIIGCITDALGLVGLAIVTDAF
SMAPILAALGIGGIGLPALQTLLSQRVDEQHQGRLQGVLASINSVTSIFGPVAFTTIFAL
TYINADGFLWLCAAALYVPCVILIMRGTAA
>gb|AAF36805_1|ARO_3002958|vanYF__Paenibacillus_popilliae_ATCC_14706__TM_#01
KDTSDDKMTA
>gb|AAF36805_1|ARO_3002958|vanYF__Paenibacillus_popilliae_ATCC_14706__TM_#02
RVDGKKYTISYD
>gb|AKQ05894_1|ARO_3004584|tet(50)__uncultured_bacterium__TM_#01
MTKHIKILVIGVGVAGPAVA
>gb|AKQ05894_1|ARO_3004584|tet(50)__uncultured_bacterium__TM_#02
KCGRYVDVKGNVLHEEQGET
>gb|AKQ05894_1|ARO_3004584|tet(50)__uncultured_bacterium__TM_#03
EFKQSVIKIEQNEDSVTVTY
>gb|AKQ05894_1|ARO_3004584|tet(50)__uncultured_bacterium__TM_#04
GLDHMELLCESNHKLVTLQSDSQADKAMAGFMFRSKHVLEDIRDEQEQKHFLHAS
>gb|AKQ05894_1|ARO_3004584|tet(50)__uncultured_bacterium__TM_#05
LKDDEVSKEIAEARSNKILAMIKSVSNSINLPQYE
>gb|QHW12375_1|ARO_3005087|msrF__Macrococcus_canis__TM_#01
KVNRYIDYGYFEQVESPKVSMADPRLLGKLNVKVDSNNLSGGEL
>gb|QHW12375_1|ARO_3005087|msrF__Macrococcus_canis__TM_#02
MIKVYSGNYSEYLRQKKVEREQQAHDFDL
>gb|QHW12375_1|ARO_3005087|msrF__Macrococcus_canis__TM_#03
LQRAAKAVEKRIEQLEVVDAPKEIHTIQFHQTHSTPPLHNKFPILGDRLTLQA
>gb|QHW12375_1|ARO_3005087|msrF__Macrococcus_canis__TM_#04
VIGFYEQMGYQFNEDKT
>gb|QHW12375_1|ARO_3005087|msrF__Macrococcus_canis__TM_#05
EYEGTIILISHDKKFVDHVSDTIYKIENKKLNLVN
>gb|CAH14033_1|ARO_3004100|LpeB__Legionella_pneumophila_str__Paris__TM_#01
QPEKLFSFGVNVDEIVNALAKNR
>gb|CAH14033_1|ARO_3004100|LpeB__Legionella_pneumophila_str__Paris__TM_#02
ANSKHPILLKSLANVALETDNSQMRVRVNGHAGVVLSINKANEANPIEVSKEIRKVIKGL
QQGLPKDLKINT
>gb|CAH14033_1|ARO_3004100|LpeB__Legionella_pneumophila_str__Paris__TM_#03
SASSKNWWPQFDNALEKLTKHYSNILQFILKHQKITLLTALISVVACFGFYNLIS
>gb|CAH14033_1|ARO_3004100|LpeB__Legionella_pneumophila_str__Paris__TM_#04
GMLDNKTGKLEKKLDAIPEANNRLTFIGDWGGSIVLPLKPHAQRHRSANQIVEK
>gb|CAH14033_1|ARO_3004100|LpeB__Legionella_pneumophila_str__Paris__TM_#05
HFRQLFDETEKLKS
>gb|CAH14033_1|ARO_3004100|LpeB__Legionella_pneumophila_str__Paris__TM_#06
TGGKKRCSKE
>gb|ABL75133_1|ARO_3004856|SCO-1__Acinetobacter_baumannii__TM_#01
MTRSALLIPLTTAAIALNAISPVYASDTHSIDDTVKQVETTLGAKVGIAVLDTGSQRAWF
>gb|ABL75133_1|ARO_3004856|SCO-1__Acinetobacter_baumannii__TM_#02
DKGQSFMNKEALIKKADLDEYAPVTSGIVGKKVSAADLCSI
>gb|ABL75133_1|ARO_3004856|SCO-1__Acinetobacter_baumannii__TM_#03
QAVTAYLRK
>gb|ABL75133_1|ARO_3004856|SCO-1__Acinetobacter_baumannii__TM_#04
ETLNKLVLGPTLGSDERKQLTTWLESNEVGD
>gb|ABL75133_1|ARO_3004856|SCO-1__Acinetobacter_baumannii__TM_#05
NGTRGVIAVMWPPKHAPIIAAIYITQTKATMEERNAAIASIGKAIAAEVLE
>gb|ADZ12699_1|ARO_3005091|RanA__Riemerella_anatipestifer_RA-GD__TM_#01
MIEVKDLRKSFND
>gb|ADZ12699_1|ARO_3005091|RanA__Riemerella_anatipestifer_RA-GD__TM_#02
KGVKEWEGNKDLIIKAENEHLIDFVYSSALFKQVRETMLRNNQTNL
>gb|BAO21229_1|ARO_3003199|AAC(6_)-Iak__Stenotrophomonas_maltophilia__TM_#01
AWAQLRLGLWPDADDPLE
>gb|BAO21229_1|ARO_3003199|AAC(6_)-Iak__Stenotrophomonas_maltophilia__TM_#02
GAVFLACAA
>gb|BAO21229_1|ARO_3003199|AAC(6_)-Iak__Stenotrophomonas_maltophilia__TM_#03
GVGRALLAAV
>gb|BAO21229_1|ARO_3003199|AAC(6_)-Iak__Stenotrophomonas_maltophilia__TM_#04
AWTRDAGCRELASDSRVEDVQ
>gb|APZ75411_1|ARO_3004861|NDM-18__Escherichia_coli__TM_#01
GLIVRDGGRVL
>gb|APZ75411_1|ARO_3004861|NDM-18__Escherichia_coli__TM_#02
DQTAQILNWIKQEINLPVAL
>gb|APZ75411_1|ARO_3004861|NDM-18__Escherichia_coli__TM_#03
NGWVEPATAPNF
>gb|AAF36806_1|ARO_3002963|vanZF__Paenibacillus_popilliae_ATCC_14706__TM_#01
GIGNVWVVGRYETLIRVSEINLLPFSSE
>gb|AAF36806_1|ARO_3002963|vanZF__Paenibacillus_popilliae_ATCC_14706__TM_#02
YTRDEKKLDNKSSLVIK
>gb|ABR14060_1|ARO_3002838|lnuD__Streptococcus_uberis__TM_#01
MVNKADAIEIILYAEEN
>gb|ABR14060_1|ARO_3002838|lnuD__Streptococcus_uberis__TM_#02
GKTFIEILKEKGFTEVIEAYTTTD
>gb|ABR14060_1|ARO_3002838|lnuD__Streptococcus_uberis__TM_#03
QGDLVFEGESYPSN
>gb|ABR14060_1|ARO_3002838|lnuD__Streptococcus_uberis__TM_#04
HDENDVHDVRLLCERYNIPVPSEYK
>gb|ENV32314_1|ARO_3001763|OXA-308__Acinetobacter_gerneri_DSM_14967___CIP_107464__TM_#01
MNKKLNLALLCFLSILCAACQSNQQLSANSHTENHNTRAAEISLLFDEM
>gb|ENV32314_1|ARO_3001763|OXA-308__Acinetobacter_gerneri_DSM_14967___CIP_107464__TM_#02
QHFQSYGNA
>gb|ENV32314_1|ARO_3001763|OXA-308__Acinetobacter_gerneri_DSM_14967___CIP_107464__TM_#03
SRGQSKLFAKSGLSMKNGQPDIGWYTGWVEQADGKIVAFSINMQMVQGLDVNSRQQATLD
>gb|AAK97755_1|ARO_3000167|tet(C)__Aeromonas_salmonicida__TM_#01
LMQESHKGERRPMP
```

```
>gb|CAH35802_1|ARO_3002983|amrB__Burkholderia_pseudomallei_K96243__TM_#01
SIEKAADNAQIIVSLTSEDGRLSG
>gb|CAH35802_1|ARO_3002983|amrB__Burkholderia_pseudomallei_K96243__TM_#02
DGFFGWFNRFVARSTHRYTRRVGRVLE
>gb|CAH35802_1|ARO_3002983|amrB__Burkholderia_pseudomallei_K96243__TM_#03
RVEEYVRTHS
>gb|CAH35802_1|ARO_3002983|amrB__Burkholderia_pseudomallei_K96243__TM_#04
ERKRARDQVQAIIAEINAHFAGTPNTMVFAINMPALPDLGL
>gb|CAH35802_1|ARO_3002983|amrB__Burkholderia_pseudomallei_K96243__TM_#05
GAFVAAREKLLAEGR
>gb|BAM10414_1|ARO_3003039|oprA__Pseudomonas_aeruginosa__TM_#01
MPLSKLSASSLALCLGLLGACSLAPRYQRPEAPIPTTYPAVPASQQAGDRARLDDWQQQF
TDPVLRQMIGQ
>gb|BAM10414_1|ARO_3003039|oprA__Pseudomonas_aeruginosa__TM_#02
ASERLPTLEASGRYERERMRGETREAGEVEQRYRVAA
>gb|BAM10414_1|ARO_3003039|oprA__Pseudomonas_aeruginosa__TM_#03
GGYVQERALYAQQRLAERTLHARENGLALVRKRYAAGMSTRIDLRSEEMLVESARATHAA
LVRERSQAVSGLQLLLGDFTGDWQDSQLDLEHLQLQ
>gb|BAM10414_1|ARO_3003039|oprA__Pseudomonas_aeruginosa__TM_#04
DLGSASSGLHGLFRGGSRVWTFS
>gb|BAM10414_1|ARO_3003039|oprA__Pseudomonas_aeruginosa__TM_#05
NRYEESIQVAFREVADALSAGDQLELQLRAQRAVRDADRERLQLVRK
>gb|BAM10414_1|ARO_3003039|oprA__Pseudomonas_aeruginosa__TM_#06
QLIHLRGLRLNNGVALYRALGGGWSQ
>gb|AAA26652_1|ARO_3002835|lnuA__Staphylococcus_haemolyticus__TM_#01
VIQKLEDIGYKIEVHWMPSRMELKHEEYGYLDIHPINLNDDGSITQANPEGGNYVFQNDW
FSE
>gb|AAA26652_1|ARO_3002835|lnuA__Staphylococcus_haemolyticus__TM_#02
TDHFDIKNLKSIT
>gb|AAM09852_1|ARO_3003070|vanXD__Enterococcus_faecium__TM_#01
ELGAALRKAQKA
>gb|AAM09852_1|ARO_3003070|vanXD__Enterococcus_faecium__TM_#02
LPENNLTKKRYYPNIKRNEMITKGYVASQSSHSRGSAIDLTIFR
>gb|AAM09852_1|ARO_3003070|vanXD__Enterococcus_faecium__TM_#03
VRSHHAASGLSEEEAGNRERLRDIMER
>gb|AQV34023_1|ARO_3004637|qnrE2__Escherichia_coli__TM_#01
AIFRNCDFSGTDLTSSEFIGCQFYDRASQQGGNFNRAQ
>gb|CCN31428_1|ARO_3005049|CrcB__Klebsiella_pneumoniae_subsp__pneumoniae_Ecl8__TM_#01
VVPRLSDGFPLDIFIANIIAAFLLGLCTSLYKRNRVNQYIHM
>gb|CCN31428_1|ARO_3005049|CrcB__Klebsiella_pneumoniae_subsp__pneumoniae_Ecl8__TM_#02
SGAVEMMNEPLSALIAIC
>gb|CCN31428_1|ARO_3005049|CrcB__Klebsiella_pneumoniae_subsp__pneumoniae_Ecl8__TM_#03
RLGSRVKPAPPMTHNRSTG
>gb|AIG22448_1|ARO_3001558|OXA-372__Citrobacter_freundii__TM_#01
HIFLVFLILCSNFALAEDKAISAIFSTEGVDGTIILKSLRGDKTITHNDARASRRFAS
>gb|AIG22448_1|ARO_3001558|OXA-372__Citrobacter_freundii__TM_#02
AVQENVVSLSGTAFRWDGKT
>gb|AIG22448_1|ARO_3001558|OXA-372__Citrobacter_freundii__TM_#03
EETYRRYL
>gb|AIG22448_1|ARO_3001558|OXA-372__Citrobacter_freundii__TM_#04
TTFWLDGSFTVSAVEQ
>gb|AIG22448_1|ARO_3001558|OXA-372__Citrobacter_freundii__TM_#05
KIYLRELPFRD
>gb|AIG22448_1|ARO_3001558|OXA-372__Citrobacter_freundii__TM_#06
VMLAEQTDSYKLYAKTGWA
>gb|AIG22448_1|ARO_3001558|OXA-372__Citrobacter_freundii__TM_#07
LRAERIIP
>gb|AAG03380_1|ARO_3002703|cmx__Corynebacterium_striatum__TM_#01
AVGVIRGVTNNVGRSETSATSPRLRVELSQ
>gb|AAG03380_1|ARO_3002703|cmx__Corynebacterium_striatum__TM_#02
RRALTKTAAEAN
>gb|AEM44648_1|ARO_3001970|CTX-M-110__Shigella_sp__SH165__TM_#01
SAPLYAQTSA
>gb|ENU21024_1|ARO_3001751|OXA-296__Acinetobacter_bohemicus_ANC_3994__TM_#01
QLFNSAHTS
>gb|CAB75346_1|ARO_3000582|L1_beta-lactamase__Stenotrophomonas_maltophilia__TM_#01
MRSTLLAFALSSLALAATLFTFDGA
>gb|CAB75346_1|ARO_3000582|L1_beta-lactamase__Stenotrophomonas_maltophilia__TM_#02
LVLIDGGMPQMASYLLTNMKARGTNTGPLRMVLLSHAHTDHAGPVAEIKRRTGAQ
>gb|CAD55718_1|ARO_3000197|tet36__Bacteroides_coprosuis_DSM_18011__TM_#01
RMLDGAILVVSAKEGIQAQTRLLFNVLQQLEIPTILFVNKIDREGVNLNQLYLEIQNSLS
KDIIFMQSVEGKELTSSCTIHYISEKNRETILEKDDLLLEKYLSDTQLSNLDYWNSMVRL
VQA
>gb|CAD55718_1|ARO_3000197|tet36__Bacteroides_coprosuis_DSM_18011__TM_#02
LYSLNGSDENLKIRGLKTFYSGDEIDVDEVFTNDIAIADHADNLMVGDYLGIMPNLFDKL
NIPSPALKSSIHPAKVENRSKLISAMNVLSVEDPSLAFSINADNNELEVSLYGATQREVI
LTLLEERFSVDAYFEEVKTIYKERLKT
>gb|CAD55718_1|ARO_3000197|tet36__Bacteroides_coprosuis_DSM_18011__TM_#03
LPIGAGLVMESEISLGYLNRSFQNAVFDGV
>gb|AAC23556_1|ARO_3002660|APH(6)-Id__Pseudomonas_aeruginosa__TM_#01
HGDYQATE
>gb|AAC23556_1|ARO_3002660|APH(6)-Id__Pseudomonas_aeruginosa__TM_#02
DYVHAAIIADQMMS
>gb|AJP77057|ARO_3003718|MSI-1__Massilia_sp__SB1-3__TM_#01
MGARSCSRAWCCSRATRSAPPGRRSGRLSCARCRRPSTGCNPSSTPPITGDIMKHIRLAC
VAAGLVAAASSVSAIG
>gb|AJP77057|ARO_3003718|MSI-1__Massilia_sp__SB1-3__TM_#02
NQGHILIDGGSDKSPPQIAARIRQLGFKPEDIRFILV
>gb|AJP77057|ARO_3003718|MSI-1__Massilia_sp__SB1-3__TM_#03
NAEVLAGAAAVPVLHSGEAGRNDPQYGGLPKMAPVARV
>gb|AJP77057|ARO_3003718|MSI-1__Massilia_sp__SB1-3__TM_#04
GGFRYSGDARYPSARADVER
>gb|AJP77057|ARO_3003718|MSI-1__Massilia_sp__SB1-3__TM_#05
TRYERRAAQGNAAFIDAGACKAYAVKARVKLQEQLARETAKP
>gb|AAK76136_1|ARO_3000025|patB__Streptococcus_pneumoniae_TIGR4__TM_#01
WQSLSGIMVNLGLLVLVLFISSVIYMCLMTRVIAE
>gb|AAK76136_1|ARO_3000025|patB__Streptococcus_pneumoniae_TIGR4__TM_#02
FNESLIQVMSNIVLYIGLILVMFSRNVTLALITIASTPLAFLMLIFIVKMARKYTNLQQK
EVGKLNAYMDESISGQKAVIVQGIQEDMMAGFLEQNERVRKATFKGRM
>gb|CAA23892_1|ARO_3002644|APH(3_)-IIa__Escherichia_coli__TM_#01
CSDAAVFRLSAQGRPVLFVKTDLSGALNELQDEAARLSWLATTGVPCAAVLDVVT
>gb|CAA23892_1|ARO_3002644|APH(3_)-IIa__Escherichia_coli__TM_#02
HLAPAEKVSIMADAMRRLHT
```

>gb|CAA23892_1|ARO_3002644|APH(3_)-IIa__Escherichia_coli__TM_#03
QAKHRIERART
>gb|AAN32638_1|ARO_3000842|EBR-1_beta-lactamase__Empedobacter_brevis__TM_#01
FSLIALIGSFAFGQIKPIQIDPINNNLFVYQTFNSFN
>gb|AAN32638_1|ARO_3000842|EBR-1_beta-lactamase__Empedobacter_brevis__TM_#02
LNIPTYATSLTNSK
>gb|AAN32638_1|ARO_3000842|EBR-1_beta-lactamase__Empedobacter_brevis__TM_#03
KFGNEKFV
>gb|AAN32638_1|ARO_3000842|EBR-1_beta-lactamase__Empedobacter_brevis__TM_#04
VEWPKTVHKLVAKH
>gb|ACT97415_1|ARO_3002999|CblA-1__mixed_culture_bacterium_AX_gF3SD01_15__TM_#01
MKAYFIAILTLFTCIATVVRAQQMSELENRIDSLLN
>gb|ACT97415_1|ARO_3002999|CblA-1__mixed_culture_bacterium_AX_gF3SD01_15__TM_#02
TDKGDMLRYNDH
>gb|ACT97415_1|ARO_3002999|CblA-1__mixed_culture_bacterium_AX_gF3SD01_15__TM_#03
PPNTYSPLRKK
>gb|ACT97415_1|ARO_3002999|CblA-1__mixed_culture_bacterium_AX_gF3SD01_15__TM_#04
YIHRLSIDSFNLSETEDGMHSSFEAVYRNWSTPSAMVRLLRTADEKELFSNKELKDFLWQ
TMIDTETGANKLKGMLPAKTVVGHKTGSSDRNADGMKTADNDAGLVILPDGRKYYIAAFV
MDSYETDEDNANIIARISRMVYDAMR
>gb|BAB56495_1|ARO_3000746|mepR__Staphylococcus_aureus_subsp__aureus_Mu50__TM_#01
MEFTYSYLFRMISHEMKQKADQKLEQFDITNEQGHTLGYLYAHQQDGLTQNDIAKALQRT
GPTVSNLLRNLERKKLIYRYVDAQDTRRKNIGLTTSGIKLVEAFTSIFDEMEQTLVSQLS
EEENEQMKANLTKMLSSLQ
>gb|ABI18382_1|ARO_3003855|ADC-10__Acinetobacter_baumannii__TM_#01
MRFKKISCLLL
>gb|ABI18382_1|ARO_3003855|ADC-10__Acinetobacter_baumannii__TM_#02
EIKKLVDQNFKPLL
>gb|ABI18382_1|ARO_3003855|ADC-10__Acinetobacter_baumannii__TM_#03
KYDVPGMAV
>gb|ABI18382_1|ARO_3003855|ADC-10__Acinetobacter_baumannii__TM_#04
DQQVLTFFK
>gb|ABI18382_1|ARO_3003855|ADC-10__Acinetobacter_baumannii__TM_#05
KVVALSMNKPF
>gb|ABI18382_1|ARO_3003855|ADC-10__Acinetobacter_baumannii__TM_#06
APASPAYGVKSTLPDML
>gb|ABI18382_1|ARO_3003855|ADC-10__Acinetobacter_baumannii__TM_#07
DIQRAINETHQG
>gb|ABI18382_1|ARO_3003855|ADC-10__Acinetobacter_baumannii__TM_#08
TMYQALGWEEFSY
>gb|ABI18382_1|ARO_3003855|ADC-10__Acinetobacter_baumannii__TM_#09
YVVFIPKENIGLVMLT
>gb|ABX00624_1|ARO_3002881|lmrC__Streptomyces_lincolnensis__TM_#01
GHVGYLPQSLPLIDGTVDEALE
>gb|ABX00624_1|ARO_3002881|lmrC__Streptomyces_lincolnensis__TM_#02
ELEQENVQRAVRGAEQELRRHKREAQA
>gb|ABX00624_1|ARO_3002881|lmrC__Streptomyces_lincolnensis__TM_#03
ASSGVPRIHAGALQRQAQESAGRAASVHQDRVSQAKAKLDEASQGMREEARLAITLPQTS
VPAGRTVLTCHEANVRYGERTLFT
>gb|ABX00624_1|ARO_3002881|lmrC__Streptomyces_lincolnensis__TM_#04
RFAPHLQDGEVRYRLAQFLFRGDRVHRTAGW
>gb|ABX00624_1|ARO_3002881|lmrC__Streptomyces_lincolnensis__TM_#05
IAEADDAWPHRDK
>gb|CAA26964_1|ARO_3000347|ErmA__Staphylococcus_aureus__TM_#01
GGLCQVTKEAVNPSENIKVIQTDILKFSFPKHIN
>gb|AAL19308_1|ARO_3000504|golS__Salmonella_enterica_subsp__enterica_serovar_Typhimurium_str__LT2__TM_#01
DRRIQNMQHMAQTL
>gb|AAL19308_1|ARO_3000504|golS__Salmonella_enterica_subsp__enterica_serovar_Typhimurium_str__LT2__TM_#02
LGQPDDSEP
>gb|AAG07106_1|ARO_3004056|ArmR__Pseudomonas_aeruginosa_PAO1__TM_#01
MSLNTPRNKPSRTETEAVAASSGRSAVGRRDYTEQLRRAARRNAWDLYGEHFY
>gb|ANK04027_1|ARO_3003838|gadW__Escherichia_coli_O25b_H4__TM_#01
LPRNLGLHSKDRLLINQSPPIQLVTAIFDSFNDPR
>gb|ANK04027_1|ARO_3003838|gadW__Escherichia_coli_O25b_H4__TM_#02
IPLLFNSISTVSGKVERLISFDIAKRWYLRDIAERMYTSESLIKKKLQ
>gb|AAG11411_2|ARO_3004680|APH(3_)-VIIIa__Streptomyces_rimosus__TM_#01
MDDALRALRGRYPGCEWVVVEDGASGAGVYRLRGGGRELFVKVAALGAGVGLLGEAERLV
WLAEVGIPVPRVVEGGGDERV
>gb|AAG11411_2|ARO_3004680|APH(3_)-VIIIa__Streptomyces_rimosus__TM_#02
LDVAVALAGLARSLH
>gb|AAG11411_2|ARO_3004680|APH(3_)-VIIIa__Streptomyces_rimosus__TM_#03
ARAVAEGSVDLEDLDEERK
>gb|AAG11411_2|ARO_3004680|APH(3_)-VIIIa__Streptomyces_rimosus__TM_#04
SAAFLREYGRGWDGA
>gb|ABA71728_1|ARO_3002937|vanSG__Enterococcus_faecalis__TM_#01
FGAFALISASLSLLSGHFSRAVVGILEIFYKDYEKALVVYTYVFRDNKEWFVMIAAFVS
>gb|ABA71728_1|ARO_3002937|vanSG__Enterococcus_faecalis__TM_#02
KEESSEDVVLSSELAATEKTINTIKHTLEQQKTA
>gb|ABA71728_1|ARO_3002937|vanSG__Enterococcus_faecalis__TM_#03
MLAEKKLNCVLKTM
>gb|ABA71728_1|ARO_3002937|vanSG__Enterococcus_faecalis__TM_#04
SITVTQNENMVH
>gb|ABA71728_1|ARO_3002937|vanSG__Enterococcus_faecalis__TM_#05
TARSEDEKIEFEVTILSS
>gb|ADQ43421_1|ARO_3002646|APH(3_)-IIc__Stenotrophomonas_maltophilia__TM_#01
MEASNPFTDGLR
>gb|ADQ43421_1|ARO_3002646|APH(3_)-IIc__Stenotrophomonas_maltophilia__TM_#02
RRVAELLADALRGLHAVPVAN
>gb|BAE78082_1|ARO_3003550|mdtP__Escherichia_coli_str__K-12_substr__W3110__TM_#01
LPVLAEEMSLLMLDSRRVIQSIQLMKSLGGGYQAGPVVEKK
>gb|CAD37801_1|ARO_3003793|SPM-1__Pseudomonas_aeruginosa__TM_#01
MNSPKSRALLGFMGAFCLLLVAGAPLSAKSSDHVDLPYNLTATKIDSDVFVVTDRDFYSS
NVLVAKMLDGTVVIVSSPFENLGTQTLMDWVAKTMKPKKVVAINTHFHLDGTGGNEIYKK
MGAETWSSDLTKQLRLEENKKDRIKAAEFYKNEDLKRRILSSHPVPADNVFDLKQGKVFS
FSNELVEVS
>gb|CAD37801_1|ARO_3003793|SPM-1__Pseudomonas_aeruginosa__TM_#02
KKKLLFGGCMIKPKELGYLGDANVKAWPDSARRLKKFDAKIVIPGHGEWGGPEMVNKTIK
VAEKAVGEMRL
>gb|AJD07405_1|ARO_3001600|OXA-409__Acinetobacter_baumannii__TM_#01
LLLITSAIFISACCSPYIV
>gb|AJD07405_1|ARO_3001600|OXA-409__Acinetobacter_baumannii__TM_#02
ANPNHSASKSD
>gb|AJD07405_1|ARO_3001600|OXA-409__Acinetobacter_baumannii__TM_#03

KAEKIKNLFNE
>gb|AJD07405_1|ARO_3001600|OXA-409__Acinetobacter_baumannii__TM_#04
HTTGVLVI
>gb|AJD07405_1|ARO_3001600|OXA-409__Acinetobacter_baumannii__TM_#05
QGQTQQSYGNDLA
>gb|AJD07405_1|ARO_3001600|OXA-409__Acinetobacter_baumannii__TM_#06
KRLFPEWEK
>gb|AJD07405_1|ARO_3001600|OXA-409__Acinetobacter_baumannii__TM_#07
MTLGDAMKASA
>gb|AJD07405_1|ARO_3001600|OXA-409__Acinetobacter_baumannii__TM_#08
KLANKTLPFS
>gb|AJD07405_1|ARO_3001600|OXA-409__Acinetobacter_baumannii__TM_#09
KVQDEVQSMLFIEEKN
>gb|AGH19769_1|ARO_3003198|rmtH__Klebsiella_pneumoniae__TM_#01
MTIEQAAADILSSKKYQLLCPDTVV
>gb|AGH19769_1|ARO_3003198|rmtH__Klebsiella_pneumoniae__TM_#02
KPKQAVERTRE
>gb|AGH19769_1|ARO_3003198|rmtH__Klebsiella_pneumoniae__TM_#03
LAPQVEKQASTALAAGDVQK
>gb|AGH19769_1|ARO_3003198|rmtH__Klebsiella_pneumoniae__TM_#04
DTYPQLYQFVFENNLPA
>gb|AGH19769_1|ARO_3003198|rmtH__Klebsiella_pneumoniae__TM_#05
MLHRQGVASVWGCDIHQGLGNVLTPYAQKHG
>gb|AGH19769_1|ARO_3003198|rmtH__Klebsiella_pneumoniae__TM_#06
VAASGDMALVFKLLPLLEREQPGAALALLRTLDAPVIC
>gb|AGH19769_1|ARO_3003198|rmtH__Klebsiella_pneumoniae__TM_#07
HQHYATWFEGLVAPHFTVQHHTLIGDELLYRIQPNPA
>gb|CAB14620_1|ARO_3002627|aadK__Bacillus_subtilis_subsp__subtilis_str__168__TM_#01
REAEDYFANNDGLVKVLLDKDSFINYK
>gb|CAB14620_1|ARO_3002627|aadK__Bacillus_subtilis_subsp__subtilis_str__168__TM_#02
KGYSFSMGKNYKFMKRYLSNKEWEELMSTYSV
>gb|CAB14620_1|ARO_3002627|aadK__Bacillus_subtilis_subsp__subtilis_str__168__TM_#03
GLAYKYPDYDEGITKYTEGIYCSVK
>gb|AAK63040_1|ARO_3002635|APH(2__)-IIa__Escherichia_coli__TM_#01
MVNLDAEIYEHLNKQIKINELRYLSSGDDSDTFLCNEQYVVKVPKRDSVRISQKRELELY
RFLENCKLSYQIPAVVYQSDRFNI
>gb|AAK63040_1|ARO_3002635|APH(2__)-IIa__Escherichia_coli__TM_#02
LLISILEKEQLLTDEMLEHIETIYENILSNAVLFKYTPCLVHNDFSANNMIFRNNRLFGV
IDFGDFN
>gb|AAK63040_1|ARO_3002635|APH(2__)-IIa__Escherichia_coli__TM_#03
QHKAPEVAERKAELNDVYWSIDQIIYGYERKDREMLIKDVSELLQTQAEMFIF
>gb|AAC72341_1|ARO_3000182|tet(Y)__IncQ_plasmid_pIE1120__TM_#01
MSKSLITALIVVA
>gb|AAC72341_1|ARO_3000182|tet(Y)__IncQ_plasmid_pIE1120__TM_#02
PAEQTAFHYGVFLSLYAFMQVFCAPV
>gb|AAC72341_1|ARO_3000182|tet(Y)__IncQ_plasmid_pIE1120__TM_#03
IILLVSFLGATIDYSIMAAAPVLWVLYIGRIISGVTGATGAIAASIIADTTKQEERARWF
GF
>gb|AAC72341_1|ARO_3000182|tet(Y)__IncQ_plasmid_pIE1120__TM_#04
DISVHAPFVAGALLNAIAFCLVAFLLPKTPSQPPEGQPAKINLFEGFRFNFAVQGLASFF
ALFFLMQLIGQAPAALWVIYGEQRLNWDIGTAGVSLAVFGAAHTFVQAVLTGTLSKRLGD
RGVLLLGMGADMCGFLLLAFITQSWMVLPAIFMLATGGIGMPALQAIISGLVCDEKQGAL
QGTLTGLTNITSIIGPVGF
>gb|CAD12765_1|ARO_3002203|IMP-12__Pseudomonas_putida__TM_#01
LPDLKIEKLE
>gb|CAD12765_1|ARO_3002203|IMP-12__Pseudomonas_putida__TM_#02
DAYLIDTPF
>gb|CAD12765_1|ARO_3002203|IMP-12__Pseudomonas_putida__TM_#03
SSHFHSDS
>gb|CAA09666_1|ARO_3004122|Klebsiella_pneumoniae_OmpK37__Klebsiella_pneumoniae__TM_#01
NAQDINVGTNNR
>gb|AAK38324_1|ARO_3000840|JOHN-1_beta-lactamase__Flavobacterium_johnsoniae_UW101__TM_#01
KPDYIISGHDDWTSK
>gb|ACJ59254_1|ARO_3000768|abeS__Acinetobacter_baumannii_AB307-0294__TM_#01
AIGWIFYKQHLDLAACIGLALMIAGIVIINVFSKNTHL
>gb|CBL58195_1|ARO_3002831|vgaC__Staphylococcus_aureus__TM_#01
MVLLEAKNIKHYIKDRLLLKIDELK
>gb|CBL58195_1|ARO_3002831|vgaC__Staphylococcus_aureus__TM_#02
EKFIPDEGTITPYAQSEILPQLKKTDAA
>gb|CBL58195_1|ARO_3002831|vgaC__Staphylococcus_aureus__TM_#03
LKQKEVEKQQQQST
>gb|CBL58195_1|ARO_3002831|vgaC__Staphylococcus_aureus__TM_#04
EKIKDQPPIKMDLPNERNLKNRVIIRVEDLEGLVPKQLLWKKATFQ
>gb|CBL58195_1|ARO_3002831|vgaC__Staphylococcus_aureus__TM_#05
MTLDVNKSILENVQNS
>gb|CBL58195_1|ARO_3002831|vgaC__Staphylococcus_aureus__TM_#06
LKALENLLNEYTGSIIFVSHDRTFTENIATRILEIRNKKIEIFDGTYQQFKNRSTKKERD
FQQEEQLLLDTKISEV
>gb|CBL58195_1|ARO_3002831|vgaC__Staphylococcus_aureus__TM_#07
NLLKEKSKLKE
>gb|BAB35162_1|ARO_3000676|H-NS__Escherichia_coli_O157_H7_str__Sakai__TM_#01
NSLAAVKSGTKAKRAQ
>gb|BAB35162_1|ARO_3000676|H-NS__Escherichia_coli_O157_H7_str__Sakai__TM_#02
KSLDDFLIKQ
>gb|AKQ05895_1|ARO_3004586|tet(51)__uncultured_bacterium__TM_#01
CANLRKGGHAVDIRGVAIDLAKSMGIYK
>gb|AKQ05895_1|ARO_3004586|tet(51)__uncultured_bacterium__TM_#02
FGFREGDDIEIL
>gb|AKQ05895_1|ARO_3004586|tet(51)__uncultured_bacterium__TM_#03
MGDVPCHFNQWVESIKQR
>gb|AKQ05895_1|ARO_3004586|tet(51)__uncultured_bacterium__TM_#04
TNLGAYFSAFSIPNYLNLNHTDVQFEANQKLISMASDKNPKIAITGFCFRAQNVLNNL
>gb|AVX51087_1|ARO_3004559|CAM-1__Pseudomonas_aeruginosa__TM_#01
MKSTAIILFLLVFSLGVFGQTGDALKISQLSGDFYIFTTYQTYKDAKVS
>gb|AVX51087_1|ARO_3004559|CAM-1__Pseudomonas_aeruginosa__TM_#02
ETQLQPLLNYIKEKHNKDVVMSVSTHFHEDRTNGIEFLRTKGVKTYTTKKTDELSQKKGY
ERAEFLLEKDTEFKIGQYKFQ
>gb|AVX51087_1|ARO_3004559|CAM-1__Pseudomonas_aeruginosa__TM_#03
EDIGNLSDANIDEWSN
>gb|AVX51087_1|ARO_3004559|CAM-1__Pseudomonas_aeruginosa__TM_#04
ASTKSLKHTLKLIKKTRKK
>gb|AFH35853_1|ARO_3001328|Escherichia_coli_mdfA__Escherichia_coli__TM_#01
RQPGMLEN

```
>gb|BAB20748_1|ARO_3002825|ErmY__Staphylococcus_aureus__TM_#01
SNLCIQTQNKVTNYDNFRIINKDILQFKFPNNKA
>gb|BAB20748_1|ARO_3002825|ErmY__Staphylococcus_aureus__TM_#02
EATVSYLIVEE
>gb|BAB20748_1|ARO_3002825|ErmY__Staphylococcus_aureus__TM_#03
SLIILKRHPSKISY
>gb|BAB20748_1|ARO_3002825|ErmY__Staphylococcus_aureus__TM_#04
YAKIKDLKNINF
>gb|ADK25050_1|ARO_3002267|IND-11__Chryseobacterium_indologenes__TM_#01
LDLLDKNKKPE
>gb|CAC32082_1|ARO_3002527|AAC(2_)-Ie__Mycobacterium_leprae__TM_#01
LRRLQQMVT
>gb|CAC32082_1|ARO_3002527|AAC(2_)-Ie__Mycobacterium_leprae__TM_#02
RHGTIIAHAAVVQRRLF
>gb|CAC32082_1|ARO_3002527|AAC(2_)-Ie__Mycobacterium_leprae__TM_#03
KDCRGRGLV
>gb|CAC32082_1|ARO_3002527|AAC(2_)-Ie__Mycobacterium_leprae__TM_#04
FGALSSSDRARRVYM
>gb|BAP34782_1|ARO_3002008|CTX-M-151__Salmonella_enterica__TM_#01
MINKRLSIALALAAMIGTPVAMALESQKPGSDSANHIQHQMVQQLSALEKSANGRLGVAV
IDTGSGAIAGWRMDEP
>gb|BAP34782_1|ARO_3002008|CTX-M-151__Salmonella_enterica__TM_#02
TPELMSQPQPVASG
>gb|BAP34782_1|ARO_3002008|CTX-M-151__Salmonella_enterica__TM_#03
RFVGKSMTFD
>gb|BAP34782_1|ARO_3002008|CTX-M-151__Salmonella_enterica__TM_#04
LALGDALGQVQREKLSH
>gb|BAP34782_1|ARO_3002008|CTX-M-151__Salmonella_enterica__TM_#05
QAESQRPVLAKAAAIVASHYVLPKG
>gb|AAK64581_1|ARO_3004568|dfrA18__Vibrio_cholerae_MO10__TM_#01
MNKEIQFSMIVARGVNGEIGQDGDLPWHVEGVRLKEDLKR
>gb|AAK64581_1|ARO_3004568|dfrA18__Vibrio_cholerae_MO10__TM_#02
DKADITERGDGTFVAWGNSCHLFEVAEHLGVTEIIVAGGAEIYNLHKDVITKVFETKVLR
AYPAADTHVDVFWESPGYDTEGRQWRVTSRGHIIENGSFTIATTYER
>gb|WP_149100971_1|ARO_3005351|DfrA39__Pseudomonas_aeruginosa__TM_#01
MNTPMIYIS
>gb|WP_149100971_1|ARO_3005351|DfrA39__Pseudomonas_aeruginosa__TM_#02
LVISRNQHYKCQDCTVVKNLDEAIAIAKEFGNELFVAGGAEIYSLAMPVAHRIYLTEISK
NFEGDVFFPEFNSADFRKISSEEVPAS
>gb|AAB41701_1|ARO_3002526|AAC(2_)-Id__Mycolicibacterium_smegmatis_MC2_155__TM_#01
MLTQHVSEARTRGA
>gb|AAB41701_1|ARO_3002526|AAC(2_)-Id__Mycolicibacterium_smegmatis_MC2_155__TM_#02
RDPSGDSDFTD
>gb|AAB41701_1|ARO_3002526|AAC(2_)-Id__Mycolicibacterium_smegmatis_MC2_155__TM_#03
DIARPMYIARGWLSWEGPTSVLTPTEGIV
>gb|AAB41701_1|ARO_3002526|AAC(2_)-Id__Mycolicibacterium_smegmatis_MC2_155__TM_#04
AREITCDWRSGDPW
>gb|AAK84316_1|ARO_3002843|vatD__Enterococcus_faecium__TM_#01
MKMYPIEGNKSVQFIKPILEKLENVE
>gb|AAK84316_1|ARO_3002843|vatD__Enterococcus_faecium__TM_#02
NGETFDKQILYHYPILNDKLK
>gb|AAK84316_1|ARO_3002843|vatD__Enterococcus_faecium__TM_#03
MLAGGNPANEIKQRFDQDT
>gb|AAK84316_1|ARO_3002843|vatD__Enterococcus_faecium__TM_#04
IREVIWKK
>gb|CBZ41939|ARO_3005097|mecC-type_BlaZ__Staphylococcus_aureus__Partial_TM_#01
MKKLIILVVLALILSACNSKNSTNND
>gb|CBZ41939|ARO_3005097|mecC-type_BlaZ__Staphylococcus_aureus__Partial_TM_#02
GQVKTKLIDLGDTTTHPSRKEPDLNFYSPKDKRDTSTPLAYGKTLKKLIADGDLSKANKD
FLLNLMF
>gb|AAD03493_1|ARO_3002565|AAC(6_)-Iu__Acinetobacter_dispersus__TM_#01
ACTEFASDAA
>gb|AAD03493_1|ARO_3002565|AAC(6_)-Iu__Acinetobacter_dispersus__TM_#02
ALGFHETERVVYFKKNI
>gb|AAB05622_1|ARO_3002921|vanRB__Enterococcus_faecalis__TM_#01
MSIRILLVEDDDHICNTVRAFLAEARYEVDACTDGNEAH
>gb|AAB05622_1|ARO_3002921|vanRB__Enterococcus_faecalis__TM_#02
VGRLTLLPEDFRVLCDGT
>gb|CEJ95855_1|ARO_3004595|erm45__Staphylococcus_fleurettii__TM_#01
MNQNIKFTQNFITNEKLLSNIMKQINIDENDIIYEVGTGKGHLTSKLAEKCKHVYSIELD
KKLYELSSNKLQDNSRVTLINQDILQFNYPYRKKYKIVGNIPFNISTQIVKDAVFRSQAS
EMYF
>gb|CEJ95855_1|ARO_3004595|erm45__Staphylococcus_fleurettii__TM_#02
MIDTRRTLSLQLQTQVYIQQLLPIPAGSFHPKPKVNCILIKLTRHISDIKDKHKKKYEFF
ISKWVNK
>gb|CEJ95855_1|ARO_3004595|erm45__Staphylococcus_fleurettii__TM_#03
HQALKHARIKDLNKISYEQVLSVFESYILFNPRK
>gb|AGV10818_1|ARO_3002669|APH(2__)-Ig__Campylobacter_coli_CVM_N29710__TM_#01
MCEFSSPQIPITDIENAMERIGSPVRELRRLDAGDDSEVLLCNGLFVIKIPKRPSVRVTQ
QREFAVYSFLKQYDLPALIPEVIFQCSEFNVMSFIPGENFGFQEYALLSEKEKEALASDM
AIFLRRLHGISVPLSEKPFCEIFEDKRKRYLEDQEQLLEVLENRKLLNAPLQKNIQTIYE
HIGQNQELFNYAACLVHNDFSSSNMVFRHNRLYGVIDFGDVI
>gb|AGV10818_1|ARO_3002669|APH(2__)-Ig__Campylobacter_coli_CVM_N29710__TM_#02
RHYGHRNPQLAERKAEINDAYWPIQQVLLGVQREDRSLFCKGYRELLAIDPDAFIL
>gb|ARO85965_1|ARO_3003115|OXA-427__Enterobacter_cloacae__TM_#01
MSRILLSSLLAAGLF
>gb|ARO85965_1|ARO_3003115|OXA-427__Enterobacter_cloacae__TM_#02
LPALPFKAGDPDFLPEWKQTTTPSRWM
>gb|AAN86837_2|ARO_3000601|Erm(38)__Mycolicibacterium_smegmatis__TM_#01
ARSAPGPASRPTEVVA
>gb|AAN86837_2|ARO_3000601|Erm(38)__Mycolicibacterium_smegmatis__TM_#02
QRLPTPVPRTWLRANGIAPNSLPRQLSAAQWAALFEQTRLTGAQRVDRPRDVQHGRAHRR
RGGEVDRPATHHKQTGPVVGQRQPQRGRDADADPDDQRTAPPVTR
>gb|BAA34540_1|ARO_3000319|mphC__Staphylococcus_aureus__TM_#01
ENDELWPRH
>gb|AAG16656_1|ARO_3002705|floR__Escherichia_coli__TM_#01
GSLNSSPS
>gb|AAG16656_1|ARO_3002705|floR__Escherichia_coli__TM_#02
AATEKSPVV
>gb|ACS44783_1|ARO_3002188|MOX-5__Aeromonas_caviae__TM_#01
LEKMQAYYRQWTPAYS
>gb|ACS44783_1|ARO_3002188|MOX-5__Aeromonas_caviae__TM_#02
LAASSMKQPF
```

```
>gb|ACS44783_1|ARO_3002188|MOX-5__Aeromonas_caviae__TM_#03
YGIKTSSADLLRFVKANI
>gb|ACS44783_1|ARO_3002188|MOX-5__Aeromonas_caviae__TM_#04
MTQGLGWE
>gb|ACS44783_1|ARO_3002188|MOX-5__Aeromonas_caviae__TM_#05
YPVSEQTLLAGNS
>gb|ACS44783_1|ARO_3002188|MOX-5__Aeromonas_caviae__TM_#06
HAILTQLAR
>gb|ABQ47844_1|ARO_3000215|mecR1__Staphylococcus_aureus_subsp__aureus_JH9__TM_#01
QNIMSHKIWLLVLVSTLIPLIPFYKISNFTFSKDMMNRNVSDTTSSVSHMLDGQQSSVTK
DLAINVNQFETSNITYMILLIWVFGSLLCLFYMIKAFRQIDVIKSSSLESSYLNERLKVC
QSKMQFYKKHITISYSSNIDNPMVFGLVKSQIVLPTVVVETMN
>gb|ABQ47844_1|ARO_3000215|mecR1__Staphylococcus_aureus_subsp__aureus_JH9__TM_#02
SLLIQAPLLSAHVQQDKYETNVSYKKLNQLAPYFKGFDGSFVLYNEREQAYSIYNEPESK
QRYSPNSTYKIYLALMAFDQNLLSLNHTEQQWDKHQYPFKEWNQDQNLNSSMKYSV
>gb|ABQ47844_1|ARO_3000215|mecR1__Staphylococcus_aureus_subsp__aureus_JH9__TM_#03
QHNMHFDNKAIEKVENSMTLK
>gb|AAA26683_1|ARO_3002840|vatA__Staphylococcus_aureus__TM_#01
MNLNNDHGPDPENIL
>gb|AAA26683_1|ARO_3002840|vatA__Staphylococcus_aureus__TM_#02
RMGWEKYMPSLKDLPL
>gb|AAA26683_1|ARO_3002840|vatA__Staphylococcus_aureus__TM_#03
LKFIRKRFSDG
>gb|AAA26683_1|ARO_3002840|vatA__Staphylococcus_aureus__TM_#04
KIINENLPFIINGDIEMLKRKRKLLDDT
>gb|AAB66655_1|ARO_3002663|APH(9)-Ib__Streptomyces_netropsis__TM_#01
MEDLPENLDQESLFQGLREFGISTTSASYAPLGFGDYHWHITGD
>gb|AAB66655_1|ARO_3002663|APH(9)-Ib__Streptomyces_netropsis__TM_#02
PAALRGLRRAMDTAVHLREQGGLPFVVAPRTTSDGASLVPLDSRYALTVFPHVSARPGEF
GQKLTERERDQVLVLLAELHGQAPPKCTPTTDMVPTGLDGVHTALAEPSGTWTGGPFSEP
ARELLAEHEATLRGRMAEFGE
>gb|AAB66655_1|ARO_3002663|APH(9)-Ib__Streptomyces_netropsis__TM_#03
DDPAALARYTE
>gb|AAB66655_1|ARO_3002663|APH(9)-Ib__Streptomyces_netropsis__TM_#04
SDTEAAWQSFAETLDHLNSEVPS
>gb|AAP27834_1|ARO_3005100|FosB2__Bacillus_anthracis_str__Ames__TM_#01
NEIKQSYTHMAFTVTNEALDHLKEVLIQNDVNILPGR
>gb|AAP27834_1|ARO_3005100|FosB2__Bacillus_anthracis_str__Ames__TM_#02
EDKKHMTFYI
>gb|CAA37477_1|ARO_3002891|otr(A)__Streptomyces_rimosus__TM_#01
DEEHVLTE
>gb|CAA37477_1|ARO_3002891|otr(A)__Streptomyces_rimosus__TM_#02
HGIRELLPSVHA
>gb|CAA37477_1|ARO_3002891|otr(A)__Streptomyces_rimosus__TM_#03
SETRATAGDIAQAW
>gb|CAA37477_1|ARO_3002891|otr(A)__Streptomyces_rimosus__TM_#04
NFFAPPSLETVIRPERPEEAGRLHAALRMLD
>gb|CAA37477_1|ARO_3002891|otr(A)__Streptomyces_rimosus__TM_#05
AGAVVRLYGEVQKEILGSTLAESFG
>gb|CAA37477_1|ARO_3002891|otr(A)__Streptomyces_rimosus__TM_#06
GAPWVCASDRPSPARA
>gb|CAA37477_1|ARO_3002891|otr(A)__Streptomyces_rimosus__TM_#07
VRSPVSAADDFRKANARLVLMDALGR
>gb|AAB41956_1|ARO_3000800|MexC__Pseudomonas_aeruginosa__TM_#01
MADLRAIGRIGALAMAIALAGCGPAEERQEAAEMVLPVEVLTVQ
>gb|AAB41956_1|ARO_3000800|MexC__Pseudomonas_aeruginosa__TM_#02
KGTLAAGDSQ
>gb|AAB41956_1|ARO_3000800|MexC__Pseudomonas_aeruginosa__TM_#03
HRSSDGSAQVMVVGADERAESRS
>gb|AAB41956_1|ARO_3000800|MexC__Pseudomonas_aeruginosa__TM_#04
QAQAQSPAPQQ
>gb|AAQ63644_1|ARO_3004452|Bacillus_clausii_chloramphenicol_acetyltransferase__Bacillus_clausii__TM_#01
MEILTLLIRDTMGSKRIFLNPYFQQQP
>gb|AAQ63644_1|ARO_3004452|Bacillus_clausii_chloramphenicol_acetyltransferase__Bacillus_clausii__TM_#02
LYTIFDNKSHTFSGIWSPNLTIFSEFHSKYENDAERYNGTRRLFPKKPIPDNP
>gb|AAQ63644_1|ARO_3004452|Bacillus_clausii_chloramphenicol_acetyltransferase__Bacillus_clausii__TM_#03
QVNDELFLPVSIQNASCLFVMATMQSVFINDLQNLVDESEDWIYLVVSDEWYY
>gb|NP_388442_1|ARO_3004476|vmlR__Bacillus_subtilis_subsp__subtilis_str__168__TM_#01
APAQGQILRKDIKLALVEQETAA
>gb|NP_388442_1|ARO_3004476|vmlR__Bacillus_subtilis_subsp__subtilis_str__168__TM_#02
LQFLIQQLK
>gb|NP_388442_1|ARO_3004476|vmlR__Bacillus_subtilis_subsp__subtilis_str__168__TM_#03
TPEYTVRFSIDTTH
>gb|NP_388442_1|ARO_3004476|vmlR__Bacillus_subtilis_subsp__subtilis_str__168__TM_#04
KELDQAFNELTKRIKELDHQDKKD
>gb|ACH58990_1|ARO_3002511|LRA-12__uncultured_bacterium_BLR12__TM_#01
MNVQNCMVKAVSVSIILFASLSLAAQKVKEPTVSN
>gb|ACH58990_1|ARO_3002511|LRA-12__uncultured_bacterium_BLR12__TM_#02
KQGNIIVNTGLAASALQIK
>gb|ACH58990_1|ARO_3002511|LRA-12__uncultured_bacterium_BLR12__TM_#03
LMADEGDAT
>gb|ACH58990_1|ARO_3002511|LRA-12__uncultured_bacterium_BLR12__TM_#04
AFGGHGSMFEPIIADRLLHDKDTIQ
>gb|ACH58990_1|ARO_3002511|LRA-12__uncultured_bacterium_BLR12__TM_#05
IEKKFSEVSSYPGIAKDYAYTLQ
>gb|ACH58990_1|ARO_3002511|LRA-12__uncultured_bacterium_BLR12__TM_#06
SMHSKHKPGDGYNPKSFMDRKGYDESLDKLQKEYEKHLNEN
>gb|ABK33456_1|ARO_3002639|APH(3__)-Ib__Pseudomonas_aeruginosa__TM_#01
RRGELAGERDRLIWLKGRGVACPEVI
>gb|AAK76137_1|ARO_3000024|patA__Streptococcus_pneumoniae_TIGR4__TM_#01
EILDAEPAMTFKDIPDEELVGSLSFENVTFTYPMDKEPMLKDVSFT
>gb|AAK76137_1|ARO_3000024|patA__Streptococcus_pneumoniae_TIGR4__TM_#02
RAANIAQASEFIHRMEKT
>gb|AAA98406_1|ARO_3001404|OXA-9__Klebsiella_pneumoniae__TM_#01
MKKILLLHMLVFVSATLPISSVASDEVETLKCTIIADAITGNTLYETGECARRV
>gb|AAA98406_1|ARO_3001404|OXA-9__Klebsiella_pneumoniae__TM_#02
QSPKSPTWELKPEYNPSPRDRTYKQ
>gb|AAA98406_1|ARO_3001404|OXA-9__Klebsiella_pneumoniae__TM_#03
VDRFTEYVKKFE
>gb|AAA98406_1|ARO_3001404|OXA-9__Klebsiella_pneumoniae__TM_#04
LMSSLTISPKEQIQFLLRFVAHKLPVSEAAYDMAYATIPQYQAAEGWAVHGKSGSGWLRD
NNGKI
>gb|CAC14596_1|ARO_3003057|smeF__Stenotrophomonas_maltophilia__TM_#01
```

TLALAGCSTLAPKNTAVAPAIPAQWPAEAAQGEVADVAAV
>gb|CAC14596_1|ARO_3003057|smeF__Stenotrophomonas_maltophilia__TM_#02
VAVTGQMDRRGTDAGVTEQ
>gb|CAC14596_1|ARO_3003057|smeF__Stenotrophomonas_maltophilia__TM_#03
AQRLKIADATLKTYEDSLRLAEARH
>gb|CAC14596_1|ARO_3003057|smeF__Stenotrophomonas_maltophilia__TM_#04
AGGQLDPALLPDSIEPQLL
>gb|AAC64365_1|ARO_3000230|ANT(2__)-Ia__Pseudomonas_aeruginosa__TM_#01
MDTTQVTLIHKILAAADER
>gb|AAC64365_1|ARO_3000230|ANT(2__)-Ia__Pseudomonas_aeruginosa__TM_#02
KHDDIDLTFPGERRGELEAIVEMLGGRVMEELDYGFLAEIGDELLDCEPAWWADEAYEIA
EAPQGSCPEAAEGVIAGRPV
>gb|AAC64365_1|ARO_3000230|ANT(2__)-Ia__Pseudomonas_aeruginosa__TM_#03
VPPVDWPTKHIESYRLACTSLGAEKVEVLRAAFRSRYAA
>gb|AAF63432|ARO_3003744|vatF__Yersinia_enterocolitica__TM_#01
MEDKPILGPDPQC
>gb|AAF63432|ARO_3003744|vatF__Yersinia_enterocolitica__TM_#02
SRDELPYKGDTH
>gb|AAF63432|ARO_3003744|vatF__Yersinia_enterocolitica__TM_#03
TLIKNRFSAEVIGKLQTIAWWDWPIDAISRNLHLIVAGDIEALARAASEIDHT
>gb|ACT97371_1|ARO_3003097|CfxA6__uncultured_organism__TM_#01
MSNYSVAELRNM
>gb|ACT97371_1|ARO_3003097|CfxA6__uncultured_organism__TM_#02
KNRKKQIVVL
>gb|ACT97371_1|ARO_3003097|CfxA6__uncultured_organism__TM_#03
IALVCIFILVFSL
>gb|ACT97371_1|ARO_3003097|CfxA6__uncultured_organism__TM_#04
VLTDSISQIVSACPGEIGVAVI
>gb|ACT97371_1|ARO_3003097|CfxA6__uncultured_organism__TM_#05
KLDPKTWSPM
>gb|ACT97371_1|ARO_3003097|CfxA6__uncultured_organism__TM_#06
MLMNRLFTE
>gb|AAC75731_1|ARO_3000516|emrR__Escherichia_coli_str__K-12_substr__MG1655__TM_#01
HSIQPSELSC
>gb|AAC75731_1|ARO_3000516|emrR__Escherichia_coli_str__K-12_substr__MG1655__TM_#02
ERRESDNDRRCLHLQLTEKGHEFLR
>gb|AAC75731_1|ARO_3000516|emrR__Escherichia_coli_str__K-12_substr__MG1655__TM_#03
STTEKDQLEQITRKLLSRLDQMEQDGVVLEAMS
>gb|AAC09015_1|ARO_3002481|AER-1__Aeromonas_hydrophila__TM_#01
MYVLSVEKPTLRNKFAAGIGVVLVCVVASFIPTPVFALDTTKLIQAVQSEESALHARVGM
TVFDSNTGT
>gb|AAC09015_1|ARO_3002481|AER-1__Aeromonas_hydrophila__TM_#02
KVDGKSLSLG
>gb|AAC09015_1|ARO_3002481|AER-1__Aeromonas_hydrophila__TM_#03
FDAIGGATGFNAYMRSIGDEE
>gb|AAC09015_1|ARO_3002481|AER-1__Aeromonas_hydrophila__TM_#04
KILLGDALSASSRSQLTQWMLD
>gb|AAC09015_1|ARO_3002481|AER-1__Aeromonas_hydrophila__TM_#05
VVLKDTVAP
>gb|ABS43901_1|ARO_3000783|cmeA__Campylobacter_jejuni_subsp__doylei_269_97__TM_#01
MKLFQKNTILVLGVVF
>gb|ABS43901_1|ARO_3000783|cmeA__Campylobacter_jejuni_subsp__doylei_269_97__TM_#02
VRNTQSGKWDLDSIHANLNLNGETVQ
>gb|AAC74149_2|ARO_3001216|mdtH__Escherichia_coli_str__K-12_substr__MG1655__TM_#01
LSMMPVGMV
>gb|CAA27626_1|ARO_3000363|EreB__Escherichia_coli__TM_#01
MRFEEWVKDKHIPFKLNHPDDNYDDFKPLRK
>gb|CAA27626_1|ARO_3000363|EreB__Escherichia_coli__TM_#02
TLLRFFIEDLGFTTFAFEFGFAEGQIINNWIHGQGTDDEIGRFLKHFYYPEELKTTFLWL
REYNKAAKEKITFLGIDIPRNGGSYLPNMEIVHDFFRTADKEALHIIDDAFNIAKKIDYF
STSQAALNLHELTDSEKCRLTSQLARVKVRLEAMAPIHIEKYGIDKYETILHYAN
>gb|CAA27626_1|ARO_3000363|EreB__Escherichia_coli__TM_#03
ILYDGFLSCLPMGQRLKNAIGDDYMSLGITSYSGHTAALYPEVDTKYGFRVDNFQLQEPN
EGSVEKAISGCGVTNSFVFFRNIPEDLQSIPNMI
>gb|ENU93232_1|ARO_3001749|OXA-294__Acinetobacter_sp__NIPH_758__TM_#01
MSKKLKLLALCATVISAATLVGCQNIQSQAQPLVLKKQ
>gb|ENU93232_1|ARO_3001749|OXA-294__Acinetobacter_sp__NIPH_758__TM_#02
QDQIATAFENIQTTGVLVTYDGKNFQ
>gb|ENU93232_1|ARO_3001749|OXA-294__Acinetobacter_sp__NIPH_758__TM_#03
PSTQQQVRDMLLIENVQ
>gb|QKT21444_1|ARO_3005085|AAC(3)-IIg__Enterobacter_cloacae__TM_#01
MNTRETIAADLSRLGVQSGA
>gb|QKT21444_1|ARO_3005085|AAC(3)-IIg__Enterobacter_cloacae__TM_#02
RRHREGLVGQAHC
>gb|CAC81324_1|ARO_3003015|dfrA19__Salmonella_enterica_subsp__enterica_serovar_Typhimurium__TM_#01
MSHPQLELIVAVDSKLGFGKGGKIPWKCKEDMARFTRISKEIRVCVIGKHTYTDMRDMQL
EKDGAEERIKEKGILPERESFVISSTLKQ
>gb|CAC81324_1|ARO_3003015|dfrA19__Salmonella_enterica_subsp__enterica_serovar_Typhimurium__TM_#02
LYENTDQRIAVIGGEKLYIQALSSATKLHMTIIPREFDCDRFIPVDPIQNNFHIDSSASE
TVEATVDETQERIHFATYVRNNQ
>gb|WP_104671188_1|ARO_3003948|efrA__Enterococcus_faecalis__TM_#01
IPKDMDYVYQQGIW
>gb|WP_104671188_1|ARO_3003948|efrA__Enterococcus_faecalis__TM_#02
REEEGVTE
>gb|ACX65640_1|ARO_3003907|cipA__Paenibacillus_sp__Y412MC10__TM_#01
MKYLSKYEKIRKILSALNQPNYRYSQITEAIFKNKIGNFEAMNNLPKPVRNELIKELGNN
VLSITPKMEQ
>gb|ACX65640_1|ARO_3003907|cipA__Paenibacillus_sp__Y412MC10__TM_#02
YIESVRLSYQT
>gb|ACX65640_1|ARO_3003907|cipA__Paenibacillus_sp__Y412MC10__TM_#03
TVLDEHIQQ
>gb|ACX65640_1|ARO_3003907|cipA__Paenibacillus_sp__Y412MC10__TM_#04
KAVADLLRE
>gb|AAC32027_1|ARO_3002817|carA__Streptomyces_thermotolerans__TM_#01
EGMRRTEAELAERPYQTGQDPE
>gb|AAC32027_1|ARO_3002817|carA__Streptomyces_thermotolerans__TM_#02
QSRETVAELTGVRVGDRLSVDSLH
>gb|AAC32027_1|ARO_3002817|carA__Streptomyces_thermotolerans__TM_#03
RPGMTVLQAFSS
>gb|AAC32027_1|ARO_3002817|carA__Streptomyces_thermotolerans__TM_#04
SRFNGAHLTLQDGRVAEFTAA
>gb|AAM70498_1|ARO_3001887|CTX-M-25__Escherichia_coli__TM_#01
EKHVNGTMT

```
>gb|AAM70498_1|ARO_3001887|CTX-M-25__Escherichia_coli__TM_#02
RRDVLAAAA
>gb|BAE77781_1|ARO_3000795|mdtE__Escherichia_coli_str__K-12_substr__W3110__TM_#01
NSAKGSLAKALSTA
>gb|BAE77781_1|ARO_3000795|mdtE__Escherichia_coli_str__K-12_substr__W3110__TM_#02
VASGQIKQVQGSTPVQLNLENGKRYS
>gb|BAE77781_1|ARO_3000795|mdtE__Escherichia_coli_str__K-12_substr__W3110__TM_#03
TALVDEGSRQNVLL
>gb|AAN28945|ARO_3003785|Chlamydia_trachomatis_intrinsic_murA_conferring_resistance_to_fosfomycin__Chlamydia_trachomatis__TM_#01
MPGIKVFGE
>gb|AAN28945|ARO_3003785|Chlamydia_trachomatis_intrinsic_murA_conferring_resistance_to_fosfomycin__Chlamydia_trachomatis__TM_#02
RTILKNVPNIEDVRQTVDLCRVLGAIVEWDQQAQVIEIH
>gb|AAN28945|ARO_3003785|Chlamydia_trachomatis_intrinsic_murA_conferring_resistance_to_fosfomycin__Chlamydia_trachomatis__TM_#03
WKKLGAEIVISDEGYWASAPNGLVGAH
>gb|AAN28945|ARO_3003785|Chlamydia_trachomatis_intrinsic_murA_conferring_resistance_to_fosfomycin__Chlamydia_trachomatis__TM_#04
QGRIFVEQARHEH
>gb|AAN28945|ARO_3003785|Chlamydia_trachomatis_intrinsic_murA_conferring_resistance_to_fosfomycin__Chlamydia_trachomatis__TM_#05
HENGIEFFYDKP
>gb|AAN28945|ARO_3003785|Chlamydia_trachomatis_intrinsic_murA_conferring_resistance_to_fosfomycin__Chlamydia_trachomatis__TM_#06
VKMGAHCDLFH
>gb|AAN28945|ARO_3003785|Chlamydia_trachomatis_intrinsic_murA_conferring_resistance_to_fosfomycin__Chlamydia_trachomatis__TM_#07
WIENTEMLDRGYTDWRGKLERLGAKVLARDAVSVYV
>gb|CAA78046_1|ARO_3002701|Rhodococcus_fascians_cmr__Rhodococcus_fascians__TM_#01
RRRALLTFLITFM
>gb|CAA78046_1|ARO_3002701|Rhodococcus_fascians_cmr__Rhodococcus_fascians__TM_#02
VTLVMLFVQG
>gb|CAA78046_1|ARO_3002701|Rhodococcus_fascians_cmr__Rhodococcus_fascians__TM_#03
LSFAVGSTLIS
>gb|CAA78046_1|ARO_3002701|Rhodococcus_fascians_cmr__Rhodococcus_fascians__TM_#04
YRAPLWTSAALV
>gb|CAA78046_1|ARO_3002701|Rhodococcus_fascians_cmr__Rhodococcus_fascians__TM_#05
LAIVIGAAT
>gb|AAA22289_1|ARO_3002672|Bacillus_pumilus_cat86__Bacillus_pumilus__TM_#01
MFKQIDENYL
>gb|AAA22289_1|ARO_3002672|Bacillus_pumilus_cat86__Bacillus_pumilus__TM_#02
KETKTFSSIWTPFDENFAQFYKSCVADIETFSK
>gb|AAA22289_1|ARO_3002672|Bacillus_pumilus_cat86__Bacillus_pumilus__TM_#03
EHCDEWLNDSLHIT
>gb|AAC43969_1|ARO_3003037|opcM__Burkholderia_cepacia__TM_#01
RSSISPGRSAISVDARQPVPVVEPY
>gb|AAC43969_1|ARO_3003037|opcM__Burkholderia_cepacia__TM_#02
VGDAASGKADVAAR
>gb|ABB43029_1|ARO_3002539|AAC(3)-IV__Escherichia_coli__TM_#01
MQYEWRKAELIGQLLNLGVTPGGVLLVHSSFRSVRPLEDGPL
>gb|ABB43029_1|ARO_3002539|AAC(3)-IV__Escherichia_coli__TM_#02
SPVTPDLGVVSDTFWRLPNVKRSA
>gb|ABB43029_1|ARO_3002539|AAC(3)-IV__Escherichia_coli__TM_#03
EQIISDPLPLPPHS
>gb|ABB43029_1|ARO_3002539|AAC(3)-IV__Escherichia_coli__TM_#04
RWLKEKSLQKEGPVGHAF
>gb|CCE73593_2|ARO_3001503|OXA-258__Achromobacter_ruhlandii__TM_#01
MTVRLVSRALGAVLFASALTLPARAD
>gb|CCE73593_2|ARO_3001503|OXA-258__Achromobacter_ruhlandii__TM_#02
QPVWNYQPA
>gb|CCE73593_2|ARO_3001503|OXA-258__Achromobacter_ruhlandii__TM_#03
DRQLVFARLLQDEGATQPNAGLRARDGLMRDWAAMVAAPRK
>gb|AAA16194_1|ARO_3002540|AAC(3)-VIa__Enterobacter_cloacae__TM_#01
MTDPRKNGDLHEPATAPATPWSKSELVRQLRDLGVRSGDMVMPHVSLRAVGPLADGPQTL
VDALIEAVGPTGNILAFVSWRDSPYEQTLGHDAPPAAIAQ
>gb|AAA16194_1|ARO_3002540|AAC(3)-VIa__Enterobacter_cloacae__TM_#02
AWLVAPHEMGAAYGPRSPIARFLAHA
>gb|AAA16194_1|ARO_3002540|AAC(3)-VIa__Enterobacter_cloacae__TM_#03
EGKRRVTYSMPLLR
>gb|AAA16194_1|ARO_3002540|AAC(3)-VIa__Enterobacter_cloacae__TM_#04
APDGPDAVERIARDYLARTRVAQGPVGGAQSRLIDAADIVSFGIEWLEARHAAPAAAALK
PKQRRD
>gb|AJP77076|ARO_3003715|PEDO-3__Pedobacter_sp__Stok-3__TM_#01
MRYLLSFLLCLSAFA
>gb|AJP77076|ARO_3003715|PEDO-3__Pedobacter_sp__Stok-3__TM_#02
VYTTYNTYKGALTD
>gb|AJP77076|ARO_3003715|PEDO-3__Pedobacter_sp__Stok-3__TM_#03
LTDEILKTNKEPRAAY
>gb|AJP77076|ARO_3003715|PEDO-3__Pedobacter_sp__Stok-3__TM_#04
IEANDLGYIGESDLAAWPKSIEKLKQKYPDTKIVITGHA
>gb|AAG33665_1|ARO_3001431|OXA-37__Acinetobacter_baumannii__TM_#01
MIIRFLALLFSAVVLVSLGHAQ
>gb|AAG33665_1|ARO_3001431|OXA-37__Acinetobacter_baumannii__TM_#02
KTHESSNWGKYFSDFNAKGTIVVVDERTNGNSTSVYNE
>gb|AAG33665_1|ARO_3001431|OXA-37__Acinetobacter_baumannii__TM_#03
RSYLEKLNYGNADPSTKSGDYWIDGNLAISAN
>gb|AAG33665_1|ARO_3001431|OXA-37__Acinetobacter_baumannii__TM_#04
AILQSVNALPPN
>gb|CAA79966_1|ARO_3003665|NmcR__Enterobacter_cloacae__TM_#01
QMLGVALFTRVPRGLQLTDEGMHLLPSITEALQMMSSAMDKFHEGKI
>gb|CAA79966_1|ARO_3003665|NmcR__Enterobacter_cloacae__TM_#02
ITAFLNENPWIDIRILTHNNVVNLAAEGIDASIRFGTGGWINTENILLFQAPHTVLCSPE
TSKKLY
>gb|CAA79966_1|ARO_3003665|NmcR__Enterobacter_cloacae__TM_#03
LMIDAVKLGDYAALVPYHMFQKELNERSVAKPFEIYATLGGYWLTLQKSRVNHNSEALNV
FKEWIIEHSREFVLKS
>gb|Q8FW76|ARO_3003772|Brucella_suis_mprF__Brucella_suis__TM_#01
MLWWDDISMSLIDDEIPSSQPQVSGRHF
>gb|Q8FW76|ARO_3003772|Brucella_suis_mprF__Brucella_suis__TM_#02
HSIPDAVPALVAR
>gb|AAG15269_1|ARO_3002528|AAC(3)-Ia__Pseudomonas_aeruginosa__TM_#01
MGIIRTCRLGPDQVKSMRAALDLFGREFGDVATYSQHQPDSDYLGNLLRSKTFIALAAFD
QEAVVGALAAYVLPR
>gb|AAG15269_1|ARO_3002528|AAC(3)-Ia__Pseudomonas_aeruginosa__TM_#02
LLKHEANA
>gb|AAC05822_1|ARO_3002846|arr-1__Mycolicibacterium_smegmatis__TM_#01
AAMREGLEDLRRKGLAVIYD
>gb|AUR80098_1|ARO_3004517|MCR-7_1__Klebsiella_pneumoniae__TM_#01
FFALVLNWPFFLRFYSVISGLEHVRAGFVISVPL
```

>gb|AUR80098_1|ARO_3004517|MCR-7_1__Klebsiella_pneumoniae__TM_#02
ASMIQNIVETNN
>gb|AUR80098_1|ARO_3004517|MCR-7_1__Klebsiella_pneumoniae__TM_#03
MVVLWSLKVRYPANWYKGLAIRAGALAF
>gb|AUR80098_1|ARO_3004517|MCR-7_1__Klebsiella_pneumoniae__TM_#04
PLGTDAKVVAK
>gb|AUR80098_1|ARO_3004517|MCR-7_1__Klebsiella_pneumoniae__TM_#05
RPPYHQRYPDKPPP
>gb|AUR80098_1|ARO_3004517|MCR-7_1__Klebsiella_pneumoniae__TM_#06
STAVYDKQLDIFSQCRTVQ
>gb|ACB05808_1|ARO_3003109|msrE__Acinetobacter_baumannii__TM_#01
VLAEATLQRFGDFAHISQLGGIEIETVEDRAMLSRLGVSNVQNDTMSGG
>gb|ACB05808_1|ARO_3003109|msrE__Acinetobacter_baumannii__TM_#02
LNGIDLLIGQLKA
>gb|ACB05808_1|ARO_3003109|msrE__Acinetobacter_baumannii__TM_#03
LMMKERERLESAVQEKRQQANRLD
>gb|ACB05808_1|ARO_3003109|msrE__Acinetobacter_baumannii__TM_#04
SMGIGANDIQKNLSD
>gb|ACB05808_1|ARO_3003109|msrE__Acinetobacter_baumannii__TM_#05
IAALETMMKSYAGTIIFVSHDKQLVDNIADIIYEIKDHKIIKTFERDC
>gb|AFK80333_1|ARO_3001313|facT__Streptomyces_sp__WAC5292__TM_#01
MNKAGQAE
>gb|AFK80333_1|ARO_3001313|facT__Streptomyces_sp__WAC5292__TM_#02
LAGILSYAWAADRF
>gb|AFK80333_1|ARO_3001313|facT__Streptomyces_sp__WAC5292__TM_#03
GPQAERVAPTVQSVSPRQASDLRDCATEALGQDDLAKVPDICSTLVQGADDGTRDT
>gb|AFK80333_1|ARO_3001313|facT__Streptomyces_sp__WAC5292__TM_#04
VLLLPHHRVRREEPAQ
>gb|AAA25683_1|ARO_3002538|AAC(3)-IIIc__Pseudomonas_aeruginosa__TM_#01
LAYADWEARYE
>gb|AAA25683_1|ARO_3002538|AAC(3)-IIIc__Pseudomonas_aeruginosa__TM_#02
APSVLVDAAAITAFGV
>gb|AAF36804_1|ARO_3002952|vanXF__Paenibacillus_popilliae_ATCC_14706__TM_#01
QLLCSIMEY
>gb|AAF36804_1|ARO_3002952|vanXF__Paenibacillus_popilliae_ATCC_14706__TM_#02
GNHLDPFSNFCGTVPLDALSP
>gb|CAJ98570_1|ARO_3003071|mphF__uncultured_bacterium__TM_#01
MLHDTDRILKLAREAGLELAPGSLRLNEM
>gb|CAJ98570_1|ARO_3003071|mphF__uncultured_bacterium__TM_#02
TDVACAAVKEAKILDYFRSRLPVA
>gb|CAJ98570_1|ARO_3003071|mphF__uncultured_bacterium__TM_#03
SLPGNPGLTFDASTYETTWHFDQNSPVYVETLGAALAQLHGLDTDDAISAGLSNLSIDAV
RENWTRDLETVEKSFEVPAARLALWRAWLADLSFWPTHAASVHGDLYVGHVMVKSDGTVC
GIIDWSEAHIGDPGIDLAGHLKVFGEASLRDLLGHYEAAGGQTWPRIVEHCKMLQSAEGI
RYAMFALKTGSAEHLEGAQGLLSAPGI
>gb|AAM09850_1|ARO_3002944|vanHD__Enterococcus_faecium__TM_#01
AVFRKLSSEYGVTVSLIEDVVSEH
>gb|AAM09850_1|ARO_3002944|vanHD__Enterococcus_faecium__TM_#02
QLGMAVGTVA
>gb|AAM09850_1|ARO_3002944|vanHD__Enterococcus_faecium__TM_#03
TKSVLRGTQKQNYCLNDC
>gb|AAP74657_1|ARO_3000600|Erm(34)__Bacillus_clausii__TM_#01
DRVLAVEYDQKCIEALQWKLVGSKNVSILHQDIMKVALPT
>gb|AAP74657_1|ARO_3000600|Erm(34)__Bacillus_clausii__TM_#02
VSPKDAYVMAWHMWFDIHYERGISRSS
>gb|AAP74657_1|ARO_3000600|Erm(34)__Bacillus_clausii__TM_#03
VRKQHPLFPYKEAKAMHDFLSYALNNPRAPLDQVLRGIFTAPQAKKVRQAIGVKPETPVA
MLHARQWAMVCDAMVRHVPKVYWPRRKR
>gb|KKE03230_1|ARO_3004113|FosA7__Salmonella_enterica_subsp__enterica_serovar_Heidelberg__TM_#01
SLTFWRDLLGLQLHAEW
>gb|KKE03230_1|ARO_3004113|FosA7__Salmonella_enterica_subsp__enterica_serovar_Heidelberg__TM_#02
TGAYLTCGDLW
>gb|KKE03230_1|ARO_3004113|FosA7__Salmonella_enterica_subsp__enterica_serovar_Heidelberg__TM_#03
PYSGMRFGPGK
>gb|AAL68645_1|ARO_3002681|catB9__Vibrio_cholerae__TM_#01
NFADARDGFT
>gb|AAL68645_1|ARO_3002681|catB9__Vibrio_cholerae__TM_#02
ESWLKESMQSLCSSDIEGLYLNWQSKART
>gb|WP_050815728_1|ARO_3004543|mphO__Brachybacterium_paraconglomeratum__TM_#01
MTETSPSSPSSATADAG
>gb|WP_050815728_1|ARO_3004543|mphO__Brachybacterium_paraconglomeratum__TM_#02
SADGTVEWHVDMASTEYARALGTFLAQLHTVDPEEAAATGIPSRTPSEVRGVWREDLTRV
AEAFP
>gb|WP_050815728_1|ARO_3004543|mphO__Brachybacterium_paraconglomeratum__TM_#03
MFHRSSAPPEAFAATLAAYV
>gb|AAD46626_1|ARO_3002596|AAC(6_)-IIc__Pseudomonas_aeruginosa__TM_#01
MSANNAAIVLRVMA
>gb|AAD46626_1|ARO_3002596|AAC(6_)-IIc__Pseudomonas_aeruginosa__TM_#02
SPEVLAKQAVV
>gb|AAD46626_1|ARO_3002596|AAC(6_)-IIc__Pseudomonas_aeruginosa__TM_#03
LRTVQSFKIKGKWS
>gb|ADA78299_1|ARO_3004801|FTU-1__Francisella_tularensis_subsp__tularensis_NE061598__TM_#01
DDSFKNLENKYD
>gb|ADA78299_1|ARO_3004801|FTU-1__Francisella_tularensis_subsp__tularensis_NE061598__TM_#02
YDMHNQGFLDKKIPINQDDIGKLGYAPITAKNVGKTLTISQLNYAAILSDSPASNILVRE
LGGLQNLNKFIKKLGDNDTIITADEPEINYTQPH
>gb|ADA78299_1|ARO_3004801|FTU-1__Francisella_tularensis_subsp__tularensis_NE061598__TM_#03
KDIYKLAFGNILDKKHKDIFIKYLQD
>gb|ADA78299_1|ARO_3004801|FTU-1__Francisella_tularensis_subsp__tularensis_NE061598__TM_#04
KNQQPIALGILYTNPND
>gb|AKI29908_1|ARO_3003603|OXA-447__Campylobacter_jejuni__TM_#01
NKILSFALN
>gb|AKI29908_1|ARO_3003603|OXA-447__Campylobacter_jejuni__TM_#02
IKNIKIREELL
>gb|AKI29908_1|ARO_3003603|OXA-447__Campylobacter_jejuni__TM_#03
NNNRIKSFY
>gb|ADK34116_1|ARO_3001662|OXA-164__Acinetobacter_baumannii__TM_#01
MKLLKILSLVCLSISIGACAEHSMSRAKTSTIPQVNNSIIDQNVQALFNEIS
>gb|ADK34116_1|ARO_3001662|OXA-164__Acinetobacter_baumannii__TM_#02
DAVFVTYDGQNIKKYGTH
>gb|ADK34116_1|ARO_3001662|OXA-164__Acinetobacter_baumannii__TM_#03
PSLMQSELQRIGYGNMQ
>gb|ADK34116_1|ARO_3001662|OXA-164__Acinetobacter_baumannii__TM_#04

IQEVKFVYDLAQ
>gb|ADK34116_1|ARO_3001662|OXA-164__Acinetobacter_baumannii__TM_#05
YVGFVEKADGQVVAFALNMQMK
>gb|CAA33795_1|ARO_3003563|RCP-1__Rhodobacter_capsulatus__TM_#01
LAETPVEALSETVARIEEQLGARVGLSLMETGTG
>gb|CAA33795_1|ARO_3003563|RCP-1__Rhodobacter_capsulatus__TM_#02
RLSLSDALPVRK
>gb|CAA33795_1|ARO_3003563|RCP-1__Rhodobacter_capsulatus__TM_#03
SHTRNLVAVIQ
>gb|APB03223_1|ARO_3003989|AAC(6_)-34__Paenibacillus_sp__LC231__TM_#01
MRIGDLIREDGGIA
>gb|APB03223_1|ARO_3003989|AAC(6_)-34__Paenibacillus_sp__LC231__TM_#02
NESDADCRT
>gb|APB03223_1|ARO_3003989|AAC(6_)-34__Paenibacillus_sp__LC231__TM_#03
MTSMVEYLVF
>gb|APB03223_1|ARO_3003989|AAC(6_)-34__Paenibacillus_sp__LC231__TM_#04
RLLEKHEQ
>gb|AAG05915_1|ARO_3004074|MuxB__Pseudomonas_aeruginosa_PAO1__TM_#01
YSLGEAVEAIRGVEASLELPLSMQG
>gb|AAS48620_1|ARO_3002372|VEB-3__Enterobacter_cloacae__TM_#01
MKIVKRILLVLLSLFFT
>gb|AAS48620_1|ARO_3002372|VEB-3__Enterobacter_cloacae__TM_#02
DNLTLKIENVLKAKNARIGVAIFNSNEKDT
>gb|AAS48620_1|ARO_3002372|VEB-3__Enterobacter_cloacae__TM_#03
KINNDFHFP
>gb|AAS48620_1|ARO_3002372|VEB-3__Enterobacter_cloacae__TM_#04
WATPTAMNKLLIDTYNNK
>gb|AAS48620_1|ARO_3002372|VEB-3__Enterobacter_cloacae__TM_#05
QLIFISVFVAESKETSEINEKIISDIAKITWNYYLNK
>gb|BAG33043_1|ARO_3003920|pgpB__Porphyromonas_gingivalis_ATCC_33277__TM_#01
MEYILEVERNLFLTLNGVQHPLLDGFFYLISAKWTWVIMSIAFLFFLFYKKPTKEAL
>gb|BAG33043_1|ARO_3003920|pgpB__Porphyromonas_gingivalis_ATCC_33277__TM_#02
IDYVKTVYG
>gb|BAG33043_1|ARO_3003920|pgpB__Porphyromonas_gingivalis_ATCC_33277__TM_#03
YISLALFTSRIFRNKFYTWTIWSV
>gb|BAG33043_1|ARO_3003920|pgpB__Porphyromonas_gingivalis_ATCC_33277__TM_#04
IAVGLIVGHFVYKVYLYARSRWLGASCPAHPSAVYAGDSIRLWTLSLIGFVFAMLCMSRQ
LTEILQYYVFLLF
>gb|AAG03548_1|ARO_3003681|TriC__Pseudomonas_aeruginosa_PAO1__TM_#01
SISRDFVDPPT
>gb|AAG03548_1|ARO_3003681|TriC__Pseudomonas_aeruginosa_PAO1__TM_#02
KEGGNILEFGEALNARMQE
>gb|AAG03548_1|ARO_3003681|TriC__Pseudomonas_aeruginosa_PAO1__TM_#03
QKKGRIAERFDSLLHLAMRRRWTTIFLTALLFGV
>gb|AAG03548_1|ARO_3003681|TriC__Pseudomonas_aeruginosa_PAO1__TM_#04
RAVMDRLEATLKDDEDID
>gb|AAG03548_1|ARO_3003681|TriC__Pseudomonas_aeruginosa_PAO1__TM_#05
KMLKIDIA
>gb|ABA71729_1|ARO_3002959|vanYG1__Enterococcus_faecalis__TM_#01
MNHMNMKHRRRKRRRNQSFLFTGILLLVVVSASSFLWYGFGNAAKKDSVIEEMPFTITQD
GMQAKEEIKKTVLETSYGGKQQVAEENHGNTQNAGTDEAWNLMLVNRDNAIPDNYE
>gb|ABA71729_1|ARO_3002959|vanYG1__Enterococcus_faecalis__TM_#02
SLYDEKIAKFKKEGYSDSEAVRQAEQWVAVPGH
>gb|AAX38178_1|ARO_3002634|APH(2__)-Ie__Enterococcus_casseliflavus__TM_#01
TYTFDQVE
>gb|AAX38178_1|ARO_3002634|APH(2__)-Ie__Enterococcus_casseliflavus__TM_#02
AIEQLYPDFTINTIEISGEGNDCIAYEIN
>gb|AAX38178_1|ARO_3002634|APH(2__)-Ie__Enterococcus_casseliflavus__TM_#03
FIFKFPKHSR
>gb|AAX38178_1|ARO_3002634|APH(2__)-Ie__Enterococcus_casseliflavus__TM_#04
LPIPEVVFTGMPSE
>gb|AAX38178_1|ARO_3002634|APH(2__)-Ie__Enterococcus_casseliflavus__TM_#05
QMSFAGFTKIKGVPLTPLLL
>gb|AAX38178_1|ARO_3002634|APH(2__)-Ie__Enterococcus_casseliflavus__TM_#06
QAAKDLARFLSELHSINISGFKSNLVLDFREKINEDNKKIKKLLSRELKG
>gb|AAX38178_1|ARO_3002634|APH(2__)-Ie__Enterococcus_casseliflavus__TM_#07
QMKKVDDFYRDIL
>gb|AAX38178_1|ARO_3002634|APH(2__)-Ie__Enterococcus_casseliflavus__TM_#08
NEIYFKYYPCLIHNDFSSDHILFDTEKNTIC
>gb|AAX38178_1|ARO_3002634|APH(2__)-Ie__Enterococcus_casseliflavus__TM_#09
NDFISLMEDDEEYGMEFVSKILNHYKHKDIPTVLEKY
>gb|AAX38178_1|ARO_3002634|APH(2__)-Ie__Enterococcus_casseliflavus__TM_#10
MKEKYWSFEKIIYGKEYGYMDWYEEGLNEIRSIKIK
>gb|SNU87672_1|ARO_3001459|OXA-154__Pandoraea_sputorum__TM_#01
RAAVVLRLASAVVAHGLLPSPAHALELSRASAAAAPSVA
>gb|SNU87672_1|ARO_3001459|OXA-154__Pandoraea_sputorum__TM_#02
AVGYGNRTIGRVN
>gb|SNU87672_1|ARO_3001459|OXA-154__Pandoraea_sputorum__TM_#03
MNGDADGPKRARIVREVLKNLKLI
>gb|ACX70402_1|ARO_3001654|OXA-143__Acinetobacter_baumannii__TM_#01
MKKFILPI
>gb|ACX70402_1|ARO_3001654|OXA-143__Acinetobacter_baumannii__TM_#02
ALSAVPVYQ
>gb|ACX70402_1|ARO_3001654|OXA-143__Acinetobacter_baumannii__TM_#03
LMQKEVKRV
>gb|ACX70402_1|ARO_3001654|OXA-143__Acinetobacter_baumannii__TM_#04
NRLPFKLETQEEV
>gb|ABD30512_1|ARO_3000839|arlS__Staphylococcus_aureus_subsp__aureus_NCTC_8325__TM_#01
MTKRKLRNNWIIVTTMITFV
>gb|ABD30512_1|ARO_3000839|arlS__Staphylococcus_aureus_subsp__aureus_NCTC_8325__TM_#02
NNLFHSKPV
>gb|ABD30512_1|ARO_3000839|arlS__Staphylococcus_aureus_subsp__aureus_NCTC_8325__TM_#03
VIKKRYKGIE
>gb|ABD30512_1|ARO_3000839|arlS__Staphylococcus_aureus_subsp__aureus_NCTC_8325__TM_#04
EIRRDGFQNKLQLNTNYEEIDNLANTFNEMMS
>gb|ABD30512_1|ARO_3000839|arlS__Staphylococcus_aureus_subsp__aureus_NCTC_8325__TM_#05
ELTKGDVNDISSEAQTVHINDEIR
>gb|ABD30512_1|ARO_3000839|arlS__Staphylococcus_aureus_subsp__aureus_NCTC_8325__TM_#06
VKNKKIKVKTRLKNKQKIIEITDHGIGIPE
>gb|ANZ79476_1|ARO_3003908|Erm(47)__Helcococcus_kunzii__TM_#01
MNRKSVRFGQNFVTSINDINKICKKIDVNSNDVY
>gb|ANZ79476_1|ARO_3003908|Erm(47)__Helcococcus_kunzii__TM_#02
NNKFSDLDNIKIINHDFMSYELPSTFKYKVFGNIPFNLSTSIIRKLSLEKYADEI

```
>gb|ANZ79476_1|ARO_3003908|Erm(47)__Helcococcus_kunzii__TM_#03
EDLNRKMGLMLAPFYEISILYNIPKRYFHPIPSVEVVLIKLKRTSYNMSMKEYIKYEDFI
EKWVKKDYNVLFTKNQLKQAIRYGNIDNLRILKVDQILSIFESYKLFNGLK
>gb|AAU93796_1|ARO_3000594|ErmR__Aeromicrobium_erythreum__Partial_TM_#01
MAGPQDRPRGRGPSSGRPQRPVGGRSQRDRDRRVLGQNFLRDPATIRRIADAADVDPDGL
>gb|AAU93796_1|ARO_3000594|ErmR__Aeromicrobium_erythreum__Partial_TM_#02
GRVRTYELDQRLARRLSTDLAQETS
>gb|AAU93796_1|ARO_3000594|ErmR__Aeromicrobium_erythreum__Partial_TM_#03
PHPEEPFQ
>gb|AAU93796_1|ARO_3000594|ErmR__Aeromicrobium_erythreum__Partial_TM_#04
ALTVTTWPTFEWQYVAKVDRTLFTPVPRVHSAIMRLRRRPQPLLRDAAARSRFADMVEIG
FVGKGGSLYRSLTREWPRSKVDSAFARADVHHDE
>gb|AAU93796_1|ARO_3000594|ErmR__Aeromicrobium_erythreum__Partial_TM_#05
QLLDGSRGGAARGPGDQRGRRGRPGGGPRPDGRAGGGPRRDAGGRRTGDGRGGRPRPPRG
GQA
>gb|SUA92210_1|ARO_3001461|OXA-156__Pandoraea_pulmonicola__TM_#01
SRWRRGAL
>gb|SUA92210_1|ARO_3001461|OXA-156__Pandoraea_pulmonicola__TM_#02
LGALASPVVFA
>gb|SUA92210_1|ARO_3001461|OXA-156__Pandoraea_pulmonicola__TM_#03
VIPWDGKPR
>gb|SUA92210_1|ARO_3001461|OXA-156__Pandoraea_pulmonicola__TM_#04
AFRVSCLPCYQ
>gb|SUA92210_1|ARO_3001461|OXA-156__Pandoraea_pulmonicola__TM_#05
SHKIPRQYAQAKLNE
>gb|SUA92210_1|ARO_3001461|OXA-156__Pandoraea_pulmonicola__TM_#06
GYGNRTIG
>gb|SUA92210_1|ARO_3001461|OXA-156__Pandoraea_pulmonicola__TM_#07
VLKDLKLI
>gb|CAD61201_1|ARO_3002277|VIM-7__Pseudomonas_aeruginosa__TM_#01
MFQIRSFLVGISAFVMAVLGSAAYSAQPGGEYPTVDDIPVGEVRLYKIGDGVWSHIATQK
LGDT
>gb|CAD61201_1|ARO_3002277|VIM-7__Pseudomonas_aeruginosa__TM_#02
LTRQLAEAAGNEVPAHSLKA
>gb|CAD61201_1|ARO_3002277|VIM-7__Pseudomonas_aeruginosa__TM_#03
GDNLVVYVPAVRVLFGGCAVHEASRE
>gb|CAA56561_1|ARO_3003553|CepS_beta-lactamase__Aeromonas_sobria__TM_#01
MKQTRALPLLALGTLLLAPLSLAAPVDPLKAVVDDA
>gb|CAA56561_1|ARO_3003553|CepS_beta-lactamase__Aeromonas_sobria__TM_#02
GGQAHYFNYGLADV
>gb|CAA56561_1|ARO_3003553|CepS_beta-lactamase__Aeromonas_sobria__TM_#03
LKANPVTK
>gb|AJE92936_1|ARO_3004541|mphK__Bacillus_subtilis_subsp__subtilis__TM_#01
TDQISAGQSGIEVIR
>gb|AJE92936_1|ARO_3004541|mphK__Bacillus_subtilis_subsp__subtilis__TM_#02
TTLWERWQKWVDDDA
>gb|AAA50325_1|ARO_3003036|oleB__Streptomyces_antibioticus__TM_#01
MQNAHRSDTGAAALTGTPEKLLPTQPET
>gb|AAA50325_1|ARO_3003036|oleB__Streptomyces_antibioticus__TM_#02
RAPGGCGYLPQT
>gb|AAA50325_1|ARO_3003036|oleB__Streptomyces_antibioticus__TM_#03
RGLREAEQALAGAEPEELEG
>gb|AAA50325_1|ARO_3003036|oleB__Streptomyces_antibioticus__TM_#04
LATGPRRNTERSN
>gb|AAA50325_1|ARO_3003036|oleB__Streptomyces_antibioticus__TM_#05
RARVEGGGTVGRGGALAELYKVTVGTRLDVPSFTVDPGE
>gb|AAA50325_1|ARO_3003036|oleB__Streptomyces_antibioticus__TM_#06
CERPERIGWLPQETEITDRQQSLLAAFAAGLPGIAEEHRGALLGFGLFRPSALG
>gb|AAA50325_1|ARO_3003036|oleB__Streptomyces_antibioticus__TM_#07
FAQRFTGRRMHMEGGRFVE
>gb|AAA23033_2|ARO_3000190|tetO__Campylobacter_jejuni__TM_#01
PMVYREMKAKLSSEIIVKQKVGQHPHINVTDNDDMEQWDAVI
>gb|AAD51348_1|ARO_3003066|smeR__Stenotrophomonas_maltophilia__TM_#01
MSTSPATSTK
>gb|AAD51348_1|ARO_3003066|smeR__Stenotrophomonas_maltophilia__TM_#02
AAAGMASEWVDDGGQVID
>gb|AAD51348_1|ARO_3003066|smeR__Stenotrophomonas_maltophilia__TM_#03
YRPDPGARANGGLHIDEPAARATWNGKGLDLTPVEYRLLRTLLATPGRIWARDELLDRLY
L
>gb|AAD51348_1|ARO_3003066|smeR__Stenotrophomonas_maltophilia__TM_#04
MEGEPIRSVYGMGYSYEP
>gb|BAJ09383_1|ARO_3003046|qacA__Staphylococcus_aureus__TM_#01
MISFFTKTTDMMTSKKRW
>gb|BAJ09383_1|ARO_3003046|qacA__Staphylococcus_aureus__TM_#02
ALVVLAVSLFVVTMDMTILIMALPELVR
>gb|BAJ09383_1|ARO_3003046|qacA__Staphylococcus_aureus__TM_#03
FIIPLSAF
>gb|BAJ09383_1|ARO_3003046|qacA__Staphylococcus_aureus__TM_#04
FAIIAVVAGLFLLPESKLSKEKSHSWDIPSTILSIAGMIGLVWSIKEFSKEGLADIIPWV
V
>gb|BAJ09383_1|ARO_3003046|qacA__Staphylococcus_aureus__TM_#05
PIAPGLAARFGPKIVLPSGIGIAAIGMFIMYFFGHPLSYSTMALALILV
>gb|BAJ09383_1|ARO_3003046|qacA__Staphylococcus_aureus__TM_#06
ASLAVASALIMLETPTS
>gb|BAJ09383_1|ARO_3003046|qacA__Staphylococcus_aureus__TM_#07
MLYRVFLDISSFSSKGI
>gb|AAA88548_1|ARO_3002597|AAC(6_)-Ie-APH(2__)-Ia__Staphylococcus_aureus__TM_#01
KNNPRAIRA
>gb|AAA88548_1|ARO_3002597|AAC(6_)-Ie-APH(2__)-Ia__Staphylococcus_aureus__TM_#02
VNNEYIFKTKFSTNKKKGYAKEKAIYNFLNTN
>gb|AAA88548_1|ARO_3002597|AAC(6_)-Ie-APH(2__)-Ia__Staphylococcus_aureus__TM_#03
LDYTDISEC
>gb|AAA88548_1|ARO_3002597|AAC(6_)-Ie-APH(2__)-Ia__Staphylococcus_aureus__TM_#04
ILLRETIYNDLTDIEKDYIESFMERLNA
>gb|AAA88548_1|ARO_3002597|AAC(6_)-Ie-APH(2__)-Ia__Staphylococcus_aureus__TM_#05
IKQEFIENGRKEIYKRTYKD
>gb|BAA03402_1|ARO_3001306|ErmW__Micromonospora_griseorubida__TM_#01
MSSIRRRHAAASLDT
>gb|BAA03402_1|ARO_3001306|ErmW__Micromonospora_griseorubida__TM_#02
CTRIAEVVSSTTAHPVLELGAGDGAITRALVAANL
>gb|BAA03402_1|ARO_3001306|ErmW__Micromonospora_griseorubida__TM_#03
TFADGVTVVHGDMLRYDFGPYPHHVVSTVPFSITTPLLRRLIGQRF
>gb|BAA03402_1|ARO_3001306|ErmW__Micromonospora_griseorubida__TM_#04
```

SWPWYEFTLVERVPKTSFD
>gb|BAA03402_1|ARO_3001306|ErmW__Micromonospora_griseorubida__TM_#05
SAPLLDDRCVGDYQNLVREVYTGPGRGLAAILRTRLPGREVDAWLRRERVDPAALPRDLK
AGHWASLYRLYREVGTRPAPAGRSVRARPGSVGPDRSLPPRGLRSGPPRARRRGGGA
>gb|AAA26548_1|ARO_3002534|AAC(3)-IIb__Serratia_marcescens__TM_#01
MNTIESITADLHG
>gb|AAA26548_1|ARO_3002534|AAC(3)-IIb__Serratia_marcescens__TM_#02
ASVVSALRA
>gb|CAA42594_1|ARO_3002690|cml__Streptomyces_lividans_1326__TM_#01
GTTSFPVLVAC
>gb|CAA42594_1|ARO_3002690|cml__Streptomyces_lividans_1326__TM_#02
FGVLKAIPAGRATAAATGGPPLRVELAA
>gb|CAA42594_1|ARO_3002690|cml__Streptomyces_lividans_1326__TM_#03
TAAPADATR
>gb|AKQ05893_1|ARO_3004582|tet(49)__uncultured_bacterium__TM_#01
NRGDLVEILIQT
>gb|AKQ05893_1|ARO_3004582|tet(49)__uncultured_bacterium__TM_#02
WIDGIKQS
>gb|AKQ05893_1|ARO_3004582|tet(49)__uncultured_bacterium__TM_#03
GLVIGADGLHSKTRRMVFNEDEYKLTNLGL
>gb|AKQ05893_1|ARO_3004582|tet(49)__uncultured_bacterium__TM_#04
SFVEANQKLGILVNESTLVYGEVSQ
>gb|AKQ05893_1|ARO_3004582|tet(49)__uncultured_bacterium__TM_#05
EIAANMISLPNYE
>gb|ABY64751_1|ARO_3002667|rmtD__Klebsiella_pneumoniae__TM_#01
RECSAKFKKEKD
>gb|ABY64751_1|ARO_3002667|rmtD__Klebsiella_pneumoniae__TM_#02
DKAAREALHGVTGAFMTEREYKRAME
>gb|ABY64751_1|ARO_3002667|rmtD__Klebsiella_pneumoniae__TM_#03
RDWEALLGM
>gb|ABY64751_1|ARO_3002667|rmtD__Klebsiella_pneumoniae__TM_#04
SMDRVFDQLFEA
>gb|ABY64751_1|ARO_3002667|rmtD__Klebsiella_pneumoniae__TM_#05
HRLPNAAI
>gb|ABY64751_1|ARO_3002667|rmtD__Klebsiella_pneumoniae__TM_#06
GVDISGQCVNVIRAFGGAEARLGDLLCEIPEDEA
>gb|ABY64751_1|ARO_3002667|rmtD__Klebsiella_pneumoniae__TM_#07
FKVLPLLERQR
>gb|ABY64751_1|ARO_3002667|rmtD__Klebsiella_pneumoniae__TM_#08
ALMRVNAEWI
>gb|ABY64751_1|ARO_3002667|rmtD__Klebsiella_pneumoniae__TM_#09
MEAHVPENRAIAARLTGENELFYVLKRK
>gb|WP_045475347_1|ARO_3005334|Trimethoprim-resistant_dihydrofolate_reductase_DfrA42__Gammaproteobacteria__Partial_TM_#01
MHKPTPLKRLSMILARDLNGAIGYEGSLAIKSDNDFAWYKKITKPFKHAVCGRVTYEEDL
PDIVKKRHRFIIITRNPDKYASTPEAQYMTLSDALKELQVWDVPGHDIICLGGAEIYKAL
LPYVSTVYLTTFFSVADEADTYFNDFTDKWESVGATWFNDCYCTRLERVCQSTNNII
>gb|AIG19992_1|ARO_3005039|PDC-55__Pseudomonas_aeruginosa__TM_#01
CLCGIAASTLLFA
>gb|AIG19992_1|ARO_3005039|PDC-55__Pseudomonas_aeruginosa__TM_#02
EAPADRLKALVDAAVQPVMKANDIPGLAVAISLKGEPHYFSYGLASKEDG
>gb|AIG19992_1|ARO_3005039|PDC-55__Pseudomonas_aeruginosa__TM_#03
QDKMRLDDRAS
>gb|AIG19992_1|ARO_3005039|PDC-55__Pseudomonas_aeruginosa__TM_#04
HWPALQGSR
>gb|AIG19992_1|ARO_3005039|PDC-55__Pseudomonas_aeruginosa__TM_#05
FPALGLEQTHLDVPEAALAQYAQGYGKDDRPLR
>gb|AIG19992_1|ARO_3005039|PDC-55__Pseudomonas_aeruginosa__TM_#06
HPERLDRPWAQALD
>gb|CCF55073_1|ARO_3003205|Erm(43)__Staphylococcus_lentus__TM_#01
CINEILKNIIIT
>gb|CCF55073_1|ARO_3003205|Erm(43)__Staphylococcus_lentus__TM_#02
ALSKVVKSVIGVEIDKSLYYNLKKDSKLQDNIQLINQDILNFQFPDNKD
>gb|CCF55073_1|ARO_3003205|Erm(43)__Staphylococcus_lentus__TM_#03
QDKNKALSLLLLPKMDVEILKVIPNK
>gb|CCF55073_1|ARO_3003205|Erm(43)__Staphylococcus_lentus__TM_#04
KHKPLISAKDEKN
>gb|CCF55073_1|ARO_3003205|Erm(43)__Staphylococcus_lentus__TM_#05
KNANVQNLNKISKQQFISIFYSYKLFN
>gb|AAB51440_1|ARO_3002824|ErmV__Streptomyces_viridochromogenes__TM_#01
DAPHVRVLGEDFLR
>gb|AAB51440_1|ARO_3002824|ErmV__Streptomyces_viridochromogenes__TM_#02
DVLVGEVPPWTWLRLHEVLGS
>gb|CAJ77857_1|ARO_3000778|adeG__Acinetobacter_baumannii_AYE__TM_#01
IQRLIQSNAVSRQELDL
>gb|AAS79458_1|ARO_3001302|chrB__Streptomyces_bikiniensis__TM_#01
GVMEAFPDAQ
>gb|AAS79458_1|ARO_3001302|chrB__Streptomyces_bikiniensis__TM_#02
RGVLLVVTPLPDHLREVIGALGLLQVDEG
>gb|AAS79458_1|ARO_3001302|chrB__Streptomyces_bikiniensis__TM_#03
LETIQRLPAPTRVTLSVRLSAYRLSA
>gb|AHG97174_1|ARO_3003011|dfrA10__Klebsiella_pneumoniae__TM_#01
MNISLIFANELITRAFGNQGKLPWQFIKEDMQFFQKTTENSVVVMGLNTWRSLPKMKKLG
RDFIVISSTITEHEVLNNNIQIFKSFESFLEAFRDTTKPINVIGGVGLLSEAIEHASTVY
MSSIHMVKPVHADVYVPVELMNKLYSDFKYPENILWVGDPIDSVYSLSIDKFVRPASLVG
VPNDINT
>gb|ABC68330_1|ARO_3002640|APH(3__)-Ic__Mycolicibacterium_fortuitum__TM_#01
NSDGSSYAKVVD
>gb|ABC68330_1|ARO_3002640|APH(3__)-Ic__Mycolicibacterium_fortuitum__TM_#02
RHGVPGPAVIDWRVTEDGGACLITSTVRGVAADRLSESALRAAWPAIVEAVRTLHALPAD
G
>gb|ABC68330_1|ARO_3002640|APH(3__)-Ic__Mycolicibacterium_fortuitum__TM_#03
GAGAVNPEFLSDEDREVPAEALLD
>gb|ANW35663_1|ARO_3003620|OXA-464__Arcobacter_butzleri__TM_#01
QVEGTLVLESLNTKKVDIYNEKRANTS
>gb|ANW35663_1|ARO_3003620|OXA-464__Arcobacter_butzleri__TM_#02
KDSIIVWDKKVREFD
>gb|ANW35663_1|ARO_3003620|OXA-464__Arcobacter_butzleri__TM_#03
TIGKDVTDFWLDESLRITAFEEIRFLKQLQANNLAFKQEDINLLKELMIDEKSENYVVR
>gb|ANW35663_1|ARO_3003620|OXA-464__Arcobacter_butzleri__TM_#04
LNIDTKTKEDLAKRKALTLEALKTKGIID
>gb|CAA26428_1|ARO_3002630|ANT(9)-Ia__Staphylococcus_aureus__TM_#01
MSNLINGKIPNQAIQTLKIVKDLF
>gb|CAA26428_1|ARO_3002630|ANT(9)-Ia__Staphylococcus_aureus__TM_#02

DSSSILVS
>gb|CAA26428_1|ARO_3002630|ANT(9)-Ia__Staphylococcus_aureus__TM_#03
EWAIPLLPKEHVTLLDIARKGYRGECDDKWEGLYSKVKALVKYMKNSIETSLN
>gb|AAA21532_1|ARO_3003559|cepA_beta-lactamase__Bacteroides_fragilis__TM_#01
MQKRLIHLSIIFFLLCPALVVAQNSPLETQLKK
>gb|AAA21532_1|ARO_3003559|cepA_beta-lactamase__Bacteroides_fragilis__TM_#02
HHQKQPLETRLL
>gb|AAA21532_1|ARO_3003559|cepA_beta-lactamase__Bacteroides_fragilis__TM_#03
PDAVNKYLHSLGIRECAVIHTENDMHKNLEF
>gb|AAA21532_1|ARO_3003559|cepA_beta-lactamase__Bacteroides_fragilis__TM_#04
ADNRENSEIIAEISRIVYEYVTQQID
>gb|CAA40146_1|ARO_3002997|LCR-1__Pseudomonas_aeruginosa__TM_#01
MLKSTLLAFGLFIALSARAENQAIA
>gb|CAA40146_1|ARO_3002997|LCR-1__Pseudomonas_aeruginosa__TM_#02
RAGVDGTIVIESLTT
>gb|CAA40146_1|ARO_3002997|LCR-1__Pseudomonas_aeruginosa__TM_#03
QRLVHNDPRAQQRYP
>gb|CAA40146_1|ARO_3002997|LCR-1__Pseudomonas_aeruginosa__TM_#04
ENQIFHWNGTQYSIAN
>gb|CAA40146_1|ARO_3002997|LCR-1__Pseudomonas_aeruginosa__TM_#05
LRVGALKYPAYIQQTNYGHLL
>gb|CAA40146_1|ARO_3002997|LCR-1__Pseudomonas_aeruginosa__TM_#06
DSLKKVMFADENAQYRLYAKTGWATR
>gb|CAA40146_1|ARO_3002997|LCR-1__Pseudomonas_aeruginosa__TM_#07
TPSVGWYVGYVEA
>gb|AIL46641_1|ARO_3000579|Chryseobacterium_meningosepticum_BlaB__Elizabethkingia_anophelis_NUHP1__TM_#01
KAFGQENPDVKIEKLKDNLYVYTTYNTFNGTKYA
>gb|AIL46641_1|ARO_3000579|Chryseobacterium_meningosepticum_BlaB__Elizabethkingia_anophelis_NUHP1__TM_#02
CPWGEDKFKSFTDEIYKKHGKKVIMNI
>gb|AIL46641_1|ARO_3000579|Chryseobacterium_meningosepticum_BlaB__Elizabethkingia_anophelis_NUHP1__TM_#03
GLEYFGKI
>gb|AIL46641_1|ARO_3000579|Chryseobacterium_meningosepticum_BlaB__Elizabethkingia_anophelis_NUHP1__TM_#04
SKDLGYIGEAYVNDWTQSVHNIQQKFSGA
>gb|AAM77075_1|ARO_3002695|cmlA5__uncultured_bacterium__TM_#01
GWRAIFAFLGLGMIAA
>gb|AAM77075_1|ARO_3002695|cmlA5__uncultured_bacterium__TM_#02
WPETRVQR
>gb|AAM77075_1|ARO_3002695|cmlA5__uncultured_bacterium__TM_#03
LQWSQLLLP
>gb|AAM77075_1|ARO_3002695|cmlA5__uncultured_bacterium__TM_#04
AGMGSFFVFF
>gb|AAM77075_1|ARO_3002695|cmlA5__uncultured_bacterium__TM_#05
FSLLFATVAIAM
>gb|AAM77075_1|ARO_3002695|cmlA5__uncultured_bacterium__TM_#06
ARFMGRVI
>gb|AAM77075_1|ARO_3002695|cmlA5__uncultured_bacterium__TM_#07
LRMGMGCLIAGAVLL
>gb|AAM77075_1|ARO_3002695|cmlA5__uncultured_bacterium__TM_#08
QSVLGFIAPMWLVG
>gb|AAM77075_1|ARO_3002695|cmlA5__uncultured_bacterium__TM_#09
GVATAVSV
>gb|AAM77075_1|ARO_3002695|cmlA5__uncultured_bacterium__TM_#10
VLGLSCVSR
>gb|AAA65954_1|ARO_3002931|vanSA__Enterococcus_faecium__TM_#01
LSILENKYDLNHLDAMKLYQYSIRNNI
>gb|AAA65954_1|ARO_3002931|vanSA__Enterococcus_faecium__TM_#02
QIELSAEMDV
>gb|AAA65954_1|ARO_3002931|vanSA__Enterococcus_faecium__TM_#03
RVELPAMPDLVDKRRS
>gb|CAA30578_1|ARO_3002652|APH(3_)-VIa__Acinetobacter_baumannii__TM_#01
MELPNIIQQFIGNSVLEP
>gb|CAA30578_1|ARO_3002652|APH(3_)-VIa__Acinetobacter_baumannii__TM_#02
SFNRNNETFFLKRSSTLYTETTYSVSREAKMLSWLS
>gb|CAA30578_1|ARO_3002652|APH(3_)-VIa__Acinetobacter_baumannii__TM_#03
KLKVPELIMTFQDEQFE
>gb|CAA30578_1|ARO_3002652|APH(3_)-VIa__Acinetobacter_baumannii__TM_#04
MITKAINAK
>gb|CAA30578_1|ARO_3002652|APH(3_)-VIa__Acinetobacter_baumannii__TM_#05
QELLAIYKE
>gb|CAA30578_1|ARO_3002652|APH(3_)-VIa__Acinetobacter_baumannii__TM_#06
AIIDCPFIS
>gb|CAA30578_1|ARO_3002652|APH(3_)-VIa__Acinetobacter_baumannii__TM_#07
IDHRLKESKFFIDNQLLD
>gb|CAA30578_1|ARO_3002652|APH(3_)-VIa__Acinetobacter_baumannii__TM_#08
ETRVEERLVFSHGDITDSNIFIDK
>gb|CAA30578_1|ARO_3002652|APH(3_)-VIa__Acinetobacter_baumannii__TM_#09
KIFLKHLKND
>gb|AWB15813_1|ARO_3005096|GPC-1__Pseudomonas_aeruginosa__TM_#01
TAGTSRAAGENLAQRLAALEARHG
>gb|AWB15813_1|ARO_3005096|GPC-1__Pseudomonas_aeruginosa__TM_#02
SAMVETLRRLLFTDVLSAR
>gb|ASK40551_1|ARO_3004332|MCR-5__Salmonella_enterica_subsp__enterica_serovar_Paratyphi_B__TM_#01
MRLSAFITFLKMRPQVRTEFLTLFISLVFTLLCNGVFWNALLAGRDSLTSGTWLMLLCTG
LLITGLQWLLLLLVATRWSVKPLLILLAVMTPAAVYFMRNYGVYLDKAMLRNLMETDVRE
ASELLQWRMLPYLLVAAVSVWWIARVRVLRTGWKQAVMMRSA
>gb|ASK40551_1|ARO_3004332|MCR-5__Salmonella_enterica_subsp__enterica_serovar_Paratyphi_B__TM_#02
GIRVLTEQASSSADEAREVVAADAHRGPQEQG
>gb|ASK40551_1|ARO_3004332|MCR-5__Salmonella_enterica_subsp__enterica_serovar_Paratyphi_B__TM_#03
LNGRRDYDERQIRRRESVLHVLNRSD
>gb|ASK40551_1|ARO_3004332|MCR-5__Salmonella_enterica_subsp__enterica_serovar_Paratyphi_B__TM_#04
SSAGHPTLCHGERCLDEILLEGLAEKITTSRSDM
>gb|ASK40551_1|ARO_3004332|MCR-5__Salmonella_enterica_subsp__enterica_serovar_Paratyphi_B__TM_#05
RRWSPTCDTTDLASCSHEAL
>gb|ASK40551_1|ARO_3004332|MCR-5__Salmonella_enterica_subsp__enterica_serovar_Paratyphi_B__TM_#06
TAAYTPELDLLATCRKGQPQ
>gb|AAB84282_1|ARO_3000181|tet(V)__Mycolicibacterium_smegmatis_MC2_155__TM_#01
VVNFVTVAVISALALLGVLKIWHMAVAAGI
>gb|AAB84282_1|ARO_3000181|tet(V)__Mycolicibacterium_smegmatis_MC2_155__TM_#02
ATAGLVPVAIAAVAFTAARMHRDEVANPLL
>gb|AAP97895_1|ARO_3005111|blaS1__Mycolicibacterium_smegmatis_MC2_155__TM_#01
DDRIADLERRNNASIGIYAVDLDSN
>gb|AAP97895_1|ARO_3005111|blaS1__Mycolicibacterium_smegmatis_MC2_155__TM_#02
VLTGDVLAPPQRQLLDE

>gb|AAP97895_1|ARO_3005111|blaS1__Mycolicibacterium_smegmatis_MC2_155__TM_#03
VRTRGDDPNADGF
>gb|AAZ42322|ARO_3005098|qacL__Vibrio_cholerae__TM_#01
MKNWIFMAVAIF
>gb|AAZ42322|ARO_3005098|qacL__Vibrio_cholerae__TM_#02
LVPSVVVVA
>gb|AAZ42322|ARO_3005098|qacL__Vibrio_cholerae__TM_#03
FWAFIGMGLIVSGVAVLNLLSKVSAH
>gb|AKQ05891_1|ARO_3004613|Tet(47)__uncultured_bacterium__TM_#01
MRCHFRCNLALPLLVFLNLSRAIKDYQT
>gb|AKQ05891_1|ARO_3004613|Tet(47)__uncultured_bacterium__TM_#02
GNRRTQVEFGRYMDTQ
>gb|AKQ05891_1|ARO_3004613|Tet(47)__uncultured_bacterium__TM_#03
CYREGEDVEIT
>gb|AKQ05891_1|ARO_3004613|Tet(47)__uncultured_bacterium__TM_#04
YLNQSIDSIKQRDGDVAVRFKDGKIEY
>gb|CAB06660_1|ARO_3004644|dfrA6_from_Proteus_mirabilis__Proteus_mirabilis__TM_#01
QADVIHLSVIHKHISGDVFFP
>gb|CAB06660_1|ARO_3004644|dfrA6_from_Proteus_mirabilis__Proteus_mirabilis__TM_#02
FKQTFEQS
>gb|CAE51638_1|ARO_3002641|APH(3_)-Ia__Serratia_marcescens__TM_#01
NSNLDADLY
>gb|CAE51638_1|ARO_3002641|APH(3_)-Ia__Serratia_marcescens__TM_#02
ELFLKHGKGSV
>gb|AAB95639_1|ARO_3000118|vgaB__Staphylococcus_aureus__TM_#01
MLKIDMKNVKKYYADKLILNIKELKIYS
>gb|AAB95639_1|ARO_3000118|vgaB__Staphylococcus_aureus__TM_#02
KGLIEIDEGNIIISEKTTIKYISQLEEPHSKIIDGKYASIFQVENK
>gb|AAB95639_1|ARO_3000118|vgaB__Staphylococcus_aureus__TM_#03
DQCSLMLV
>gb|AAB95639_1|ARO_3000118|vgaB__Staphylococcus_aureus__TM_#04
TNTFKEYRDT
>gb|AAB95639_1|ARO_3000118|vgaB__Staphylococcus_aureus__TM_#05
KIFEIENGYIREFIGNYTNYIEQKEMLLRKQQEEYEKYNSKRKQLEQAIKLKENKAQGMI
KPPSKTMGTSESRIWKMQHATKQKKMHRNTKSLETRIDKLNHVEKIKELPSIKMDLPNRE
QFHGRNVISLKNLSIKFNNQF
>gb|AAB95639_1|ARO_3000118|vgaB__Staphylococcus_aureus__TM_#06
EKVESVIISPSVKIGYVSQNLDVLQSHK
>gb|AAB95639_1|ARO_3000118|vgaB__Staphylococcus_aureus__TM_#07
IARIVLARLHFYRNDVH
>gb|AAB95639_1|ARO_3000118|vgaB__Staphylococcus_aureus__TM_#08
EGVVLFASHDKKFIQNLAEQLL
>gb|AAA26832_1|ARO_3002827|tlrC__Streptomyces_fradiae__TM_#01
VVLSQVSLAI
>gb|AAA26832_1|ARO_3002827|tlrC__Streptomyces_fradiae__TM_#02
GLPPRATVQDAIDLAMTELRVLEAELRRTEAALAEAATD
>gb|AAA26832_1|ARO_3002827|tlrC__Streptomyces_fradiae__TM_#03
RLSLTARIATADGPGEAPAAELDGVVVGSRLRVPK
>gb|AAA26832_1|ARO_3002827|tlrC__Streptomyces_fradiae__TM_#04
KLTVLEAFAHNRPGDRDEQADRR
>gb|AAA26832_1|ARO_3002827|tlrC__Streptomyces_fradiae__TM_#05
REGVVSGAR
>gb|AAT46346_1|ARO_3004292|Laribacter_hongkongensis_ampC_beta-lactamase__Laribacter_hongkongensis__TM_#01
MKKRITPFSRFASKGLFACSAGMLLVTVAHAANTAAAPAGMDAMVQTVMQAHQIPGMAIA
IIQPGKTTYHNYGVASRETGQPVRE
>gb|AAT46346_1|ARO_3004292|Laribacter_hongkongensis_ampC_beta-lactamase__Laribacter_hongkongensis__TM_#02
PFTALVAQRAETEGRIDLSAPASRYVAALRGSAFDRI
>gb|AAT46346_1|ARO_3004292|Laribacter_hongkongensis_ampC_beta-lactamase__Laribacter_hongkongensis__TM_#03
LAASQATGESFAGLLGTTVLHPLGMNSTYLQVPPEARSRYAMGYTAAGKAVRVSPGPLDE
ETYGVKSTTADMAGFLLAHMDPARSKGALQSALQQTRVPVYCAGQTRQGLGWESYQDWKN
LDVLLAGNSNQMVFEPQPVKACPAGTMNDPDVWV
>gb|AAT46346_1|ARO_3004292|Laribacter_hongkongensis_ampC_beta-lactamase__Laribacter_hongkongensis__TM_#04
RLAHGILTALH
>gb|AGZ55247_1|ARO_3002814|clbA__Bacillus_amyloliquefaciens_CC178__TM_#01
MQQKNKYIRIQEFLKQNKFP
>gb|AGZ55247_1|ARO_3002814|clbA__Bacillus_amyloliquefaciens_CC178__TM_#02
FNEITVLPKSLR
>gb|AGZ55247_1|ARO_3002814|clbA__Bacillus_amyloliquefaciens_CC178__TM_#03
LIEEFGESILNI
>gb|AGZ55247_1|ARO_3002814|clbA__Bacillus_amyloliquefaciens_CC178__TM_#04
LFALSPRRLSISTIGIIP
>gb|AGZ55247_1|ARO_3002814|clbA__Bacillus_amyloliquefaciens_CC178__TM_#05
LMPINERYPL
>gb|AGZ55247_1|ARO_3002814|clbA__Bacillus_amyloliquefaciens_CC178__TM_#06
VMDTLDEHIRVTS
>gb|AGZ55247_1|ARO_3002814|clbA__Bacillus_amyloliquefaciens_CC178__TM_#07
HVNIIRYNPT
>gb|AFJ11385_1|ARO_3002666|rmtF__Klebsiella_pneumoniae__TM_#01
KKARALLARWNEGDESALAA
>gb|AFJ11385_1|ARO_3002666|rmtF__Klebsiella_pneumoniae__TM_#02
PGADEWMRRVSPFLGADA
>gb|AFJ11385_1|ARO_3002666|rmtF__Klebsiella_pneumoniae__TM_#03
ILLGSMGVTNALGMDIHLGCVRLVNETARARGWHTRARACDLLSEIPAEEADA
>gb|AFJ11385_1|ARO_3002666|rmtF__Klebsiella_pneumoniae__TM_#04
RAAELLASLRAPRLVVTFPTRTLGGRGVG
>gb|AAP69916_1|ARO_3001780|OXA-85__Fusobacterium_nucleatum__TM_#01
MLLFMFSIISFGNENQFMKEIFERKGLNGTFVVYDLKNDKIDYYNLDRANERFYP
>gb|AAP69916_1|ARO_3001780|OXA-85__Fusobacterium_nucleatum__TM_#02
GLENGIVKNVDEMFYYYDGS
>gb|AAP69916_1|ARO_3001780|OXA-85__Fusobacterium_nucleatum__TM_#03
KKLARELGKERMQEGLNKLNYGNKEIGSEIDK
>gb|AAP69916_1|ARO_3001780|OXA-85__Fusobacterium_nucleatum__TM_#04
NLLSQSKLPFKLENQEQVKDITILEKKDDFILHGKTGWATDNIVVPIGWFVGWIETSDNI
YSFAINLDISDSKFLPKREEIVREYFKNINVIK
>gb|AAL75563_1|ARO_3000168|tet(D)__Shigella_flexneri_Y__TM_#01
LMVFFIFKPAVQTEEKPAEQKQ
>gb|AAL75563_1|ARO_3000168|tet(D)__Shigella_flexneri_Y__TM_#02
KRLSEKTIIFAGFIADATAFLLMSAITS
>gb|AAL75563_1|ARO_3000168|tet(D)__Shigella_flexneri_Y__TM_#03
GLLLAICLLIRKPAPVAATC
>gb|WP_102607462_1|ARO_3005324|OXA-570__Ralstonia_insidiosa__TM_#01
RWSKTFALAL
>gb|WP_102607462_1|ARO_3005324|OXA-570__Ralstonia_insidiosa__TM_#02

FEEARFTSRLALKQLPVKPRTWDL
>gb|WP_102607462_1|ARO_3005324|OXA-570__Ralstonia_insidiosa__TM_#03
LEADMAKRI
>gb|WP_102607462_1|ARO_3005324|OXA-570__Ralstonia_insidiosa__TM_#04
LGKRLMQAL
>gb|AAK63041_1|ARO_3004629|AAC(6_)-Im__Escherichia_coli__TM_#01
MLEKKRVSFRPMNEDDLV
>gb|AAK63041_1|ARO_3004629|AAC(6_)-Im__Escherichia_coli__TM_#02
HTQKTIREHYTEQWADEIYRVIIEYDTIPIGYAQIYRIQGELFDEYNYHETEE
>gb|AAK63041_1|ARO_3004629|AAC(6_)-Im__Escherichia_coli__TM_#03
AEYCRVVCQYLRTEMDADAVILDPRKNNLRAVRAYQKAGFK
>gb|AAK99775_1|ARO_3000614|mefE__Streptococcus_pneumoniae_R6__TM_#01
MKIDKKNE
>gb|AAK99775_1|ARO_3000614|mefE__Streptococcus_pneumoniae_R6__TM_#02
SFIVPVNDEVVTKDKMTIGGVLN
>gb|AAK99775_1|ARO_3000614|mefE__Streptococcus_pneumoniae_R6__TM_#03
VINKTLSSKRLMVFLSCSGLMLMLSTPLYFLFQNF
>gb|AHA41501_1|ARO_3002954|vanXO__Rhodococcus_hoagii__TM_#01
KPIESKNREL
>gb|ACD45689_1|ARO_3004550|dfrA27__Vibrio_cholerae_non-O1_non-O139__TM_#01
MKISLMAAKARNGVIGC
>gb|ACD45689_1|ARO_3004550|dfrA27__Vibrio_cholerae_non-O1_non-O139__TM_#02
VVFPSIEAAM
>gb|ACD45689_1|ARO_3004550|dfrA27__Vibrio_cholerae_non-O1_non-O139__TM_#03
LKTLTNHVVVSGGGEIYKSLIAHADTLHISTIDSEPEGNVFFPEIPK
>gb|ACD45689_1|ARO_3004550|dfrA27__Vibrio_cholerae_non-O1_non-O139__TM_#04
FNVVFEQEFHSNINYRYQIWQRG
>gb|AAO83986_1|ARO_3002623|ANT(4_)-Ia__Bacillus_clausii__TM_#01
MNMNGPASMAQKERLQTCQEIAKRLHEVYGND
>gb|AAO83986_1|ARO_3002623|ANT(4_)-Ia__Bacillus_clausii__TM_#02
KDAATVEDR
>gb|AAO83986_1|ARO_3002623|ANT(4_)-Ia__Bacillus_clausii__TM_#03
EGFFQRLRL
>gb|AAO83986_1|ARO_3002623|ANT(4_)-Ia__Bacillus_clausii__TM_#04
NRNGPSTYLPSLALRFAHYGAMLIGLHNQT
>gb|AAO83986_1|ARO_3002623|ANT(4_)-Ia__Bacillus_clausii__TM_#05
HRPKGFDHVAELAMSGDLAQPAKIVSACEDFWKGLVAWAAEHDYVIHSKRIPF
>gb|AAC37034_1|ARO_3000522|ErmG__Bacteroides_thetaiotaomicron__TM_#01
SKLCEVTRNKLLNYPNYQIVNDDILKFTFPSHNP
>gb|AAC37034_1|ARO_3000522|ErmG__Bacteroides_thetaiotaomicron__TM_#02
PAKMAFKERKK
>gb|AAC37034_1|ARO_3000522|ErmG__Bacteroides_thetaiotaomicron__TM_#03
YDINNISFEQFVSLFNSYKIFNG
>gb|AEG78825_1|ARO_3002872|FosA3__Escherichia_coli__TM_#01
ASSLAFYQ
>gb|AAA99504_1|ARO_3002987|bcrA__Bacillus_licheniformis__TM_#01
KKRNRKYLEFQLSDQNKAVVLMEQHFDIHDYEVHQDGI
>gb|AAN34365_1|ARO_3002604|aadA4__Acinetobacter_baumannii__TM_#01
MGEFFPAQ
>gb|AAN34365_1|ARO_3002604|aadA4__Acinetobacter_baumannii__TM_#02
LLVTVSAAPNDSLRQALMLDLLKVSSPPG
>gb|AAN34365_1|ARO_3002604|aadA4__Acinetobacter_baumannii__TM_#03
EHFSKALFDTI
>gb|AAN34365_1|ARO_3002604|aadA4__Acinetobacter_baumannii__TM_#04
AMRVEETAAFVRYAKATIERILR
>gb|CAA37806_1|ARO_3002684|catII__Haemophilus_influenzae__TM_#01
MNFTRIDLNTWNRREHFALYRQ
>gb|CAA37806_1|ARO_3002684|catII__Haemophilus_influenzae__TM_#02
VNQFPEFRMA
>gb|CAA37806_1|ARO_3002684|catII__Haemophilus_influenzae__TM_#03
SEFMAGYNAV
>gb|CAA37806_1|ARO_3002684|catII__Haemophilus_influenzae__TM_#04
TGNDDYFAPVFTMAKFQQE
>gb|CAH14032_1|ARO_3004099|LpeA__Legionella_pneumophila_str__Paris__TM_#01
YSKVTPESIPNKLVEVEPIKSHN
>gb|CAH14032_1|ARO_3004099|LpeA__Legionella_pneumophila_str__Paris__TM_#02
NPDLEKNLQLSLSAVELAKAQYERITPLI
>gb|CAH14032_1|ARO_3004099|LpeA__Legionella_pneumophila_str__Paris__TM_#03
ATLNEGQPVYVLGKR
>gb|CAH14032_1|ARO_3004099|LpeA__Legionella_pneumophila_str__Paris__TM_#04
DDCLIGATTSVELLVAEKNNTIVIPFQAIFLRNSK
>gb|CAH14032_1|ARO_3004099|LpeA__Legionella_pneumophila_str__Paris__TM_#05
EDKIEIVEGLKAGQQLVTKGQERLYPEMTVDIYHPATSSS
>gb|AAL05554_1|ARO_3003206|lsaE__Enterococcus_faecalis__TM_#01
MLKQKELIANVKNLTES
>gb|AAL05554_1|ARO_3003206|lsaE__Enterococcus_faecalis__TM_#02
SITSNFDSSNWLFDVAPYLMLYK
>gb|AAL05554_1|ARO_3003206|lsaE__Enterococcus_faecalis__TM_#03
NYLSEISGAR
>gb|AAL05554_1|ARO_3003206|lsaE__Enterococcus_faecalis__TM_#04
LENYKKFAKTTARLDKVELFEAYKNSLLLVMDLQSHLIEQYNLKVTHDILERLLNYISE
>gb|BAB72072_1|ARO_3002202|IMP-11__Acinetobacter_baumannii__TM_#01
LFGGCFVKPYGLGNL
>gb|BAB72072_1|ARO_3002202|IMP-11__Acinetobacter_baumannii__TM_#02
SLLKLTWEQ
>gb|EOR02560_1|ARO_3004479|OXA-664__Acinetobacter_tandoii_DSM_14970___CIP_107469__TM_#01
AVVFVTYDGEKLQRFGNDLH
>gb|EOR02560_1|ARO_3004479|OXA-664__Acinetobacter_tandoii_DSM_14970___CIP_107469__TM_#02
TTEVFVWDGKARALK
>gb|EOR02560_1|ARO_3004479|OXA-664__Acinetobacter_tandoii_DSM_14970___CIP_107469__TM_#03
TLARRIGLPLMQKELHRVD
>gb|EOR02560_1|ARO_3004479|OXA-664__Acinetobacter_tandoii_DSM_14970___CIP_107469__TM_#04
ADVKPQVG
>gb|AAA22851_1|ARO_3000179|tet(L)__Geobacillus_stearothermophilus__TM_#01
LVTLNVYKHSQRDF
>gb|CAB61635_1|ARO_3004788|FONA-1__Serratia_fonticola__TM_#01
ANAKANIQQQLSELEK
>gb|CAB61635_1|ARO_3004788|FONA-1__Serratia_fonticola__TM_#02
TDKNLLAKRMEIKQ
>gb|AAF67494_2|ARO_3002522|novA__Streptomyces_niveus__TM_#01
MKSALSTWKPSDRPPDPTLPEP
>gb|AAF67494_2|ARO_3002522|novA__Streptomyces_niveus__TM_#02
GAPAPAPVPARDERVGAA

>gb|AUW34365_1|ARO_3004444|RSA-1__uncultured_bacterium__TM_#01
MGMQLARSTILTVLLCLPIAVTATTKEEIQKIERQRNLTVGIALVDDGGTLL
>gb|AUW34365_1|ARO_3004444|RSA-1__uncultured_bacterium__TM_#02
TLKQIESGKWSAAERLSYSAGQLDAYAPAAKRYLPTGYITVAEANQASVQ
>gb|AUW34365_1|ARO_3004444|RSA-1__uncultured_bacterium__TM_#03
RIVAKLVYGNYLSTAGREQLQRLLIGNNTGDSRIRAGIASGWTTGDKTGSCPNGGRNDAA
FLVSPDGRRFALTVYLNAPSLDDKARNEVVATVARLAVESIR
>gb|ABU39980_1|ARO_3004749|BCL-1__Bacillus_clausii__TM_#01
MKRSFFMLKTKITSSILVGACLLIGCSNGNEQPVSNEPEPEESVETGEAVFKALEEEYAA
RLGVFALDTGTGQTVSYRSDERFTYASAHKPLAVAVLLQQKSIEELEQLITYSAD
>gb|ABU39980_1|ARO_3004749|BCL-1__Bacillus_clausii__TM_#02
IQDTSTPEALAK
>gb|ABU39980_1|ARO_3004749|BCL-1__Bacillus_clausii__TM_#03
KEPIILAVLSSKDEK
>gb|ABU39980_1|ARO_3004749|BCL-1__Bacillus_clausii__TM_#04
VINLLAQTE
>gb|AAC69328_1|ARO_3001265|Erm(30)__Streptomyces_venezuelae__TM_#01
MAMRDSIPRRADRDTLRRELGQNFLQDDRAVRNLVTHV
>gb|AAC69328_1|ARO_3001265|Erm(30)__Streptomyces_venezuelae__TM_#02
HVRRKFEGE
>gb|AAC69328_1|ARO_3001265|Erm(30)__Streptomyces_venezuelae__TM_#03
SLLESTNWQSAALIVQWEVARKRAGRSGGSLLTTSWAPWYEFAVHDRVRASSFRPMPR
>gb|AAC69328_1|ARO_3001265|Erm(30)__Streptomyces_venezuelae__TM_#04
PQPLLPESASRAFQNFAEAVFTGPGRGLAEILRRHIPKRTYRSLADRHGIPDGGLPKDLT
LTQWIALFQASQPSYAPGAP
>gb|BAG75524_1|ARO_3003954|efmA__Enterococcus_faecium__TM_#01
MENEQSVVLTNWKRNYL
>gb|BAG75524_1|ARO_3003954|efmA__Enterococcus_faecium__TM_#02
ALFSVVPV
>gb|BAG75524_1|ARO_3003954|efmA__Enterococcus_faecium__TM_#03
KIPKVSPEILEVPLTIFKDAKFGLQQLMDNKGLWYITINGAFVMLLFMPAISLYPLMTLD
YFGGSVGQAGAVEVVYAVGMLLGGALISFIGTWKDRMKPIIIAYIIMGLTIGASGLVPND
SQGFLYFLILNAGAGC
>gb|BAG75524_1|ARO_3003954|efmA__Enterococcus_faecium__TM_#04
LGVEKMFLFSGIGILLCGIVLFLTSA
>gb|ABS43151_1|ARO_3000784|cmeB__Campylobacter_jejuni_subsp__doylei_269_97__TM_#01
TAKMPDAVKKLGVTVRKTSSATLAAISMYSSDGSMS
>gb|AGT57825|ARO_3003762|lnuE__synthetic_construct__TM_#01
MGKNNVTEKHLFYILDLLKDLQ
>gb|AGT57825|ARO_3003762|lnuE__synthetic_construct__TM_#02
LVKKLKEIGYITV
>gb|AGT57825|ARO_3003762|lnuE__synthetic_construct__TM_#03
KKDGTATQADPKGGFYLFEKDWFTT
>gb|AGT57825|ARO_3003762|lnuE__synthetic_construct__TM_#04
KDQFDIKNLNSINQVKKEGHFSNDF
>gb|AAA26412_1|ARO_3002642|APH(3_)-Ib__Plasmid_RP4__TM_#01
MNDIDREEPCAAAAVPESMAAHVMGYKWARDKVGQSGCAVYRLHSKSGGSDLFLKHGKDA
F
>gb|AAA26412_1|ARO_3002642|APH(3_)-Ib__Plasmid_RP4__TM_#02
ISVPSVVSFVRTPNQAWLLTTAIHGKTAYQV
>gb|AAA26412_1|ARO_3002642|APH(3_)-Ib__Plasmid_RP4__TM_#03
VQQWTTHAGLPERGSIEAGVVDVDDFDK
>gb|ACT66697_1|ARO_3000846|SIM-1_beta-lactamase__Acinetobacter_bereziniae__TM_#01
MRTLLILCLFGTLNTAFAEEAQPDLKIEKIEEGIYLHTSFQEYKGFGIVKKQGLVVLDNH
KAYLIDTPASAGDTEKLVNWLEKNDFTVNG
>gb|ACT66697_1|ARO_3000846|SIM-1_beta-lactamase__Acinetobacter_bereziniae__TM_#02
KLTNELLNKNGKTQAKHSFDKE
>gb|ABV89601_1|ARO_3004826|LAP-2__Enterobacter_cloacae__TM_#01
MKKIRLIIISLLAGMCTPALSTPVNVTDTIQSTEDHIKGRVGFTEIDFLSGKVLSSHRR
>gb|ABV89601_1|ARO_3004826|LAP-2__Enterobacter_cloacae__TM_#02
KGLEQLERRITYNKH
>gb|ABV89601_1|ARO_3004826|LAP-2__Enterobacter_cloacae__TM_#03
HFLRSTGDSYTRLDRHEPS
>gb|ABV89601_1|ARO_3004826|LAP-2__Enterobacter_cloacae__TM_#04
SVLTEKSRKKLISWMQEDKVGGPLF
>gb|AAB63533_1|ARO_3002556|AAC(6_)-Ii__Enterococcus_faecium__TM_#01
PVLKDQLSDLLRLTWPEEYGDSSAEEVEEMMNPERIAVAAVDQDE
>gb|AAB63533_1|ARO_3002556|AAC(6_)-Ii__Enterococcus_faecium__TM_#02
LDHGTTLSQTDLYVHTFDKVASIQNLR
>gb|AAB63533_1|ARO_3002556|AAC(6_)-Ii__Enterococcus_faecium__TM_#03
KIVGVLPNANGWDKPDIWMAKTIIPRPDSQ
>gb|CAL34518_1|ARO_3000526|cmeR__Campylobacter_jejuni_subsp__jejuni_NCTC_11168___ATCC_700819__TM_#01
MNSNRTPSQ
>gb|CAL34518_1|ARO_3000526|cmeR__Campylobacter_jejuni_subsp__jejuni_NCTC_11168___ATCC_700819__TM_#02
YSKTQEIENGTLKEILTSFGLAFIEIFNQPEAVAFGKIIYSQVYDKDRHLANWIENNQQN
FSYNILMGFFKQQNNSYMKKNAEKLAVL
>gb|AAD04032_1|ARO_3002892|otr(B)__Streptomyces_rimosus__TM_#01
MSSANPGPAGTAD
>gb|AAD04032_1|ARO_3002892|otr(B)__Streptomyces_rimosus__TM_#02
DMAESGQGAGINLDDTSLLNGIDARLMQPVTDSFAHGFHIMFLAGGV
>gb|AAD04032_1|ARO_3002892|otr(B)__Streptomyces_rimosus__TM_#03
ERPAESGAGAKNGPLPASDA
>gb|BAA07390_1|ARO_3002893|tcr3__Kitasatospora_aureofaciens__TM_#01
MGMANATSQTGEAVADEAGGPAGFTHRQIITAL
>gb|BAA07390_1|ARO_3002893|tcr3__Kitasatospora_aureofaciens__TM_#02
QTVQAWVITGYLVSSTI
>gb|BAA07390_1|ARO_3002893|tcr3__Kitasatospora_aureofaciens__TM_#03
IVGSAACAMANSMETLAIARVLQGFGGAGLMSLPTAVIADLAPVRERGRYFSYLMMAWVA
ASVLGPLVGGLFAGAGEIL
>gb|BAA07390_1|ARO_3002893|tcr3__Kitasatospora_aureofaciens__TM_#04
SVRKALNLPHRRVDHPIDFRGALTLALC
>gb|BAA07390_1|ARO_3002893|tcr3__Kitasatospora_aureofaciens__TM_#05
TLFAVSLIGLVLFVLAERARGLEAMVPLRLFRRGGITMAT
>gb|BAA07390_1|ARO_3002893|tcr3__Kitasatospora_aureofaciens__TM_#06
VAGLVIIPVMTGAIVSQTICAKIIKKWNRYKKPAIVGLGSMAGALLSLSAAGADTPLAVI
VVIAAWLGF
>gb|BAA07390_1|ARO_3002893|tcr3__Kitasatospora_aureofaciens__TM_#07
LDGADPDEAVRRALSDPG
>gb|AAG05916_1|ARO_3004073|MuxA__Pseudomonas_aeruginosa_PAO1__TM_#01
PASAPSSDGRPGRGGKPGAALPKANALTVGVARVEQGDLALHFNALGTVTAFN
>gb|AAG05916_1|ARO_3004073|MuxA__Pseudomonas_aeruginosa_PAO1__TM_#02
VRQLQGTIRTNQGQVD
>gb|AAG05916_1|ARO_3004073|MuxA__Pseudomonas_aeruginosa_PAO1__TM_#03

VEQMNGPGKLTVTALDRNQDKVLAE
>gb|AAG05916_1|ARO_3004073|MuxA__Pseudomonas_aeruginosa_PAO1__TM_#04
VVGADNKVSQRSVAIGTSENERVVVE
>gb|AAG05916_1|ARO_3004073|MuxA__Pseudomonas_aeruginosa_PAO1__TM_#05
EASPQVLEGEPQKPQTGRPSGLQGDSVGSGSAE
>gb|AMR06225_1|ARO_3003767|mphM__Bacillus_thuringiensis__TM_#01
EKVKAKFDVGK
>gb|AMR06225_1|ARO_3003767|mphM__Bacillus_thuringiensis__TM_#02
ETLQVNDR
>gb|AAA26699_1|ARO_3002649|APH(3_)-Va__Streptomyces_fradiae__TM_#01
MDDSTLRRKYPHHEWHA
>gb|AAA26699_1|ARO_3002649|APH(3_)-Va__Streptomyces_fradiae__TM_#02
DLSGEADRLEWLHR
>gb|ACJ41547_2|ARO_3004574|Acinetobacter_baumannii_AbaQ__Acinetobacter_baumannii_AB0057__TM_#01
MDFEKDVIRTVTFKLIPALVILYL
>gb|ACJ41547_2|ARO_3004574|Acinetobacter_baumannii_AbaQ__Acinetobacter_baumannii_AB0057__TM_#02
AVGLPAVLLALPTFLWLPDNIDKVKWLSI
>gb|ACJ41547_2|ARO_3004574|Acinetobacter_baumannii_AbaQ__Acinetobacter_baumannii_AB0057__TM_#03
IQIGFLSSIPYIFGIIGLLIIPRSTDRLNDRYGHLSFLYALGACAMFLSGWLNSPVMQLA
ALAVVAFCLFSS
>gb|ACJ41547_2|ARO_3004574|Acinetobacter_baumannii_AbaQ__Acinetobacter_baumannii_AB0057__TM_#04
LLKEYTGNMAAGLYFLSIVMLFGLILTYIVYAKLERQKTQTVNIQKPL
>gb|CBJ02047_1|ARO_3004611|Escherichia_coli_ampC1_beta-lactamase__Escherichia_coli_ETEC_H10407__TM_#01
RIAKYIPGFADSPNDT
>gb|CBJ02047_1|ARO_3004611|Escherichia_coli_ampC1_beta-lactamase__Escherichia_coli_ETEC_H10407__TM_#02
MTIVMLSNKPHSPVADPQKNPNMFESGQ
>gb|AGD91915_1|ARO_3001778|OXA-232__Escherichia_coli__TM_#01
IGMPAVAKEWQENKSWNAHF
>gb|AGD91915_1|ARO_3001778|OXA-232__Escherichia_coli__TM_#02
VLKQEKIIP
>gb|BAF34030_1|ARO_3005036|BLMT__Escherichia_coli__TM_#01
ERLGFGIVFRDA
>gb|BAF34030_1|ARO_3005036|BLMT__Escherichia_coli__TM_#02
RQCKSVGIQETSSGYPRIHAPELQEWGGT
>gb|BAB36671_1|ARO_3000832|evgA__Escherichia_coli_O157_H7_str__Sakai__TM_#01
IKNDIEILAELTEGGSAVQR
>gb|BAB36671_1|ARO_3000832|evgA__Escherichia_coli_O157_H7_str__Sakai__TM_#02
VNGIQVLETLRKRQYSGIIIIVSAKNDHFYGKHCADA
>gb|BAB36671_1|ARO_3000832|evgA__Escherichia_coli_O157_H7_str__Sakai__TM_#03
SLTSDQQKLDSLSKQEISVMRYILDGKDN
>gb|AEP40503_1|ARO_3002929|vanRN__Enterococcus_faecium__TM_#01
MDTIVIVDDEK
>gb|AEP40503_1|ARO_3002929|vanRN__Enterococcus_faecium__TM_#02
KVMTFYSGKEALDYIDQNGA
>gb|AEP40503_1|ARO_3002929|vanRN__Enterococcus_faecium__TM_#03
QRYDQPSHSQSDEEFTKEGLV
>gb|CAA76544_1|ARO_3003836|qacH__Staphylococcus_saprophyticus__TM_#01
MPYLYLLLSIVS
>gb|CAA76544_1|ARO_3003836|qacH__Staphylococcus_saprophyticus__TM_#02
LYPTITTIISFLICFYFLSKTMQH
>gb|CAA76544_1|ARO_3003836|qacH__Staphylococcus_saprophyticus__TM_#03
LTTIVSVLIFKEQINLISIISIILIIFGVVLLNTFGSSH
>gb|CBL58181_1|ARO_3003918|apmA__Staphylococcus_aureus__TM_#01
VPTRRLLRALKPFKFHTADRVDPQYHYVVAVTDDDLTDFLSDEQSKSFQYANDYLTFDDE
GGELPFERMCFNVPVGRQTYFGDGVVGACENGYIKSIGQFTSINGTAEIHA
>gb|CBL58181_1|ARO_3003918|apmA__Staphylococcus_aureus__TM_#02
NFFNEESMAVFQEKLRKDPKHPYAYSKEPMTIGSDVYIGAHAFINASTVTSIGDGAIIGS
GAVVLENVPPFAVVVGVPARIKRYRFSKEMIETLLRVKWWDWSIEEINENVDALISPELF
MKKYGSL
>gb|APY23733_1|ARO_3004734|ARL-1__Staphylococcus_arlettae__TM_#01
MKKFFTIFVLLCVCFAYTTA
>gb|APY23733_1|ARO_3004734|ARL-1__Staphylococcus_arlettae__TM_#02
LTKLEHKNDATVGVYGINTATG
>gb|APY23733_1|ARO_3004734|ARL-1__Staphylococcus_arlettae__TM_#03
TYSHNADT
>gb|APY23733_1|ARO_3004734|ARL-1__Staphylococcus_arlettae__TM_#04
ITSGLLLQQ
>gb|APY23733_1|ARO_3004734|ARL-1__Staphylococcus_arlettae__TM_#05
SPEALNKTVTIKESDIVAYSPVTEQYVGKTMTLRQLISAAMLQSDNTASN
>gb|APY23733_1|ARO_3004734|ARL-1__Staphylococcus_arlettae__TM_#06
STADTSTPRATAH
>gb|APY23733_1|ARO_3004734|ARL-1__Staphylococcus_arlettae__TM_#07
LLTTDAVAPQQRKFLQNLM
>gb|APY23733_1|ARO_3004734|ARL-1__Staphylococcus_arlettae__TM_#08
SLIKKGVP
>gb|APY23733_1|ARO_3004734|ARL-1__Staphylococcus_arlettae__TM_#09
SYKVGDKSGQGTTYGTRNDVA
>gb|APY23733_1|ARO_3004734|ARL-1__Staphylococcus_arlettae__TM_#10
IYPKHQTKPI
>gb|BAD89844_2|ARO_3000753|abeM__Acinetobacter_baumannii__TM_#01
LMPFFLHV
>gb|BAD89844_2|ARO_3000753|abeM__Acinetobacter_baumannii__TM_#02
ASAYRNTSIFSRFDKISLT
>gb|BAD89844_2|ARO_3000753|abeM__Acinetobacter_baumannii__TM_#03
YQVQKIGLSTAVFFALLTMSFIALGREQIVSVYTQDIN
>gb|BAD89844_2|ARO_3000753|abeM__Acinetobacter_baumannii__TM_#04
SRLYLNTKRLSQT
>gb|BAA11237_1|ARO_3000254|emrY__Escherichia_coli__TM_#01
LSVTFFSLSSLMCSL
>gb|BAA11237_1|ARO_3000254|emrY__Escherichia_coli__TM_#02
NAIWAGLAYAPIGIMPLLI
>gb|BAA11237_1|ARO_3000254|emrY__Escherichia_coli__TM_#03
LMYAVCYYWRSVTF
>gb|BAA11237_1|ARO_3000254|emrY__Escherichia_coli__TM_#04
ATIDQFNPVFNSSSQIMDK
>gb|CAB55427_1|ARO_3004775|CME-1__Elizabethkingia_meningoseptica__TM_#01
MKKIILLFILTSQL
>gb|CAB55427_1|ARO_3004775|CME-1__Elizabethkingia_meningoseptica__TM_#02
VKTLYANYTTTASMVKTLKAFYKGMFLSKRSTIFLMDIMTKTNTGMSKLPGLLPKVRMAR
KTGSSGKMKNGLTIAENDS
>gb|CAB55427_1|ARO_3004775|CME-1__Elizabethkingia_meningoseptica__TM_#03
KDSMESEEVNCGMIAQVSKIVWDALNKKK
>gb|BAE78116_1|ARO_3003576|eptA__Escherichia_coli_str__K-12_substr__W3110__TM_#01

MLKRLLKRPSLNLLAWLLLAA
>gb|AHV80711_1|ARO_3002863|dfrA8__Salmonella_enterica_subsp__enterica_serovar_Typhimurium__TM_#01
MIELHAILAATANGCIGKDNALPWPPLKGDLARFKKL
>gb|AHV80711_1|ARO_3002863|dfrA8__Salmonella_enterica_subsp__enterica_serovar_Typhimurium__TM_#02
VKLEGRTCIVMTRQALELPGVRDANGAIFVNNVSDAMRFAQEESVGDVAYVIGGAEIFKR
LALMITQIELTFVKRLYEGDTYVDLAEMVKDYEQNGMEEHDLHTYFTYRKKELTE
>gb|AAN63647_1|ARO_3000843|MUS-1_beta-lactamase__Myroides_odoratimimus__TM_#01
MHRILSVITMLICTTLVHAQSDKLKIKQLNDNMYIYTTYQEFQGVTYSSNSMYVLTDEG
>gb|AAN63647_1|ARO_3000843|MUS-1_beta-lactamase__Myroides_odoratimimus__TM_#02
ILIDTPWDKDQYEPLLEYIRSN
>gb|AAN63647_1|ARO_3000843|MUS-1_beta-lactamase__Myroides_odoratimimus__TM_#03
IAWPKTIEAVKQKFKNAKVIIPGHDEWDM
>gb|AAN63647_1|ARO_3000843|MUS-1_beta-lactamase__Myroides_odoratimimus__TM_#04
QQHSTKND
>gb|CAE48335_2|ARO_3002599|AAC(6_)-30_AAC(6_)-Ib__fusion_protein__Pseudomonas_aeruginosa__TM_#01
TQFFNGEVEEPNEVLLAVTEENDAIAHIELSLRYDIDGLT
>gb|CAE48335_2|ARO_3002599|AAC(6_)-30_AAC(6_)-Ib__fusion_protein__Pseudomonas_aeruginosa__TM_#02
EERHRAAGVVLKLLRAAEFWARDQGCLAFASDRDDRVIIYARYTGAPPNNS
>gb|AAC61670_1|ARO_3001308|VgbB__Staphylococcus_cohnii__TM_#01
NDKTIQEYQL
>gb|AAC61670_1|ARO_3001308|VgbB__Staphylococcus_cohnii__TM_#02
KCKIGKLNLNE
>gb|CAA37605_1|ARO_3002638|APH(3__)-Ia__Streptomyces_griseus__TM_#01
MSDHPGPGAVTPELFGV
>gb|CAA37605_1|ARO_3002638|APH(3__)-Ia__Streptomyces_griseus__TM_#02
GADDLRTAWGA
>gb|AAA25550_1|ARO_3003105|dfrA3__Plasmid_pAZ1__TM_#01
MLISLIAALAHNN
>gb|AAA25550_1|ARO_3003105|dfrA3__Plasmid_pAZ1__TM_#02
QWQAEGVEVAPSLDAALALLTDCEEA
>gb|AAA25550_1|ARO_3003105|dfrA3__Plasmid_pAZ1__TM_#03
YIDAQLNGDTHFPDYLSLGWQELERSTHPADDKNSYACEFVTLSRQR
>gb|CAA39184_1|ARO_3002536|AAC(3)-IIIa__Pseudomonas_aeruginosa__TM_#01
MTDLNIPHTHAHLVDAFQALGIRAGQAL
>gb|CAA39184_1|ARO_3002536|AAC(3)-IIIa__Pseudomonas_aeruginosa__TM_#02
DSLPDALKAVYLEQH
>gb|CAA39184_1|ARO_3002536|AAC(3)-IIIa__Pseudomonas_aeruginosa__TM_#03
CVHRSANPEASMVAVGRQAALLTANHALDYGYGVESPLAKLVAIE
>gb|CAA39184_1|ARO_3002536|AAC(3)-IIIa__Pseudomonas_aeruginosa__TM_#04
KMRHKNVVRYPCPILRDGRKVWVTVEDYDTGDPHDDYSFEQIARDYVAQGGGTRGKVGDA
DAYLFAAQDLTRFAVQWLESRFGDSASYG
>gb|AYD68552_1|ARO_3005018|LMB-1__Citrobacter_freundii__TM_#01
MTLAKSFRCFCLVTTLSSMAMLAGCGGTAPVTVLSPPPADWVNSCK
>gb|AYD68552_1|ARO_3005018|LMB-1__Citrobacter_freundii__TM_#02
NAYRASLH
>gb|AYD68552_1|ARO_3005018|LMB-1__Citrobacter_freundii__TM_#03
TQGCVVYADAITQRLEQRLIKETTQ
>gb|AKQ05892_1|ARO_3004581|tet(48)__uncultured_bacterium__TM_#01
MTFLFKEFKGVFKMKKVLVVGAGVAGLAVCYWLKE
>gb|AKQ05892_1|ARO_3004581|tet(48)__uncultured_bacterium__TM_#02
NALRKGGYGVDIFGIAVDIAKKMSVYEKICAMRTQLEHGFYVNADGHTLV
>gb|AKQ05892_1|ARO_3004581|tet(48)__uncultured_bacterium__TM_#03
GEEVEILREDLIEILKQAIKDIPCHFNQRIKRIKQDGKHVEVTFKDNK
>gb|AKQ05892_1|ARO_3004581|tet(48)__uncultured_bacterium__TM_#04
FTFDKEEYDLIDFGCYSAIFSLPNYLKLRQSEIAFDANQKFISVSSDKNPTIALASLMFH
SNRGDNIRNEKDQKSFFKDAFIDLGWETNNLLQYMEESNDFYFDVAT
>gb|AKQ05892_1|ARO_3004581|tet(48)__uncultured_bacterium__TM_#05
ITAFERYNM
>gb|AKQ05892_1|ARO_3004581|tet(48)__uncultured_bacterium__TM_#06
LGAWINETFLLEDAVSKEAVEARTDNIIKKISAISNVIKLPEYSAYK
>gb|AHY03238_1|ARO_3002789|QnrD2__Salmonella_enterica_subsp__enterica_serovar_Hadar__TM_#01
MEKHFINEKFSRDQFTGNR
>gb|AHY03238_1|ARO_3002789|QnrD2__Salmonella_enterica_subsp__enterica_serovar_Hadar__TM_#02
VITGAVFRGSDLSCGEFSSFDWSLADFTGCDLTGGALGELDARR
>gb|AHY03238_1|ARO_3002789|QnrD2__Salmonella_enterica_subsp__enterica_serovar_Hadar__TM_#03
NLDGVKLDGEQA
>gb|AHY03238_1|ARO_3002789|QnrD2__Salmonella_enterica_subsp__enterica_serovar_Hadar__TM_#04
QLVESLGVIVHR
>gb|AAG08375_1|ARO_3004038|Pseudomonas_aeruginosa_emrE__Pseudomonas_aeruginosa_PAO1__TM_#01
AEVVATTSLKAVAGFSKPLPLLLVVGGYVLAFSMLVLVMRT
>gb|AAG08375_1|ARO_3004038|Pseudomonas_aeruginosa_emrE__Pseudomonas_aeruginosa_PAO1__TM_#02
MFVYGQRLDPAALL
>gb|ACE77058_1|ARO_3002366|PER-4__Proteus_vulgaris__TM_#01
SAQSPLLK
>gb|ACE77058_1|ARO_3002366|PER-4__Proteus_vulgaris__TM_#02
GPDDLEPLL
>gb|ACE77058_1|ARO_3002366|PER-4__Proteus_vulgaris__TM_#03
NPFEKFPMQSVFK
>gb|ACE77058_1|ARO_3002366|PER-4__Proteus_vulgaris__TM_#04
HLAMLVLHQVDQGKLDLN
>gb|ACE77058_1|ARO_3002366|PER-4__Proteus_vulgaris__TM_#05
VQQLLQYSVS
>gb|ACE77058_1|ARO_3002366|PER-4__Proteus_vulgaris__TM_#06
VVANEAQMHADDQVQY
>gb|ACE77058_1|ARO_3002366|PER-4__Proteus_vulgaris__TM_#07
AGKTAATND
>gb|ACE77058_1|ARO_3002366|PER-4__Proteus_vulgaris__TM_#08
LPDGRPLLVAVFVKDSAES
>gb|ACE77058_1|ARO_3002366|PER-4__Proteus_vulgaris__TM_#09
RTNEAIIAQVAQ
>gb|ACE77058_1|ARO_3002366|PER-4__Proteus_vulgaris__TM_#10
AYQFELKKLSA
>gb|AAF86219_1|ARO_3000375|ErmB__Enterococcus_faecium__Partial_TM_#01
EKVLNQIIKQLNLKETDTVYEIGTGKGHLTTKLAKISKQVTSIELDSHLFNLSSEKLKLN
TRV
>gb|AAF86219_1|ARO_3000375|ErmB__Enterococcus_faecium__Partial_TM_#02
TTDVPDKYWKLYT
>gb|EHS19134_1|ARO_3004661|Staphylococcus_aureus_FosB__Staphylococcus_aureus_subsp__aureus_IS-88__TM_#01
MLKSINHICFSVRNLNDSIHFYRDILLGKLLLTGK
>gb|EHS19134_1|ARO_3004661|Staphylococcus_aureus_FosB__Staphylococcus_aureus_subsp__aureus_IS-88__TM_#02
SEFKYWHQRLKDNNVNILE
>gb|ALU64000|ARO_3003713|VCC-1__Vibrio_cholerae__TM_#01
MKRIAMYVALSISTSTAFADEHNKNMADIEAAFEGRVGVYAINTGSGKAY

>gb|ALU64000|ARO_3003713|VCC-1__Vibrio_cholerae__TM_#02
MDQDSPGVLLEKVNYH
>gb|ALU64000|ARO_3003713|VCC-1__Vibrio_cholerae__TM_#03
TEKFQSQGMAVG
>gb|ALU64000|ARO_3003713|VCC-1__Vibrio_cholerae__TM_#04
EKYIKGPEGMTQFMNSIGDTK
>gb|ALU64000|ARO_3003713|VCC-1__Vibrio_cholerae__TM_#05
ISNTVLDNYHQEIFKK
>gb|ALU64000|ARO_3003713|VCC-1__Vibrio_cholerae__TM_#06
KYGTANDHAFILQGNNA
>gb|ALU64000|ARO_3003713|VCC-1__Vibrio_cholerae__TM_#07
GEHMKHDDEVIAKAARIAIENVK
>gb|AAA73865_1|ARO_3002674|Clostridium_butyricum_catB__Clostridium_butyricum__TM_#01
YEIKLKNIKF
>gb|AAA73865_1|ARO_3002674|Clostridium_butyricum_catB__Clostridium_butyricum__TM_#02
KSFLRFYSDYLDDIKNYGNIMKFTPKSNEPD
>gb|AAA73865_1|ARO_3002674|Clostridium_butyricum_catB__Clostridium_butyricum__TM_#03
RFINEMQELAFSFQEWLENK
>gb|AAR32134_1|ARO_3001792|OXA-62__Pandoraea_pnomenusa__TM_#01
MNTIISRRWRAGLWRRLVGAVVLPATLAATPAAYAADVPKAA
>gb|AAR32134_1|ARO_3001792|OXA-62__Pandoraea_pnomenusa__TM_#02
GRITERADWGKL
>gb|AAR32134_1|ARO_3001792|OXA-62__Pandoraea_pnomenusa__TM_#03
VLKDLKLI
>gb|AFO59566_1|ARO_3002226|IMP-35__Pseudomonas_aeruginosa__TM_#01
DVYVHTSFE
>gb|AAB05625_1|ARO_3002964|vanWB__Enterococcus_faecalis__TM_#01
CYAQTIGEKTLPY
>gb|AAB05625_1|ARO_3002964|vanWB__Enterococcus_faecalis__TM_#02
WVTLDDEKIIGQVF
>gb|AAB60941_1|ARO_3005099|23S_rRNA_(adenine(2058)-N(6))-methyltransferase_Erm(A)__Streptococcus_pyogenes__TM_#01
KKAVEPFQNIKVIHEDILKFS
>gb|AAB60941_1|ARO_3005099|23S_rRNA_(adenine(2058)-N(6))-methyltransferase_Erm(A)__Streptococcus_pyogenes__TM_#02
KPFILKKDYKKYRF
>gb|AAB60941_1|ARO_3005099|23S_rRNA_(adenine(2058)-N(6))-methyltransferase_Erm(A)__Streptococcus_pyogenes__TM_#03
LRQVLKHANVTDLDKLSN
>gb|AAR96051_1|ARO_3002894|otrC__Streptomyces_rimosus__TM_#01
MTRKTISN
>gb|AAR96051_1|ARO_3002894|otrC__Streptomyces_rimosus__TM_#02
RGALPAHV
>gb|AAR96051_1|ARO_3002894|otrC__Streptomyces_rimosus__TM_#03
GPDAGRRPWRF
>gb|AAR96051_1|ARO_3002894|otrC__Streptomyces_rimosus__TM_#04
TWCANRRALRRTIG
>gb|AAR96051_1|ARO_3002894|otrC__Streptomyces_rimosus__TM_#05
TISGHQHPSAQ
>gb|AAR96051_1|ARO_3002894|otrC__Streptomyces_rimosus__TM_#06
ADGGPQDGPQDQQGVQDKQYEEVPA
>gb|BAF80809_1|ARO_3002665|npmA__Escherichia_coli__TM_#01
MLILKGTKTVDLSKDELTEIIGQFDRVHIDLGTGDGRNIYKLAINDQNTFYIGIDPVKEN
LFDISKKIIKKPSKGGLSNVVFVIAAAESLPFELKNIADSISILFPWGTLLEYVIKPNRD
ILSNVADLAKKEAHFEFVTTYSDSYEEAEIKKRGLPLLSKAYFLSEQYKAELSNSGFRID
DVKELDNEYVKQFNSLWAKRLAFGRKRSFFRVSGHVSKH
>gb|ACI28880_1|ARO_3002575|AAC(6_)-Iai__Pseudomonas_aeruginosa__TM_#01
MKYTIIDIKDSET
>gb|ACI28880_1|ARO_3002575|AAC(6_)-Iai__Pseudomonas_aeruginosa__TM_#02
EISPESWPTLQKAKEDVIECIEG
>gb|ACI28880_1|ARO_3002575|AAC(6_)-Iai__Pseudomonas_aeruginosa__TM_#03
THHNMGFGKILINEIEKKARERNLE
>gb|ACI28880_1|ARO_3002575|AAC(6_)-Iai__Pseudomonas_aeruginosa__TM_#04
IELNNENILQEIKNIRNLE
>gb|ADI55014_1|ARO_3002801|QnrVC4__Aeromonas_caviae__TM_#01
MDKTDQLYVQADFSHQD
>gb|CAE00499_1|ARO_3003835|cdeA__Clostridioides_difficile__TM_#01
MENLFTRKFTTF
>gb|CAE00499_1|ARO_3003835|cdeA__Clostridioides_difficile__TM_#02
NRQDEANSTFSFIVLFSLIVGILFTVISYFFIKEISILLGATDKLLPYCITYGKVMILCT
PFYILKFIFEYFARTDGNSK
>gb|CAE00499_1|ARO_3003835|cdeA__Clostridioides_difficile__TM_#03
YFGMGLLGAAVATAIGIILTCVLGIIYFLSNKSTLKLRKPKTDFRLIRD
>gb|CAE00499_1|ARO_3003835|cdeA__Clostridioides_difficile__TM_#04
VVALKLAGENGLAALTIVLYAHFLMTSVYLGFAAGVSPLISYNFGAENSDKLKETFKHSL
KFIFISSLLVFIIALVFAPFIVRVFVNPDNTVFKLALQ
>gb|CAE00499_1|ARO_3003835|cdeA__Clostridioides_difficile__TM_#05
NMTGLWLTVPFAEVITIFISILFIKKYKGRYKY
>gb|BAD10948_2|ARO_3002563|AAC(6_)-Isa__Streptomyces_albulus__TM_#01
MELRGDDVVLRPVA
>gb|BAD10948_2|ARO_3002563|AAC(6_)-Isa__Streptomyces_albulus__TM_#02
AGMLAIVFEGEVVGAIQFY
>gb|ABG49324_1|ARO_3002609|aadA9__Corynebacterium_sp__L2-79-05__TM_#01
SNSIHTGISRQLSQARDVIKRHLAST
>gb|ABG49324_1|ARO_3002609|aadA9__Corynebacterium_sp__L2-79-05__TM_#02
ATRRSLMLDFLNISAPPCESSIL
>gb|ABG49324_1|ARO_3002609|aadA9__Corynebacterium_sp__L2-79-05__TM_#03
VAQKVFMPVPEHDFLQVLSDTLKLWNTHEDWEN
>gb|ABG49324_1|ARO_3002609|aadA9__Corynebacterium_sp__L2-79-05__TM_#04
TETGGIVPKDVAAEWVLERLPAEHKPILVEARQAYLGLCKDSLALRADETSAFIGYAKSA
VADLLEKRKSQTSHICDGAKNV
>gb|ANZ79240_1|ARO_3004035|tetA(60)__uncultured_bacterium__TM_#01
MNDLLKVIINFIKKHPMRYLVSFILMIGSSIAAVYPARIIGQVVDKIVASELNAEWLGTQ
LVILVGIILVAYITESIWTYFIFIGYYEIQKELRVKLLRNNLRKKIPFYAHFRTGEIITR
SSEDVTTIGDMMGFGMFALMNSTLLMSVSIYMMVTTISLPLTIAAILPLPILSYLVYKWG
FDLEEEYNKAQNAVSQLNN
>gb|ANZ79240_1|ARO_3004035|tetA(60)__uncultured_bacterium__TM_#02
AMMDEFRAKTKKAMKQNIIVTEIESRFIPLAQLFMMISFTIALFYGGYLVSTGAILVGDV
IAFQVYMGAIMWPMFMIGDIITNYKRGKVATERINEVLKHD
>gb|EGP45232_2|ARO_3004143|AxyX__Achromobacter_insuavis_AXX-A__TM_#01
MTHRVPFRTLAFASVLVLVSACSKQEAPEA
>gb|EGP45232_2|ARO_3004143|AxyX__Achromobacter_insuavis_AXX-A__TM_#02
DLVSDRAISERDHAESVAQERQARAEVALAKANLQSARLRLE
>gb|EGP45232_2|ARO_3004143|AxyX__Achromobacter_insuavis_AXX-A__TM_#03
MQLQKQIRAGALQGVAPDKM
>gb|EGP45232_2|ARO_3004143|AxyX__Achromobacter_insuavis_AXX-A__TM_#04

NADGAHVLVAGDDGELRSVAVTAHRLLGPNWVVTEGLAGGERVVVENAAQLAPGQK
>gb|EHL92831_1|ARO_3004588|Klebsiella_pneumoniae_KpnG__Klebsiella_sp__4_1_44FAA__TM_#01
EVNTTDRDGEMLASQVRSSPV
>gb|EHL92831_1|ARO_3004588|Klebsiella_pneumoniae_KpnG__Klebsiella_sp__4_1_44FAA__TM_#02
EIIQANAG
>gb|AAA26685_1|ARO_3002542|AAC(3)-VIIIa__Streptomyces_fradiae__TM_#01
RGNGRVPEALRHQ
>gb|AAA26685_1|ARO_3002542|AAC(3)-VIIIa__Streptomyces_fradiae__TM_#02
SRGVPYGRVVPEGVVPFT
>gb|AND91341_1|ARO_3000856|BJP-1_beta-lactamase__Bradyrhizobium_diazoefficiens_USDA_110__TM_#01
MRRLTAALCALTLLSTGAQAQTIKDFLAVA
>gb|AND91341_1|ARO_3000856|BJP-1_beta-lactamase__Bradyrhizobium_diazoefficiens_USDA_110__TM_#02
AKRAEMKDGAPNPFIKPGELV
>gb|AHF82024_1|ARO_3004648|AQU-3__Aeromonas_dhakensis__TM_#01
MKQTSPLS
>gb|AHF82024_1|ARO_3004648|AQU-3__Aeromonas_dhakensis__TM_#02
LALSALLLSPLTQAAPADPLVGVVD
>gb|AHF82024_1|ARO_3004648|AQU-3__Aeromonas_dhakensis__TM_#03
VIRPLVKEHRIPGMAVAV
>gb|AHF82024_1|ARO_3004648|AQU-3__Aeromonas_dhakensis__TM_#04
ANINGVDDKGLQQAIALTHQ
>gb|CAA40897_1|ARO_3004548|dfrA9__Escherichia_coli__TM_#01
MASLNMIVAVNKTGGIGFENQIPWHEPEDLKHFKAVTMNS
>gb|CAA40897_1|ARO_3004548|dfrA9__Escherichia_coli__TM_#02
LHVVVSKTVPPTQNTDQVVYVSTYQIAVRTASLLVDKPEYSQIFVIGGKSAYENLAAYVD
KLYLTRVQLNTQQDTELDLSLFKSWKLVSEVPTITENKTKLIFQIWINPNPISEEPTC
>gb|AAA26775_1|ARO_3003564|EXO_beta-lactamase__Streptomyces_albus__TM_#01
VSDAERRLAGLERASGARLGVYAYDTGS
>gb|AAA26775_1|ARO_3003564|EXO_beta-lactamase__Streptomyces_albus__TM_#02
LYTQDDVEQADGAGPETGKPQNLANAQLTVEELCEVSITA
>gb|WP_000949574_1|ARO_3005348|DfrA36__Gammaproteobacteria__TM_#01
MLSKSDILLQFIYFYNTFIFLIAFF
>gb|WP_000949574_1|ARO_3005348|DfrA36__Gammaproteobacteria__TM_#02
AILTRNTAFEAPNCTVFHSMEGCLKHYENEDKRT
>gb|WP_000949574_1|ARO_3005348|DfrA36__Gammaproteobacteria__TM_#03
NRVDEMFITFVDHTFGADTFFPSIDFSLWNEEVLRVHEADSKNAYNFTVKKFTKKLS
>gb|ABX54689_1|ARO_3002973|vanTmL__Enterococcus_faecalis__TM_#01
MKKQNTGVNNFRLIAAAMVVAIHCFPFQTISKELDTLVTLTVFRIAVPFFFMVSGYYLLG
PIPSSATNTYQINNYIKKQLKVYTFAIVLYLPLAFYSQSITLDMSIISFIKQLLFNGFFY
HLWFFPAWVLGLLIVQFLLKRMNIQTVLFITFVA
>gb|ABX54689_1|ARO_3002973|vanTmL__Enterococcus_faecalis__TM_#02
WGIVKQVPFFFRFYNAIFQLFGYTRNGLFYAPLFFALGAYLYKMNIKNFNSARNNYLLLL
FSIEMILESYFLHL
>gb|ABX54689_1|ARO_3002973|vanTmL__Enterococcus_faecalis__TM_#03
FVMTLVFIKIYNWSPKNNLLNSSQLSLGVYLIHPYIIAVIHSISIYVSIFTNSIINYLSV
LLISYLTIRLIL
>gb|ACL82960_1|ARO_3002947|vanHM__Enterococcus_faecium__TM_#01
SEQDEADVFQEISSRFGVTPTIVSSPISETNVMLAPKNK
>gb|ACL82960_1|ARO_3002947|vanHM__Enterococcus_faecium__TM_#02
GHVLAYGNNKEATANYVSF
>gb|ABA71733_1|ARO_3002972|vanTG__Enterococcus_faecalis__TM_#01
RKAFLIAS
>gb|ABA71733_1|ARO_3002972|vanTG__Enterococcus_faecalis__TM_#02
KSVSCLNV
>gb|ABA71733_1|ARO_3002972|vanTG__Enterococcus_faecalis__TM_#03
QNRLSLKRSIVGFIVCFALMFGEA
>gb|ABA71733_1|ARO_3002972|vanTG__Enterococcus_faecalis__TM_#04
IIYIIHPFMIVVIRLFAK
>gb|ABA71733_1|ARO_3002972|vanTG__Enterococcus_faecalis__TM_#05
QIGSFYKVLDWLKSS
>gb|ABA71733_1|ARO_3002972|vanTG__Enterococcus_faecalis__TM_#06
VGVALYGVLSSTN
>gb|AKA86814|ARO_3003746|optrA__Enterococcus_faecalis__TM_#01
LTYQKEYETMIRSMGFTEADYKKP
>gb|AKA86814|ARO_3003746|optrA__Enterococcus_faecalis__TM_#02
LQQIEIERITRLIERFRYKPTKAKMVQSKIKLLQRMQILNAPDQYDTKTYMSKFQPRISS
SRQVLSASELVIGYDTPLAKVNFNLERGQKLG
>gb|AKA86814|ARO_3003746|optrA__Enterococcus_faecalis__TM_#03
GVAALSGDFKFGYNVEISYFDQQLAQISGDDTLFEIFQSEYPELNDTEVRTALGSFQ
>gb|AKA86814|ARO_3003746|optrA__Enterococcus_faecalis__TM_#04
KVRLTLCKLLYKR
>gb|AKA86814|ARO_3003746|optrA__Enterococcus_faecalis__TM_#05
FDKDGVEFVQSTYGEYEKKRMNSEKPFNNIKVEQKVEKNNTVKGDRNSIEKEKVKKEKRI
EKLEVLINQYDEELERLNKIISEPNNSSDYIVLTEIQKSIDDVKRCQGNYFNEW
>gb|AAB19430_2|ARO_3002242|CARB-3__Pseudomonas_aeruginosa__TM_#01
GDLRDTTTP
>gb|AAB19430_2|ARO_3002242|CARB-3__Pseudomonas_aeruginosa__TM_#02
RNDAIVKIG
>gb|CAQ53840_1|ARO_3000853|AIM-1__Pseudomonas_aeruginosa__TM_#01
MKRRFTLLGSVVALALSSTALASDAPASRGCADDAGWNDPAM
>gb|CAQ53840_1|ARO_3000853|AIM-1__Pseudomonas_aeruginosa__TM_#02
LPDRTDPQFEVAEPVAPVANIVTLADDGVVSVGPLALT
>gb|CAQ53840_1|ARO_3000853|AIM-1__Pseudomonas_aeruginosa__TM_#03
AAPLMDTTACRRYAQGARQRLEKRLAEEAATSPSSGARP
>gb|BAE85481_1|ARO_3003727|vanKI__Desulfitobacterium_hafniense_Y51__TM_#01
MLNVDKISE
>gb|BAE85481_1|ARO_3003727|vanKI__Desulfitobacterium_hafniense_Y51__TM_#02
LVAYLRSKQVFSIKIDPPVVQAKWSAPTIKTFLGQAREQGSKGKVLRDLPPDEDYTTVQQ
VQQQLRQMGWRKQRGDTGFAATQPQYVYRLPLE
>gb|BAE85481_1|ARO_3003727|vanKI__Desulfitobacterium_hafniense_Y51__TM_#03
RLGVKVRVGTEQDLPAFYELLKVTSERDHFKVRSFSYFSNLYRSLKAEAADRIALYLAED
EEELLAATLAVHSNA
>gb|BAE85481_1|ARO_3003727|vanKI__Desulfitobacterium_hafniense_Y51__TM_#04
YRAFQLYLKRR
>gb|AIA08936_1|ARO_3000444|rphA__Streptomyces_sp__WAC4747__TM_#01
EPTPAETDGGAPPAGDGAEPDEADPSIVTELIERSRRSLAELEREIG
>gb|AIA08936_1|ARO_3000444|rphA__Streptomyces_sp__WAC4747__TM_#02
YLTFDEFEEAVRV
>gb|ACM47284|ARO_3002934|vanSD__Enterococcus_faecium__TM_#01
ACSILIIAGVYLFILKDNFANVVVAILDSFIYHDRDEAVVV
>gb|ACM47284|ARO_3002934|vanSD__Enterococcus_faecium__TM_#02
VMGVFFMIFRRYLDSISKYFKEINRGIDTLVNEDAN
>gb|ACM47284|ARO_3002934|vanSD__Enterococcus_faecium__TM_#03

TKRKTDAE
>gb|ACM47284|ARO_3002934|vanSD__Enterococcus_faecium__TM_#04
LIFQISRLCTAKSI
>gb|CAC14595_1|ARO_3003056|smeE__Stenotrophomonas_maltophilia__TM_#01
DAALEDMKG
>gb|CAC14595_1|ARO_3003056|smeE__Stenotrophomonas_maltophilia__TM_#02
HRPWRFMGIVAALF
>gb|AIG22447_1|ARO_3002191|MOX-9__Citrobacter_freundii__TM_#01
MQQQVRMTLLMASTLLWAGLAQATADTQA
>gb|AIG22447_1|ARO_3002191|MOX-9__Citrobacter_freundii__TM_#02
TGTGDEAMQQAIALTHKGV
>gb|AIG22447_1|ARO_3002191|MOX-9__Citrobacter_freundii__TM_#03
ETAHAILSKLAE
>gb|AAO43110_1|ARO_3000300|lsaA__Enterococcus_faecalis_ATCC_29212__TM_#01
MSKIELKQLSFAYDNQEALLFDQANI
>gb|AAO43110_1|ARO_3000300|lsaA__Enterococcus_faecalis_ATCC_29212__TM_#02
LHQVDFVYFPQTVAEEQQLTYYVLQEVTSFEQWKLERELT
>gb|AAO43110_1|ARO_3000300|lsaA__Enterococcus_faecalis_ATCC_29212__TM_#03
TLYQGDFSIYEEQKKLRDAFELAENEKIKKEVNRLKETARKKAEWSMNREGDKYGNAK
>gb|AAO43110_1|ARO_3000300|lsaA__Enterococcus_faecalis_ATCC_29212__TM_#04
HIQQRAETQLAEKEKLLKDLEYIDPLSMDYQPTHHKTLLTVEELRLGYEKNWLFTPISFS
INAGEIVGITGKNGSGKSSLIQYLLGDFSGDSEGEATLAH
>gb|AAO43110_1|ARO_3000300|lsaA__Enterococcus_faecalis_ATCC_29212__TM_#05
LILSVRPAMLVIEHDAHFMKKITDKKIALKS
>gb|AAP93922_1|ARO_3000569|tet(41)__Serratia_marcescens__TM_#01
MKKPMLVILL
>gb|AAP93922_1|ARO_3000569|tet(41)__Serratia_marcescens__TM_#02
LIGLAADAVGLALLSVATRGWAPFALLPFFAAGGMALPALQALMAHKVDDD
>gb|AAP93922_1|ARO_3000569|tet(41)__Serratia_marcescens__TM_#03
LAAALYLVVPPLLARSRARDAAP
>gb|EGP45230|ARO_3004142|OprZ__Achromobacter_insuavis_AXX-A__TM_#01
YFNQRSLAEQLRLTDD
>gb|EGP45230|ARO_3004142|OprZ__Achromobacter_insuavis_AXX-A__TM_#02
ELTREASLARHALGLLAGDFALPLGVDP
>gb|EGP45230|ARO_3004142|OprZ__Achromobacter_insuavis_AXX-A__TM_#03
DRFSDLFSGGTGGWSFA
>gb|AAL59753_1|ARO_3000412|sul2__Vibrio_cholerae__TM_#01
SLDSYQPATQ
>gb|AAL59753_1|ARO_3000412|sul2__Vibrio_cholerae__TM_#02
THEPRPLR
>gb|CAP07796_1|ARO_3000860|rmtB__Escherichia_coli__TM_#01
MNINDALTSILASKKYRALCPDTVR
>gb|CAP07796_1|ARO_3000860|rmtB__Escherichia_coli__TM_#02
TLYDFIFSAETPR
>gb|CAP07796_1|ARO_3000860|rmtB__Escherichia_coli__TM_#03
PAEAGDLALIFKLLPLLEREQAGSAMALLQSLN
>gb|CAP07796_1|ARO_3000860|rmtB__Escherichia_coli__TM_#04
AEFEIEDKKTIGTELIYLIKKNG
>gb|ACK75961_1|ARO_3002787|QnrC__Proteus_mirabilis__TM_#01
MNYSHKTYDQIDFSGQDLSSHHFSHCKFFGCNFNRVNLRDAKFMGCTFIESNDFEGCNFI
YADLRDASFMNCMLSMANFQ
>gb|ACK75961_1|ARO_3002787|QnrC__Proteus_mirabilis__TM_#02
SFSDDFWEQCRIQGCDLTHSELN
>gb|BAD14386_1|ARO_3002573|AAC(6_)-Iae__Pseudomonas_aeruginosa__TM_#01
MKYNIVNIKDSEK
>gb|BAD14386_1|ARO_3002573|AAC(6_)-Iae__Pseudomonas_aeruginosa__TM_#02
HINFDSWPSLQKATETVIECISA
>gb|BAD14386_1|ARO_3002573|AAC(6_)-Iae__Pseudomonas_aeruginosa__TM_#03
KHQNKGFGKILIFETEKKAKERNLE
>gb|BAD14386_1|ARO_3002573|AAC(6_)-Iae__Pseudomonas_aeruginosa__TM_#04
SELNNENIFHE
>gb|CCC86795_1|ARO_3001209|mecC__Staphylococcus_aureus_subsp__aureus_LGA251__TM_#01
MKKIYISVLVLLLIMIIITWLFKDDDIEKTISSIEKGNYNEVYKNSSEKSKLAYGEEEIV
DRNKKIYKDLSVNNLKITNHEI
>gb|CCC86795_1|ARO_3001209|mecC__Staphylococcus_aureus_subsp__aureus_LGA251__TM_#02
RPDVIVPGLKNG
>gb|CCC86795_1|ARO_3001209|mecC__Staphylococcus_aureus_subsp__aureus_LGA251__TM_#03
KEKYDDIARDLQIDTK
>gb|CCC86795_1|ARO_3001209|mecC__Staphylococcus_aureus_subsp__aureus_LGA251__TM_#04
KTKSQIWKKDIIPKKDID
>gb|ABW06859_1|ARO_3000603|Erm(41)__Mycobacteroides_abscessus__TM_#01
HLRSRFAEEDVRVAEADLLAF
>gb|ABW06859_1|ARO_3000603|Erm(41)__Mycobacteroides_abscessus__TM_#02
VTSALIRSLLTPESRLLAADLVLQRGAVHKHAKRAPVRHWTLRAGITLPRSAFHH
>gb|AAT90846_1|ARO_3003556|SLB-1__Shewanella_livingstonensis__TM_#01
MLSLPSYSHEVEPTSTTIQSVTSSLEGQLSISKLADGVYLHHSYKNVSNFGLVEANGLVV
IK
>gb|AAT90846_1|ARO_3003556|SLB-1__Shewanella_livingstonensis__TM_#02
TENDKTTAKSTFTGM
>gb|AAT90846_1|ARO_3003556|SLB-1__Shewanella_livingstonensis__TM_#03
PQVKIVVPGHGQVGDKALLEHTIELLIPKNETVNSSS
>gb|CAA75663_1|ARO_3000175|tet(H)__Pasteurella_multocida__TM_#01
RTPENQTASNTVTV
>gb|CAA75663_1|ARO_3000175|tet(H)__Pasteurella_multocida__TM_#02
KLAQKWGEKTTIMISMSIDMM
>gb|CAA75663_1|ARO_3000175|tet(H)__Pasteurella_multocida__TM_#03
LMGAILYAMLLITAYFHQRKTTPKAVISTP
>gb|ACQ82816_1|ARO_3004577|Acinetobacter_baumannii_AmvA__Acinetobacter_baumannii__TM_#01
AMLIPATLS
>gb|ACQ82816_1|ARO_3004577|Acinetobacter_baumannii_AmvA__Acinetobacter_baumannii__TM_#02
LAVLVMIVMIIPKQQEKTDQPINLG
>gb|ACQ82816_1|ARO_3004577|Acinetobacter_baumannii_AmvA__Acinetobacter_baumannii__TM_#03
RSQKRATTPMIDLE
>gb|ACQ82816_1|ARO_3004577|Acinetobacter_baumannii_AmvA__Acinetobacter_baumannii__TM_#04
GICLNKWGLRRVSSLGILVSALSLWGLAQLNFSTDHFLAWTCMVFLGFSIEIALLASTAA
IMSSV
>gb|ACQ82816_1|ARO_3004577|Acinetobacter_baumannii_AmvA__Acinetobacter_baumannii__TM_#05
SRSIILPAELPSNLIEKASISIGETMQLASNLENPLGGQLIVVAQQAFSYAHSWVLTLSA
ICFFLLTVFVWFSFPKKVN
>gb|AEP40501_1|ARO_3002969|vanXYN__Enterococcus_faecium__TM_#01
MHNFYLQLVNQQHPWKSFNHSPQLVQATYAEEKILIDSKVNHQFNQLLETLQLTDRIMIV
DGHRTVAEQKHLWNYSLNAHGVNYTKSYVAS
>gb|AEP40501_1|ARO_3002969|vanXYN__Enterococcus_faecium__TM_#02

TEHDLIAPRFEGPEAEL
>gb|AEP40501_1|ARO_3002969|vanXYN__Enterococcus_faecium__TM_#03
KHQIEAVS
>gb|BAC67143_1|ARO_3000166|tet(B)__Gram-negative_bacterium_TC71__TM_#01
DIANHFGVL
>gb|BAC67143_1|ARO_3000166|tet(B)__Gram-negative_bacterium_TC71__TM_#02
SFGLGLIAGPIIGGFAGEISP
>gb|BAC67143_1|ARO_3000166|tet(B)__Gram-negative_bacterium_TC71__TM_#03
LLHSVFQAFVAGRIATKWGE
>gb|ACF17980_1|ARO_3002616|aadA16__Escherichia_coli__TM_#01
LRALEVTIVV
>gb|ACF17980_1|ARO_3002616|aadA16__Escherichia_coli__TM_#02
VQHQPVLLEA
>gb|AAA88560_1|ARO_3004699|sta__Streptomyces_lavendulae__TM_#01
MTTTHGSTYEFRSAR
>gb|AAA88560_1|ARO_3004699|sta__Streptomyces_lavendulae__TM_#02
GGSDGEDGA
>gb|AAA88560_1|ARO_3004699|sta__Streptomyces_lavendulae__TM_#03
FCGLDSALYQGTASEGEHALYMSMPCP
>gb|CAC33832_1|ARO_3000851|THIN-B_beta-lactamase__Janthinobacterium_lividum__TM_#01
KPDTPVDCDSCKAWNGEVT
>gb|CAC33832_1|ARO_3000851|THIN-B_beta-lactamase__Janthinobacterium_lividum__TM_#02
KVKVVGEGDAIKLGPLN
>gb|CAC33832_1|ARO_3000851|THIN-B_beta-lactamase__Janthinobacterium_lividum__TM_#03
VHPDSTGVLDKAAKRSGEHNPFIDANACRAYAATADAMLTKRLAKERGVALPAAAPAAQH
AH
>gb|AAK37618_1|ARO_3005008|Txr__Vibrio_harveyi__TM_#01
MKVQKCKATIHRRSLDADPLL
>gb|AAK37618_1|ARO_3005008|Txr__Vibrio_harveyi__TM_#02
QIHKVGADKIIDVDCRVIAASHVDLKSEVIQGGFREDLFYRQ
>gb|AAK37618_1|ARO_3005008|Txr__Vibrio_harveyi__TM_#03
VILSERFLEELSNGKVQSMSPEVKK
>gb|BAM16262_1|ARO_3004628|AAC(2_)-IIa__Burkholderia_glumae__TM_#01
MKDRSHDDSMAEVCRNTSENHWLKTDYRTLFRLCPDG
>gb|BAM16262_1|ARO_3004628|AAC(2_)-IIa__Burkholderia_glumae__TM_#02
ADVPDDIALKLEELASVEPPFTPPAIPKHLERYLSLLGSDGPVTHDLGLIYELPHAQQYP
SKARLIGSGSEEGESLMQSWAEDR
>gb|AJD73064_1|ARO_3001301|RlmA(II)__Streptococcus_pneumoniae__TM_#01
SQTASLTKT
>gb|EOD99669_1|ARO_3002875|dfrE__Enterococcus_faecalis_EnGen0074__TM_#01
TLVLGRATFEGMGCRPLPNRTTIVLTSNPDYRAEGVLVMHSVEEILAYADNYEGVTVIGG
GSVVFKELIPA
>gb|EOD99669_1|ARO_3002875|dfrE__Enterococcus_faecalis_EnGen0074__TM_#02
FVWEKVATVPGVVDEKNLYAHDYETYHRNDK
>gb|ABV26705_1|ARO_3002849|arr-4__Pseudomonas_aeruginosa__TM_#01
NDWIPTSH
>gb|ABV26705_1|ARO_3002849|arr-4__Pseudomonas_aeruginosa__TM_#02
GDLLSPGHPSHFEQGR
>gb|ABV26705_1|ARO_3002849|arr-4__Pseudomonas_aeruginosa__TM_#03
LKHIYFAAL
>gb|ABV26705_1|ARO_3002849|arr-4__Pseudomonas_aeruginosa__TM_#04
VEDWQGHSP
>gb|ABV26705_1|ARO_3002849|arr-4__Pseudomonas_aeruginosa__TM_#05
VLQGMLASLEDLQRRGLAIIED
>gb|ALE30770_1|ARO_3004797|FRI-1__Enterobacter_cloacae__TM_#01
LCLPLNSFASQ
>gb|ALE30770_1|ARO_3004797|FRI-1__Enterobacter_cloacae__TM_#02
FGGRIGVYILN
>gb|ALE30770_1|ARO_3004797|FRI-1__Enterobacter_cloacae__TM_#03
KSVSLDDM
>gb|ALE30770_1|ARO_3004797|FRI-1__Enterobacter_cloacae__TM_#04
NIAFGSVLDAKNKSLLQ
>gb|ALE30770_1|ARO_3004797|FRI-1__Enterobacter_cloacae__TM_#05
VGDKTGTCG
>gb|ALE30770_1|ARO_3004797|FRI-1__Enterobacter_cloacae__TM_#06
DANSPAVMAVYTTRPNQNDKHDE
>gb|CAA45050_1|ARO_3002828|srmB__Streptomyces_ambofaciens__TM_#01
GESDENGSERELSAGLQR
>gb|CAA45050_1|ARO_3002828|srmB__Streptomyces_ambofaciens__TM_#02
GDGRIAEFSAG
>gb|AAA27471_1|ARO_3000205|tetX__Bacteroides_fragilis__TM_#01
NDTVIWDRKLV
>gb|AAA27471_1|ARO_3000205|tetX__Bacteroides_fragilis__TM_#02
SETADLVI
>gb|AAA27471_1|ARO_3000205|tetX__Bacteroides_fragilis__TM_#03
LLFANPNNNGAL
>gb|AAA27471_1|ARO_3000205|tetX__Bacteroides_fragilis__TM_#04
GISFKTPDEWK
>gb|AAW34150_1|ARO_3004191|APH(2__)-If__Campylobacter_jejuni__TM_#01
MDIKKIIEEKCNIVVDSIKLIGEGYDSKAYIVNN
>gb|CAG34233_2|ARO_3003019|dfrA23__Salmonella_enterica_subsp__enterica_serovar_Typhimurium__TM_#01
MPTVEIIVAVDPVGGFGRNGQIPWTCKEDMKRFTT
>gb|CAG34233_2|ARO_3003019|dfrA23__Salmonella_enterica_subsp__enterica_serovar_Typhimurium__TM_#02
KDMLDMQMKKEGAEERIKEKGILPERESYVVSSTLKP
>gb|CAG34233_2|ARO_3003019|dfrA23__Salmonella_enterica_subsp__enterica_serovar_Typhimurium__TM_#03
QYHDSDQRIAVIGGEKLYVQALASATKVHMTVMHKPYNCDRTLPMSYIDKKFVAGQGSIT
IQTAVDGETHPVKFITYERARP
>gb|AAA21889_1|ARO_3002554|AAC(6_)-Ig__Acinetobacter_haemolyticus__TM_#01
ELRNKLWSDS
>gb|AAA21889_1|ARO_3002554|AAC(6_)-Ig__Acinetobacter_haemolyticus__TM_#02
YALQLLAYSDHQAIAMLEASIRF
>gb|AAA21889_1|ARO_3002554|AAC(6_)-Ig__Acinetobacter_haemolyticus__TM_#03
AHRRSGVATMLIRQAEV
>gb|AAD03494_1|ARO_3002566|AAC(6_)-Iv__Acinetobacter_sp__631__TM_#01
ESQLSDWLVLRCLLWPDHE
>gb|AAD03494_1|ARO_3002566|AAC(6_)-Iv__Acinetobacter_sp__631__TM_#02
LQEMRQLITQAHC
>gb|AAD03494_1|ARO_3002566|AAC(6_)-Iv__Acinetobacter_sp__631__TM_#03
GLVQHVEIWAKQF
>gb|AAD03494_1|ARO_3002566|AAC(6_)-Iv__Acinetobacter_sp__631__TM_#04
CTEFASDA
>gb|AAK52604_1|ARO_3001425|OXA-31__Pseudomonas_aeruginosa__TM_#01
IIYSSASASTDISTVAS

>gb|AAK52604_1|ARO_3001425|OXA-31__Pseudomonas_aeruginosa__TM_#02
LFEGTEGCFLLYD
>gb|AAK52604_1|ARO_3001425|OXA-31__Pseudomonas_aeruginosa__TM_#03
EIAQFNKAKCA
>gb|AAK52604_1|ARO_3001425|OXA-31__Pseudomonas_aeruginosa__TM_#04
IDQKTIFKWDK
>gb|AAK52604_1|ARO_3001425|OXA-31__Pseudomonas_aeruginosa__TM_#05
PKGMEIWNSNHTPKTWMQFSVVWVSQEITQ
>gb|AAK52604_1|ARO_3001425|OXA-31__Pseudomonas_aeruginosa__TM_#06
QFLRKIINHNLPV
>gb|AAK52604_1|ARO_3001425|OXA-31__Pseudomonas_aeruginosa__TM_#07
NSAIENTI
>gb|AAK52604_1|ARO_3001425|OXA-31__Pseudomonas_aeruginosa__TM_#08
NSTKLYGKTGAGFTANRTLQNGWFEGFIISKSGHKYVFVSALTG
>gb|AAK52604_1|ARO_3001425|OXA-31__Pseudomonas_aeruginosa__TM_#09
LGSNLTSSIKAKK
>gb|AAK52604_1|ARO_3001425|OXA-31__Pseudomonas_aeruginosa__TM_#10
AITILNTLNL
>gb|AGC50807_1|ARO_3002307|VIM-37__Pseudomonas_aeruginosa__TM_#01
VIPGHGLPGGLDLL
>gb|AAG03546_1|ARO_3003679|TriA__Pseudomonas_aeruginosa_PAO1__TM_#01
MSDARGAFHSKGRWSRMALPAILCAGLLVGCGAEPPAEEHVRVLAQTVKMAEFASATSI
>gb|AAG03546_1|ARO_3003679|TriA__Pseudomonas_aeruginosa_PAO1__TM_#02
ENAQAAVAAQQAQSKLADLNYQRQKA
>gb|AAG03546_1|ARO_3003679|TriA__Pseudomonas_aeruginosa_PAO1__TM_#03
SVRSAQSSLK
>gb|AAG03546_1|ARO_3003679|TriA__Pseudomonas_aeruginosa_PAO1__TM_#04
DVDGQRITVSLLGKPEVTASGKVREITPTVDERSGTLKVKVGLDSVPAEMSLGSVVNASV
AAPAEHSVVLPWSALSKVGEQPAVWLLDQQGKARLQPVRVARYASEKVVIDGGLEAGQTV
VTVGGQLLHPGQVVEVAQPPQPTQSTASRDAVGGGQP
>gb|AAM45855_1|ARO_3004780|DES-1__Desulfovibrio_desulfuricans__TM_#01
MHSRSSYSRRYVLAGLCALPFASLSAGLIFNSSADAASLAAINGKTLQKKLAELEAASGG
RLGVAARSSNGGKS
>gb|AAM45855_1|ARO_3004780|DES-1__Desulfovibrio_desulfuricans__TM_#02
DKPGILEQRIHFAQI
>gb|AAM45855_1|ARO_3004780|DES-1__Desulfovibrio_desulfuricans__TM_#03
CNTLNGLLCGNLLKAPARER
>gb|AAM45855_1|ARO_3004780|DES-1__Desulfovibrio_desulfuricans__TM_#04
GKATKPGKNKG
>gb|AAM45855_1|ARO_3004780|DES-1__Desulfovibrio_desulfuricans__TM_#05
SATRLVCAAEGLAMPLDNMY
>gb|ACH58985_1|ARO_3002485|LRA-2__uncultured_bacterium_BLR2__TM_#01
MMDGIKKKTAAGAAAGSLLMMLGVFATPAAGGEAAFKDCPQCAQWNQQRKPFRIYGNTYF
VGTA
>gb|ACH58985_1|ARO_3002485|LRA-2__uncultured_bacterium_BLR2__TM_#02
QSGAQVYALRTAEAVLRTGRLTQDDPQSASKTATITPVPQVWVVQDDQLLGVGALRMR
>gb|ACH58985_1|ARO_3002485|LRA-2__uncultured_bacterium_BLR2__TM_#03
DGNCLKMIYADSLSAVAAGKYRFKDHPEVLQAFASSFSRAESA
>gb|ACH58985_1|ARO_3002485|LRA-2__uncultured_bacterium_BLR2__TM_#04
DASQLFQRLDPEGGTRAASIKDDTACRRYVQAARDTLARKLASEG
>gb|AAB36568_1|ARO_3002700|cmlv__Streptomyces_venezuelae_ATCC_10712__TM_#01
MPSPSAEPTTSTPTPDAGP
>gb|AAB36568_1|ARO_3002700|cmlv__Streptomyces_venezuelae_ATCC_10712__TM_#02
RVPAGAFLGEHLGWA
>gb|AAB36568_1|ARO_3002700|cmlv__Streptomyces_venezuelae_ATCC_10712__TM_#03
GTAALALRLTKRPAPGHVVARSRGA
>gb|AAL82588_1|ARO_3002600|AAC(3)-Ib_AAC(6_)-Ib____Pseudomonas_aeruginosa__TM_#01
MSIIATVKIGPDEISAMRAVLDLFGKEFEDIPTYSDRQPTNEYLANLLHSETFIALAAFD
RG
>gb|AAL82588_1|ARO_3002600|AAC(3)-Ib_AAC(6_)-Ib____Pseudomonas_aeruginosa__TM_#02
LGVATALISHLKRVAV
>gb|CBH51824_1|ARO_3002629|ANT(6)-Ib__Campylobacter_fetus_subsp__fetus__TM_#01
MKMRTEKQIYDTILNF
>gb|CBH51824_1|ARO_3002629|ANT(6)-Ib__Campylobacter_fetus_subsp__fetus__TM_#02
TNIPVPTDEDYYIEH
>gb|CBH51824_1|ARO_3002629|ANT(6)-Ib__Campylobacter_fetus_subsp__fetus__TM_#03
QGYSFSLGKNYKFLERYISPELWKKILATYNMGSYTEMWKSLELCMGIFRMVSKEVAQCL
NYLYPDYDKNISNYVIRQKEKYQR
>gb|ACH59005_1|ARO_3002513|LRA-19__uncultured_bacterium_BLR19__TM_#01
MNSEMSQTSFKIRILVTCLLSIAQLTMAQQVQVTEPPITNQ
>gb|ACH59005_1|ARO_3002513|LRA-19__uncultured_bacterium_BLR19__TM_#02
NAQLMIDEKDSP
>gb|ACH59005_1|ARO_3002513|LRA-19__uncultured_bacterium_BLR19__TM_#03
ELFGSTGSTYEPVKADRLLKNGDKIT
>gb|ACH59005_1|ARO_3002513|LRA-19__uncultured_bacterium_BLR19__TM_#04
SKKFSDIPTYPGIAEDYTYTFDAMKKVH
>gb|ACH59005_1|ARO_3002513|LRA-19__uncultured_bacterium_BLR19__TM_#05
EAYNPGVFIDRAGYDKAVGDLEDKFSKKKQADK
>gb|BAA32493_1|ARO_3000423|FomA__Streptomyces_wedmorensis__TM_#01
SLDDDAVTPFARNFARLAETYR
>gb|BAA32493_1|ARO_3000423|FomA__Streptomyces_wedmorensis__TM_#02
DHDSTHAF
>gb|BAA32493_1|ARO_3000423|FomA__Streptomyces_wedmorensis__TM_#03
EKLRGIGVDAFPLQLAAMCTLRNGIPQLRSEVLRDVLDHGALPVLAGDALFDEHGKLWAF
SSDRVPEVLLPMVE
>gb|BAA32493_1|ARO_3000423|FomA__Streptomyces_wedmorensis__TM_#04
TILPEVDARSPEQAYAALWGSSEWDATGAMHT
>gb|BAA32493_1|ARO_3000423|FomA__Streptomyces_wedmorensis__TM_#05
SDLEFLTAPFSSWPAHVRSTRITTTASA
>gb|AHA41504_1|ARO_3002941|vanSO__Rhodococcus_hoagii__TM_#01
SVAVRLPAAQRRP
>gb|CDL65151_1|ARO_3003971|erm(44)__Staphylococcus_xylosus__TM_#01
NEILNETNIGI
>gb|CDL65151_1|ARO_3003971|erm(44)__Staphylococcus_xylosus__TM_#02
YMSNIARFITSIEIDKALYCNLKNDISLSTNIELVNKDILIYEFPKYKQ
>gb|CDL65151_1|ARO_3003971|erm(44)__Staphylococcus_xylosus__TM_#03
LYESNAEYNYLI
>gb|CDL65151_1|ARO_3003971|erm(44)__Staphylococcus_xylosus__TM_#04
ILKVIPNS
>gb|ABA71731_1|ARO_3002909|vanG__Enterococcus_faecalis__TM_#01
TNKFDIIPIGITRSGEWYHYTGEKEKILNNTWFEDSKNLCPVVVSQNRSVKGFLEIASDK
YRII
>gb|ABA71731_1|ARO_3002909|vanG__Enterococcus_faecalis__TM_#02

RFNEEAAMKEIEAN
>gb|AGU01679_2|ARO_3003894|Rm3_beta-lactamase__uncultured_bacterium__TM_#01
MSLTPPRALVLALLASPGTQAQTPAPATPPTPGCEVCATWNADQAPFRL
>gb|AGU01679_2|ARO_3003894|Rm3_beta-lactamase__uncultured_bacterium__TM_#02
PRCLNMVYADSINA
>gb|AGU01679_2|ARO_3003894|Rm3_beta-lactamase__uncultured_bacterium__TM_#03
ADLRHSFETLEKLPCDV
>gb|AGU01679_2|ARO_3003894|Rm3_beta-lactamase__uncultured_bacterium__TM_#04
GGSDAFVDPQACRAYVAAARTLLDSRLDQEKQQ
>gb|BAA34299_1|ARO_3003034|mexX__Pseudomonas_aeruginosa_PAO1__TM_#01
MDRLAARLLAALVALFLLGCEEAADAGKTAE
>gb|BAA34299_1|ARO_3003034|mexX__Pseudomonas_aeruginosa_PAO1__TM_#02
PAPIGITS
>gb|BAA34299_1|ARO_3003034|mexX__Pseudomonas_aeruginosa_PAO1__TM_#03
PGRGQPRAAADKLKA
>gb|BAA34299_1|ARO_3003034|mexX__Pseudomonas_aeruginosa_PAO1__TM_#04
KSRHAAGDPRRPGEGCRRQGHRRAPGPGRRQRVPLAGE
>gb|BAA34299_1|ARO_3003034|mexX__Pseudomonas_aeruginosa_PAO1__TM_#05
DWIISRGLKGGEWVIVENAAQHAAGSSVQAVVRQPASADAPSPLAASPAGQ
>gb|CAC14594_1|ARO_3003055|smeD__Stenotrophomonas_maltophilia__TM_#01
MLLSRIRPFA
>gb|CAD91341_1|ARO_3004675|aphA15__Pseudomonas_aeruginosa__TM_#01
MTVALDEVSELKNL
>gb|CAD91341_1|ARO_3004675|aphA15__Pseudomonas_aeruginosa__TM_#02
DARVIRVLFPDRNTAYLKYASGSSAQEILQEHQRTRWLRTRALVPEVISYVSTSTVTILL
TKALIGHNAADAA
>gb|CAD91341_1|ARO_3004675|aphA15__Pseudomonas_aeruginosa__TM_#03
IVVAEMARALRDLHSISPDDCPFDERLHLRLKLASGRLEAGLVDEEDFDHARQGMLARDV
YEQLF
>gb|CAD91341_1|ARO_3004675|aphA15__Pseudomonas_aeruginosa__TM_#04
PENFIFQGNAFVGFIDCGRVGLADKYQDLALASRNIDAVFGPELTNQFFIEYGEPNPNIA
KIEYYRILDEFF
>gb|ADX95999_1|ARO_3002873|FosB3__Enterococcus_faecium__TM_#01
MIKGINHITYSVSNIAKSIEFYRDIL
>gb|ADX95999_1|ARO_3002873|FosB3__Enterococcus_faecium__TM_#02
ADILVESET
>gb|ADX95999_1|ARO_3002873|FosB3__Enterococcus_faecium__TM_#03
SDNDFEDWY
>gb|ADX95999_1|ARO_3002873|FosB3__Enterococcus_faecium__TM_#04
WLKENEVNILEG
>gb|ADX95999_1|ARO_3002873|FosB3__Enterococcus_faecium__TM_#05
EAKPHMNFYI
>gb|AAL19306_1|ARO_3000789|mdsA__Salmonella_enterica_subsp__enterica_serovar_Typhimurium_str__LT2__TM_#01
PPSVPVAKALSRTLAPTAEFTGFLAAPE
>gb|AAL19306_1|ARO_3000789|mdsA__Salmonella_enterica_subsp__enterica_serovar_Typhimurium_str__LT2__TM_#02
ASGAVSRKNADDVTATRNARQ
>gb|AAL19306_1|ARO_3000789|mdsA__Salmonella_enterica_subsp__enterica_serovar_Typhimurium_str__LT2__TM_#03
KNPPVVNMGLTTDNGLPYQGVLDFMGNQMNRSTGTIRARAVIPDPDGMLSPGLFARISLP
IGEPRETV
>gb|AAL19306_1|ARO_3000789|mdsA__Salmonella_enterica_subsp__enterica_serovar_Typhimurium_str__LT2__TM_#04
FRVVTQGVL
>gb|QGQ32905_1|ARO_3005015|VMB-1__Vibrio_alginolyticus__TM_#01
MKYIILFLLLIPSIVFANNEGTKELKLKKLSDNVYQHISYKRVEPWGLIGASGLVVINGT
EAHMIDTPWTTQGTKQLIEWIEAKGLTIKSAVVTHFHEDASGDIPLLNDLKIKTYATSLT
NKLLKLNQKEVSSDEISSNTFEFIDGVAS
>gb|QGQ32905_1|ARO_3005015|VMB-1__Vibrio_alginolyticus__TM_#02
YTGDANISEWPNSMQKVINRYPDAKL
>gb|QGQ32905_1|ARO_3005015|VMB-1__Vibrio_alginolyticus__TM_#03
AASNKKINKD
>gb|ABY55281_1|ARO_3004498|dfrB4__Escherichia_coli__TM_#01
MNEGKNEVSTSAAGRFAFPSNATFALGDRVRKKSGAAWQGRIVGWYCTT
>gb|ABY55281_1|ARO_3004498|dfrB4__Escherichia_coli__TM_#02
DSVQIYPMTALERVA
>gb|CDO12042_1|ARO_3005053|ArnT__Klebsiella_pneumoniae__TM_#01
VGLLGLIAVLVVSPWGF
>gb|CAD91132_1|ARO_3002839|lnuF__Escherichia_coli__TM_#01
CHEDARIIAALMFGSFAI
>gb|CAD91132_1|ARO_3002839|lnuF__Escherichia_coli__TM_#02
HFENFDQRSWLNAVSPVAAYFP
>gb|CAD91132_1|ARO_3002839|lnuF__Escherichia_coli__TM_#03
KSDIPVIST
>gb|CAD91132_1|ARO_3002839|lnuF__Escherichia_coli__TM_#04
SGELSRYASALVG
>gb|CAD91132_1|ARO_3002839|lnuF__Escherichia_coli__TM_#05
REGAPLVEGLVLNLISLMLFGANLLNRGEYARAWALLSKAHENLLKLVRLHEGATDHWPT
PSRALEKD
>gb|CAD91132_1|ARO_3002839|lnuF__Escherichia_coli__TM_#06
SEDSYNRYLACT
>gb|CAD91132_1|ARO_3002839|lnuF__Escherichia_coli__TM_#07
PLNIELPR
>gb|CAD91132_1|ARO_3002839|lnuF__Escherichia_coli__TM_#08
KRLLNESATPHNK
>gb|BAG12271_1|ARO_3004674|FosD__Staphylococcus_aureus__TM_#01
MIQSINHICYSVSDLKNSIR
>gb|BAG12271_1|ARO_3004674|FosD__Staphylococcus_aureus__TM_#02
DESEFNDWYQWFKE
>gb|CAA10975_1|ARO_3000194|tetW__Butyrivibrio_fibrisolvens__TM_#01
EKYKLETVV
>gb|AAA20117_1|ARO_3000195|tetB(P)__Clostridium_perfringens__TM_#01
AVYIKKLFDTCI
>gb|AAA20117_1|ARO_3000195|tetB(P)__Clostridium_perfringens__TM_#02
FASNDCESDLSGVVFKIERTSK
>gb|AAA20117_1|ARO_3000195|tetB(P)__Clostridium_perfringens__TM_#03
RLENGGVVEAQR
>gb|AAA20117_1|ARO_3000195|tetB(P)__Clostridium_perfringens__TM_#04
FGASIMHMQEDLNPFWATVGLE
>gb|ACZ72746_1|ARO_3004808|GOB-16__Elizabethkingia_meningoseptica__TM_#01
MRNFAILFFLLITFSWKAQVVKEPENTN
>gb|ACZ72746_1|ARO_3004808|GOB-16__Elizabethkingia_meningoseptica__TM_#02
EGDPYNPQVFMDKANYFAFLNSLETDYLEKIKNDSQKK
>gb|AAD51347_1|ARO_3003067|smeS__Stenotrophomonas_maltophilia__TM_#01
MAFAMAKFQ
>gb|AAD51347_1|ARO_3003067|smeS__Stenotrophomonas_maltophilia__TM_#02

```
EHLHGDGDRWARLLRPDLAHGHEGPVPSLSDQTGVPSRLGLFDAQHRFVAGNPDATSDDE
PHAVQVDGQTVGWLGMVPFQTVIATNDLNFYNTQVRAWWVIGIALLLVTVLLAWLVSRAL
RQRLAKL
>gb|AAD51347_1|ARO_3003067|smeS__Stenotrophomonas_maltophilia__TM_#03
ERTSDDELDALVNDFNRMAQALDDTERN
>gb|AAD51347_1|ARO_3003067|smeS__Stenotrophomonas_maltophilia__TM_#04
RANLVGLQGEIRQLGKLIDDLHDLSMTQSGGLAYRFAPLDLVALLRSELNGMRVRFANAG
LALEEDLPAT
>gb|AAD51347_1|ARO_3003067|smeS__Stenotrophomonas_maltophilia__TM_#05
ARVPAGVQLVVEDTAPGVPPDKC
>gb|AAD51347_1|ARO_3003067|smeS__Stenotrophomonas_maltophilia__TM_#06
ILAHHGVIHAAPSPLGGLRVVITLPEPA
>gb|CCC86797_1|ARO_3005046|MecC-type_methicillin_resistance_repressor_MecI__Staphylococcus_aureus_subsp__aureus_LGA251__TM_#01
MTREGYDISASEWEIMNTIWNKKLISANDVIEIVQKHKEWSPKTIRTLINRLYKKKFIDR
TSRNKIFEYFPIVEEKDMKYKTSKVFLDKVYEGGLNSLVLNFVENEELSEDDIEELKNIL
NNKY
>gb|AMS25623_1|ARO_3005366|KPC-34__Klebsiella_pneumoniae__TM_#01
MSLYRRLVLLSCLSWPLAGFSATALTNLVAEPF
>gb|AMS25623_1|ARO_3005366|KPC-34__Klebsiella_pneumoniae__TM_#02
IRYGKNALV
>gb|AMS25623_1|ARO_3005366|KPC-34__Klebsiella_pneumoniae__TM_#03
WSPISEKYLTTG
>gb|AMS25623_1|ARO_3005366|KPC-34__Klebsiella_pneumoniae__TM_#04
VYTRAPNKPNKDDKHSEAKDDK
>gb|ABF33001_1|ARO_3003969|lmrP__Streptococcus_pyogenes_MGAS9429__TM_#01
CLFVVIALLGIYFTVVSAKMKKV
>gb|CAJ77855_1|ARO_3000779|adeH__Acinetobacter_baumannii_AYE__TM_#01
MVITSKQN
>gb|CAJ77855_1|ARO_3000779|adeH__Acinetobacter_baumannii_AYE__TM_#02
KVIVPVKFKESDPKLEDNNWKIAQPADQ
>gb|CAJ77855_1|ARO_3000779|adeH__Acinetobacter_baumannii_AYE__TM_#03
LDTEQAIYNRTIKLLGETRDLMQLRFKNGLVSELDVSRAQTELATAQTT
>gb|CAJ77855_1|ARO_3000779|adeH__Acinetobacter_baumannii_AYE__TM_#04
VQPLTANS
>gb|OOS23853_1|ARO_3005073|ROB-3__Moraxella_pluranimalium__TM_#01
SNPQPASAPVQQSATQATFQQT
>gb|OOS23853_1|ARO_3005073|ROB-3__Moraxella_pluranimalium__TM_#02
ANLEQQYQARIGVYVWDTETGHS
>gb|OOS23853_1|ARO_3005073|ROB-3__Moraxella_pluranimalium__TM_#03
SLPEKDLNRTISYSQKDLVSYSPETQKYVGKGMTIAQLCEAAVRFSDN
>gb|OOS23853_1|ARO_3005073|ROB-3__Moraxella_pluranimalium__TM_#04
ATNLLLKELGGVEQYQRILRQLGDNVTH
>gb|OOS23853_1|ARO_3005073|ROB-3__Moraxella_pluranimalium__TM_#05
NRLEPDLNQAKPNDIRDTSTPKQMAMNLNAYLLGNTLTESQ
>gb|OOS23853_1|ARO_3005073|ROB-3__Moraxella_pluranimalium__TM_#06
KYGVRNDIAVVRI
>gb|OOS23853_1|ARO_3005073|ROB-3__Moraxella_pluranimalium__TM_#07
TQFTEEAKFNNKLVED
>gb|OOS23853_1|ARO_3005073|ROB-3__Moraxella_pluranimalium__TM_#08
AKQVFHTLQLN
>gb|AVL76727_1|ARO_3004663|FusF__Staphylococcus_cohnii__TM_#01
TTKIDIYQQFHQIDDTLTEAIEKLMNIRITKVQVDKILETLQTYVIPFEHPSKKQVEKTF
RKIKKLKSPLISDEILL
>gb|AVL76727_1|ARO_3004663|FusF__Staphylococcus_cohnii__TM_#02
EQGTLTGFYGDIANQTVKGYCAICNKESN
>gb|AVL76727_1|ARO_3004663|FusF__Staphylococcus_cohnii__TM_#03
IKCNQQLSDITQFYQFVDKIHS
>gb|QIB98918_1|ARO_3005012|IDC-1__sediment_metagenome__TM_#01
MPRTESVPSKSLVVRTLLLVFACLFPMAVPAVED
>gb|QIB98918_1|ARO_3005012|IDC-1__sediment_metagenome__TM_#02
VDAAILPLMSQHDIPGM
>gb|QIB98918_1|ARO_3005012|IDC-1__sediment_metagenome__TM_#03
VGLILDGQPYVVTYGVASKE
>gb|QIB98918_1|ARO_3005012|IDC-1__sediment_metagenome__TM_#04
YFRNWTPLAPPGTRREYSNASPGLLG
>gb|QIB98918_1|ARO_3005012|IDC-1__sediment_metagenome__TM_#05
VAASALDDDFATLMQ
>gb|QIB98918_1|ARO_3005012|IDC-1__sediment_metagenome__TM_#06
TVFPAFGMTDSFIHVPDRKMPDYAWGYRKDR
>gb|QIB98918_1|ARO_3005012|IDC-1__sediment_metagenome__TM_#07
VRVNEGPLDEQAYGVKTTVSDLLRFVQANIDP
>gb|QIB98918_1|ARO_3005012|IDC-1__sediment_metagenome__TM_#08
AVEATQVGYFRAGTLVQGLGWEK
>gb|BAD63613_1|ARO_3002816|clbC__Bacillus_clausii_KSM-K16__TM_#01
MKVVNHATKYERLKHFLNALNEPT
>gb|BAD63613_1|ARO_3002816|clbC__Bacillus_clausii_KSM-K16__TM_#02
AFNKMTTLPKALRESLINEFGPSILTVEPVLETTSQQVTKVLLKVAGNNQVEAVRMH
>gb|BAD63613_1|ARO_3002816|clbC__Bacillus_clausii_KSM-K16__TM_#03
TFCSTGAIGLKQ
>gb|BAD63613_1|ARO_3002816|clbC__Bacillus_clausii_KSM-K16__TM_#04
RIFDALNVLVDRQ
>gb|BAD63613_1|ARO_3002816|clbC__Bacillus_clausii_KSM-K16__TM_#05
DQVMNVLDQHIHE
>gb|BAD63613_1|ARO_3002816|clbC__Bacillus_clausii_KSM-K16__TM_#06
EKHAEALVKRILNNRYP
>gb|BAD63613_1|ARO_3002816|clbC__Bacillus_clausii_KSM-K16__TM_#07
GTPENYGQTIEEKLQTFYRVVKSARIPVTIRSQFGR
>gb|APM84516_1|ARO_3005150|PAC-1__Pseudomonas_aeruginosa__TM_#01
MRCNKNVLSVVLLGALSLSAGNAFGQVSQADVDAVIRPLMSKYKIPGMAVALSVDGQHT
>gb|APM84516_1|ARO_3005150|PAC-1__Pseudomonas_aeruginosa__TM_#02
QWSDQASHY
>gb|APM84516_1|ARO_3005150|PAC-1__Pseudomonas_aeruginosa__TM_#03
AWYQAWQPTA
>gb|APM84516_1|ARO_3005150|PAC-1__Pseudomonas_aeruginosa__TM_#04
QKPFSEAMEQDLLAPLGMKHSWVKVPENQMAE
>gb|APM84516_1|ARO_3005150|PAC-1__Pseudomonas_aeruginosa__TM_#05
DLNMAITPPSPAWQQAITETHK
>gb|APM84516_1|ARO_3005150|PAC-1__Pseudomonas_aeruginosa__TM_#06
ETLLAGNSSERIMKGLGAKPLTPPQAG
>gb|APM84516_1|ARO_3005150|PAC-1__Pseudomonas_aeruginosa__TM_#07
KTALILLANKWYPNDARIEAAYELVQRLKK
>gb|AAP43109_1|ARO_3003922|oqxA__Escherichia_coli__TM_#01
RGQGASSDN
```

>gb|DAC81085_1|ARO_3005088|msrH__Macrococcus_canis__TM_#01
SRGKVNQYVEYGYFEQLEGPIAKATNPRLLGKLQVKENAS
>gb|DAC81085_1|ARO_3005088|msrH__Macrococcus_canis__TM_#02
TVKIYSGNYSDYMRQKQIERTQQVYNHEQ
>gb|DAC81085_1|ARO_3005088|msrH__Macrococcus_canis__TM_#03
SLEKMKKAEKIAKST
>gb|DAC81085_1|ARO_3005088|msrH__Macrococcus_canis__TM_#04
NQLQVVEAPNDDYVIHFHSTNNISKLHNKFPIMGDCITLEID
>gb|DAC81085_1|ARO_3005088|msrH__Macrococcus_canis__TM_#05
EIGIYEQMGYQFDEDKTVLSYIKDR
>gb|DAC81085_1|ARO_3005088|msrH__Macrococcus_canis__TM_#06
KEYKGTVIIISHDKKFVEHVSDIVYRIENQKLKLVN
>gb|AAG07065_1|ARO_3003692|mexJ__Pseudomonas_aeruginosa_PAO1__TM_#01
MYRHIPLVALSLFSSLFLAACGNGTPPPAAARPAIVVQPQPAGEVSQAFPGEIRARH
>gb|AAG07065_1|ARO_3003692|mexJ__Pseudomonas_aeruginosa_PAO1__TM_#02
RRYRTLLDRNLVSHSQFENIQNS
>gb|AAG07065_1|ARO_3003692|mexJ__Pseudomonas_aeruginosa_PAO1__TM_#03
ATPAELGQSARVYVAAAEA
>gb|AAG07065_1|ARO_3003692|mexJ__Pseudomonas_aeruginosa_PAO1__TM_#04
VVEPGSSTLRRQA
>gb|AAG07065_1|ARO_3003692|mexJ__Pseudomonas_aeruginosa_PAO1__TM_#05
ANRTVKLAAKE
>gb|ABA42118_2|ARO_3000866|oleI__Streptomyces_antibioticus__TM_#01
MTSEHRSASVTPRHISFFNIPG
>gb|ABA42118_2|ARO_3000866|oleI__Streptomyces_antibioticus__TM_#02
AAQVKAAGATPVVYDSILPKESNPEESWPEDQESAMGLFLDEAVRV
>gb|ABA42118_2|ARO_3000866|oleI__Streptomyces_antibioticus__TM_#03
RPDLIVYDIASWPAPVLGRKWDIPF
>gb|ABA42118_2|ARO_3000866|oleI__Streptomyces_antibioticus__TM_#04
VPAVQDPTADRGEEAAAPAGTGDAEEGAEAEDGLVRFFTRLSAFLEEHGVDTPATEFLIA
PNRCIVALPRTFQIKGDTVGDNYTFVGPTYGDRSHQGTWEGPGDGRPVLLIALGSAFTDH
LDFYRTCLSAVDG
>gb|ABA42118_2|ARO_3000866|oleI__Streptomyces_antibioticus__TM_#05
NAVPMVAVPQIAEQTMNAERIVELGLGRHIPRDQVTAEKLREAVLAVASDPGVAERLAAV
RQEIREAGGARAAA
>gb|BAJ10053_1|ARO_3002874|FosC2__Escherichia_coli__TM_#01
EATWICLSCDEVHPSQDYCHIAFDVSEENFEPVTKKLREAHVV
>gb|BAJ10053_1|ARO_3002874|FosC2__Escherichia_coli__TM_#02
QSRLESLKSKPYQGLVWL
>gb|CCQ22388_1|ARO_3000421|norB__Listeria_monocytogenes__TM_#01
NTKAKFDSFGLVLFVIAMVCLNLIITRGATFGWTSPITITMLVVFLV
>gb|CCQ22388_1|ARO_3000421|norB__Listeria_monocytogenes__TM_#02
SGITAVGIALMALTFIPG
>gb|CCQ22388_1|ARO_3000421|norB__Listeria_monocytogenes__TM_#03
VVSLISVAITTPSAKKALELKAAKE
>gb|AAA99931_1|ARO_3003060|tsnR__Streptomyces_laurentii__TM_#01
MANLDVIV
>gb|AAA99931_1|ARO_3003060|tsnR__Streptomyces_laurentii__TM_#02
TQSIRAGVEFTEVYGLDTVPFPGDLLAACEKRGIR
>gb|AAA99931_1|ARO_3003060|tsnR__Streptomyces_laurentii__TM_#03
PAGRFADLES
>gb|AAA99931_1|ARO_3003060|tsnR__Streptomyces_laurentii__TM_#04
RSALGAAGIVLVDSG
>gb|AAA99931_1|ARO_3003060|tsnR__Streptomyces_laurentii__TM_#05
AFFRDGGMRPVVFEADGKLSIGELD
>gb|AAA99931_1|ARO_3003060|tsnR__Streptomyces_laurentii__TM_#06
RRNLSRPRG
>gb|CBH51823_1|ARO_3000556|tet44__Campylobacter_fetus_subsp__fetus__TM_#01
EKYIVGEKLTIQELM
>gb|CBH51823_1|ARO_3000556|tet44__Campylobacter_fetus_subsp__fetus__TM_#02
KTVCSGDIFI
>gb|AAB41957_1|ARO_3000801|MexD__Pseudomonas_aeruginosa__TM_#01
PARRCRAPGTRQGELQHFLATERHAHRGRGYPAVARGQRDPDPT
>gb|AAB41957_1|ARO_3000801|MexD__Pseudomonas_aeruginosa__TM_#02
ELYVPNAAGNLVPLSAFVSVK
>gb|AAB41957_1|ARO_3000801|MexD__Pseudomonas_aeruginosa__TM_#03
PAPIEQAASAGE
>gb|AKQ05899_1|ARO_3004591|tet(55)__uncultured_bacterium__TM_#01
LCYWLNHYGFQPTLVEKNQSTRKGGYAIDLRGIAVDVAKQMGIYDSVCAMRTSLQCVRYV
DAA
>gb|AKQ05899_1|ARO_3004591|tet(55)__uncultured_bacterium__TM_#02
KTITDIPCFYDHAIESLTQHDDHVTVQFKNGKT
>gb|AKQ05899_1|ARO_3004591|tet(55)__uncultured_bacterium__TM_#03
SKDDYHLRNLGC
>gb|AKQ05899_1|ARO_3004591|tet(55)__uncultured_bacterium__TM_#04
QLDHCETLLEAKQKLVSITSDKDSTKAFAGFMFRSSNSPNYIRDEASQKDFLRENFTNHG
WESNKLLSLMNDA
>gb|AKQ05899_1|ARO_3004591|tet(55)__uncultured_bacterium__TM_#05
TDHVAAFARYNELLKPYVEANQA
>gb|AKQ05899_1|ARO_3004591|tet(55)__uncultured_bacterium__TM_#06
ADEPLSAEQAEERNNIVLGIMKKATHAIELPEY
>gb|ADC55560_1|ARO_3000858|armA__Pseudomonas_aeruginosa__TM_#01
MDKNDVVKKILESKKYENLDSDIVEKVVSISEKKYKLKEVENYSKKKLHQIWG
>gb|ADC55560_1|ARO_3000858|armA__Pseudomonas_aeruginosa__TM_#02
RVATLNDFYTYVFGNIKHVSSILDFGCGFNPLALYQWNENEKIIYHAYDIDRAEIAFLSS
IIGKLKTTIKYRFLNKESDVYKGTYDVVFLLKMLPVLKQQDVNILDFLQLFHTQNFVISF
PIKSLSGKEKGMEENYQLWFESFTKGWIKILDSKVIGNELVYITSGFQK
>gb|ABV63006_2|ARO_3004759|BPU-1__Bacillus_pumilus_SAFR-032__TM_#01
MKKKYIKRLFSCIMLMAFCISQPSSTEARSIAWSVDEFF
>gb|ACH58997_1|ARO_3002492|LRA-18__uncultured_bacterium_BLR18__TM_#01
MLKRIRLPQLALALAALFPLAAYAAPDAA
>gb|ACH58997_1|ARO_3002492|LRA-18__uncultured_bacterium_BLR18__TM_#02
KYVPQLQGSALDG
>gb|ACH58997_1|ARO_3002492|LRA-18__uncultured_bacterium_BLR18__TM_#03
NLKTREQLFSYFQHWKPDAAPGK
>gb|ACH58997_1|ARO_3002492|LRA-18__uncultured_bacterium_BLR18__TM_#04
HYAWGYSKDQAVRVQPDL
>gb|ACH58997_1|ARO_3002492|LRA-18__uncultured_bacterium_BLR18__TM_#05
QIDPSRLAAPMRRAVQ
>gb|ACH58997_1|ARO_3002492|LRA-18__uncultured_bacterium_BLR18__TM_#06
QGNSTDMAWKPQPVQAIQPVQTA
>gb|ACH58997_1|ARO_3002492|LRA-18__uncultured_bacterium_BLR18__TM_#07
RAYPNDARIKLAYAILNQLAPAAN

>gb|AHH86051_1|ARO_3004063|EdeQ__Brevibacillus_brevis__TM_#01
EPTPEQSEFVAPNLYSIAESKFQTTFVPLAIYHDDTMVGF
>gb|AHH86051_1|ARO_3004063|EdeQ__Brevibacillus_brevis__TM_#02
RLGYGRTAISQVIEILKAKEDCQKIVIGYAPANVAAENLYASLGFQKNGMVLFGETIAEL
NF
>gb|AAG03547_1|ARO_3003680|TriB__Pseudomonas_aeruginosa_PAO1__TM_#01
TNGRIASRLFDVGDF
>gb|AAG03547_1|ARO_3003680|TriB__Pseudomonas_aeruginosa_PAO1__TM_#02
QARLDDARTRLKTSQASF
>gb|AAG03547_1|ARO_3003680|TriB__Pseudomonas_aeruginosa_PAO1__TM_#03
RLVTDFDGVITTWH
>gb|AAG03547_1|ARO_3003680|TriB__Pseudomonas_aeruginosa_PAO1__TM_#04
TEVAESLPADARFLVSAQLDPQARTTGSIRELG
>gb|AAG03547_1|ARO_3003680|TriB__Pseudomonas_aeruginosa_PAO1__TM_#05
EAFRLGSTIQVQLSSAGSVRSVLPA
>gb|AAG07763_1|ARO_3003031|mexW__Pseudomonas_aeruginosa_PAO1__TM_#01
RNGQLVALSTLIET
>gb|AGC92784_1|ARO_3000848|DIM-1_beta-lactamase__Enterobacter_sp__SL1__TM_#01
MRTHFTALLLLFSLSSLANDEVPELRIEKVKENIFLHT
>gb|AGC92784_1|ARO_3000848|DIM-1_beta-lactamase__Enterobacter_sp__SL1__TM_#02
KGNAFIVDTPWSDRDTETLVHWIRKNGYELLGSVSTHW
>gb|AGC92784_1|ARO_3000848|DIM-1_beta-lactamase__Enterobacter_sp__SL1__TM_#03
DQSISTYATTSTNHLLKENKKEPAKYTLKGNESTLVDG
>gb|AGC92784_1|ARO_3000848|DIM-1_beta-lactamase__Enterobacter_sp__SL1__TM_#04
DSEGLGYTGEAHIDQWSRSAQNALSRYSEAQ
>gb|AGC92784_1|ARO_3000848|DIM-1_beta-lactamase__Enterobacter_sp__SL1__TM_#05
IALLKHTKSLAETASNKSIQPNANASAD
>gb|AGE00988_1|ARO_3002668|rmtG__Klebsiella_pneumoniae__TM_#01
MRDPLFEKLA
>gb|AGE00988_1|ARO_3002668|rmtG__Klebsiella_pneumoniae__TM_#02
LTECRAKYRREKEIDKAAREKLHGITAAFMTDAEYRRAMEIAV
>gb|AGE00988_1|ARO_3002668|rmtG__Klebsiella_pneumoniae__TM_#03
QNRYPEMRVTGIDISGQCVRVLRALGVDARLGDLLAENAIPRARYSVALLFKILPLLDRQ
SAGAARRILEAVNADALIC
>gb|AGE00988_1|ARO_3002668|rmtG__Klebsiella_pneumoniae__TM_#04
AVHYAAWMRDQLPEKWR
>gb|AAG06465_1|ARO_3005063|cprR__Pseudomonas_aeruginosa_PAO1__TM_#01
MHIHVLVVEDNFDLAGTVIDYLEAAGVVCDHARDGQAGLNLARANRYDVILLDIMLPRIN
GRQVCRQLREAGL
>gb|AAG06465_1|ARO_3005063|cprR__Pseudomonas_aeruginosa_PAO1__TM_#02
QRLQVDDLVMDLDSRQASRGGTPLALSPTAWKILECLMRASPALVTREQLGRSVWGDEPP
ESNTLNVHMHHLRSTVDKGFATPLIHTLHSVGFQLERK
>gb|QEE23059_1|ARO_3005060|OXA-837__Cupriavidus_gilardii__TM_#01
MKSRTEAQSLHAHG
>gb|QEE23059_1|ARO_3005060|OXA-837__Cupriavidus_gilardii__TM_#02
ALADVAPASAVAAPSEVARGDLMSHFRALDVDGCFALFDVKANRMTLVNASRAK
>gb|QEE23059_1|ARO_3005060|OXA-837__Cupriavidus_gilardii__TM_#03
ADPDQIQPYGGGKTRFPQWQRDMNLREAIAMSNVPVYQGIARRIGMQRMQTWVDRLDYGN
RQLGK
>gb|QEE23059_1|ARO_3005060|OXA-837__Cupriavidus_gilardii__TM_#04
SGGVHAFALNMDLQREELAPKRMAIARAMMAELGVLPVENGQARKMS
>gb|BAD12078_1|ARO_3002572|AAC(6_)-Iad__Acinetobacter_pittii__TM_#01
MIRKATVQDPPLLARLAMNVWKESSLKELVAEFEQMTKSNDAVAFILFIEDQ
>gb|BAD12078_1|ARO_3002572|AAC(6_)-Iad__Acinetobacter_pittii__TM_#02
KEFRHRGYASELLLKCEDWVKTK
>gb|BAD12078_1|ARO_3002572|AAC(6_)-Iad__Acinetobacter_pittii__TM_#03
KVGFTEANRMICFTKQL
>gb|CAA77936_1|ARO_3004039|Escherichia_coli_emrE__Escherichia_coli__TM_#01
MNPYIYLGGAILA
>gb|CAA77936_1|ARO_3004039|Escherichia_coli_emrE__Escherichia_coli__TM_#02
CYCASFWLLAQTLAY
>gb|CAA77936_1|ARO_3004039|Escherichia_coli_emrE__Escherichia_coli__TM_#03
LSWGFFGQRLDLPAIIGMM
>gb|SOX29786_1|ARO_3004730|tva(A)__Brachyspira_hyodysenteriae__TM_#01
SSYSGYLTSIFKIDYNYLYRFDTLSFGERKRLQIASALYS
>gb|SOX29786_1|ARO_3004730|tva(A)__Brachyspira_hyodysenteriae__TM_#02
HDAKGKIDAARLAGKDSRLATKAKQAKSLYNNTVMEMESLYTKKREVMDMEFIGERYKGK
FLFYLEAGETKINSIVLRHPELIVKKDSRIGIEGVNGAGKTSLLNYIIETMY
>gb|SOX29786_1|ARO_3004730|tva(A)__Brachyspira_hyodysenteriae__TM_#03
ISFNCALIIVSHNRNFIKNAVNTLWSIKIEYNYSILDIKNTI
>gb|ARB93503_1|ARO_3004603|tet(56)__Legionella_longbeachae__TM_#01
IFMDGRIEQ
>gb|AAC32025_1|ARO_3002650|APH(3_)-Vb__Streptomyces_ribosidificus__TM_#01
MESTLRRTYPHHTWHL
>gb|AAC32025_1|ARO_3002650|APH(3_)-Vb__Streptomyces_ribosidificus__TM_#02
HLDGEADRLDWLARHGISVPRVVERGADDTT
>gb|AAC32025_1|ARO_3002650|APH(3_)-Vb__Streptomyces_ribosidificus__TM_#03
AAASEEWPEDERAAVVDAIAEMARTLHELPVSECPFDRRLDVTG
>gb|AAD37403_1|ARO_3003202|TLA-1__Escherichia_coli__TM_#01
KGTDSLKN
>gb|AAD37403_1|ARO_3003202|TLA-1__Escherichia_coli__TM_#02
IEDNFKLNVNEKHH
>gb|AAD37403_1|ARO_3003202|TLA-1__Escherichia_coli__TM_#03
KLDKENIS
>gb|AAD37403_1|ARO_3003202|TLA-1__Escherichia_coli__TM_#04
DKKLFVKKS
>gb|AAD37403_1|ARO_3003202|TLA-1__Escherichia_coli__TM_#05
SDNNGCDILFRFVGGTNKVHNFISKLGV
>gb|AAD37403_1|ARO_3003202|TLA-1__Escherichia_coli__TM_#06
TNWTTPDATVQLLKKFYKNEILSKNSYD
>gb|AAD37403_1|ARO_3003202|TLA-1__Escherichia_coli__TM_#07
SEKSDVNEKIIAEICKSVWDYLVKDGK
>gb|CAA41211_1|ARO_3001400|OXA-5__Pseudomonas_aeruginosa__TM_#01
MKTIAAYLVLVF
>gb|CAA41211_1|ARO_3001400|OXA-5__Pseudomonas_aeruginosa__TM_#02
TALSESISENLAWNKEFSSESVHGVFVLCKSSSNSCTTNNA
>gb|CAA41211_1|ARO_3001400|OXA-5__Pseudomonas_aeruginosa__TM_#03
RQVFKWDGKPRAMKQWEKDL
>gb|CAA41211_1|ARO_3001400|OXA-5__Pseudomonas_aeruginosa__TM_#04
LRGAIQVSAVPVFQQIAR
>gb|CAA41211_1|ARO_3001400|OXA-5__Pseudomonas_aeruginosa__TM_#05
RMQKYLNL
>gb|CAA41211_1|ARO_3001400|OXA-5__Pseudomonas_aeruginosa__TM_#06

KFLESLYLNNLPASKANQLIVKEAIVTEATPEYIVHSKTGYSGVGTES
>gb|CAA41211_1|ARO_3001400|OXA-5__Pseudomonas_aeruginosa__TM_#07
TKIMASEGIIIGG
>gb|AAA83417_1|ARO_3001407|OXA-12__Aeromonas_sobria__TM_#01
MSRLLLSGLL
>gb|AAA83417_1|ARO_3001407|OXA-12__Aeromonas_sobria__TM_#02
GCFLYADGNGQ
>gb|AAA83417_1|ARO_3001407|OXA-12__Aeromonas_sobria__TM_#03
WLPAWRETTTPRRWETY
>gb|AXF35727_1|ARO_3005121|lsa(D)__Lactococcus_garvieae__TM_#01
VHLEIKTNKSFVYFPQTINDKD
>gb|AXF35727_1|ARO_3005121|lsa(D)__Lactococcus_garvieae__TM_#02
EEKRLRDLTEKAQDDKLRKEIGRLKQTAREKEAWSRNLEATKSRKKRGFDSETKRVDKGF
IGRKAANMMQKSKNLEKRMKEDIADKELLLKNWEEVPGLEMSVLESHQKRLLTVENLAAG
FEDF
>gb|AXF35727_1|ARO_3005121|lsa(D)__Lactococcus_garvieae__TM_#03
QVADAQIELIKSSV
>gb|AAG03816_1|ARO_3000379|OprM__Pseudomonas_aeruginosa_PAO1__TM_#01
VTQQQTAKKEDPQA
>gb|AEV91554_1|ARO_3001714|OXA-215__Acinetobacter_haemolyticus__TM_#01
MKLSKLYTLTVLIGFGLSGVACQHIHTPV
>gb|AEV91554_1|ARO_3001714|OXA-215__Acinetobacter_haemolyticus__TM_#02
FNQIENDQTKQ
>gb|AEV91554_1|ARO_3001714|OXA-215__Acinetobacter_haemolyticus__TM_#03
AYKAYGNDLNRAK
>gb|AEV91554_1|ARO_3001714|OXA-215__Acinetobacter_haemolyticus__TM_#04
PKTQQQVIDMLLVDEIR
>gb|AEV91554_1|ARO_3001714|OXA-215__Acinetobacter_haemolyticus__TM_#05
WTGWIEDPNGK
>gb|AAR03105_1|ARO_3001813|OXA-55__Shewanella_algae__TM_#01
MNKGLHRKRLSKRLLLPMLLCLLAQQTQAVAAEQTKVSDVCSEVTAEGWQEVRRWDKLFE
SAGVKGSLLLWDQKRSLGLSNNLSRAAEG
>gb|AAR03105_1|ARO_3001813|OXA-55__Shewanella_algae__TM_#02
ETSRFSWDGKVREIAVWNRDQSFR
>gb|AAR03105_1|ARO_3001813|OXA-55__Shewanella_algae__TM_#03
EIGPKVMAAMVRQLEYGNQDIGGQADSFWLDGQLRITAFQQVDFLRQLHDNK
>gb|AAR03105_1|ARO_3001813|OXA-55__Shewanella_algae__TM_#04
TVYFAVNLDLASASQLPLRQQLVKQVLKQEQLLP
>gb|CAE50926_1|ARO_3002590|AAC(6_)-Iih__Enterococcus_durans__TM_#01
QPMKEVERLLED
>gb|CAE50926_1|ARO_3002590|AAC(6_)-Iih__Enterococcus_durans__TM_#02
MYRKQQVGTRLVSYLEKEIASQGGIVVYLGTDDVEGQTSLAIEEDLFEDTFDKLETIQNR
K
>gb|CAE50926_1|ARO_3002590|AAC(6_)-Iih__Enterococcus_durans__TM_#03
IARKHGSE
>gb|AAF42063_1|ARO_3000810|mtrC__Neisseria_meningitidis_MC58__TM_#01
MAFYAFKAMRAAAL
>gb|AAF42063_1|ARO_3000810|mtrC__Neisseria_meningitidis_MC58__TM_#02
RQIAEGKLLAADGVIAVGIKFDDGTVYPEKGRLLFADPA
>gb|AAF42063_1|ARO_3000810|mtrC__Neisseria_meningitidis_MC58__TM_#03
AQQQGTNWI
>gb|AAF42063_1|ARO_3000810|mtrC__Neisseria_meningitidis_MC58__TM_#04
GITGAKKVTPKEWAS
>gb|AAG29765_2|ARO_3002260|IND-4__Chryseobacterium_indologenes__TM_#01
MRKNVRIFTVLS
>gb|AAM09853_1|ARO_3002957|vanYD__Enterococcus_faecium__TM_#01
MERQNNNENQYGRNRRKDKRKKLFFYRAACAMLGLLIVCVIFGAVYFLRESKDPVLPSKE
NTKTGKDYSFLADGQSEDESPISEPAISNRANAIDLNIIAANAIVMNKDTDALLYQKKRH
GQNCAGQYSKDDYGVDRA
>gb|EEL41021_1|ARO_3003072|mphL__Bacillus_cereus_Rock3-29__TM_#01
ASMEQRMNRVKEQYY
>gb|AAM76670_1|ARO_3002625|ANT(4_)-IIb__Pseudomonas_aeruginosa__TM_#01
MQHTIARWVDRLREEYADAVAILLKGSYARGDAATWSDIDFDVLVSTQDVEDYRTWIEPV
GDRLVHISAAVEWVTGWERDTVDPSSWSYGLPTQETTRLMWAINDETRRRMDRPYKT
>gb|AAM76670_1|ARO_3002625|ANT(4_)-IIb__Pseudomonas_aeruginosa__TM_#02
IARGDDLGVYQSAQTVAKLVPTLLIPINPPVT
>gb|AAM76670_1|ARO_3002625|ANT(4_)-IIb__Pseudomonas_aeruginosa__TM_#03
VGFAADWLTCLGLVEERSARSTAAAAERMVRGVLEMLPTDPDLLGEDIARLMNAGLLEKY
VQQ
>gb|AAA98296_1|ARO_3000250|ErmC__Staphylococcus_epidermidis__TM_#01
MNEKNIKH
>gb|AAA98296_1|ARO_3000250|ErmC__Staphylococcus_epidermidis__TM_#02
HPKPKVNSSLIRLNRKKSRISHKDKQK
>gb|CAC41008_1|ARO_3004041|Klebsiella_pneumoniae_acrA__Klebsiella_pneumoniae__TM_#01
KSPLRSAAGS
>gb|AEP40502_2|ARO_3002975|vanTN__Enterococcus_faecium__TM_#01
YLKIKKFLKKQVKLYVLATLIYLPLAFYSGVITFKTSVIQFFQLIFFE
>gb|AEP40502_2|ARO_3002975|vanTN__Enterococcus_faecium__TM_#02
YCLLQRFTLRQVLLVTFLF
>gb|AEP40502_2|ARO_3002975|vanTN__Enterococcus_faecium__TM_#03
SFHQSEWNMRTSKAKYFLLIASLGMLVESYLLHSFSSPKHDSA
>gb|AEP40502_2|ARO_3002975|vanTN__Enterococcus_faecium__TM_#04
NWQPTRVIADASTISLGIYVLHPYVIAVVHTLAKKITILNNSLIYYLCVSLLTSLIILYV
HSKKKKTTKNQA
>gb|AEP40502_2|ARO_3002975|vanTN__Enterococcus_faecium__TM_#05
KSKILILGYTPSI
>gb|AEP40502_2|ARO_3002975|vanTN__Enterococcus_faecium__TM_#06
YGYLSEHNGDSKINLNLQPILDVQALLVSKKWVAAGEYLGYSLDTKLVSPKLIGIVSIGY
ADGVPRELSHNEFY
>gb|EPF70268_1|ARO_3004478|OXA-665__Acinetobacter_rudis_CIP_110305__TM_#01
MKKNLFACLVLSTALTQVACSTLQTTADPSTQASTAQQSIKSYFDEVQTKGVIVIKQDG
>gb|EPF70268_1|ARO_3004478|OXA-665__Acinetobacter_rudis_CIP_110305__TM_#02
AQEVKRLN
>gb|EPF70268_1|ARO_3004478|OXA-665__Acinetobacter_rudis_CIP_110305__TM_#03
TVQQMILLQ
>gb|EPF70268_1|ARO_3004478|OXA-665__Acinetobacter_rudis_CIP_110305__TM_#04
PKMSGSIRNEITLKALENLGVI
>gb|ENV34035|ARO_3004087|APH(3_)-IX__Acinetobacter_gerneri_DSM_14967___CIP_107464__TM_#01
MINDMKISLPQSLKSFIGNQPLQK
>gb|ENV34035|ARO_3004087|APH(3_)-IX__Acinetobacter_gerneri_DSM_14967___CIP_107464__TM_#02
SFTKNNEKYYLKTTELIYAQTTYSIIREAKILDWLDGKLNVPELVLMDTDHENEYMISKA
VPAKPLQDFTGKSDQFFIDIYTDALAQLQSISIKNCPFISNKKFRLAEAEFFIENGLLDE
LDDDEKDLKLWSSYQNFAEFLDDLKQQNFQEEYVFSHGDLTDSNVFLSHDAQIY

>gb|ENV34035|ARO_3004087|APH(3_)-IX__Acinetobacter_gerneri_DSM_14967___CIP_107464__TM_#03
SLREDCSEDAALQFLNHLAEDDSF
>gb|AAA88552_1|ARO_3002541|AAC(3)-VIIa__Streptomyces_rimosus__TM_#01
EAFEIIGRDMR
>gb|AAZ04368_1|ARO_3002385|BEL-1__Pseudomonas_aeruginosa__TM_#01
MKLLLYPLLLFLVIPAFAQADFEHAISDLEAHNQAKIGVALVSENGNLIQ
>gb|AAZ04368_1|ARO_3002385|BEL-1__Pseudomonas_aeruginosa__TM_#02
AGEENPERKLHYDSAFLEEYAPAAKRYVATGYMTVTEAIQSALQ
>gb|AAZ04368_1|ARO_3002385|BEL-1__Pseudomonas_aeruginosa__TM_#03
PLLTKYFRSLGDKVSR
>gb|AAZ04368_1|ARO_3002385|BEL-1__Pseudomonas_aeruginosa__TM_#04
QTVSKLIFGDTLTYKSKGQLRRLLIGNQTGDKTIRAGLPDSWVTGDKTGSCANGGRNDVA
FFITTAGKKYVLSVYTNAPELQGEERALLIASVAKLARQYVVH
>gb|AAF36802_1|ARO_3002945|vanHF__Paenibacillus_popilliae_ATCC_14706__TM_#01
GHSKEAAANYVSL
>gb|AAC76296_1|ARO_3000656|acrS__Escherichia_coli_str__K-12_substr__MG1655__TM_#01
KIPRQQALLK
>gb|AAC76296_1|ARO_3000656|acrS__Escherichia_coli_str__K-12_substr__MG1655__TM_#02
PQTLREVLQACQQQGCVANNLDLDVVMIIIDGAFSGIVQNWLMNMAG
>gb|AAC76296_1|ARO_3000656|acrS__Escherichia_coli_str__K-12_substr__MG1655__TM_#03
FMPDENITKLIHQTNELSVM
>gb|AAF86641_1|ARO_3002922|vanRC__Enterococcus_gallinarum__TM_#01
VTTFLQNEGFSVQPFYDGTSAIAY
>gb|AAF86641_1|ARO_3002922|vanRC__Enterococcus_gallinarum__TM_#02
HSTASPTVEEYEKDGLILKINSHQCILYGKEVFL
>gb|BAE06006_1|ARO_3003705|mexN__Pseudomonas_aeruginosa__TM_#01
EIIASDPAVQ
>gb|BAH63252_1|ARO_3004583|Klebsiella_pneumoniae_KpnF__Klebsiella_pneumoniae_subsp__pneumoniae_NTUH-K2044__TM_#01
MQQFEWIHAAWLAIAIVLEIIANVFLKFSDGFRRKIYGILSLAAVLGAFSALSQAVK
>gb|AFP97030_1|ARO_3001797|OXA-46__Pseudomonas_aeruginosa__TM_#01
ILLSTFFLTSFV
>gb|AFP97030_1|ARO_3001797|OXA-46__Pseudomonas_aeruginosa__TM_#02
RSDWKKFFSDL
>gb|AFP97030_1|ARO_3001797|OXA-46__Pseudomonas_aeruginosa__TM_#03
RSFAGHNQ
>gb|AFP97030_1|ARO_3001797|OXA-46__Pseudomonas_aeruginosa__TM_#04
YLKQIDYGN
>gb|AFP97030_1|ARO_3001797|OXA-46__Pseudomonas_aeruginosa__TM_#05
FLRKLYRNQLPF
>gb|AFP97030_1|ARO_3001797|OXA-46__Pseudomonas_aeruginosa__TM_#06
VEHQRLVK
>gb|AAA65955_1|ARO_3002942|vanHA__Enterococcus_faecium__TM_#01
NANVSESNAK
>gb|AAF61331_1|ARO_3002966|vanXYC__Enterococcus_gallinarum__TM_#01
MNTLQLINKNHPLKKNQEPPHLVLAPFSDHDVYLQPEVAKQWERLVRATGLEKDIRLVDG
YRTEKEQRRLWEYSLKENGLAYTKQ
>gb|AAF61331_1|ARO_3002966|vanXYC__Enterococcus_gallinarum__TM_#02
VGLKKQED
>gb|AAF61331_1|ARO_3002966|vanXYC__Enterococcus_gallinarum__TM_#03
TAQKWTLEEYHDYLAQTVRQFA
>gb|APB03224_1|ARO_3003994|cpaA__Paenibacillus_sp__LC231__TM_#01
MPLRITAMTETYADQIMQ
>gb|APB03224_1|ARO_3003994|cpaA__Paenibacillus_sp__LC231__TM_#02
KEGQLFGFCCTGSSAQIPIA
>gb|APB03224_1|ARO_3003994|cpaA__Paenibacillus_sp__LC231__TM_#03
STGQCRGKEFLSFVLASIAEFHKRQ
>gb|APB03224_1|ARO_3003994|cpaA__Paenibacillus_sp__LC231__TM_#04
EVATFDYGGTTFITMIKKPGSGL
>gb|ABA70720_1|ARO_3005035|Yrc-1__Yersinia_ruckeri__TM_#01
TFATPQTEKKLSGIVDNV
>gb|AAN17791_1|ARO_3004294|BUT-1__Buttiauxella_agrestis__TM_#01
MCRTLCHVTYGRFSMMKKTLCCALVLSASFSAFAAQKTLSDKQLEEAVNQTLKPMIT
>gb|AAN17791_1|ARO_3004294|BUT-1__Buttiauxella_agrestis__TM_#02
YNMNPQQVKQPTLQKGLEI
>gb|AAN17791_1|ARO_3004294|BUT-1__Buttiauxella_agrestis__TM_#03
ATVINGSDNKVALAASPVTAIEPPVAPVKAS
>gb|AAN17791_1|ARO_3004294|BUT-1__Buttiauxella_agrestis__TM_#04
NTLLNTLQ
>gb|CAA52372_1|ARO_3002656|APH(4)-Ib__Burkholderia_pseudomallei__TM_#01
MLQTSKKKSGHDESWANADAHKWRGERKRDNRKIVLSGTTKLLFVAEEQFQLIPPPSYCV
SLVPKLPSNVTQPLFEYCFAPRILFFYALKKMTQHTKLCKLSSLIWREMWAISSRLQWQC
VCAARRITMRNGGWKFIEMLSCWSDMVHKHESVLISTLPSFINFLVGPFRSAGAEPGGMH
RRVDPPRPLSPALIEAFDGVMQLSGAPSRGVTPTPRGPDALGRITDSRGGSEAGYRFNMC
NRAVPSAALPIGEVLDIGEFSGKRTYLAAVHRARE
>gb|CAA52372_1|ARO_3002656|APH(4)-Ib__Burkholderia_pseudomallei__TM_#02
CTGMAHAIAAADLSHTSGFAPFGPQ
>gb|PLT17746_1|ARO_3003599|OXA-443__Ralstonia_mannitolilytica__TM_#01
LAALATFAHAEHP
>gb|PLT17746_1|ARO_3003599|OXA-443__Ralstonia_mannitolilytica__TM_#02
HTPVEHFR
>gb|PLT17746_1|ARO_3003599|OXA-443__Ralstonia_mannitolilytica__TM_#03
VNRQLPVSA
>gb|PLT17746_1|ARO_3003599|OXA-443__Ralstonia_mannitolilytica__TM_#04
DRTVQTWQVPGGW
>gb|PLT17746_1|ARO_3003599|OXA-443__Ralstonia_mannitolilytica__TM_#05
GKTGTAGPAPGNT
>gb|PLT17746_1|ARO_3003599|OXA-443__Ralstonia_mannitolilytica__TM_#06
TYVFANLIQDDK
>gb|PLT17746_1|ARO_3003599|OXA-443__Ralstonia_mannitolilytica__TM_#07
EPTSGGIRSRDA
>gb|AJO67548_1|ARO_3005006|tet(57)__Providencia_sp__Y14__TM_#01
VIFKDQRKAVIQDAQHGDQPSPIPFMQIIKPIIQ
>gb|AJO67548_1|ARO_3005006|tet(57)__Providencia_sp__Y14__TM_#02
VIFGQTLTM
>gb|AJO67548_1|ARO_3005006|tet(57)__Providencia_sp__Y14__TM_#03
PTIKQQTDN
>gb|AKQ05897_1|ARO_3004589|tet(53)__uncultured_bacterium__TM_#01
EGQALDFRGVAIDIVKGMNIYEKMCNMHKQLEVGRYVDT
>gb|AKQ05897_1|ARO_3004589|tet(53)__uncultured_bacterium__TM_#02
GVCFGVFSIL
>gb|AKQ05897_1|ARO_3004589|tet(53)__uncultured_bacterium__TM_#03
KNEQQQLLRDAFQD
>gb|AKQ05897_1|ARO_3004589|tet(53)__uncultured_bacterium__TM_#04

LVEACHKLGVL
>gb|AEG74318_1|ARO_3002795|QnrS6__Aeromonas_hydrophila__TM_#01
DFRRANLRD
>gb|AEG74318_1|ARO_3002795|QnrS6__Aeromonas_hydrophila__TM_#02
NCKFIEQGDIEGC
>gb|AEG74318_1|ARO_3002795|QnrS6__Aeromonas_hydrophila__TM_#03
DASFQQCQLAMANFSNANCYGIE
>gb|AEG74318_1|ARO_3002795|QnrS6__Aeromonas_hydrophila__TM_#04
CDLKGANF
>gb|AEG74318_1|ARO_3002795|QnrS6__Aeromonas_hydrophila__TM_#05
LSYANMERVCLE
>gb|AEG74318_1|ARO_3002795|QnrS6__Aeromonas_hydrophila__TM_#06
LEALGIVV
>gb|CAA63264_1|ARO_3002012|CMY-1__Klebsiella_pneumoniae__TM_#01
QLDDKASRHAPWLKGS
>gb|CAA63264_1|ARO_3002012|CMY-1__Klebsiella_pneumoniae__TM_#02
LMEQTLLPGLG
>gb|CAA63264_1|ARO_3002012|CMY-1__Klebsiella_pneumoniae__TM_#03
HHTYVNVPKQ
>gb|CAA63264_1|ARO_3002012|CMY-1__Klebsiella_pneumoniae__TM_#04
LEANPTAAPRESG
>gb|AAL55262_1|ARO_3004766|CGA-1__Chryseobacterium_gleum__TM_#01
MKKTTLLFLLISAFSL
>gb|AFO53532_1|ARO_3002883|rgt1438__Streptomyces_sp__WAC1438__TM_#01
RDLGIGSAHPDPVV
>gb|AFO53532_1|ARO_3002883|rgt1438__Streptomyces_sp__WAC1438__TM_#02
VETADRAGRPVSP
>gb|CAI57696_1|ARO_3002614|aadA14__Pasteurella_multocida__TM_#01
MTNKPPESIAEQVSEARSILENHL
>gb|CAI57696_1|ARO_3002614|aadA14__Pasteurella_multocida__TM_#02
NESTRAALMSDLLAVSAFPGTDSKRRALEVTVLTQ
>gb|CAI57696_1|ARO_3002614|aadA14__Pasteurella_multocida__TM_#03
VVDVQRSLLETLTLWTTEADWK
>gb|CAI57696_1|ARO_3002614|aadA14__Pasteurella_multocida__TM_#04
VAAADWALQRLPREIKS
>gb|CAI57696_1|ARO_3002614|aadA14__Pasteurella_multocida__TM_#05
DLRNHIHSSVTAKLQ
>gb|APB03222_1|ARO_3003992|rphB__Paenibacillus_sp__LC231__TM_#01
AMLNRLTMLS
>gb|APB03222_1|ARO_3003992|rphB__Paenibacillus_sp__LC231__TM_#02
IAENEQTPGKNNISSAGYQTPIDHDPTIVSDLME
>gb|APB03222_1|ARO_3003992|rphB__Paenibacillus_sp__LC231__TM_#03
KLLTFDGGQQAQD
>gb|AAQ76277_1|ARO_3001796|OXA-50__Pseudomonas_aeruginosa__TM_#01
MRPLLFSALLLLSG
>gb|AAQ76277_1|ARO_3001796|OXA-50__Pseudomonas_aeruginosa__TM_#02
QASEWNDS
>gb|AAQ76277_1|ARO_3001796|OXA-50__Pseudomonas_aeruginosa__TM_#03
AVDKLFGAAGVKGTFVLYDVQRQ
>gb|AAQ76277_1|ARO_3001796|OXA-50__Pseudomonas_aeruginosa__TM_#04
YVGHDRERAETRFVPASTYKVAN
>gb|AAQ76277_1|ARO_3001796|OXA-50__Pseudomonas_aeruginosa__TM_#05
RFKAWEHDMSLR
>gb|AAQ76277_1|ARO_3001796|OXA-50__Pseudomonas_aeruginosa__TM_#06
LGYGNAEIGQ
>gb|AAQ76277_1|ARO_3001796|OXA-50__Pseudomonas_aeruginosa__TM_#07
FLLRLAQGELPFPAPVQSTVRAMTLLES
>gb|ASF80997_1|ARO_3005164|dfrA35__Escherichia_coli__TM_#01
MISIVVAKSA
>gb|ASF80997_1|ARO_3005164|dfrA35__Escherichia_coli__TM_#02
VIISKQPVPIEWASKVVWVNSIQQAMDYVRGLDGMIKTFIIGGSEIYRQFISLVDQVYLT
EVGAEIEGDATFQPLDEHEWTLKTWWVVPDQSSKDQFRYQRKLYVRKVLDE
>gb|AGV28567_1|ARO_3000559|adeN__Acinetobacter_baumannii__TM_#01
MHDPVLESHHLVCEKPQTRRGI
>gb|AGV28567_1|ARO_3000559|adeN__Acinetobacter_baumannii__TM_#02
LDVQNTIAQALLISHQSGEIT
>gb|QEQ43476_1|ARO_3005061|AAC(3)-IVb__Cupriavidus_gilardii__TM_#01
MVTQLRALGVPPGAVLLVHASFRSIRPVQGGPQGLIEALREAA
>gb|QEQ43476_1|ARO_3005061|AAC(3)-IVb__Cupriavidus_gilardii__TM_#02
GDDDDAPFDPAATPAAADLGAVADAFWRLPDVVRSH
>gb|QEQ43476_1|ARO_3005061|AAC(3)-IVb__Cupriavidus_gilardii__TM_#03
GVPHHCTVLRDGKPARIDYLENDHCCQRFDLVDGWLKEKGLQREGPVGNAGARLMRARDI
VDVVRQQLARDPLVFLHPPQVGCEECDAARRSVSTK
>gb|ABH10964_1|ARO_3002830|vgaALC__Staphylococcus_haemolyticus__TM_#01
EVTRNYIRQALD
>gb|ABH10964_1|ARO_3002830|vgaALC__Staphylococcus_haemolyticus__TM_#02
VEQKELERHREELE
>gb|ABH10964_1|ARO_3002830|vgaALC__Staphylococcus_haemolyticus__TM_#03
KQKKLRKTVKSLETRLEKLE
>gb|ABH10964_1|ARO_3002830|vgaALC__Staphylococcus_haemolyticus__TM_#04
VEKRNELPPLKMDLVNLESVKNRTIIRGEDVSGTIEGRVLWKAKSFSIRGGDKMAIIGSN
GTGKTTFIKKIVHGNP
>gb|ABH10964_1|ARO_3002830|vgaALC__Staphylococcus_haemolyticus__TM_#05
DGTYEQFKQAEKP
>gb|ABH10964_1|ARO_3002830|vgaALC__Staphylococcus_haemolyticus__TM_#06
NLINEKRNLDK
>gb|AFK13828_1|ARO_3000823|ramA__Enterobacter_cloacae__TM_#01
NLHQPLRIEEIARH
>gb|AFK13828_1|ARO_3000823|ramA__Enterobacter_cloacae__TM_#02
LLAARDLRESDA
>gb|AEQ93536_1|ARO_3004468|CrpP__Pseudomonas_aeruginosa__TM_#01
MSKKATGTDKLDRRHFNDPHRTVRAIGAEAARKGLRVFDCPYSHPAMRASWLKGFAQEQQ
QQLDF
>gb|BAN78519_1|ARO_3004638|AAC(6_)-Iag__Pseudomonas_aeruginosa__TM_#01
MSKLGPTKPAPSMANTPVGNVVPCKTPDHPGWLELRLQLWPDGSTEEFLPEMAAACAEPD
RFGQFLFLSPGGLAEGLVEV
>gb|BAN78519_1|ARO_3004638|AAC(6_)-Iag__Pseudomonas_aeruginosa__TM_#02
VFVVPASRGLGIARALVAAAEGWARDRGCTEFASDAEVSNVG
>gb|AAB28795_1|ARO_3000178|tet(K)__Staphylococcus_aureus__TM_#01
SYLLILPMITIVTIPFLIKVMVPGKSTKNTLDIVGIVLMSISIICFMLFTTNYNWTFLIL
FTIFFVIFIKHISRVSNPFINPKLGKNIPFMLGLFSGGLIFS
>gb|AAB28795_1|ARO_3000178|tet(K)__Staphylococcus_aureus__TM_#02
TIYHVNVATIGNSVIFPGTMSVIVFGYFGGFLVDRKGSLFVFILGSLSISISFLTIAFFV

EFSMWLTTFM
>gb|AAB28795_1|ARO_3000178|tet(K)__Staphylococcus_aureus__TM_#03
KIVSSSLSE
>gb|AAB28795_1|ARO_3000178|tet(K)__Staphylococcus_aureus__TM_#04
VLEFINYSSGVYSNILVAMAILIILCCLLTIIVFKRSEKQFE
>gb|AAD51344_1|ARO_3003051|smeA__Stenotrophomonas_maltophilia__TM_#01
DRTDAPAMPEVGVIIASAQPLALQQTLPGRAVPFEISEVRPQIGGLIRQRLFTEGQ
>gb|AAD51344_1|ARO_3003051|smeA__Stenotrophomonas_maltophilia__TM_#02
TVLSAQPKAERTRALVSMDAASKQDADDATSALKQAQAN
>gb|AAD51344_1|ARO_3003051|smeA__Stenotrophomonas_maltophilia__TM_#03
LVKAIDGKAQVKVLLEDGSTYAHEGTLEFVGSAVDPGTGNVK
>gb|AAD51344_1|ARO_3003051|smeA__Stenotrophomonas_maltophilia__TM_#04
ARALLVPQKAVVRNERGEPLLRLLDAKDHVVERRVSTGQVVGNQWQITSGLKAGERVIVS
NGSAVSLGQQVKAVAPTTAQLAAMPAVDPNGNTDEKSH
>gb|WP_082741435_1|ARO_3004569|ICR-Mo__Moraxella_osloensis__TM_#01
MVHLDKVSNR
>gb|WP_082741435_1|ARO_3004569|ICR-Mo__Moraxella_osloensis__TM_#02
IKRSILQRG
>gb|WP_082741435_1|ARO_3004569|ICR-Mo__Moraxella_osloensis__TM_#03
LNGYNRTTFPQMAATAGVTNFNQ
>gb|WP_082741435_1|ARO_3004569|ICR-Mo__Moraxella_osloensis__TM_#04
LPAADFVDYKTARNNTMCNT
>gb|WP_082741435_1|ARO_3004569|ICR-Mo__Moraxella_osloensis__TM_#05
LAKCDPQSVINAFDNALLATDDFLAKTVNWLDKYDSTH
>gb|AAA98096_1|ARO_3000595|ErmT__Plasmid_pGT633__TM_#01
EILRNVHLNT
>gb|AAA98096_1|ARO_3000595|ErmT__Plasmid_pGT633__TM_#02
IKLFSKNQFYQALKYAR
>gb|AAN85115_1|ARO_3004642|dfrA3b__Citrobacter_freundii__Partial_TM_#01
GGGLPWDTLKDDLQFFKRLTEGTDLVMGASTYRTLPLLPTNNRQFIVVSNTEEPSLNVHV
VSPEHFKAFLSKTSRNLTIIGGSSLLTVDILSKMDKIIMTTVYGSFDADVYLPTEVVSYV
TGKASNATLFNNSDAKMAVYYG
>gb|CAA53189|ARO_3000521|mupA__Staphylococcus_aureus__TM_#01
SKIRVENEYYILATDLINSIITEKYEIIDTFSGSNLINLKYIPPFESDGLV
>gb|CAA53189|ARO_3000521|mupA__Staphylococcus_aureus__TM_#02
LVLERDLDFLNVITREGVYNDRFPELVGNKAKNSDIEIIKLLSKKQ
>gb|CAA53189|ARO_3000521|mupA__Staphylococcus_aureus__TM_#03
HYNLTGKSVHLQDYPQYKESFINQALEDEMHTVIKIVELSRQARKNADLKIKQPLSKMVI
KPNSQLNLSFLPNYYSIIKDELNIKNIELTDNINDYITYELKLNFSSVGPKLGNKTKNIQ
TLIDSLSEYDKKSLIESNNFKSLSSDAELTKDDFIIKTLPKDSYQLSEDNDCVILLDKNL
SPELIREGHA
>gb|CAA53189|ARO_3000521|mupA__Staphylococcus_aureus__TM_#04
KKNLPINQRIDIY
>gb|AAK01167_1|ARO_3002819|msrC__Enterococcus_faecium__TM_#01
MENLAVNITNLQVRFGNQLELSIDSLRVYQ
>gb|AAK01167_1|ARO_3002819|msrC__Enterococcus_faecium__TM_#02
IPNQGKIQTEITFNYLPQLTYLAEAKDLNLELASHFQLKLEETSERKWSGGEERKIELIR
LLSSYEQG
>gb|AAK01167_1|ARO_3002819|msrC__Enterococcus_faecium__TM_#03
KVWEVKDGEIREFPE
>gb|AAK01167_1|ARO_3002819|msrC__Enterococcus_faecium__TM_#04
NTKTLERRLQKIGETTKPQQMKQIRFPVPKSLELHNRYPIMGQNIQLERSGRTLLINS
>gb|AAL27443_1|ARO_3002967|vanXYE__Enterococcus_faecalis__TM_#01
MKKNYLRLINENNEIKDSERPSHLVQAPFAQTNILVDPMVAIQLEKLIKTTGLDSQIITI
DGYRSKETQQALWDETIQEKGLEFAH
>gb|AAL27443_1|ARO_3002967|vanXYE__Enterococcus_faecalis__TM_#02
SIMVQQNWVLEEYIEFIESIRGTAYEA
>gb|CAA03986_1|ARO_3000343|tap__Mycolicibacterium_fortuitum__TM_#01
CCSILAISVLRLEGAGAPDRSVLTEAVL
>gb|CAA03986_1|ARO_3000343|tap__Mycolicibacterium_fortuitum__TM_#02
RLRELDLASKP
>gb|ABS29619_1|ARO_3000841|CGB-1_beta-lactamase__Chryseobacterium_gleum__TM_#01
LANAQDTQVRDFVIEPQIQ
>gb|ABS29619_1|ARO_3000841|CGB-1_beta-lactamase__Chryseobacterium_gleum__TM_#02
KAAADLGYTGEANVAQWPK
>gb|AAC32026_1|ARO_3002823|ErmH__Streptomyces_thermotolerans__TM_#01
MAALLKRILRRRMAEKRSGRGRMAAARTTGAQSRKTAQRSGRSEADRRRRVHGQNFLVDR
ETVQRFVRFADPDPGEVVLEVGAGN
>gb|AAC32026_1|ARO_3002823|ErmH__Streptomyces_thermotolerans__TM_#02
RHFADRLREATAEDPRIEVVAGDFLKTSQPKVPFSVVGNIPFGN
>gb|AAC32026_1|ARO_3002823|ErmH__Streptomyces_thermotolerans__TM_#03
ATWPEVEWRMGERISRRWFRPVPAVDSAVLRLERRPVPLIPPGLMHDFRDLVETGFTGKG
GSLDASLRRRFPARRVAAGFRRARLEQGVVVAYVTPGQWITLFEELHGR
>gb|CAA71441_1|ARO_3003095|imiS__Aeromonas_veronii__TM_#01
ASFWGGSV
>gb|CAA71441_1|ARO_3003095|imiS__Aeromonas_veronii__TM_#02
KSDWAEIVA
>gb|CAA71441_1|ARO_3003095|imiS__Aeromonas_veronii__TM_#03
PDGIFVYFPD
>gb|CAA71441_1|ARO_3003095|imiS__Aeromonas_veronii__TM_#04
ERLKAMKLPIK
>gb|CAA71441_1|ARO_3003095|imiS__Aeromonas_veronii__TM_#05
GGHDSPLHGPELIDHY
>gb|CAA37699_1|ARO_3003565|r39_beta-lactamase__Actinomadura_sp__R39__TM_#01
MLFPTARRTGFAALAALALVPAAACSGSAAPAEAEPASAEVTAEDLSG
>gb|CAA37699_1|ARO_3003565|r39_beta-lactamase__Actinomadura_sp__R39__TM_#02
EDMRELGDDVISADRI
>gb|CAA37699_1|ARO_3003565|r39_beta-lactamase__Actinomadura_sp__R39__TM_#03
EEGPRDVLTEMLLN
>gb|CAA37699_1|ARO_3003565|r39_beta-lactamase__Actinomadura_sp__R39__TM_#04
DDPIVIAVMSTREQEDAEFDNALVSGATEVVVEALAP
>gb|AAA26549_1|ARO_3002549|AAC(6_)-Ic__Serratia_marcescens__TM_#01
MIVICDHDNLDAWLALRTALWPSGSPEDHRAEMREILASPHHTAFMARGLDGAF
>gb|AAA26549_1|ARO_3002549|AAC(6_)-Ic__Serratia_marcescens__TM_#02
AERARRQGWAARLIAQVQEWAKQQGCSELASDTDIANLDSQRL
>gb|AET10445_1|ARO_3004033|tetB(46)__Streptococcus_australis__TM_#01
MKVLKRLLSRITLYPTVFLAGFICLLLATIFSELSPFLLQKMIDGPLTALTHGGGQGDLL
QMGGFYLLVLSLGQLISYMGNRILLHGSNQVTAS
>gb|AET10445_1|ARO_3004033|tetB(46)__Streptococcus_australis__TM_#02
ILVIYLVRFLFILGILFYLS
>gb|AET10445_1|ARO_3004033|tetB(46)__Streptococcus_australis__TM_#03
YLYKVMTDQ
>gb|AET10445_1|ARO_3004033|tetB(46)__Streptococcus_australis__TM_#04

```
HQEPGVVEEFEATTQKMLGANDRILLADSIASWTLTELLKFLVIAGILTIAGISFLQGNI
GVTAGFLFININY
>gb|AET10445_1|ARO_3004033|tetB(46)__Streptococcus_australis__TM_#05
HIDRNAVKDALKKVGAWPF
>gb|AAL86999_1|ARO_3004782|ERP-1__Erwinia_persicina__TM_#01
MTILLQRRQLLVAGAALALTASLTPLNVFAAGDSLQRQLAALETEVNGRIGLSLIDSASQ
QA
>gb|AAL86999_1|ARO_3004782|ERP-1__Erwinia_persicina__TM_#02
SESQPALMQQTLHWTPADHLSYMPVTAKHP
>gb|AAL86999_1|ARO_3004782|ERP-1__Erwinia_persicina__TM_#03
TLLGGPASVTRL
>gb|AAL86999_1|ARO_3004782|ERP-1__Erwinia_persicina__TM_#04
SHSVQQLLVKSGLQTAQQQ
>gb|AAL86999_1|ARO_3004782|ERP-1__Erwinia_persicina__TM_#05
KNAIAAALPAGWEIG
>gb|AAL86999_1|ARO_3004782|ERP-1__Erwinia_persicina__TM_#06
PGKAPLILAIYFTQHAPEAKSRQDVLAKAAAIALKSVI
>gb|ENU91137|ARO_3004090|ANT(3__)-IIb__Acinetobacter_sp__NIPH_758__TM_#01
MSEQFQLQQLQEYLHALF
>gb|ENU91137|ARO_3004090|ANT(3__)-IIb__Acinetobacter_sp__NIPH_758__TM_#02
HVQRQQLAQALLTLSHPIGGLQRALEVTILLKEEVISGR
>gb|ENU91137|ARO_3004090|ANT(3__)-IIb__Acinetobacter_sp__NIPH_758__TM_#03
VDGGELSAQN
>gb|ENU91137|ARO_3004090|ANT(3__)-IIb__Acinetobacter_sp__NIPH_758__TM_#04
SLGEIVSKSHAAQWVIAQLEEKD
>gb|ENU91137|ARO_3004090|ANT(3__)-IIb__Acinetobacter_sp__NIPH_758__TM_#05
MTKQDWPSQHQVLQPIVNFLSQHIETFFDKKGLKIKQ
>gb|AJP77059|ARO_3003670|PEDO-1__Pedobacter_sp__SI-33__TM_#01
MKIKFIFCLLLFPLFVFSQNVQEPTD
>gb|AJP77059|ARO_3003670|PEDO-1__Pedobacter_sp__SI-33__TM_#02
RSSAAQIKKNVEL
>gb|AJP77059|ARO_3003670|PEDO-1__Pedobacter_sp__SI-33__TM_#03
SVMKDGGRTDYALGNGKTTYAPIIPDRLLKDGDQIQ
>gb|AJP77059|ARO_3003670|PEDO-1__Pedobacter_sp__SI-33__TM_#04
EKDYAYTL
>gb|AJP77059|ARO_3003670|PEDO-1__Pedobacter_sp__SI-33__TM_#05
CLHEKHKPGDKYNPTAFIDRAGYDKVLNSLQLEFDKKIGKK
>gb|APB03225_1|ARO_3003990|VgbC__Paenibacillus_sp__LC231__TM_#01
NSSPYGLAEGPDGA
>gb|APB03225_1|ARO_3003990|VgbC__Paenibacillus_sp__LC231__TM_#02
CIGRITVDGEITEYRIPTEQ
>gb|APB03225_1|ARO_3003990|VgbC__Paenibacillus_sp__LC231__TM_#03
QIGRITTAGEITEFKLPSG
>gb|APB03225_1|ARO_3003990|VgbC__Paenibacillus_sp__LC231__TM_#04
CTGDITEYPIPTPSAEPHGITVDSGGEVWFAEECDQIGRFTIQY
>gb|AMP42492_1|ARO_3004441|tet(59)__uncultured_bacterium_IN-14__TM_#01
LSFLVIMLIFKDNKIKNTEKNTTETAENSRPFLQVIKPVILLLFIFFMT
>gb|AMP42492_1|ARO_3004441|tet(59)__uncultured_bacterium_IN-14__TM_#02
FIFGQTLASWDGWIWMIGAIMYVL
>gb|AMP42492_1|ARO_3004441|tet(59)__uncultured_bacterium_IN-14__TM_#03
TKKIVKIAKLPAS
>gb|ACX92987_1|ARO_3002845|vatH__Enterococcus_faecium__TM_#01
MAEKLKGPNSNEMYPIAGNKSVQFVKPSL
>gb|ACX92987_1|ARO_3002845|vatH__Enterococcus_faecium__TM_#02
QELLKIKWWDFEDQVISDNIDAILSLDVEALNNISKEND
>gb|ABG86067_1|ARO_3003773|Clostridium_perfringens_mprF__Clostridium_perfringens_SM101__TM_#01
LLIISGIYPSIFYKIKFLDNIYSLSFLRFSHRASILIGLMLIMTSKEVFFKVKRAYYVTL
TLLIVGGAFAFIKDLDYKEGIFILGVIILLILSKKSFYRKSIPIKVTKLSGILIVLSIVM
IIFASFIHKFNIHFSKNYKYYIDFFHSTKGYLRIALFTYISFIIFVIIWYLTMPKIEDDE
RYMDADLEKVSKFFKEIDYGTIFSHLVYLKDKKVFWANEGESLIMYSKYKDKIIVLGDPI
ATKENLYSCIEEFQAFTNLYGYDVVFYEIEEKNFSTYHDAGYYFFKLGEEARIDLEEFNL
>gb|QHW12307_1|ARO_3005089|mef(D)__Macrococcus_canis__TM_#01
YEINISFVILILAI
>gb|QHW12307_1|ARO_3005089|mef(D)__Macrococcus_canis__TM_#02
SGILLFIYIPKIQNQNQQSIDSKG
>gb|QHW12307_1|ARO_3005089|mef(D)__Macrococcus_canis__TM_#03
FLGILGDKFNKINTMTTGILLMGIALFLTGILSPSLFYFFVVLAGLVGF
>gb|QHW12307_1|ARO_3005089|mef(D)__Macrococcus_canis__TM_#04
TSISLLATPLGYVIAGLLIEITNVSTLFSIIGVLIIFNGIIINRLK
>gb|AAP15294_1|ARO_3002675|catB2__Pasteurella_multocida__TM_#01
NYFESPFKGK
>gb|AAP15294_1|ARO_3002675|catB2__Pasteurella_multocida__TM_#02
EISMLLDM
>gb|AAP15294_1|ARO_3002675|catB2__Pasteurella_multocida__TM_#03
WWDWPLEQIKEAMP
>gb|AAA65956_1|ARO_3000010|vanA__Enterococcus_faecium__TM_#01
KKNHEYEINH
>gb|ENU37733|ARO_3004091|ANT(3__)-IIc__Acinetobacter_parvus_DSM_16617___CIP_108168__TM_#01
FGPSLTQWSV
>gb|AVZ47168_1|ARO_3004693|MCR-3_12__Escherichia_coli__TM_#01
MPSLIKIKIVPL
>gb|AVZ47168_1|ARO_3004693|MCR-3_12__Escherichia_coli__TM_#02
FFLALYFAF
>gb|AVZ47168_1|ARO_3004693|MCR-3_12__Escherichia_coli__TM_#03
LNWRGVLHFYEILYKLE
>gb|AVZ47168_1|ARO_3004693|MCR-3_12__Escherichia_coli__TM_#04
GFIPAILLFFV
>gb|AVZ47168_1|ARO_3004693|MCR-3_12__Escherichia_coli__TM_#05
FAPDDQTRVPMQVWMSPGF
>gb|AVZ47168_1|ARO_3004693|MCR-3_12__Escherichia_coli__TM_#06
RYSHDNIFSSVLGIWDVKT
>gb|ABX30738_1|ARO_3000621|PC1_beta-lactamase_(blaZ)__Staphylococcus_aureus_subsp__aureus_USA300_TCH959__TM_#01
ELNDLEKKYNAHIGVYALDTKSGKEVKFNSDKRFAYASTSKAINSAILLEQVPYNKLNKK
VH
>gb|ABX30738_1|ARO_3000621|PC1_beta-lactamase_(blaZ)__Staphylococcus_aureus_subsp__aureus_USA300_TCH959__TM_#02
KNKNFLLDLM
>gb|AAV80464_1|ARO_3000565|tet(38)__Staphylococcus_aureus__TM_#01
IGALIAIIVFALYIKNAQ
>gb|AAV80464_1|ARO_3000565|tet(38)__Staphylococcus_aureus__TM_#02
EYLNSKQAIITAII
>gb|AAV80464_1|ARO_3000565|tet(38)__Staphylococcus_aureus__TM_#03
DALSSHFGIILIILGLMSIVGLVLFVILNRWTQSEK
>gb|AJP77058|ARO_3003719|MSI-OXA__Massilia_sp__SB1-3__TM_#01
```

MSHTFIIDRLGASMIKQILAALLLISPLLAQAAEWKESAQVARLFKQEGVSGTFVVY
>gb|AJP77058|ARO_3003719|MSI-OXA__Massilia_sp__SB1-3__TM_#02
HGAVANVDEVVPYGGKPVARPEWAR
>gb|AJP77058|ARO_3003719|MSI-OXA__Massilia_sp__SB1-3__TM_#03
EFLTKLAQGTLPFTNVAMAAVREISRQDGAPELYAKTGWGSRPGEA
>gb|AJP77058|ARO_3003719|MSI-OXA__Massilia_sp__SB1-3__TM_#04
KKDGKLYAFALNMDLPDGAQDKRVSLAKAALRELGLL
>gb|AJI75632_1|ARO_3004795|FPH-1__Francisella_philomiragia_subsp__philomiragia_ATCC_25015__TM_#01
MKRLLFTSLY
>gb|AJI75632_1|ARO_3004795|FPH-1__Francisella_philomiragia_subsp__philomiragia_ATCC_25015__TM_#02
HKGFLDKKILITQDDIGTLGYAPVTGKNIG
>gb|AJI75632_1|ARO_3004795|FPH-1__Francisella_philomiragia_subsp__philomiragia_ATCC_25015__TM_#03
LVAKLGDKDTIIKN
>gb|AJI75632_1|ARO_3004795|FPH-1__Francisella_philomiragia_subsp__philomiragia_ATCC_25015__TM_#04
GYLQKNNTGANRIAYS
>gb|AJI75632_1|ARO_3004795|FPH-1__Francisella_philomiragia_subsp__philomiragia_ATCC_25015__TM_#05
PPFALSILYTNPSDVKAPSNEKIIQQASKLVSESIAKKDS
>gb|BAQ22025_1|ARO_3003200|AAC(6_)-Ian__Serratia_marcescens__TM_#01
MLVQQGRLAIRALQKSDAPVMLRWLQDERVLEFYEGRDKHFDLQTVIEVFIEDQGETTPC
LVLLDDKPLGYVQFYPLDSEDKQALELPVEDV
>gb|BAQ22025_1|ARO_3003200|AAC(6_)-Ian__Serratia_marcescens__TM_#02
GLGLGTELVSLVRDYLITDKAAQRLVLDPQSRNS
>gb|BAQ22025_1|ARO_3003200|AAC(6_)-Ian__Serratia_marcescens__TM_#03
RLLPAHEMHEGQLQDCWLMQYYPARSSLMASSRPKI
>gb|CAH35803_1|ARO_3002982|amrA__Burkholderia_pseudomallei_K96243__TM_#01
MKYEWARTRRLSAALAVAAFVAAGCGKHESEHDAAAPREASVVTVKK
>gb|CAH35803_1|ARO_3002982|amrA__Burkholderia_pseudomallei_K96243__TM_#02
FKAARDAAAGALEKARAAH
>gb|CAH35803_1|ARO_3002982|amrA__Burkholderia_pseudomallei_K96243__TM_#03
KSGRAAGIAQQDVEVTLVRPDGSTYAR
>gb|CAH35803_1|ARO_3002982|amrA__Burkholderia_pseudomallei_K96243__TM_#04
SATVKVVGQ
>gb|CAH35803_1|ARO_3002982|amrA__Burkholderia_pseudomallei_K96243__TM_#05
VDAAQFEAGTTVKALERGAAAQPASGAAAASAPGRRST
>gb|ACJ63262_1|ARO_3003107|mefB__Escherichia_coli__TM_#01
LADGLVAVSSIILGAAFLLVETPPIWFIYI
>gb|ACJ63262_1|ARO_3003107|mefB__Escherichia_coli__TM_#02
FVPADMLTKAGGWGNMIQSISNMMGPVLGAALMSFLPISSIMIVDILGAAFAIVCLLFVI
IPDITQTNEKMSVLSDMKQGFIAMKANKPLMAVFS
>gb|ACJ63262_1|ARO_3003107|mefB__Escherichia_coli__TM_#03
FMASLAIGLMGLATLISGALPTSGFWIFVICCFFLGASGTFMNVPV
>gb|AAG07592_1|ARO_3000806|mexG__Pseudomonas_aeruginosa_PAO1__TM_#01
MQRFIDNSLESNWLWLTARICLALMFVASGLAKLFDYQAS
>gb|AAG07592_1|ARO_3000806|mexG__Pseudomonas_aeruginosa_PAO1__TM_#02
ALVLLDRK
>gb|AAG07592_1|ARO_3000806|mexG__Pseudomonas_aeruginosa_PAO1__TM_#03
SKTGVEAKLA
>gb|AAG07592_1|ARO_3000806|mexG__Pseudomonas_aeruginosa_PAO1__TM_#04
ATAIASAQRQRLRQDVSVAATYQKA
>gb|ADV91011_1|ARO_3000245|RbpA__Mycolicibacterium_smegmatis_MC2_155__TM_#01
LCRNGLEGTLIEGDVPEP
>gb|ADV91011_1|ARO_3000245|RbpA__Mycolicibacterium_smegmatis_MC2_155__TM_#02
DLIKAKRRGTGS
>gb|AAA76822_1|ARO_3002654|APH(3_)-VIIa__Campylobacter_jejuni__TM_#01
MKYIDEIQILGKCSEGMSPAEVYKCQLKNTVCYLKKIDDIFSKTTYSVKREAEMMMWLSD
KLKVPDVIEYGVREHSEYLIMSELRGKHIDCFIDHPIKYIECLVNALHQLQAIDIRNCPF
SSKIDVRLKELKYLLDNRIADIDVSNWEDTTEFDDPMTLYQWLCENQPQEELCLSHGDMS
ANFFVSHDGIYFYDLARCGVADKWLDIAFCVREIREYYPDSDYEKFFFNMLGLEPDYKKI
NYYILLDEMF
>gb|AAD12753_1|ARO_3000177|tet(J)__Proteus_mirabilis__TM_#01
FIFSLLCFQETQTTKISTEISALNQDTAPHSTTG
>gb|AAD12753_1|ARO_3000177|tet(J)__Proteus_mirabilis__TM_#02
VRFAWHTTEVGLSLAF
>gb|AAD12753_1|ARO_3000177|tet(J)__Proteus_mirabilis__TM_#03
RNTVIISMSIDAF
>gb|AAD12753_1|ARO_3000177|tet(J)__Proteus_mirabilis__TM_#04
FIGAMLYSGLLVASYFKQKSPILKKFPS
>gb|CAA31895_1|ARO_3002533|AAC(3)-IIa__Plasmid_pWP113a__TM_#01
KLGRHREGVVGFAQC
>gb|AAT90847_1|ARO_3003557|SFB-1__Shewanella_frigidimarina__TM_#01
MISAPSFAHENEQQTDQSNTDAVKKPQQQPTELFLSPLVPDVYLHQSYKQ
>gb|AAT90847_1|ARO_3003557|SFB-1__Shewanella_frigidimarina__TM_#02
WVSDKTQRLLTANKLSTASHTFRTKQHTLQQQ
>gb|AAT90847_1|ARO_3003557|SFB-1__Shewanella_frigidimarina__TM_#03
SHTIDLLTQ
>gb|APB03218_1|ARO_3003984|BahA__Paenibacillus_sp__LC231__TM_#01
MEIHKEKEK
>gb|APB03218_1|ARO_3003984|BahA__Paenibacillus_sp__LC231__TM_#02
PLSPGWKGASLGLGATG
>gb|APB03218_1|ARO_3003984|BahA__Paenibacillus_sp__LC231__TM_#03
VGKFIVGTFLFLFAAALISGLASM
>gb|APB03218_1|ARO_3003984|BahA__Paenibacillus_sp__LC231__TM_#04
IMFLFCFIVQPGVSAVFIVSLAIVLSLFGALAYKFAAGSYKQVSKTRKIGAMACLSLITI
AIGAGSFWLIRAGDDAAPDITLK
>gb|APB03218_1|ARO_3003984|BahA__Paenibacillus_sp__LC231__TM_#05
HEDIDLQSTTLPGGSLRGENLQQWKEEKVKTKWGEAD
>gb|APB03218_1|ARO_3003984|BahA__Paenibacillus_sp__LC231__TM_#06
NGVETGRDSSIVFSLADARKTEESG
>gb|APB03215_1|ARO_3003981|tetB(48)__Paenibacillus_sp__LC231__TM_#01
MSSSMIQPGT
>gb|AAB58555_1|ARO_3001413|OXA-18__Pseudomonas_aeruginosa__TM_#01
MQRSLSMSGKRHFIFAVSFVISTVCLTFSPANAAQKLSCTLVIDEASGDLLHREGS
>gb|AAB58555_1|ARO_3001413|OXA-18__Pseudomonas_aeruginosa__TM_#02
SQQKPTDPTIWL
>gb|AAB58555_1|ARO_3001413|OXA-18__Pseudomonas_aeruginosa__TM_#03
SRFSDYVQR
>gb|AAB58555_1|ARO_3001413|OXA-18__Pseudomonas_aeruginosa__TM_#04
DALEMTKAVVPHFEAGDWDVQGKTGTGSLSDAKGGKAPIGWFIGWATRDDRRVVFARLTV
GARKGEQPAGPAARDEFLNTLPALSENF
>gb|CDO13836_1|ARO_3005044|OmpA__Klebsiella_pneumoniae__TM_#01
MRSFVTSWR
>gb|AAC60781_1|ARO_3003048|rosA__Yersinia_enterocolitica_(type_O_8)__TM_#01
MTDRSETELPPSVNTQPFDNTK

>gb|AAC60781_1|ARO_3003048|rosA__Yersinia_enterocolitica_(type_O_8)__TM_#02
ARLPVAATVWLSLFLGGRQFRQR
>gb|AAC60781_1|ARO_3003048|rosA__Yersinia_enterocolitica_(type_O_8)__TM_#03
YGKVVKVSSAKILPKKTVIS
>gb|BAA15221_2|ARO_3000263|marA__Escherichia_coli_str__K-12_substr__W3110__TM_#01
DWIEDNLESPLSLEK
>gb|BAA15221_2|ARO_3000263|marA__Escherichia_coli_str__K-12_substr__W3110__TM_#02
HSLGQYIRSRKMTEIAQKLKESNEPILY
>gb|BAA15221_2|ARO_3000263|marA__Escherichia_coli_str__K-12_substr__W3110__TM_#03
KNYFDVPPHKYRMTNMQGESRFLHPLNHYNS
>gb|CAA62365_1|ARO_3002645|APH(3_)-IIb__Pseudomonas_aeruginosa__TM_#01
MHDAATSMPPQAPSTWADYLAGYRWRGQGEGCSAATVHRLEAARRPTLFVKQEVLSAHAE
LPA
>gb|CAA62365_1|ARO_3002645|APH(3_)-IIb__Pseudomonas_aeruginosa__TM_#02
SDGRQWLLMSAMPGDTLSALAQRDELEPERLVRLVAAA
>gb|CAA62365_1|ARO_3002645|APH(3_)-IIb__Pseudomonas_aeruginosa__TM_#03
RLERRLDTVRQRVEAGLVDEADFDDDHRGRSATELYRLLLDR
>gb|CAA62365_1|ARO_3002645|APH(3_)-IIb__Pseudomonas_aeruginosa__TM_#04
AAWAEAFLVE
>gb|AAX14802_1|ARO_3000781|adeJ__Acinetobacter_baumannii__TM_#01
VIDTMTNFFMN
>gb|AAF86642_1|ARO_3002933|vanSC__Enterococcus_gallinarum__TM_#01
MKNRNPLIRKLLTQYFVTTGILLAFLVMIPLVIRFIAGTRTWYGTEPIYYILRFFADRWL
FCVAIGALLIWFGTTIYYMTKAIGYLNETIQATTQLIEEPSKRITLSSHLID
>gb|AAF86642_1|ARO_3002933|vanSC__Enterococcus_gallinarum__TM_#02
NAMRNRYTEIALQKAQRLEL
>gb|AAF86642_1|ARO_3002933|vanSC__Enterococcus_gallinarum__TM_#03
VLQTETTDLSLMLEQLTFEFLPLLEEKNLNWQLNLQKNVLATVDTEKIA
>gb|AAF86642_1|ARO_3002933|vanSC__Enterococcus_gallinarum__TM_#04
LLELVESDSIHIRLTNRGKTIPEEMIGRL
>gb|AAF86642_1|ARO_3002933|vanSC__Enterococcus_gallinarum__TM_#05
LLASGGDISAESKDETIIFNVRLPKPANN
>gb|AAA85213_1|ARO_3002866|dfrD__Listeria_monocytogenes__TM_#01
KDCEIAHSIEAA
>gb|AAA85213_1|ARO_3002866|dfrD__Listeria_monocytogenes__TM_#02
VVNFDDWKEV
>gb|WP_063964000_1|ARO_3004102|kamB__Streptoalloteichus_tenebrarius__TM_#01
MRRVVGKRVQEFSDAEFEQLRSQYDDVVLDVGTGDGKHPYKVARQNPSRLVVALDADKSR
MEK
>gb|WP_063964000_1|ARO_3004102|kamB__Streptoalloteichus_tenebrarius__TM_#02
RLPPLSGVG
>gb|WP_063964000_1|ARO_3004102|kamB__Streptoalloteichus_tenebrarius__TM_#03
SPEMLRGMAAVCR
>gb|WP_063964000_1|ARO_3004102|kamB__Streptoalloteichus_tenebrarius__TM_#04
DEWLAPRYAEAGWKLADCRYLEP
>gb|BAE77778_1|ARO_3000508|gadX__Escherichia_coli_str__K-12_substr__W3110__TM_#01
VSSLLVHHCSRDIPVFQEVAQLSQNKNLRYAEMLRKRALIFALLSVFLEDEHFIPLLLNV
LQPNMRTRVCT
>gb|CAE50925_1|ARO_3002589|AAC(6_)-Iid__Enterococcus_hirae__TM_#01
PMKEVEQLMAP
>gb|CAE50925_1|ARO_3002589|AAC(6_)-Iid__Enterococcus_hirae__TM_#02
YGGLVIYLGTDDVEGQTNLVETDLFEDTFAKLQEIKNI
>gb|CAE50925_1|ARO_3002589|AAC(6_)-Iid__Enterococcus_hirae__TM_#03
QPDIWLAKRVAKREPTE
>gb|AAG44836_1|ARO_3004740|AST-1__Nocardia_asteroides__TM_#01
MTFSALPFRRADRRRLLAAALAACALTLTAACDSGTVTVPVTD
>gb|AAR92235_1|ARO_3000602|Erm(39)__Mycolicibacterium_fortuitum__TM_#01
LHRARRLADRTTAEVIAT
>gb|AAR92235_1|ARO_3000602|Erm(39)__Mycolicibacterium_fortuitum__TM_#02
QRRAEPLLPWADRRA
>gb|AAR92235_1|ARO_3000602|Erm(39)__Mycolicibacterium_fortuitum__TM_#03
HPRWLSANGIHPSALPRALTARQWVALFDAAG
>gb|BAJ05825_1|ARO_3002263|IND-7__Chryseobacterium_indologenes__TM_#01
QVKDFVIEPPI
>gb|BAJ05825_1|ARO_3002263|IND-7__Chryseobacterium_indologenes__TM_#02
IYKTFGVFGGKEYSAN
>gb|BAJ05825_1|ARO_3002263|IND-7__Chryseobacterium_indologenes__TM_#03
LFDVPWEK
>gb|BAJ05825_1|ARO_3002263|IND-7__Chryseobacterium_indologenes__TM_#04
QYQSLMDTI
>gb|BAJ05825_1|ARO_3002263|IND-7__Chryseobacterium_indologenes__TM_#05
VLDGGCLVKS
>gb|BAJ05825_1|ARO_3002263|IND-7__Chryseobacterium_indologenes__TM_#06
ATDLGYIKEAN
>gb|AAB41059_1|ARO_3002884|iri__Rhodococcus_hoagii__TM_#01
STSVFPASRTS
>gb|AAB41059_1|ARO_3002884|iri__Rhodococcus_hoagii__TM_#02
FRVVVPAAEVADGRATPT
>gb|AAB41059_1|ARO_3002884|iri__Rhodococcus_hoagii__TM_#03
DLLGRRLRNIALTRGNLYD
>gb|AAB41059_1|ARO_3002884|iri__Rhodococcus_hoagii__TM_#04
VDGWSDRADHIVDT
>gb|AAB41059_1|ARO_3002884|iri__Rhodococcus_hoagii__TM_#05
AELDTQLSTWFGRSARDRA
>gb|BAE83690_1|ARO_3003724|vanWI__Desulfitobacterium_hafniense_Y51__TM_#01
MLRRKPWVFLLVILFSQLSLSILLGATVTYGAGYAKAPEGLTVWEKDLGGMTKDEAYAVL
AEVIPKAVVYDRTVYFLELNQTD
>gb|BAE83690_1|ARO_3003724|vanWI__Desulfitobacterium_hafniense_Y51__TM_#02
RTIPSPELLNQEEVLAQLRKFALDIDQPGKAAEAYYENGEIVIEEGSLGVRLDVDKSWEQ
LQQSIGMETVPLVTEVIV
>gb|BAE83690_1|ARO_3003724|vanWI__Desulfitobacterium_hafniense_Y51__TM_#03
PSFHERVTNVRLAAE
>gb|BAE83690_1|ARO_3003724|vanWI__Desulfitobacterium_hafniense_Y51__TM_#04
TTQGYLLLNAATGGNWIRVRIFGVADSEHPALDEPDGYPVKPREWSK
>gb|CWV56762_1|ARO_3000198|FosX__Listeria_monocytogenes__TM_#01
MISGLSHITLIVKDLNKTTAFLQNIFNAEEIYSSGDKTFSL
>gb|CWV56762_1|ARO_3000198|FosX__Listeria_monocytogenes__TM_#02
SLQERTYNHIAFQIQSEEVDEYTERIKALGVEMKPE
>gb|BAJ17544_1|ARO_3002069|CMY-59__Escherichia_coli__Partial_TM_#01
GPGHLFAFNYGTDF
>gb|BAJ17544_1|ARO_3002069|CMY-59__Escherichia_coli__Partial_TM_#02
SPGQLELDAEAYGVKSSV
>gb|BAJ17544_1|ARO_3002069|CMY-59__Escherichia_coli__Partial_TM_#03

GSDSKVALA
>gb|ENX09209_1|ARO_3001752|OXA-297__Acinetobacter_sp__NIPH_1847__TM_#01
MLFNFIVMPKKLKLL
>gb|ENX09209_1|ARO_3001752|OXA-297__Acinetobacter_sp__NIPH_1847__TM_#02
LSVVVMPSIILLGCQNIQPHVQ
>gb|ENX09209_1|ARO_3001752|OXA-297__Acinetobacter_sp__NIPH_1847__TM_#03
LVTQKQTEDQIATAFENIQTSGVLVTYDGKAIQKYGNA
>gb|ENX09209_1|ARO_3001752|OXA-297__Acinetobacter_sp__NIPH_1847__TM_#04
KQLAFDSSTQQQVKDMLLIEDI
>gb|ABG91835_1|ARO_3002860|dfrA17__Pseudomonas_aeruginosa__TM_#01
MKISLISA
>gb|ABG91835_1|ARO_3002860|dfrA17__Pseudomonas_aeruginosa__TM_#02
SNENVLVFPSIE
>gb|ABG91835_1|ARO_3002860|dfrA17__Pseudomonas_aeruginosa__TM_#03
SLIEKADIIHLST
>gb|ABG91835_1|ARO_3002860|dfrA17__Pseudomonas_aeruginosa__TM_#04
HVEVEGDI
>gb|AAA03550_1|ARO_3002523|AAC(2_)-Ia__Providencia_stuartii__TM_#01
MGIEYRSLHTSQLTLSEKEALYDLLIEGFEGDFSHDDFAHTLGGMHVMAFDQQKLVGHVA
IIQRHMALDNT
>gb|AAA03550_1|ARO_3002523|AAC(2_)-Ia__Providencia_stuartii__TM_#02
QLGLLSASDDGQKLYHSVGWQIWKGKLFELKQGS
>gb|ACJ63260_1|ARO_3000413|sul3__Escherichia_coli__TM_#01
PDTTEVGVVEEIKRLKPVIKALKEKGISISVDTFKPEVQSFCIEQKVDFINDIQGFPYPE
IYSGLAKSDCKLVLMHSVQRIGAATKVETNPEEVFTSMMEFFKERIAALVE
>gb|ACJ63260_1|ARO_3000413|sul3__Escherichia_coli__TM_#02
LVLKRFPEIQEAFNLQ
>gb|WP_114268491_1|ARO_3005323|OXA-926__Klebsiella__TM_#01
MCNRILQVAAVSVV
>gb|WP_114268491_1|ARO_3005323|OXA-926__Klebsiella__TM_#02
RVLLQRGSACS
>gb|WP_114268491_1|ARO_3005323|OXA-926__Klebsiella__TM_#03
QYVKAFQYGNADVSAVPVDDPGQSGAWVMSSLRISPK
>gb|WP_114268491_1|ARO_3005323|OXA-926__Klebsiella__TM_#04
LAFARLIQDEQAIKPNAGLRARDELLRLLPAIDH
>gb|BAD11815_1|ARO_3002593|AAC(6_)-Ib-SK__Streptomyces_kanamyceticus__TM_#01
MRAHRSCCIRRRGLGHNAGVELNGEKVLLRPVLDSDVKK
>gb|BAD11815_1|ARO_3002593|AAC(6_)-Ib-SK__Streptomyces_kanamyceticus__TM_#02
DYEEMLAITLDGEVIGAVQYE
>gb|BAD11815_1|ARO_3002593|AAC(6_)-Ib-SK__Streptomyces_kanamyceticus__TM_#03
SRHGLGLGTDTVRTVARW
>gb|AAF86220_1|ARO_3002844|vatE__Enterococcus_faecium__TM_#01
GPRFEPEVIQALENLAWWNKDIEWITANVPKLMQTTPTLELINSLMEK
>gb|AJP77054|ARO_3003716|CPS-1__Chryseobacterium_sp__Stok-1__TM_#01
MRNLTLLFLLCL
>gb|AAA22904_1|ARO_3000578|CcrA_beta-lactamase__Bacteroides_fragilis__TM_#01
MKTVFILISMLFPVAVMAQKSVKISDDISITQLSDKVYTYVSLAEIEGWGMVPSNGMIVI
NNHQAALLDTPINDAQTEMLVNWVTDSLHAKVTTFIPNHWHGDCIGGLGYLQRKGVQSYA
NQMTIDLAKEKGLPVPEHGFTDSLTVSLDGMPLQCY
>gb|AAA22904_1|ARO_3000578|CcrA_beta-lactamase__Bacteroides_fragilis__TM_#02
TENILFGGCMLKDNQATSIGNISDADVTAWPKTLDKVKAKFPSARYVVPGHGDYGGTELI
EHTKQIVNQYIESTSKP
>gb|AAU95768_1|ARO_3000026|mepA__Staphylococcus_aureus__TM_#01
IALGLIVILVTLPFSDQIAAILGARGETLALTSN
>gb|AAU95768_1|ARO_3000026|mepA__Staphylococcus_aureus__TM_#02
MKNSDVVSVNIKLA
>gb|AAU95768_1|ARO_3000026|mepA__Staphylococcus_aureus__TM_#03
LQGAIIIPVLFIMNALFGLTGVIWSLLIAESLCALA
>gb|ABX54692_1|ARO_3002938|vanSL__Enterococcus_faecalis__TM_#01
MKSKAETTTIKQILIKYLVTIGLSMLAYLVFLLTILIIMRNFVWDGTEPIYRVLHFFYRL
FNFEGILIIGVILILFVVTLFFVMKIIGYLKQIIEATKQLLEKPEQRVKLSSGLFELQEE
MNQLREKNNADNRAAKEAEK
>gb|ABX54692_1|ARO_3002938|vanSL__Enterococcus_faecalis__TM_#02
VQTRAKYTNIALSKAFRLEELLSEFFDVTRFNLTNLTINEELVDLSVMLEQISYEFLPIL
EEKKLSWNLHVESNIKSLLDPG
>gb|ABX54692_1|ARO_3002938|vanSL__Enterococcus_faecalis__TM_#03
TIIDLSLEKKESQAIFKITNRTY
>gb|ABX54692_1|ARO_3002938|vanSL__Enterococcus_faecalis__TM_#04
PFYRMDTSRSSSTGGTGLGLPIVREIIEASKGTINVSSSNNEMTFIIYLPYID
>gb|AAB49832_1|ARO_3002636|APH(2__)-IIIa__Enterococcus_gallinarum__TM_#01
MKQNKLHYTTMIMTQFPDISIQ
>gb|AAB49832_1|ARO_3002636|APH(2__)-IIIa__Enterococcus_gallinarum__TM_#02
VGCVKVNIPQYVYIGKRSDGNPFVGYRKVQGQILGEDGMAVFPDDAKDRLALQLAEFMNE
LSAFPVETAISAGVPVTNLKNKILLLSEAVEDQVFPLLDESLRDYLTLRFQSYMTHPV
>gb|AAB49832_1|ARO_3002636|APH(2__)-IIIa__Enterococcus_gallinarum__TM_#03
LFTRQVMAYRGEVDLDTHIRKVSLFVTFDQVSYLLEGLRARDQDWISEGLELLEEDKANN
FGANSA
>gb|AAW66497_1|ARO_3000566|tet(39)__Acinetobacter_sp__LUH5605__TM_#01
MKKSLSVILITIF
>gb|AAW66497_1|ARO_3000566|tet(39)__Acinetobacter_sp__LUH5605__TM_#02
ATADYLLMAAAPSLLW
>gb|AAW66497_1|ARO_3000566|tet(39)__Acinetobacter_sp__LUH5605__TM_#03
FAAAFMNGINLIMTAVLLKESKHSNKMTEKVQEQSILKKLSYLITQPNMAPLLGIFLIIT
LVSQVPATLWVIYGQDRYGWSIFIAGVSLASYGICHSIAQAFAIAPMVKRFGEKNTLLCG
IACDAIGLLLLSIAVEEWVPFALLPLFALGGVAVPALQAMMSRGISDERQGELQGLLSSF
NSLGAIIGPVLVTSLYFMTQASAPGMVWALAAILYVITLPLLLKYRLNKYSGVP
>gb|QHR93773_1|ARO_3005024|FosL1__Escherichia_coli__TM_#01
QSLERSVSFYKDTLGFRLAARWKN
>gb|QHR93773_1|ARO_3005024|FosL1__Escherichia_coli__TM_#02
KRSAQPWPEYTHYAFSALEQDFSSFVAKLIQKGVVEWKK
>gb|QHR93773_1|ARO_3005024|FosL1__Escherichia_coli__TM_#03
SRLKQCRVHPYTDMEIFD
>gb|AAG05188_1|ARO_3005068|ParR__Pseudomonas_aeruginosa_PAO1__TM_#01
AAFAAFLDFKPQ
>gb|AAG05188_1|ARO_3005068|ParR__Pseudomonas_aeruginosa_PAO1__TM_#02
HAPLPASPES
>gb|AAW38464_1|ARO_3004572|Staphylococcus_aureus_LmrS__Staphylococcus_aureus_subsp__aureus_COL__TM_#01
MAKVELTT
>gb|AAW38464_1|ARO_3004572|Staphylococcus_aureus_LmrS__Staphylococcus_aureus_subsp__aureus_COL__TM_#02
RVFKNRTFAL
>gb|AAW38464_1|ARO_3004572|Staphylococcus_aureus_LmrS__Staphylococcus_aureus_subsp__aureus_COL__TM_#03
LMSFGAKIFL
>gb|ADO96486_1|ARO_3003953|hmrM__Haemophilus_influenzae_R2846__TM_#01

LGVSIPLGLLIYFCEIPLQYMQMESKMSDL
>gb|ADO96486_1|ARO_3003953|hmrM__Haemophilus_influenzae_R2846__TM_#02
TNTQERSLKVFSQLIEMPNP
>gb|ADO96486_1|ARO_3003953|hmrM__Haemophilus_influenzae_R2846__TM_#03
QNAKKIGYAALLLGLTVTIVTAL
>gb|ADO96486_1|ARO_3003953|hmrM__Haemophilus_influenzae_R2846__TM_#04
ANLLLFAALYQFSDTIQMVVGGI
>gb|AAG29813_1|ARO_3002380|SME-2__Serratia_marcescens__TM_#01
MSNKVNFKTASFLFSVCLALSAFNAHANKSDAAAKQIKKLEEDFDGRIGVFAIDTGSGN
>gb|AAG29813_1|ARO_3002380|SME-2__Serratia_marcescens__TM_#02
VQQKKLDINQKVKYESRDLEY
>gb|AAG29813_1|ARO_3002380|SME-2__Serratia_marcescens__TM_#03
KYKGSGMTLGDMASAALQYSDNGATNIIMERFLGGPE
>gb|AAG29813_1|ARO_3002380|SME-2__Serratia_marcescens__TM_#04
SKDDKHSDKTIAEASRIAIQAID
>gb|AAA25655_1|ARO_3004657|catA4__Proteus_mirabilis__TM_#01
MDTKRVGILV
>gb|AAA25655_1|ARO_3004657|catA4__Proteus_mirabilis__TM_#02
QNGYKFYPTFIYIISLLVNK
>gb|AAA25655_1|ARO_3004657|catA4__Proteus_mirabilis__TM_#03
GYNIFHEQTETFSSLWSYYHKDINRFLKTYSEDIAQYGDDLAYFPK
>gb|ABG48543_1|ARO_3005052|LpsA__Haemophilus_influenzae__TM_#01
LDRSPKAKLS
>gb|ABG48543_1|ARO_3005052|LpsA__Haemophilus_influenzae__TM_#02
SDDIHVLKLEANGKMFFKQPKSVKCDRNVYPMTV
>gb|ABG48543_1|ARO_3005052|LpsA__Haemophilus_influenzae__TM_#03
LVKNKPLDVAVDSLVFEDFLHF
>gb|ADA62098_1|ARO_3003905|ANT(4_)-Ib__Staphylococcus_aureus__TM_#01
MNGPIIMTREERMKIVHEIKERILDKYGDDVK
>gb|ADA62098_1|ARO_3003905|ANT(4_)-Ib__Staphylococcus_aureus__TM_#02
MCVMSTEEAEFSHEWTTGEWKVEVNFDSEEILLDYASQVES
>gb|ADA62098_1|ARO_3003905|ANT(4_)-Ib__Staphylococcus_aureus__TM_#03
SGGYLEKVYQTAKSVEAQTFHDAICALIVEELFEYAGKWRNIRVQGPTTFLPSLTVQVAM
AGAMLIGLHHRICYTTSASVLTEAVKQSDLPSGYDHLCQFVMSGQLSDSEKLLESLENFW
NGIQEWTERHGYIVDVSKRIPF
>gb|ABN48311_1|ARO_3001918|CTX-M-56__Escherichia_coli__TM_#01
MMTQSIRRSMLTVMATLPLLFSSAT
>gb|ABN48311_1|ARO_3001918|CTX-M-56__Escherichia_coli__TM_#02
KNLTLGKALAE
>gb|ABN48311_1|ARO_3001918|CTX-M-56__Escherichia_coli__TM_#03
LAAAAKIVTHGF
>gb|AAB23649_1|ARO_3004460|Vibrio_anguillarum_chloramphenicol_acetyltransferase__Vibrio_anguillarum__TM_#01
MEFRLVDLKTWKRKEYFTHYFES
>gb|AAB23649_1|ARO_3004460|Vibrio_anguillarum_chloramphenicol_acetyltransferase__Vibrio_anguillarum__TM_#02
TIKTGKAKLYPALLYAVSTVVNRHEEFRMTVDDEGQIGIFSEMMPCYTIFQKDTEMFSNI
WTEYIGDYTEFCKQYEKDMQQYGENK
>gb|AAB23649_1|ARO_3004460|Vibrio_anguillarum_chloramphenicol_acetyltransferase__Vibrio_anguillarum__TM_#03
QSLQNHGGDEE
>gb|AEH41427_1|ARO_3002218|IMP-27__Proteus_mirabilis__TM_#01
VFCSITVAGETLPNLRVEKLEE
>gb|QIM09823_1|ARO_3005056|Tet(X6)__Proteus_genomosp__6__TM_#01
AYGREAQAESIINETEMFSLDFSFQKLMNL
>gb|ABI20451_1|ARO_3003741|mphE__uncultured_bacterium__TM_#01
VFALDTKGQQWLLRIPRRDGMREQIK
>gb|ABI20451_1|ARO_3003741|mphE__uncultured_bacterium__TM_#02
TELVAYPILKDNPVLNLDAETYEII
>gb|ABI20451_1|ARO_3003741|mphE__uncultured_bacterium__TM_#03
FEIHSIPEKEVRENDLKIMKPSDLRPEIANNLQLVKSEIGISEQLETRYRKWLDNDV
>gb|ABI20451_1|ARO_3003741|mphE__uncultured_bacterium__TM_#04
MYGLFALETQNESLIVGAKAQLGVI
>gb|AAD08227_1|ARO_3003964|hp1181__Helicobacter_pylori_26695__TM_#01
MGILSDKIGRKVVVMVCLLLFLAGSLVCFIANDIVW
>gb|AAD08227_1|ARO_3003964|hp1181__Helicobacter_pylori_26695__TM_#02
SYQIKNIKAYQPNSKALYLLYLSSFFEKA
>gb|AAD08227_1|ARO_3003964|hp1181__Helicobacter_pylori_26695__TM_#03
DESFLILVYVPGALLGVLSMGI
>gb|AAD08227_1|ARO_3003964|hp1181__Helicobacter_pylori_26695__TM_#04
YLWLFIVGVAF
>gb|AAD08227_1|ARO_3003964|hp1181__Helicobacter_pylori_26695__TM_#05
VSNTSLIVVALGLIWGLS
>gb|AAD08227_1|ARO_3003964|hp1181__Helicobacter_pylori_26695__TM_#06
NEEQFETLE
>gb|CAA90891_1|ARO_3004762|BRO-2__Moraxella_catarrhalis_ATCC_43617__TM_#01
MMQRRHFLQKTLLALPIIFSGNLLTGCKTNLSDDYLPDDKITNNPNLLQNKLKEILPIWE
NKFNAKIGMTIIADNGELSSHRGNEYFPVNSTIKAFIASHILLLVDKEKLDLNEKIIIKE
SDLIEYSPVCKKYFDENKPISISELCEATITLSDNGSANILLDKIGGLTAFNQFLKEIGA
DMVLANNEPLLNRSHYGETSDTAKPIPYTKSLKALIVGNILSNQSKEQLITWLINDKVAD
NLLRKY
>gb|CAA90891_1|ARO_3004762|BRO-2__Moraxella_catarrhalis_ATCC_43617__TM_#02
ENNKPYFISLFITQPHDGKSL
>gb|CAA90891_1|ARO_3004762|BRO-2__Moraxella_catarrhalis_ATCC_43617__TM_#03
FKNQKDEIMAQIGKEIYPFL
>gb|AAA27459_1|ARO_3004683|aadS__Transposon_Tn4551__TM_#01
EYKYPDKLENDIRKYLAGLKPKT
>gb|CAA29136_1|ARO_3002658|APH(6)-Ib__Streptomyces_glaucescens__TM_#01
ELTADGASAS
>gb|CAA29136_1|ARO_3002658|APH(6)-Ib__Streptomyces_glaucescens__TM_#02
ITGLRTWNGH
>gb|CAA29136_1|ARO_3002658|APH(6)-Ib__Streptomyces_glaucescens__TM_#03
SAALDPAAVTLAQALRGH
>gb|AAA25680_1|ARO_3002595|AAC(6_)-IIb__Pseudomonas_fluorescens__TM_#01
MHPGVVTLRPMTED
>gb|AAA25680_1|ARO_3002595|AAC(6_)-IIb__Pseudomonas_fluorescens__TM_#02
RPSLEEVKEDYRPSALAEEGVTPYIGLLDGTPFA
>gb|AAA25680_1|ARO_3002595|AAC(6_)-IIb__Pseudomonas_fluorescens__TM_#03
SGLLGRGYGTR
>gb|AAA25680_1|ARO_3002595|AAC(6_)-IIb__Pseudomonas_fluorescens__TM_#04
VVSTPDGPAMYMLHERPLVNGLRSAA
>gb|ABA28305_2|ARO_3000462|mgtA__Streptomyces_lividans__TM_#01
TARVLGRRWEVPVIS
>gb|AET35493_1|ARO_3001777|OXA-347__uncultured_bacterium__TM_#01
MKNILFVVFISMIFLFVCCNTTTNKNIIETEISDFDKILDSFQVNGSILIYDNDKNTFYS
NDFDWAKN

>gb|AET35493_1|ARO_3001777|OXA-347__uncultured_bacterium__TM_#02
ENDTTILKWNGEQRKMDIWEKDLSF
>gb|AET35493_1|ARO_3001777|OXA-347__uncultured_bacterium__TM_#03
IKMKEYLEKFEYKN
>gb|AAP22374_1|ARO_3002248|CARB-9__Vibrio_cholerae_non-O1_non-O139__TM_#01
MKSLLVFALLMPSVVFASSSKFQSVEQEIKGIESSLSARIGVAILDTQNGES
>gb|AAP22374_1|ARO_3002248|CARB-9__Vibrio_cholerae_non-O1_non-O139__TM_#02
KPITLSDAC
>gb|AAP22374_1|ARO_3002248|CARB-9__Vibrio_cholerae_non-O1_non-O139__TM_#03
ATMTTSDNTAANIVINAVGDPK
>gb|AAP22374_1|ARO_3002248|CARB-9__Vibrio_cholerae_non-O1_non-O139__TM_#04
TSTLNQLLFGSTLSEAS
>gb|AAF36803_1|ARO_3002908|vanF__Paenibacillus_popilliae_ATCC_14706__TM_#01
LVPANLSAEKRIK
>gb|ABO40886_1|ARO_3004645|dfrI__Yersinia_ruckeri__TM_#01
MNQNRDDHNRAADREKTAEKRGENQC
>gb|ABO40886_1|ARO_3004645|dfrI__Yersinia_ruckeri__TM_#02
SLEKEALPGCLIYSDLSVAIAALKKEPEVEEIMIMGGAQIYRAALPMM
>gb|ABO40886_1|ARO_3004645|dfrI__Yersinia_ruckeri__TM_#03
TQMPPFDFSHATLIFEEKHF
>gb|AAO82019_1|ARO_3004254|vanVB__Enterococcus_faecalis_V583__TM_#01
MFTEKFCADGICFIMRAKNEIDHIFSELYSVPNC
>gb|AAO82019_1|ARO_3004254|vanVB__Enterococcus_faecalis_V583__TM_#02
PLVICTPILIG
>gb|AAO82019_1|ARO_3004254|vanVB__Enterococcus_faecalis_V583__TM_#03
RSIVNRLRAEQKENQQKQVVLALLIHSELFDSGFR
>gb|AAA98298_1|ARO_3002545|AAC(6_)-Ia__Plasmid_R__TM_#01
MNYQIVNIAECSNYQLEAANILTEAFNDLGNNSWPDMTSATKEVKECIESPNLCFGLLIN
NS
>gb|AAA98298_1|ARO_3002545|AAC(6_)-Ia__Plasmid_R__TM_#02
PDYQNKGIGKILLKELENRAREQ
>gb|AAA98298_1|ARO_3002545|AAC(6_)-Ia__Plasmid_R__TM_#03
YRTSLSLITITEDNIFDS
>gb|AAG06466_1|ARO_3005064|cprS__Pseudomonas_aeruginosa_PAO1__TM_#01
MKRGLSLIPIVGGGVTLILAGVLLVYTRMLGDYGETGALYLLSMMMEEEGLYFAQRYQED
PATPAPDSYYFKGSVGTAGLPPKLREMLDTPPYKSIGAMQLLGNWDDDDEEEDDDAPSDD
AYVVVRQPLADGKTLYLYDNDA
>gb|AAG06466_1|ARO_3005064|cprS__Pseudomonas_aeruginosa_PAO1__TM_#02
LLDQPGRAPSPHALQRIRRSALG
>gb|AAG06466_1|ARO_3005064|cprS__Pseudomonas_aeruginosa_PAO1__TM_#03
SGELRDDGHIEVGRLLEELLEEQQALSQRRGLTFHLDVEPHSLPQTRARIIIGNLLRNAL
QYSDEGVVEIVVRDRSLL
>gb|CAB13166_1|ARO_3003063|ykkC__Bacillus_subtilis_subsp__subtilis_str__168__TM_#01
ALTWSGTAIGIIFSFYLLMKATHS
>gb|CAB13166_1|ARO_3003063|ykkC__Bacillus_subtilis_subsp__subtilis_str__168__TM_#02
AGTVLSEIVLFHEPVGWPKLL
>gb|CAB13166_1|ARO_3003063|ykkC__Bacillus_subtilis_subsp__subtilis_str__168__TM_#03
QDETEEKGGEA
>gb|ALX99516_1|ARO_3003811|adeC__Acinetobacter_baumannii__TM_#01
MIAEKSYKEFISD
>gb|ALX99516_1|ARO_3003811|adeC__Acinetobacter_baumannii__TM_#02
SQLPTIGVTGNAVRQVSPSINPNNPV
>gb|ALX99516_1|ARO_3003811|adeC__Acinetobacter_baumannii__TM_#03
SNITQVWLNYAFAQ
>gb|ALX99516_1|ARO_3003811|adeC__Acinetobacter_baumannii__TM_#04
QQAKNLLDLLAGHPVPQNLLPDHAIQNITF
>gb|ALX99516_1|ARO_3003811|adeC__Acinetobacter_baumannii__TM_#05
TAATYKLSMARYKAGVDSYF
>gb|ALX99516_1|ARO_3003811|adeC__Acinetobacter_baumannii__TM_#06
IKLNNQIEIYKVLGGGISKV
>gb|AAO04716_1|ARO_3002865|dfrC__Staphylococcus_epidermidis_ATCC_12228__TM_#01
MTLSIIVAHDK
>gb|AAO04716_1|ARO_3002865|dfrC__Staphylococcus_epidermidis_ATCC_12228__TM_#02
NQASFHHEGVDVINSLDEIKE
>gb|AWN09461_1|ARO_3004571|PNGM-1__uncultured_bacterium__TM_#01
MAGGKVTSSTGIAPKRYVYYPGSEE
>gb|AWN09461_1|ARO_3004571|PNGM-1__uncultured_bacterium__TM_#02
AAWVVELGNGDKFIVDIGSGSMANIQSLMIPAN
>gb|AWN09461_1|ARO_3004571|PNGM-1__uncultured_bacterium__TM_#03
RPGDINVHEFDYRALNEVVYQENGVTFRSWPCIHAGDGPVSFALEWNGYKVVFGGDTAPN
IWYPEYAKGADLAIHECWMTSDQMMTKYNQPAQLALRINLD
>gb|AWN09461_1|ARO_3004571|PNGM-1__uncultured_bacterium__TM_#04
TGVRENYA
>gb|AWN09461_1|ARO_3004571|PNGM-1__uncultured_bacterium__TM_#05
HAWDVAGPSEDLAPDRNRASEYTQYILDGRLNVDEANAHWKQEFMGRTGLTTEDLGVGS
>gb|AEZ05106_1|ARO_3002587|AAC(6_)-33__Escherichia_coli__TM_#01
FCEIGESNEYII
>gb|AEZ05106_1|ARO_3002587|AAC(6_)-33__Escherichia_coli__TM_#02
AARILTKSFLDIGN
>gb|AEZ05106_1|ARO_3002587|AAC(6_)-33__Escherichia_coli__TM_#03
SWPDMKSATKEVEECIEKPNICLGIHENEK
>gb|AEZ05106_1|ARO_3002587|AAC(6_)-33__Escherichia_coli__TM_#04
STQYQNKGIGRLLINELEK
>gb|AFK80745_1|ARO_3002350|GES-21__uncultured_bacterium__TM_#01
MRFIHALLLA
>gb|AFK80745_1|ARO_3002350|GES-21__uncultured_bacterium__TM_#02
IAHSAYASEKLTFKTDLEKLEREKAAQIGVAIVDPQGEIVAGHR
>gb|AFK80745_1|ARO_3002350|GES-21__uncultured_bacterium__TM_#03
FPLAALVFERIDSGTERGDRKLSYGPDMIV
>gb|AFK80745_1|ARO_3002350|GES-21__uncultured_bacterium__TM_#04
WSPATERFLASGHMTVLEAAQAAVQ
>gb|AFK80745_1|ARO_3002350|GES-21__uncultured_bacterium__TM_#05
AAMTQYFRK
>gb|AFK80745_1|ARO_3002350|GES-21__uncultured_bacterium__TM_#06
KAQERDYAVAVYTTAPKLSAVERDELVASVGQVITQLILSTDK
>gb|AAN28721_1|ARO_3002871|tet37__uncultured_bacterium__TM_#01
NALKKLAGI
>gb|AAN28721_1|ARO_3002871|tet37__uncultured_bacterium__TM_#02
EKVVNLYDRMSKYETD
>gb|AAN28721_1|ARO_3002871|tet37__uncultured_bacterium__TM_#03
DRTPSMYEYIWIKENREAKVVSSFAANIYLGWGR
>gb|AEP40500_1|ARO_3002912|vanN__Enterococcus_faecium__TM_#01
MKKIALIFGGTSAEYEVSLKSAASVLSVLENLNVEIYRIGIASNGKWYLTFSDNETIAND

LWLQDKKLNEITPSFDGRGFYDQAEKVYFKPDVLFPMLHGGTGENGTLQGVFECMQIP
>gb|AEP40500_1|ARO_3002912|vanN__Enterococcus_faecium__TM_#02
YLLHQFAKSVGVMSTPTQLISSTDEQQVIKNFTELY
>gb|AEP40500_1|ARO_3002912|vanN__Enterococcus_faecium__TM_#03
HTEAELTKALTEAFQFSQTVILQKAVSGVEIGCAILGNDQ
>gb|AEP40500_1|ARO_3002912|vanN__Enterococcus_faecium__TM_#04
ATDFFDYTEKYQMTTAKLTVPAKIPVATSREIKRQAQLLYQLLGCQ
>gb|AEP40500_1|ARO_3002912|vanN__Enterococcus_faecium__TM_#05
TGITYQELISTLITLAEDK
>gb|AIT76106_1|ARO_3002146|DHA-15__Klebsiella_pneumoniae__TM_#01
SALLAFSAPGFSAADNVA
>gb|AIT76106_1|ARO_3002146|DHA-15__Klebsiella_pneumoniae__TM_#02
FYQQWQPS
>gb|AIT76106_1|ARO_3002146|DHA-15__Klebsiella_pneumoniae__TM_#03
AAGMPYEQLLT
>gb|AIT76106_1|ARO_3002146|DHA-15__Klebsiella_pneumoniae__TM_#04
PLGLSHTFITVP
>gb|AIT76106_1|ARO_3002146|DHA-15__Klebsiella_pneumoniae__TM_#05
PSRAGNADLEMAMYL
>gb|AIT76106_1|ARO_3002146|DHA-15__Klebsiella_pneumoniae__TM_#06
NGVTNEVALQPHPVTDNQVQPYN
>gb|AAA20116_1|ARO_3000180|tetA(P)__Clostridium_perfringens__TM_#01
EIYIKGAQAGQIGAF
>gb|AAA20116_1|ARO_3000180|tetA(P)__Clostridium_perfringens__TM_#02
KPIFSAWLNGHIDDNSRATVLSINGQMNSLGQILGGPIIGIIATNISVSIGIVCTSLLVT
PVLVLYIVAMIIDKKVDDRVGGIDYEENN
>gb|BBJ69931_1|ARO_3005031|FRI-8__Enterobacter_sp__18A13__TM_#01
FGGRIGVYLLNT
>gb|BBJ69931_1|ARO_3005031|FRI-8__Enterobacter_sp__18A13__TM_#02
NGKEFSYRQ
>gb|BBJ69931_1|ARO_3005031|FRI-8__Enterobacter_sp__18A13__TM_#03
VFLAASVLK
>gb|BBJ69931_1|ARO_3005031|FRI-8__Enterobacter_sp__18A13__TM_#04
RVMEKHSPVSEKY
>gb|BBJ69931_1|ARO_3005031|FRI-8__Enterobacter_sp__18A13__TM_#05
MSLKNIAFGS
>gb|BBJ69931_1|ARO_3005031|FRI-8__Enterobacter_sp__18A13__TM_#06
AKNKALLQDWL
>gb|BBJ69931_1|ARO_3005031|FRI-8__Enterobacter_sp__18A13__TM_#07
GNTTGNARVRAAVPDK
>gb|BBJ69931_1|ARO_3005031|FRI-8__Enterobacter_sp__18A13__TM_#08
NSPAVIAVYTTR
>gb|BBJ69931_1|ARO_3005031|FRI-8__Enterobacter_sp__18A13__TM_#09
NQNDKHDET
>gb|BBJ69931_1|ARO_3005031|FRI-8__Enterobacter_sp__18A13__TM_#10
IKNAAKIAI
>gb|CAK55557_1|ARO_3002585|AAC(6_)-31__Pseudomonas_putida__TM_#01
AEVLEQYLPSALAK
>gb|CAK55557_1|ARO_3002585|AAC(6_)-31__Pseudomonas_putida__TM_#02
CALVEMLFKDA
>gb|AAL14439_1|ARO_3000774|adeA__Acinetobacter_baumannii__TM_#01
DSKEVAQAEP
>gb|AAL14439_1|ARO_3000774|adeA__Acinetobacter_baumannii__TM_#02
ARLKVQLERYEQLLPSNAISKQEV
>gb|AAL14439_1|ARO_3000774|adeA__Acinetobacter_baumannii__TM_#03
QGTAEIRPIEIGQQYEQFYIANKGLKVGDRVVVEGIERIKPNQKLA
>gb|AAV85981_1|ARO_3000533|macA__Neisseria_gonorrhoeae__TM_#01
SGGYNSSTDTASNAVYYYARSF
>gb|AAV85981_1|ARO_3000533|macA__Neisseria_gonorrhoeae__TM_#02
DKVVISEITAAEQQESGERALGGPPRR
>gb|QEQ43477_1|ARO_3005062|ANT(3__)-Ib__Cupriavidus_gilardii__TM_#01
MPPPANEPVPAEVQPILDVVVR
>gb|QEQ43477_1|ARO_3005062|ANT(3__)-Ib__Cupriavidus_gilardii__TM_#02
SRQSREDLVAALMEVSGARAGRGPARNAEVTVVVLGDIAPWRH
>gb|QEQ43477_1|ARO_3005062|ANT(3__)-Ib__Cupriavidus_gilardii__TM_#03
TADPDLTLVLATALQSHRALMGPGLAEFLPAIPYADIRRAMADSLPGLVAN
>gb|QEQ43477_1|ARO_3005062|ANT(3__)-Ib__Cupriavidus_gilardii__TM_#04
PERRALLATARDAYRGHVADEAWHECRADVSEWVSEVSQAIARTLHDSVKA
>gb|AAC75314_1|ARO_3003578|pmrF__Escherichia_coli_str__K-12_substr__MG1655__TM_#01
MLVEASQAE
>gb|AAC75314_1|ARO_3003578|pmrF__Escherichia_coli_str__K-12_substr__MG1655__TM_#02
IGGFSIAV
>gb|AAY32951_1|ARO_3002837|lnuC__Streptococcus_agalactiae__TM_#01
MVNITDVNQIFQFAIDA
>gb|AAY32951_1|ARO_3002837|lnuC__Streptococcus_agalactiae__TM_#02
YQSRAHNDIDIFVEKNDYQNFIEIMK
>gb|AAY32951_1|ARO_3002837|lnuC__Streptococcus_agalactiae__TM_#03
DEGEILYDGDCFPVETLSGKGRIEEIEVSCIEPYS
>gb|AAY32951_1|ARO_3002837|lnuC__Streptococcus_agalactiae__TM_#04
DENDAHDVKLLCETLHIEIPNEYR
>gb|ABW76138_1|ARO_3001767|OXA-137__Brachyspira_pilosicoli__TM_#01
MSKKNFILIFIFVIL
>gb|ABW76138_1|ARO_3001767|OXA-137__Brachyspira_pilosicoli__TM_#02
SNETTLIDN
>gb|ABW76138_1|ARO_3001767|OXA-137__Brachyspira_pilosicoli__TM_#03
FTNSNAEGTLVIYNLNDDKYIIHNKE
>gb|ABW76138_1|ARO_3001767|OXA-137__Brachyspira_pilosicoli__TM_#04
NEKAVKDVDEVFYK
>gb|ABW76138_1|ARO_3001767|OXA-137__Brachyspira_pilosicoli__TM_#05
RYAIKNSQVPAYKELARRIG
>gb|ABW76138_1|ARO_3001767|OXA-137__Brachyspira_pilosicoli__TM_#06
KMKENIEKLDFGNK
>gb|ABW76138_1|ARO_3001767|OXA-137__Brachyspira_pilosicoli__TM_#07
EGPLEISAMEQ
>gb|ABW76138_1|ARO_3001767|OXA-137__Brachyspira_pilosicoli__TM_#08
LLTKLAQNEL
>gb|ABW76138_1|ARO_3001767|OXA-137__Brachyspira_pilosicoli__TM_#09
YPIEIQKA
>gb|ABW76138_1|ARO_3001767|OXA-137__Brachyspira_pilosicoli__TM_#10
TLHGKTGL
>gb|ABW76138_1|ARO_3001767|OXA-137__Brachyspira_pilosicoli__TM_#11
NMTTEPIGW
>gb|ABW76138_1|ARO_3001767|OXA-137__Brachyspira_pilosicoli__TM_#12

NIYVFALNIDNINSDDLAKRINI
>gb|ABW76138_1|ARO_3001767|OXA-137__Brachyspira_pilosicoli__TM_#13
LKALNLLK
>gb|AAB08924_1|ARO_3004650|tetU__Enterococcus_faecium__TM_#01
QESLDSFASPHFLPIDIKPIDKIVIEGLIAEPSNWSIIARHTKYKYRNLLKQESQ
>gb|ADM92605_1|ARO_3000553|adeR__Acinetobacter_baumannii__TM_#01
SVIRAMNGKQAIELHASQP
>gb|ADM92605_1|ARO_3000553|adeR__Acinetobacter_baumannii__TM_#02
ANKATNKNKL
>gb|BAO79518_1|ARO_3003207|FosK__Acinetobacter_soli__TM_#01
TWEAGAYFTAGDTWVCLSVGE
>gb|BAO79518_1|ARO_3003207|FosK__Acinetobacter_soli__TM_#02
RELVELHARLKEAGVEEWKQNTSEGNSVYLLDPNGHRIELHCGTLATRLAELEKSPYKRL
VWC
>gb|AAS76623_1|ARO_3004606|erm(40)__Mycolicibacterium_mageritense__Partial_TM_#01
RRRAERLARRTTAHVVTADFLRYRLPPTE
>gb|AAS76623_1|ARO_3004606|erm(40)__Mycolicibacterium_mageritense__Partial_TM_#02
EPLIDGADRR
>gb|AAS76623_1|ARO_3004606|erm(40)__Mycolicibacterium_mageritense__Partial_TM_#03
VGPRVPRHWLRHNGIT
>gb|AAS76623_1|ARO_3004606|erm(40)__Mycolicibacterium_mageritense__Partial_TM_#04
EVTSEAKRC
>gb|AJP77071|ARO_3003714|PEDO-2__Pedobacter_sp__ALS-14__TM_#01
MKKIFLMVLFSCSLLGFAQTVTEPA
>gb|AJP77071|ARO_3003714|PEDO-2__Pedobacter_sp__ALS-14__TM_#02
SIIKENIKA
>gb|AJP77071|ARO_3003714|PEDO-2__Pedobacter_sp__ALS-14__TM_#03
LETGGKLDYELGKYGISFK
>gb|AJP77071|ARO_3003714|PEDO-2__Pedobacter_sp__ALS-14__TM_#04
ASYKDIQKDYTE
>gb|AJP77071|ARO_3003714|PEDO-2__Pedobacter_sp__ALS-14__TM_#05
APYNPKIFMDKSKFFKNLEDLENIFLEKIKN
>gb|AGC29882_1|ARO_3002853|arr-8__Klebsiella_oxytoca__TM_#01
MIKDWIPTTHENCKKMQ
>gb|AGC29882_1|ARO_3002853|arr-8__Klebsiella_oxytoca__TM_#02
NTHFEEGRTLKHVYFSAMLEPAIWGAELAVSLSGLDGRGY
>gb|AGC29882_1|ARO_3002853|arr-8__Klebsiella_oxytoca__TM_#03
SEPLQIVGVVEEWEGHSAEALKAMLDSLENLERNGLHVIYD
>gb|AAB58160_1|ARO_3003009|ceoA__Burkholderia_cepacia__TM_#01
KREPEGRRSRAGNGAHQSRLYAHHRAGVGPRVARGNHARQRRVGRRVGRAADDAGIGVAD
LRVVRRRRADLPAIHQRRAQRPQGAGRARPRERNRLLAQRRDRFGRQPARHVVRHDPRAR
PLRQRGRHPGPGPLRTREGGRQRAARGAARRRRGDQHRPGQEVRVRRRPAGPRVVSRSAA
RDAARQPARDRERAVGRRPRGRERHAARASGRAGEAAHGPDDGRRCAVRAVASTAKPAAP
AKADS
>gb|BAL14456_1|ARO_3000854|SMB-1_beta-lactamase__Serratia_marcescens__TM_#01
MKIIASLILAAFASVAQAQDRDWSSPQQPFTIYGNTHYVGTGGISAVLLSSPQGHILVDG
TTEKGAQVVAANIRAMGFKLS
>gb|BAL14456_1|ARO_3000854|SMB-1_beta-lactamase__Serratia_marcescens__TM_#02
LTGATVLAGAANVDTLRTGVSPKS
>gb|BAL14456_1|ARO_3000854|SMB-1_beta-lactamase__Serratia_marcescens__TM_#03
KAHATPGHTEGGITWTWQSCEQGKCKDV
>gb|BAL14456_1|ARO_3000854|SMB-1_beta-lactamase__Serratia_marcescens__TM_#04
SLRGSFEAVEK
>gb|BAL14456_1|ARO_3000854|SMB-1_beta-lactamase__Serratia_marcescens__TM_#05
TRQQRAAKEGNSAYVDNGACRA
>gb|AAG07593_1|ARO_3000807|mexH__Pseudomonas_aeruginosa_PAO1__TM_#01
YPPVKVALASVERRVVPRVFDGV
>gb|AAG07593_1|ARO_3000807|mexH__Pseudomonas_aeruginosa_PAO1__TM_#02
KAQLRNAEILHARARKLVERNVASQEQLDNAVAARDMALGAVRQTQAL
>gb|AAG07593_1|ARO_3000807|mexH__Pseudomonas_aeruginosa_PAO1__TM_#03
RVSRKADAPS
>gb|AAG07593_1|ARO_3000807|mexH__Pseudomonas_aeruginosa_PAO1__TM_#04
QDGDRPLSAKRVSVRIGERWDGRVEI
>gb|CBY88983_1|ARO_3002833|vgaE__Staphylococcus_aureus_subsp__aureus_ST398__TM_#01
MLLFEGTS
>gb|CBY88983_1|ARO_3002833|vgaE__Staphylococcus_aureus_subsp__aureus_ST398__TM_#02
LFDIDLIQVH
>gb|CBY88983_1|ARO_3002833|vgaE__Staphylococcus_aureus_subsp__aureus_ST398__TM_#03
FTSVKLVPQFKE
>gb|CBY88983_1|ARO_3002833|vgaE__Staphylococcus_aureus_subsp__aureus_ST398__TM_#04
EKSGGEITQQYLQ
>gb|CBY88983_1|ARO_3002833|vgaE__Staphylococcus_aureus_subsp__aureus_ST398__TM_#05
LDEPTTHLDT
>gb|CBY88983_1|ARO_3002833|vgaE__Staphylococcus_aureus_subsp__aureus_ST398__TM_#06
VVVSHDRTFLNNVCT
>gb|CBY88983_1|ARO_3002833|vgaE__Staphylococcus_aureus_subsp__aureus_ST398__TM_#07
FEKYEREK
>gb|CBY88983_1|ARO_3002833|vgaE__Staphylococcus_aureus_subsp__aureus_ST398__TM_#08
AIRQKEER
>gb|CBY88983_1|ARO_3002833|vgaE__Staphylococcus_aureus_subsp__aureus_ST398__TM_#09
HYANIQKKLR
>gb|CBY88983_1|ARO_3002833|vgaE__Staphylococcus_aureus_subsp__aureus_ST398__TM_#10
SAKALETRLEQLD
>gb|CBY88983_1|ARO_3002833|vgaE__Staphylococcus_aureus_subsp__aureus_ST398__TM_#11
DKVKELPEIKMD
>gb|CBY88983_1|ARO_3002833|vgaE__Staphylococcus_aureus_subsp__aureus_ST398__TM_#12
LTNQSVLRAENIKGE
>gb|CBY88983_1|ARO_3002833|vgaE__Staphylococcus_aureus_subsp__aureus_ST398__TM_#13
IGYFSQHLTIL
>gb|CBY88983_1|ARO_3002833|vgaE__Staphylococcus_aureus_subsp__aureus_ST398__TM_#14
LEALETLMKSYHGTILFVTHDR
>gb|CBY88983_1|ARO_3002833|vgaE__Staphylococcus_aureus_subsp__aureus_ST398__TM_#15
NIATKIIDIKDGKITVFDGSYEAYEEWLEN
>gb|CBY88983_1|ARO_3002833|vgaE__Staphylococcus_aureus_subsp__aureus_ST398__TM_#16
TKSNNDDQLLLIETKISDVLGRLSLEPS
>gb|CBY88983_1|ARO_3002833|vgaE__Staphylococcus_aureus_subsp__aureus_ST398__TM_#17
ELEDEFQRLLKEKKELT
>gb|ALA99191_1|ARO_3001462|OXA-157__Pandoraea_norimbergensis__TM_#01
MMMLSRWRRSAVVLRIAA
>gb|ALA99191_1|ARO_3001462|OXA-157__Pandoraea_norimbergensis__TM_#02
GRTGGYQAYDSTRANQ
>gb|ALA99191_1|ARO_3001462|OXA-157__Pandoraea_norimbergensis__TM_#03
VFLQRLARGQ

>gb|ALA99191_1|ARO_3001462|OXA-157__Pandoraea_norimbergensis__TM_#04
IERDGNITTVALNMDMR
>gb|AAD25063_1|ARO_3000183|tet(Z)__Corynebacterium_glutamicum__TM_#01
DQVGAPDDMIPLHVGLLTALYAI
>gb|AAD25063_1|ARO_3000183|tet(Z)__Corynebacterium_glutamicum__TM_#02
VSPHLPFLVAAALAGITLVLSASLLRETRPPGSNGSHAQQPGTAKRTAVPGMLILLAVFG
IVQFIGQAPGSTWVLFTQQRLDWNPVEVGVSLSIFGMVQVFVQAALTGRIVSRIGETRAI
LVGIAADAIGLIGLALIASTWAMLPILAALGLGSITLPALQTLLSRRAPEQQQGRLQGTL
ASLNSLTSIIGPVTFTGIFALTRTNADGTLWICAAALYVLCALLMIRETCASRRSR
>gb|ATL63232_1|ARO_3004544|mphJ__Brevibacillus_brevis_Vm4__TM_#01
MSKNNVEHMLALAKNNGILVDPTTVKVNESGLDFLAIFASTIDGIPWVLRQPRRDDVVET
ARYEKRVLDLVAKHLRVEVPDWQVHTSEFIAYPILG
>gb|ATL63232_1|ARO_3004544|mphJ__Brevibacillus_brevis_Vm4__TM_#02
YARASLREKMNEIKRVFGVSGA
>gb|ATL63232_1|ARO_3004544|mphJ__Brevibacillus_brevis_Vm4__TM_#03
VYLLFGDTGLADFIQRYEK
>gb|ATL63232_1|ARO_3004544|mphJ__Brevibacillus_brevis_Vm4__TM_#04
IMARQALGVDENGKEITS
>gb|AAC16242_1|ARO_3005069|rsmA__Pseudomonas_aeruginosa__TM_#01
MLILTRRVGETLMVGDDVTVTVLGVKGNQVRIGVNAPKEVAVHREEIYQRIQKEKDQEPN
H
>gb|AAA87229_1|ARO_3002558|AAC(6_)-Ik__Acinetobacter_sp__CIP-A165__TM_#01
KLRIKLWNDL
>gb|AAA87229_1|ARO_3002558|AAC(6_)-Ik__Acinetobacter_sp__CIP-A165__TM_#02
HALQLLVYSDDH
>gb|BAC20579_1|ARO_3000859|rmtA__Pseudomonas_aeruginosa__TM_#01
MSFDDALASILSSKKYRSLCPDTVRRILD
>gb|BAC20579_1|ARO_3000859|rmtA__Pseudomonas_aeruginosa__TM_#02
CLYDFIFSGGVP
>gb|BAC20579_1|ARO_3000859|rmtA__Pseudomonas_aeruginosa__TM_#03
HHQGLDFTFALQDVMCTPPTETGDLALVFK
>gb|BAC20579_1|ARO_3000859|rmtA__Pseudomonas_aeruginosa__TM_#04
DEFEIEDTKTIGIELVYMIKRNK
>gb|CFB60580_1|ARO_3001458|OXA-153__Pandoraea_apista__TM_#01
MKKIISRWRRGVFGLRVALAVVSPMVFAVPAHATEAAGGAAGTKA
>gb|CFB60580_1|ARO_3001458|OXA-153__Pandoraea_apista__TM_#02
GKYWNAAMDLRT
>gb|CFB60580_1|ARO_3001458|OXA-153__Pandoraea_apista__TM_#03
QFAQSKLNEAGYGNHTIGR
>gb|AAD03490_1|ARO_3002561|AAC(6_)-Ir__Acinetobacter_colistiniresistens__TM_#01
LEMRQLLEQPHTLQLLSYNDQ
>gb|AAD03490_1|ARO_3002561|AAC(6_)-Ir__Acinetobacter_colistiniresistens__TM_#02
CVVYFKKNIS
>gb|AKQ05896_1|ARO_3004587|tet(52)__uncultured_bacterium__TM_#01
YNMRTQLEFGRHVDP
>gb|AKQ05896_1|ARO_3004587|tet(52)__uncultured_bacterium__TM_#02
GFREGQDVEIIRSDLVEILIDSIEGVPLQFNQTIESVKQTDAV
>gb|AKQ05896_1|ARO_3004587|tet(52)__uncultured_bacterium__TM_#03
HEYKLNDLGA
>gb|AKQ05896_1|ARO_3004587|tet(52)__uncultured_bacterium__TM_#04
CEANQKLLSITSDKNRKMAEVAFSFRGQNVLNNVRDESEQRRV
>gb|AKQ05896_1|ARO_3004587|tet(52)__uncultured_bacterium__TM_#05
SDSFTQVI
>gb|AKQ05896_1|ARO_3004587|tet(52)__uncultured_bacterium__TM_#06
KRAFDRYNEILRPFIEANQKLGVLVNESFLVR
>gb|AKQ05896_1|ARO_3004587|tet(52)__uncultured_bacterium__TM_#07
NIMEQVKIASNMIVLQDYANPH
>gb|ABA81355_1|ARO_3004291|Rhodobacter_sphaeroides_ampC_beta-lactamase__Rhodobacter_sphaeroides_2_4_1__TM_#01
MKHLSPLSILLMVGALTPALAQDTTPSFESAAAAAFESVIEEH
>gb|ABA81355_1|ARO_3004291|Rhodobacter_sphaeroides_ampC_beta-lactamase__Rhodobacter_sphaeroides_2_4_1__TM_#02
GVLGMSYADASQTVIFPALGLKSTWIDVPTDAMG
>gb|ABA81355_1|ARO_3004291|Rhodobacter_sphaeroides_ampC_beta-lactamase__Rhodobacter_sphaeroides_2_4_1__TM_#03
TGTASPEVQT
>gb|ABA81355_1|ARO_3004291|Rhodobacter_sphaeroides_ampC_beta-lactamase__Rhodobacter_sphaeroides_2_4_1__TM_#04
YDFILQPQPVDEVDTTPDRR
>gb|CAL84423_1|ARO_3004146|cfrC__Clostridium_botulinum_A_str__ATCC_3502__TM_#01
EFGKNVSSVTPIFSQD
>gb|CAL84423_1|ARO_3004146|cfrC__Clostridium_botulinum_A_str__ATCC_3502__TM_#02
EAIIGLLRNRGS
>gb|ACL13298_1|ARO_3003110|catB10__Pseudomonas_aeruginosa__TM_#01
MTNYFESPFKGKL
>gb|ACL13298_1|ARO_3003110|catB10__Pseudomonas_aeruginosa__TM_#02
KMNWWDWPTEKIEEAMPLLCSSNIVGLHRYWQGFAV
>gb|ADP36409_1|ARO_3003730|Bifidobacteria_intrinsic_ileS_conferring_resistance_to_mupirocin__Bifidobacterium_bifidum_PRL2010__TM_#01
MSETTNSH
>gb|ADP36409_1|ARO_3003730|Bifidobacteria_intrinsic_ileS_conferring_resistance_to_mupirocin__Bifidobacterium_bifidum_PRL2010__TM_#02
VLTVPSANVAD
>gb|AAF40764_1|ARO_3003962|farB__Neisseria_meningitidis_MC58__TM_#01
YRETETVKM
>gb|AAF40764_1|ARO_3003962|farB__Neisseria_meningitidis_MC58__TM_#02
GQIAAAGSLSNFLRVL
>gb|AAF40764_1|ARO_3003962|farB__Neisseria_meningitidis_MC58__TM_#03
RFAEHITPYSATLHETAAHLSQHGVSDIQTL
>gb|AAR84672_1|ARO_3002925|vanRF__Paenibacillus_popilliae_ATCC_14706__TM_#01
EILFLLASNPNKVYR
>gb|AAT01092_1|ARO_3001773|OXA-61__Campylobacter_jejuni__TM_#01
MKKITLFLLFLNLVFGQDKILNNWFKEYN
>gb|AAT01092_1|ARO_3001773|OXA-61__Campylobacter_jejuni__TM_#02
SGTFVFYDGKTWASNDF
>gb|AAT01092_1|ARO_3001773|OXA-61__Campylobacter_jejuni__TM_#03
RRIGIKTMQE
>gb|AAT01092_1|ARO_3001773|OXA-61__Campylobacter_jejuni__TM_#04
QEAMNSVKE
>gb|AAT01092_1|ARO_3001773|OXA-61__Campylobacter_jejuni__TM_#05
ENKYKAFALNLDIDKFEDLYKREKILEKYLDELVKK
>gb|ACH58987_1|ARO_3002510|LRA-3__uncultured_bacterium_BLR3__TM_#01
MKSKLLLAAALAALAGTSMAAQAAELQYKP
>gb|ACH58987_1|ARO_3002510|LRA-3__uncultured_bacterium_BLR3__TM_#02
GDHVGDLAAFQKDAPAAKTYMNFRDAPTI
>gb|ACH58987_1|ARO_3002510|LRA-3__uncultured_bacterium_BLR3__TM_#03
EGRGFYPYHPVK
>gb|ACH58987_1|ARO_3002510|LRA-3__uncultured_bacterium_BLR3__TM_#04
TVQDGGRN

>gb|ACH58987_1|ARO_3002510|LRA-3__uncultured_bacterium_BLR3__TM_#05
ASTLKEQATYT
>gb|ACH58987_1|ARO_3002510|LRA-3__uncultured_bacterium_BLR3__TM_#06
LKIANATKA
>gb|AXA30439_1|ARO_3002459|LEN-9__Klebsiella_variicola__TM_#01
EMDLASGRTL
>gb|AXA30439_1|ARO_3002459|LEN-9__Klebsiella_variicola__TM_#02
YRQQDLVDYSPVSEKHL
>gb|AXA30439_1|ARO_3002459|LEN-9__Klebsiella_variicola__TM_#03
GELCAAAIT
>gb|AXA30439_1|ARO_3002459|LEN-9__Klebsiella_variicola__TM_#04
QLLQWMVDD
>gb|AXA30439_1|ARO_3002459|LEN-9__Klebsiella_variicola__TM_#05
GWFIADKTGA
>gb|AXA30439_1|ARO_3002459|LEN-9__Klebsiella_variicola__TM_#06
RGIVALLGP
>gb|AXA30439_1|ARO_3002459|LEN-9__Klebsiella_variicola__TM_#07
IGAALIEHWQR
>gb|ABO11759_2|ARO_3004573|Acinetobacter_baumannii_AbaF__Acinetobacter_baumannii_ATCC_17978__TM_#01
QLLEKQQLGIIEVL
>gb|ABO11759_2|ARO_3004573|Acinetobacter_baumannii_AbaF__Acinetobacter_baumannii_ATCC_17978__TM_#02
HLSDIFGRKPIYLIGAVLTAFWGFVGFPLMDTGNDWLIML
>gb|CAQ03505_1|ARO_3001338|SHV-100__Klebsiella_pneumoniae__TM_#01
RGARGIVA
>gb|BAF91108_1|ARO_3000847|KHM-1_beta-lactamase__Citrobacter_freundii__TM_#01
MKIALVISFGLLLFTNMVCADDSLPELDIQKIEDGVYLYTAYEKIE
>gb|BAF91108_1|ARO_3000847|KHM-1_beta-lactamase__Citrobacter_freundii__TM_#02
KWIDAQGFTAKASISTHFHTDSTG
>gb|BAF91108_1|ARO_3000847|KHM-1_beta-lactamase__Citrobacter_freundii__TM_#03
KNKGEEQATHSFG
>gb|BAF91108_1|ARO_3000847|KHM-1_beta-lactamase__Citrobacter_freundii__TM_#04
DASLLEKTRQRAVEALAAKK
>gb|APB03226_1|ARO_3003991|mphI__Paenibacillus_sp__LC231__TM_#01
KHGVNLIPE
>gb|APB03226_1|ARO_3003991|mphI__Paenibacillus_sp__LC231__TM_#02
GTGARWILRKPRRP
>gb|APB03226_1|ARO_3003991|mphI__Paenibacillus_sp__LC231__TM_#03
HENPGDGFIRSLAE
>gb|APB03226_1|ARO_3003991|mphI__Paenibacillus_sp__LC231__TM_#04
QEVRDETARNMEDIKSR
>gb|APB03226_1|ARO_3003991|mphI__Paenibacillus_sp__LC231__TM_#05
EEDSYWPT
>gb|APB03226_1|ARO_3003991|mphI__Paenibacillus_sp__LC231__TM_#06
ASPAKDFVLYYAIYGEHNLRVLLDRYEQAGGKVWPRMFDHIVEQHAAYPVLIAQFALLTG
QEEYMTMARNALGLTE
>gb|WP_071846200_1|ARO_3005350|DfrB9__Enterobacteriaceae__TM_#01
MNQSSNCISTPVVGQFALPFQP
>gb|ENU35234_1|ARO_3001734|OXA-279__Acinetobacter_parvus_DSM_16617___CIP_108168__TM_#01
MPKILKHLGLCASVMIGLTLLGCQNLQAPTQSAVSKKHDQTEIASLFQHAQTVGVFVTYD
GQTLQEYGNA
>gb|ENU35234_1|ARO_3001734|OXA-279__Acinetobacter_parvus_DSM_16617___CIP_108168__TM_#02
KQLTFDQSVQEQVKQMVLVD
>gb|ADI87853_1|ARO_3004666|pexA__uncultured_bacterium_Ak20-3__TM_#01
MKKFVLFCERNGIALIPIVLILLGCGLWPEMELLVPSLPDMQRAFNIQDAQIQQLLTANF
VGFLIGVLFAGPLCDSAGRRTVMMIGTIGYLVSSVLCPFCNDFVLLMIARFFQGLFMTGP
VIAGGVLLMEATEGVKQIFWMSIGNAAITFCMAAGPIVGSWINTGFGYVGNLWSILILGL
IGCLPALFLVPESLPVEKRAAFHPKLLFKGYFALLKDFRFMCLAIPMCALAAAYWIYVGV
SALYMVNQLGIAQEMFGRYQGPIVGCFSIISLGSSKLLQRFGLMKCLRAGIVSMFTGMLL
>gb|ACD35503_1|ARO_3000572|tet(42)__Micrococcus_sp__SMCC_G887__TM_#01
MTSPTSLTRRDQ
>gb|ACD35503_1|ARO_3000572|tet(42)__Micrococcus_sp__SMCC_G887__TM_#02
GNRITAIK
>gb|ACD35503_1|ARO_3000572|tet(42)__Micrococcus_sp__SMCC_G887__TM_#03
PAGETTGDAPAALVETAG
>gb|CAA39038_1|ARO_3002553|AAC(6_)-If__Enterobacter_cloacae__TM_#01
MDEASLSMWVGLRSQLWPDHSYEDHILDSQHILSCPDKYVSFLAINNQSQAIAFADAAVR
H
>gb|CAA39038_1|ARO_3002553|AAC(6_)-If__Enterobacter_cloacae__TM_#02
IPEQRGHGVAKLLVAAVQDWGVAKGCTEMASDAALDNHISY
>gb|AQX82857_1|ARO_3004185|mecD__Macrococcus_caseolyticus__TM_#01
MKNIKVKILIVCSLCLISFFLYNLLKENEIDKIFSSIENRNVDEINENITFLSRNTFSKK
QRYDRMNHIDNSLGIKKVNITDIKLLEEIVDTRKYSANMHYDSKFGKFTKKGYFEFEKNG
ESKRWELNWTPEVIIPGLTATNEVRVEELKSSRGEIVDRNGIPLAIDGEHYQVGIDPKNY
NKKDSKQIAKLLNINESTLKNKLKQSWVKDGVFVPIKSYVELSDEIKNKIPEYGLSVNKI
KGRTYPLKEASAHLLGYIGEINADELNDPKFKGYDSHSIVGKTGIEYMYDKELQNRDGLI
>gb|BAE06009_1|ARO_3003700|opmE__Pseudomonas_aeruginosa__TM_#01
ESLIQRALAANL
>gb|BAE06009_1|ARO_3003700|opmE__Pseudomonas_aeruginosa__TM_#02
MSWFQLQGIEAELAVVHDIAG
>gb|BAE06009_1|ARO_3003700|opmE__Pseudomonas_aeruginosa__TM_#03
TRNALAVLLAEAPQAFSPPVARASGERLTLR
>gb|BAE06009_1|ARO_3003700|opmE__Pseudomonas_aeruginosa__TM_#04
RGSIGLVAGNLDALDESGTSFNVLNPVIRWALLDRGRVWA
>gb|BAE06009_1|ARO_3003700|opmE__Pseudomonas_aeruginosa__TM_#05
RRCGVATDDTSPGVARQRDSRS
>gb|AGN74946|ARO_3003749|salA__Staphylococcus_sciuri_subsp__sciuri__TM_#01
MLFLFEEKALEVEHKVLIPELTFS
>gb|AGN74946|ARO_3003749|salA__Staphylococcus_sciuri_subsp__sciuri__TM_#02
KVIHQDQSVDSAMMEQDLTPYYDWTVMDYIIESYPEIAKIRL
>gb|AGN74946|ARO_3003749|salA__Staphylococcus_sciuri_subsp__sciuri__TM_#03
LQYEKQQKEQA
>gb|AGN74946|ARO_3003749|salA__Staphylococcus_sciuri_subsp__sciuri__TM_#04
QQLNRKLDQEHIPNPHKKEKTFSIQHHNFKSHYLVQFN
>gb|AGN74946|ARO_3003749|salA__Staphylococcus_sciuri_subsp__sciuri__TM_#05
FYIKRNQNVIVE
>gb|AGN74946|ARO_3003749|salA__Staphylococcus_sciuri_subsp__sciuri__TM_#06
TKGDITVHPELEIGYFSQDFENLNMHHTVLDEILE
>gb|AGN74946|ARO_3003749|salA__Staphylococcus_sciuri_subsp__sciuri__TM_#07
VLIVSHDNY
>gb|BAE06007_1|ARO_3003698|mexP__Pseudomonas_aeruginosa__TM_#01
MNLRHFIRITATLGVAALIAGCGESAPPGAASA
>gb|BAE06007_1|ARO_3003698|mexP__Pseudomonas_aeruginosa__TM_#02
QQLFLIDPRVFK

>gb|BAE06007_1|ARO_3003698|mexP__Pseudomonas_aeruginosa__TM_#03
TRGREGEQAPRVKVALLTDE
>gb|BAE06007_1|ARO_3003698|mexP__Pseudomonas_aeruginosa__TM_#04
RLAEIDGTPVDLSKTVGAAQ
>gb|ABA71732_1|ARO_3003069|vanXYG__Enterococcus_faecalis__TM_#01
MMKTIELEKEEIYCGNLLLVNKNYPLRDNNVKGLVPADIRFPNILMKRDVANVLQLIFEK
ISAGNSIVPVSGYRSLEEQTAIYDG
>gb|ABA71732_1|ARO_3003069|vanXYG__Enterococcus_faecalis__TM_#02
EFRRAAPDYGFTQRYAR
>gb|ABA71732_1|ARO_3003069|vanXYG__Enterococcus_faecalis__TM_#03
GFSLEEYTQFIKAYLEDNKYLFEQAHRAEIEIYYVPAKDDKTLIKIPENCVYQISGNNID
GFVVTIWRKTDD
>gb|ACJ41739_1|ARO_3000780|adeI__Acinetobacter_baumannii_AB0057__TM_#01
TLENAKASLLQQQANLA
>gb|ACJ41739_1|ARO_3000780|adeI__Acinetobacter_baumannii_AB0057__TM_#02
ETSGVQGQN
>gb|CAA66307_2|ARO_3000592|ErmN__Streptomyces_fradiae__TM_#01
MPSRPRTDSPHRHEGPAGPARLDRDEARRVWGQNFFRSAGSARRFARQLTGAESAGNDSV
TVEVGPGAGRITKELVRDGHPIVAVEVDPHWADRLAELELPNLTVVNDDFTTWPLPDGPL
RFIGNLPFG
>gb|CAA66307_2|ARO_3000592|ErmN__Streptomyces_fradiae__TM_#02
ALGPDRCREGVFLLQKQYTRKRTGAY
>gb|CAA66307_2|ARO_3000592|ErmN__Streptomyces_fradiae__TM_#03
RRGLGFPRQEFAPVPGSDTETLLVRSRPRPLAPWSRHAAYQRFVEDVFNTSRLTIGEAAR
ALDRRAGPGWLRGARVPPGLRVKDITAEQWADLFHACTPPPARRISPQRRR
>gb|AFN69318_1|ARO_3004658|catA8__Listeria_monocytogenes__TM_#01
EYFNHYFNQQTTYSVTKE
>gb|AFN69318_1|ARO_3004658|catA8__Listeria_monocytogenes__TM_#02
DITLLKSMIK
>gb|AFN69318_1|ARO_3004658|catA8__Listeria_monocytogenes__TM_#03
KGYELYPALI
>gb|AFN69318_1|ARO_3004658|catA8__Listeria_monocytogenes__TM_#04
NKVFRTGI
>gb|AFN69318_1|ARO_3004658|catA8__Listeria_monocytogenes__TM_#05
SFNSFYNSYK
>gb|AFN69318_1|ARO_3004658|catA8__Listeria_monocytogenes__TM_#06
YKDKNEMFPKK
>gb|AFN69318_1|ARO_3004658|catA8__Listeria_monocytogenes__TM_#07
IDFSSFNLNIGNNSR
>gb|ACH58980_1|ARO_3002482|LRA-1__uncultured_bacterium_BLR1__TM_#01
SAQLAALEA
>gb|ACH58980_1|ARO_3002482|LRA-1__uncultured_bacterium_BLR1__TM_#02
GDLLARRVNYSKGDLVSYSPITEK
>gb|AHE40505_1|ARO_3004542|mphN__Exiguobacterium_sp__S3-2__TM_#01
MALEIMEEHVSFQVPNWSIFDDE
>gb|AHE40505_1|ARO_3004542|mphN__Exiguobacterium_sp__S3-2__TM_#02
VEQQDYIWSFDKENTPQSYYQSLGKVLAELHTLPHRHFKEIGIKTLYARDLKSSMKIRME
KVRQKYHVNSE
>gb|AHE40505_1|ARO_3004542|mphN__Exiguobacterium_sp__S3-2__TM_#03
NKNFEVSGLIDWTEISIADTSVDFLSHL
>gb|AHE40505_1|ARO_3004542|mphN__Exiguobacterium_sp__S3-2__TM_#04
DLQDMRETAAHMLAQNV
>gb|AAA24786_1|ARO_3000368|vanC__Enterococcus_gallinarum__TM_#01
MKKIAVLFGGNSPEYSVSLTSAASVIQAIDPLKYEVMTIGIAPTMDWYWYQGNLANVRND
TWLEDHKNCHQLTFSSQGFILGEKRIV
>gb|AAA24786_1|ARO_3000368|vanC__Enterococcus_gallinarum__TM_#02
LADTMGIASAPTLLLSRYENDPATIDRFIQD
>gb|AAA24786_1|ARO_3000368|vanC__Enterococcus_gallinarum__TM_#03
DKTALQSALTTAFAYGSTVLIQKA
>gb|AAA24786_1|ARO_3000368|vanC__Enterococcus_gallinarum__TM_#04
TITVPAPLPLALESQIKEQAQLLYRNLGLTGLARIDFFVTNQ
>gb|AAA24786_1|ARO_3000368|vanC__Enterococcus_gallinarum__TM_#05
EVGLSYEILVEQLIALAEEDKR
>gb|AAA99505_1|ARO_3002988|bcrB__Bacillus_licheniformis__TM_#01
MAKKAKYPDVPIRFSETFSDTNLYIVLL
>gb|AAA99505_1|ARO_3002988|bcrB__Bacillus_licheniformis__TM_#02
LFGLIGQFEGLSSAVLIEGFKQFMIGGALLFFLVS
>gb|AAA99505_1|ARO_3002988|bcrB__Bacillus_licheniformis__TM_#03
IIISMVSIMVYGTEYSALFPWSAVWVIASGTFFPEYPPEYSFISVA
>gb|ADZ12700_1|ARO_3005090|RanB__Riemerella_anatipestifer_RA-GD__TM_#01
MLKLSKRMLTSVGEYVLLLGKVIKRPQKHN
>gb|ADZ12700_1|ARO_3005090|RanB__Riemerella_anatipestifer_RA-GD__TM_#02
YGGYLAGVATGNWTEADYITGIQRYIPD
>gb|CAI43424_1|ARO_3002415|OXY-6-3__Klebsiella_oxytoca__TM_#01
AAAAVPLLLAS
>gb|CAI43424_1|ARO_3002415|OXY-6-3__Klebsiella_oxytoca__TM_#02
SGGRLGVALINT
>gb|CAI43424_1|ARO_3002415|OXY-6-3__Klebsiella_oxytoca__TM_#03
VVNKRLEI
>gb|CAI43424_1|ARO_3002415|OXY-6-3__Klebsiella_oxytoca__TM_#04
YLGGPEKVTAFA
>gb|CAI43424_1|ARO_3002415|OXY-6-3__Klebsiella_oxytoca__TM_#05
TPLAMAESLRKLTLG
>gb|AAP26981_1|ARO_3005045|SatA__Bacillus_anthracis_str__Ames__TM_#01
LQNDNEELV
>gb|AAP26981_1|ARO_3005045|SatA__Bacillus_anthracis_str__Ames__TM_#02
TVDKKYRTLGVGKRLIAQAKQWAKE
>gb|AAC61671_1|ARO_3002842|vatC__Staphylococcus_cohnii__TM_#01
MKWQNQQGPNPEEIYPIEGNKHVQFIKPS
>gb|AAC61671_1|ARO_3002842|vatC__Staphylococcus_cohnii__TM_#02
SQVLYHYEL
>gb|AAC61671_1|ARO_3002842|vatC__Staphylococcus_cohnii__TM_#03
SRLIKIRFSKEKIAALLKVRWWDLEIETINENIDCILNGDIKKVKRS
>gb|KIX81495_1|ARO_3002813|lmrB__Bacillus_subtilis__TM_#01
METTAKASQQ
>gb|KIX81495_1|ARO_3002813|lmrB__Bacillus_subtilis__TM_#02
LALVFGIAYMQNVSET
>gb|KIX81495_1|ARO_3002813|lmrB__Bacillus_subtilis__TM_#03
PTVIVSLIVGVVG
>gb|KIX81495_1|ARO_3002813|lmrB__Bacillus_subtilis__TM_#04
SNVTTTSTA
>gb|KIX81495_1|ARO_3002813|lmrB__Bacillus_subtilis__TM_#05
TVKNPADPAVIPQALTAGVQHAFVFAMIVAIIGLIGAFFMKRVKVDH

```
>gb|AGW16411_1|ARO_3001517|OXA-329__Acinetobacter_calcoaceticus__TM_#01
SLFDQAQT
>gb|AGW16411_1|ARO_3001517|OXA-329__Acinetobacter_calcoaceticus__TM_#02
GVLVIKRG
>gb|AGW16411_1|ARO_3001517|OXA-329__Acinetobacter_calcoaceticus__TM_#03
VQEQVQSM
>gb|AGW16411_1|ARO_3001517|OXA-329__Acinetobacter_calcoaceticus__TM_#04
AFSLNLEM
>gb|AGW16411_1|ARO_3001517|OXA-329__Acinetobacter_calcoaceticus__TM_#05
YKGLEQLG
>gb|AAG04518_1|ARO_3000149|fosA__Pseudomonas_aeruginosa_PAO1__TM_#01
EPQYGGPAA
>gb|AAG04518_1|ARO_3000149|fosA__Pseudomonas_aeruginosa_PAO1__TM_#02
AADFARFAAQLRAHG
>gb|CTQ57092_1|ARO_3004682|aadA27__Acinetobacter_lwoffii__TM_#01
LKEQILSGS
>gb|CTQ57092_1|ARO_3004682|aadA27__Acinetobacter_lwoffii__TM_#02
NKKSSLIK
>gb|AJE60855_1|ARO_3003209|FosA5__Escherichia_coli__TM_#01
PSVAFYQQL
>gb|AJE60855_1|ARO_3003209|FosA5__Escherichia_coli__TM_#02
SFAARLEAAGVA
>gb|AJE60855_1|ARO_3003209|FosA5__Escherichia_coli__TM_#03
WKLNRSEGASH
>gb|CAD58780_1|ARO_3001794|OXA-45__Pseudomonas_aeruginosa__TM_#01
MRGKHTVILGAALSALFAGAAGAQMLECTLVADAASGQELYRKGACDKAFA
>gb|CAD58780_1|ARO_3001794|OXA-45__Pseudomonas_aeruginosa__TM_#02
KRFAAYVAGFGYGNGDISGEPGKS
>gb|CAD58780_1|ARO_3001794|OXA-45__Pseudomonas_aeruginosa__TM_#03
KISPEGQVRFVRDLLSAKLPASKDAQQMTVSILPHFAAGDWAVQGKTGTGSFIDARGAKA
PLGWFIGWATHEERRVVFARMTAGGKKGEQPAGPAARDAFLKALPDLAKRF
>gb|BAA12910_1|ARO_3000318|mphB__Escherichia_coli__TM_#01
EGLEALIL
>gb|CAA71947_1|ARO_3002157|FOX-3__Klebsiella_oxytoca__TM_#01
LGSLLLAPCTYA
>gb|CAA71947_1|ARO_3002157|FOX-3__Klebsiella_oxytoca__TM_#02
HAPWLKGSA
>gb|CAA71947_1|ARO_3002157|FOX-3__Klebsiella_oxytoca__TM_#03
WSPVYPAGTHRQY
>gb|CAA71947_1|ARO_3002157|FOX-3__Klebsiella_oxytoca__TM_#04
LAANSLGQ
>gb|CAA71947_1|ARO_3002157|FOX-3__Klebsiella_oxytoca__TM_#05
GLHHTYIQVPESA
>gb|CAA71947_1|ARO_3002157|FOX-3__Klebsiella_oxytoca__TM_#06
NYAYGYSKE
>gb|CAA71947_1|ARO_3002157|FOX-3__Klebsiella_oxytoca__TM_#07
IALTHTGF
>gb|CAA71947_1|ARO_3002157|FOX-3__Klebsiella_oxytoca__TM_#08
MTQGLGWESY
>gb|CAA71947_1|ARO_3002157|FOX-3__Klebsiella_oxytoca__TM_#09
AILSQLAE
>gb|AAN00989_1|ARO_3003774|Streptococcus_agalactiae_mprF__Streptococcus_agalactiae_2603V_R__TM_#01
MLKKLIEKVKSLTSVIKIVFFISVLVLIIVEMIHLKRTISVEQLKSVFGQLSPMNLFLII
LVGVIAVLPTTGYDFVLNGLLR
>gb|AAN00989_1|ARO_3003774|Streptococcus_agalactiae_mprF__Streptococcus_agalactiae_2603V_R__TM_#02
VIALIMSHIFHAKASVDYYYLVLIGASMYFPVIYWISGHKGSHYFGDMPSSTRIK
>gb|AAN00989_1|ARO_3003774|Streptococcus_agalactiae_mprF__Streptococcus_agalactiae_2603V_R__TM_#03
YLMGIHLPVYKILPLFC
>gb|AAN00989_1|ARO_3003774|Streptococcus_agalactiae_mprF__Streptococcus_agalactiae_2603V_R__TM_#04
HYLGSQINQRYENVPKELVSTVLQTMVSHLMRILGAFLIFSTAFFENITYIMWLQKLGLD
PLQEQMLWQFPGLLLGVCFILLARTIDQKVKNAFPIAIIWITLTLFYLNLGHISWRLSFW
FILLLLGLLVIKPTLYKKQFIYSWEE
>gb|WP_197749399_1|ARO_3005349|DfrA38__Acinetobacter_baumannii__TM_#01
MNCCIVVGIGR
>gb|WP_197749399_1|ARO_3005349|DfrA38__Acinetobacter_baumannii__TM_#02
KAEGALVIHDWSELEQHLSADKTCFIIGGSEIFKQALDAGLVNEMYITHIDATFEGADVF
FPYVNWENWTEEDILHYTKDEKNPYSFTIKKYSK
>gb|AFI56422_1|ARO_3004777|CMH-1__Enterobacter_cloacae__TM_#01
PGTTRLYAN
>gb|AFI56422_1|ARO_3004777|CMH-1__Enterobacter_cloacae__TM_#02
LAVKPSGM
>gb|AFI56422_1|ARO_3004777|CMH-1__Enterobacter_cloacae__TM_#03
RVFKPLKL
>gb|CCP42991_1|ARO_3002525|AAC(2_)-Ic__Mycobacterium_tuberculosis_H37Rv__TM_#01
DIRQMVTG
>gb|CCP42991_1|ARO_3002525|AAC(2_)-Ic__Mycobacterium_tuberculosis_H37Rv__TM_#02
TVFVLPID
>gb|BAD59497_1|ARO_3000501|Nocardia_rifampin_resistant_beta-subunit_of_RNA_polymerase_(rpoB2)__Nocardia_farcinica_IFM_10152__TM_#01
MLEGRIVTVSSRTESPLAAPGVPGAPRRLSFARIREPLAVPGLLDIQTESFGWLIGAPDW
CARAAARGTEPVA
>gb|BAD59497_1|ARO_3000501|Nocardia_rifampin_resistant_beta-subunit_of_RNA_polymerase_(rpoB2)__Nocardia_farcinica_IFM_10152__TM_#02
APLIMASLAKDNV
>gb|BAD59497_1|ARO_3000501|Nocardia_rifampin_resistant_beta-subunit_of_RNA_polymerase_(rpoB2)__Nocardia_farcinica_IFM_10152__TM_#03
AHGVDDDHERTAANLGINLSRAESITETELSG
>gb|EGP45231_1|ARO_3004144|AxyY__Achromobacter_insuavis_AXX-A__TM_#01
AAMRIWPDPAKLTALSLTPGDIVSALRSHNARVTIGELGNQAVPKDAPLNASIVAGESLH
TPEQFANIP
>gb|EGP45231_1|ARO_3004144|AxyY__Achromobacter_insuavis_AXX-A__TM_#02
RDASQHVDAVVKRINKAFADRKNLM
>gb|EGP45231_1|ARO_3004144|AxyY__Achromobacter_insuavis_AXX-A__TM_#03
KLLAKAAE
>gb|EGP45231_1|ARO_3004144|AxyY__Achromobacter_insuavis_AXX-A__TM_#04
MVQADGKRRVDVD
>gb|EGP45231_1|ARO_3004144|AxyY__Achromobacter_insuavis_AXX-A__TM_#05
RGHSSGEAMRAMETLAAELPR
>gb|EGP45231_1|ARO_3004144|AxyY__Achromobacter_insuavis_AXX-A__TM_#06
VGMRARPARPDGREPLETTP
>gb|AAR21614_1|ARO_3002529|AAC(3)-Id__Salmonella_enterica_subsp__enterica_serovar_Newport__TM_#01
MSVEIIHLTG
>gb|AAR21614_1|ARO_3002529|AAC(3)-Id__Salmonella_enterica_subsp__enterica_serovar_Newport__TM_#02
EAFNDQDSYARNKPSSSYLQKLLSTSSFIALAAVDEQKVIGAI
>gb|AAR21614_1|ARO_3002529|AAC(3)-Id__Salmonella_enterica_subsp__enterica_serovar_Newport__TM_#03
VEDQPAIELYK
```

```
>gb|AAL02176_1|ARO_3004669|emtA__Enterococcus_faecium__Partial_TM_#01
KAMEGINKR
>gb|AAL02176_1|ARO_3004669|emtA__Enterococcus_faecium__Partial_TM_#02
ESKDVFVDF
>gb|CAB96921_1|ARO_3000606|FEZ-1_beta-lactamase__Fluoribacter_gormanii__TM_#01
MKKVLSLTALMMVLNHSSFAYPMPN
>gb|CAB96921_1|ARO_3000606|FEZ-1_beta-lactamase__Fluoribacter_gormanii__TM_#02
VILSGGKSDFHYANDSSTYFTQSTVDKVLHDGERVE
>gb|CAB96921_1|ARO_3000606|FEZ-1_beta-lactamase__Fluoribacter_gormanii__TM_#03
LKDHGKQYQAVIIGSIGVNPGYKLVDNITYPKIAEDYKHSIKVLESMRCDIFLGSHAGMF
DLKNKYVLLQKGQNNPFVDPTGCKNYIEQKANDFYTELKKQETA
>gb|CAX52582_1|ARO_3003324|Bacillus_subtilis_mprF__Bacillus_subtilis_subsp__subtilis_str__168__TM_#01
MVINGLERTDLFMLVLIGLLAVAAMSLYDYVLKY
>gb|CAX52582_1|ARO_3003324|Bacillus_subtilis_mprF__Bacillus_subtilis_subsp__subtilis_str__168__TM_#02
MMFYKEHTKDH
>gb|CAX52582_1|ARO_3003324|Bacillus_subtilis_mprF__Bacillus_subtilis_subsp__subtilis_str__168__TM_#03
FVAARVLPVDEVIHEKPWLWAVVIGFALILPLSLAVSKIKDRKAGDEENADKVKNPIFAY
IGASVVEWLMAGTVIYFALFAMGIHADIRY
>gb|CAX52582_1|ARO_3003324|Bacillus_subtilis_mprF__Bacillus_subtilis_subsp__subtilis_str__168__TM_#04
QLGYHQEAIV
>gb|CAX52582_1|ARO_3003324|Bacillus_subtilis_mprF__Bacillus_subtilis_subsp__subtilis_str__168__TM_#05
LETNPRIAPAIETTNVLLVVQRAVLVRILQ
>gb|CAX52582_1|ARO_3003324|Bacillus_subtilis_mprF__Bacillus_subtilis_subsp__subtilis_str__168__TM_#06
KKVLRHEYFVHS
>gb|CAX52582_1|ARO_3003324|Bacillus_subtilis_mprF__Bacillus_subtilis_subsp__subtilis_str__168__TM_#07
VVYHKRTKPIGEKADPERLAA
>gb|CAX52582_1|ARO_3003324|Bacillus_subtilis_mprF__Bacillus_subtilis_subsp__subtilis_str__168__TM_#08
HPPFSDAFLE
>gb|CAX52582_1|ARO_3003324|Bacillus_subtilis_mprF__Bacillus_subtilis_subsp__subtilis_str__168__TM_#09
PSYLQKAPIAYMKNAEGE
>gb|AAA22562_1|ARO_3002878|BcII__Bacillus_cereus__TM_#01
PFVSTISSVQAER
>gb|AAA22562_1|ARO_3002878|BcII__Bacillus_cereus__TM_#02
KTLKERGIKAHST
>gb|AAA22562_1|ARO_3002878|BcII__Bacillus_cereus__TM_#03
LTAELAKKNGYEEPLGDLQ
>gb|AAA22562_1|ARO_3002878|BcII__Bacillus_cereus__TM_#04
GGCLVKSAS
>gb|AAA22562_1|ARO_3002878|BcII__Bacillus_cereus__TM_#05
DAYVNEWSTSIENVLKRY
>gb|AAA22562_1|ARO_3002878|BcII__Bacillus_cereus__TM_#06
GLLLHTLDLLK
>gb|AAC73788_1|ARO_3003841|kdpE__Escherichia_coli_str__K-12_substr__MG1655__TM_#01
APDPLVKFSDVTVDLAARV
>gb|AEA07977_1|ARO_3004641|aacA43__Klebsiella_pneumoniae__TM_#01
MIYNIINIADSEKNKEDAARILYSAFRGKGKDAWPTLDSAREEIAECIASPNICLGITLD
DRLVGWGGLRPMYET
>gb|AEA07977_1|ARO_3004641|aacA43__Klebsiella_pneumoniae__TM_#02
DPDYQGNGLGRLLLSKIESTATTNR
>gb|AEA07977_1|ARO_3004641|aacA43__Klebsiella_pneumoniae__TM_#03
TLSTSLSMTDIDESNIFQEIKNII
>gb|CAB60001_1|ARO_3000326|ErmE__Saccharopolyspora_erythraea_NRRL_2338__TM_#01
MSSSDEQPRPRRRN
>gb|CAB60001_1|ARO_3000326|ErmE__Saccharopolyspora_erythraea_NRRL_2338__TM_#02
TAELRPDLP
>gb|CAB60001_1|ARO_3000326|ErmE__Saccharopolyspora_erythraea_NRRL_2338__TM_#03
EPPPEPFA
>gb|CAB60001_1|ARO_3000326|ErmE__Saccharopolyspora_erythraea_NRRL_2338__TM_#04
EFVEKVDRRLFKPVPKVDSAIMRLRRRAEPLLEGAALERYESMVELCFTGVGGNIQASLL
RKYPRRRVEAALDHAGVGGGAVVAYVRPEQWLRLFERLDQKNEPRGGQPQRGRRTGGRDH
GDRRTGGQDRGDRRTGGRDHRDRQASGHGDRRSSGRNRDDGRTGEREQGDQGGRRGPSGG
GRTGGRPGRRGGPGQR
>gb|QFQ03949_1|ARO_3005051|LpsB__Acinetobacter_baumannii__TM_#01
SELQNTIQSKG
>gb|QFQ03949_1|ARO_3005051|LpsB__Acinetobacter_baumannii__TM_#02
EAENEKALLEIVKHHIEQPQT
>gb|AAD09860_1|ARO_3000561|tet(30)__Agrobacterium_fabrum_str__C58__TM_#01
FVALFVLPESRKAGPGKFA
>gb|AAD09860_1|ARO_3000561|tet(30)__Agrobacterium_fabrum_str__C58__TM_#02
LPLFATVKTPKAVAA
>gb|OPG86592_1|ARO_3004626|Erm(49)__Bifidobacterium_breve__TM_#01
HKARKVVAIEFDEELYEKLKNKFQSNNKVDIIYGDILNYTPRIPSYC
>gb|OPG86592_1|ARO_3004626|Erm(49)__Bifidobacterium_breve__TM_#02
NPYGAETLRSMLYKPFFDMDLKYRFDPSDFKPAPQARIVLASFERKQFPDVKKEEEKLYK
>gb|OPG86592_1|ARO_3004626|Erm(49)__Bifidobacterium_breve__TM_#03
QLVVNSFNNMNK
>gb|OPG86592_1|ARO_3004626|Erm(49)__Bifidobacterium_breve__TM_#04
SNRKRKPYHRNNV
>gb|AAT68578_1|ARO_3004803|GOB-11__Elizabethkingia_meningoseptica__TM_#01
AQVVKEPENM
>gb|AAT68578_1|ARO_3004803|GOB-11__Elizabethkingia_meningoseptica__TM_#02
GKYGVTFKP
>gb|AAT68578_1|ARO_3004803|GOB-11__Elizabethkingia_meningoseptica__TM_#03
TLKDQDKI
>gb|AAT68578_1|ARO_3004803|GOB-11__Elizabethkingia_meningoseptica__TM_#04
IVDKKFSEV
>gb|AAT68578_1|ARO_3004803|GOB-11__Elizabethkingia_meningoseptica__TM_#05
IQSDYAYTF
>gb|AAT68578_1|ARO_3004803|GOB-11__Elizabethkingia_meningoseptica__TM_#06
YNPQLFMDK
>gb|AEP40504_1|ARO_3002940|vanSN__Enterococcus_faecium__TM_#01
MKNKLNDPLIKRILLRYVSTVLLAIGIYGGVLLLLLFLFRLRTWYGDEPFYLFLRTLYIR
FNLIGLVSSGAFLLLLMITLVYIFKLIGYLNETITATKQLLEAPEQRIQLSTELFT
>gb|AEP40504_1|ARO_3002940|vanSN__Enterococcus_faecium__TM_#02
NNNQANRAAKV
>gb|AEP40504_1|ARO_3002940|vanSN__Enterococcus_faecium__TM_#03
TELRAKYTDIALDKALRLEELIGEFFEVTQFNLTKLTI
>gb|AEP40504_1|ARO_3002940|vanSN__Enterococcus_faecium__TM_#04
NEKGLKWQLAIDKGIKAEVDPNKMG
>gb|AEP40504_1|ARO_3002940|vanSN__Enterococcus_faecium__TM_#05
SNSTIHLSLEKNGQNLELKITNETHTLPEEKLTQIFEP
>gb|AEP40504_1|ARO_3002940|vanSN__Enterococcus_faecium__TM_#06
SGGRIHAQSSNNQMTFTLTLPISE
```

>gb|AAD51346_1|ARO_3003053|smeC__Stenotrophomonas_maltophilia__TM_#01
SGSVTRSRVSDANSETGV
>gb|AAD51346_1|ARO_3003053|smeC__Stenotrophomonas_maltophilia__TM_#02
EQSLAFTQQTLDSQHQTLQRTEARHAQGLASGLDLSQVQTSVEAARGALAKLQAQQAQDR
DALQLLVGAPLDPALLPTAQALDGSVALAP
>gb|AAD51346_1|ARO_3003053|smeC__Stenotrophomonas_maltophilia__TM_#03
YGHSSTALSTLFSAGTRG
>gb|AAD51346_1|ARO_3003053|smeC__Stenotrophomonas_maltophilia__TM_#04
QAFSEVADALATRDHLTAQLDAQRALVAD
>gb|AAF94716_1|ARO_3004289|Vibrio_cholerae_varG__Vibrio_cholerae_O1_biovar_El_Tor_str__N16961__TM_#01
MFVSHLSFPHLIEERKMKLSTLALAPITAALL
>gb|AAF94716_1|ARO_3004289|Vibrio_cholerae_varG__Vibrio_cholerae_O1_biovar_El_Tor_str__N16961__TM_#02
QSITNKPITTIMYSHYHLDHLGGGNQLVDLIKKNYPKVDKIRVIASQTVADKINQHAEVG
ENGVKTPKVPAPNDIYDLT
>gb|BAA23130_1|ARO_3002493|SRT-1__Serratia_marcescens__TM_#01
TAQAAQQQDIDAVIQPLMK
>gb|BAA23130_1|ARO_3002493|SRT-1__Serratia_marcescens__TM_#02
QQGKLSFN
>gb|BAA23130_1|ARO_3002493|SRT-1__Serratia_marcescens__TM_#03
QAMEQGMLPALGMSHTYVQVPAAQMANYAQ
>gb|CAA36304_1|ARO_3000251|msrA__Staphylococcus_epidermidis__TM_#01
EGIDCSPKV
>gb|CAA36304_1|ARO_3000251|msrA__Staphylococcus_epidermidis__TM_#02
KHVSDKKWELTGQSIHDIT
>gb|AAF01499_1|ARO_3000193|tetT__Streptococcus_pyogenes__TM_#01
KEGIQVQTKVIFNTLV
>gb|AAF01499_1|ARO_3000193|tetT__Streptococcus_pyogenes__TM_#02
KGVCLDEIYTQIQE
>gb|AAF01499_1|ARO_3000193|tetT__Streptococcus_pyogenes__TM_#03
TKSYNTEDLLSAYVYKIDRDEK
>gb|AAF01499_1|ARO_3000193|tetT__Streptococcus_pyogenes__TM_#04
CDLSKRSKLIE
>gb|AAF01499_1|ARO_3000193|tetT__Streptococcus_pyogenes__TM_#05
RKMRASIEDIIAKGEETTLSGKIPVDTSKSYQSELLSYSNGKGIFITEPYGYDIYNDKPI
INDIGNDN
>gb|AIT76110_1|ARO_3002239|IMP-48__Pseudomonas_aeruginosa__TM_#01
TKHGLVVLVKN
>gb|AIT76110_1|ARO_3002239|IMP-48__Pseudomonas_aeruginosa__TM_#02
IKGSISTHFHG
>gb|ENV18923_1|ARO_3001731|OXA-275__Acinetobacter_guillouiae_NIPH_991__TM_#01
KIKSSVLIILSSVAFSGCVSNANLHDPASSQRTSEIPLLFNY
>gb|AEY83581|ARO_3000510|mupB__Staphylococcus_aureus__TM_#01
MENENIIEEQKILNFWKEEN
>gb|AEY83581|ARO_3000510|mupB__Staphylococcus_aureus__TM_#02
KDKNEIEK
>gb|AEY83581|ARO_3000510|mupB__Staphylococcus_aureus__TM_#03
MYEKQWREFSELIGYWVDMEKPYKTMDNTYIESIWYILSD
>gb|AEY83581|ARO_3000510|mupB__Staphylococcus_aureus__TM_#04
ILKFPILDSDEN
>gb|AEY83581|ARO_3000510|mupB__Staphylococcus_aureus__TM_#05
AEEIYVKVNYDNEIFIIMESLLQSVFKDEDNIDIVSKHKGKEFVGKEYLAPFPNKSLMNN
ENSYKVLPADFVTN
>gb|AEY83581|ARO_3000510|mupB__Staphylococcus_aureus__TM_#06
IPFINVIDSRGKYNQDSPIFKGELAKESDINIIKELTHL
>gb|AEY83581|ARO_3000510|mupB__Staphylococcus_aureus__TM_#07
YKNEIKENNQKIE
>gb|AEY83581|ARO_3000510|mupB__Staphylococcus_aureus__TM_#08
NMIDWNIG
>gb|AEY83581|ARO_3000510|mupB__Staphylococcus_aureus__TM_#09
TCSHEFSPKSINDLIQHSIEDIPSDIELHRPYIDNVKCKCQNCGGDMC
>gb|AEY83581|ARO_3000510|mupB__Staphylococcus_aureus__TM_#10
FKGEAPYKN
>gb|AEY83581|ARO_3000510|mupB__Staphylococcus_aureus__TM_#11
KTYGADSLRWTLVSDSVPWTNKRFSENMV
>gb|ABX54690_1|ARO_3002974|vanTrL__Enterococcus_faecalis__TM_#01
MVENKMRAYKEFYVESLLHNVQVIKKNIPKSTKIMAVVKANAYGINAVNVAIILEYI
>gb|ABX54690_1|ARO_3002974|vanTrL__Enterococcus_faecalis__TM_#02
TSNILILGYTTPTKVDDLIHYE
>gb|ABX54690_1|ARO_3002974|vanTrL__Enterococcus_faecalis__TM_#03
FLNKTGKKIM
>gb|ABX54690_1|ARO_3002974|vanTrL__Enterococcus_faecalis__TM_#04
LEEICPIFNYPFLKIK
>gb|ABX54690_1|ARO_3002974|vanTrL__Enterococcus_faecalis__TM_#05
LSEEGKQRTIKQISRYNTIIAELKRKRVDV
>gb|ABX54690_1|ARO_3002974|vanTrL__Enterococcus_faecalis__TM_#06
NDHNVKLHLDLQPVVAVKAQLISKKKIAPGEYIGYGTDTQLTSSKT
>gb|ABX54690_1|ARO_3002974|vanTrL__Enterococcus_faecalis__TM_#07
EYCVVFEDKQIPQIGR
>gb|ABX54690_1|ARO_3002974|vanTrL__Enterococcus_faecalis__TM_#08
NCSDIPLGVMVDVLPNIEEISQIQSTITNEIISCLGSRLGMEVK
>gb|AAC44793_1|ARO_3002524|AAC(2_)-Ib__Mycolicibacterium_fortuitum__TM_#01
MPFQDVSAP
>gb|AAA26779_1|ARO_3004652|Erm(O)-lrm__Streptomyces_lividans__TM_#01
PAPRVDAG
>gb|APU52409_1|ARO_3004085|lnuG__Enterococcus_faecalis__TM_#01
MLKQKELMARVKELVQSDE
>gb|APU52409_1|ARO_3004085|lnuG__Enterococcus_faecalis__TM_#02
TISTFDSAKWLN
>gb|APU52409_1|ARO_3004085|lnuG__Enterococcus_faecalis__TM_#03
FNMTKNLEKEISPENYEKFKKTTARLNELELYEAYKNSLLLVMELRNLVEKQYQLTISDD
FLGKLFNYMNE
>gb|BAE06005_1|ARO_3003704|mexM__Pseudomonas_aeruginosa__TM_#01
IVARVERRDVEQQV
>gb|BAE06005_1|ARO_3003704|mexM__Pseudomonas_aeruginosa__TM_#02
QLTRLLVSEGQMVEAGELLATIDDRAVVAALEQAQASRASNQAQLKSAEQDLQRYRSLYA
ERAVSRQLLDQQQATVDQLRATLKANDATINAERVRLSYTRITSP
>gb|BAE06005_1|ARO_3003704|mexM__Pseudomonas_aeruginosa__TM_#03
DGGSALGEGRLLTIDNQIDSSTGTIRVRASFDNRQARLWPGQFVAVSLHTGVRRDQ
>gb|BAE06005_1|ARO_3003704|mexM__Pseudomonas_aeruginosa__TM_#04
VADDRVEAVPVRVLQDIDGLSVVEGLASGDQVVVDGHSRLMPGALVDIQEPRPSLAQATE
RRP
>gb|AAC44316_1|ARO_3005010|qacEdelta1__Pseudomonas_aeruginosa__TM_#01
VIITAIAWLL

>gb|ACX92986_2|ARO_3002832|vgaD__Enterococcus_faecium__TM_#01
MLILEANHIEKSINDRKLLDVTHLQIHY
>gb|ACX92986_2|ARO_3002832|vgaD__Enterococcus_faecium__TM_#02
HNIKELEKHFS
>gb|ACX92986_2|ARO_3002832|vgaD__Enterococcus_faecium__TM_#03
GEVKEIHGNYTSYVKQKELLRRQQQEEYEKYITKKKQLERAVTMKEQKAQ
>gb|ACX92986_2|ARO_3002832|vgaD__Enterococcus_faecium__TM_#04
KKPKDYPAVKMKLSNQDQIQGRNVLRVKDLSVSFGNHVLWTDASFT
>gb|ACX92986_2|ARO_3002832|vgaD__Enterococcus_faecium__TM_#05
EGAVLFVSHDCRFVQNIASKIIELSDQKVIEFLGSYKAFRERSQETERDYMKEELLKIEI
KLTQMISEMNDEASNELEKEFQMLIHERNQLRNQVNN
>gb|ABA71726_1|ARO_3004253|vanUG__Enterococcus_faecalis__TM_#01
MRVSYNKLWKLLIDRDMKKGELREAVGVSKSTFAKLGKNENVSLTVLLAICEYLNCDFGD
IIEALPETPDKERDS
>gb|AAK63223_1|ARO_3003561|Sed1_beta-lactamase__Citrobacter_sedlakii__TM_#01
MLKERFRQTVFIAAAVMPFIFSSTSLHAQATSDVQQV
>gb|AAX14803_1|ARO_3000782|adeK__Acinetobacter_baumannii__TM_#01
TQPVKRIAQQN
>gb|AAX14803_1|ARO_3000782|adeK__Acinetobacter_baumannii__TM_#02
VHQPSSAELKKQ
>gb|AAF94731_1|ARO_3004364|almG__Vibrio_cholerae_O1_biovar_El_Tor_str__N16961__TM_#01
MRIFWLKALTSGLFYLPLSIKNGLCRLVAKPISRKKMAAALTQLNYALPDLGDVKKQAIV
EQSTRLSLKNLLGFCHLKRYQYQVEQPDLVQEILDNQG
>gb|AAF94731_1|ARO_3004364|almG__Vibrio_cholerae_O1_biovar_El_Tor_str__N16961__TM_#02
WLNQQGKKAVTIFGAGSSGDRPDENAMISQAAKLAGVPYLLRKQNLMLELAQRIKQGEWV
VLHTDMRTEGVPVRWFGQATQLSATPFFLAHKLACPIYFHYALSEGMTQRLHFSRFALHQ
TDDLSRNIAQDAQQLADMMQQAITAHPEQWIWLYRRFK
>gb|AUW34359_1|ARO_3004445|RSA-2__uncultured_bacterium__TM_#01
MIKKIISGACLVLLAGCVLGVKPAGKETGFMIDSGRFAGMDGCAIVFDARMGKIAGVYGE
KRCKERVT
>gb|AUW34359_1|ARO_3004445|RSA-2__uncultured_bacterium__TM_#02
SDESTVLKWDGVQWPFDSWNQDQTAA
>gb|AUW34359_1|ARO_3004445|RSA-2__uncultured_bacterium__TM_#03
PMLGLEKIKAYLKA
>gb|AUW34359_1|ARO_3004445|RSA-2__uncultured_bacterium__TM_#04
SGLTSAWLTITKSDTNPDKGSLKISAYEELEFFRRFWRGALPVSGAAVEKTKKMIYLETS
PGGYALHGKTGSGYLDGLTGDFGWFAGHVEGKGREYFVVTAVTRNGNAADARIPGLVAKE
LAKNILKDNSVW
>gb|AAM09849_1|ARO_3000005|vanD__Enterococcus_faecium__TM_#01
KDGGYESQPV
>gb|AAM09849_1|ARO_3000005|vanD__Enterococcus_faecium__TM_#02
EVPGFRVLQKGDSLEAE
>gb|AAM09849_1|ARO_3000005|vanD__Enterococcus_faecium__TM_#03
TEAGKYDSKILVEEAVSGSEVGCAILGNGND
>gb|AAM09849_1|ARO_3000005|vanD__Enterococcus_faecium__TM_#04
ALPDEVREQIQE
>gb|AAA86871_1|ARO_3002841|vatB__Staphylococcus_aureus__TM_#01
MKYGPDPNSIYPHEEIKS
>gb|AAA86871_1|ARO_3002841|vatB__Staphylococcus_aureus__TM_#02
QIKWWDWSAQKIFSNLETLCSSDLEKIKSIRD
>gb|ADB56658_1|ARO_3004753|BIC-1__Pseudomonas_fluorescens__TM_#01
MARPSKLALSFSLLLPFLPFTSFAETWPQGDIARQKIVKLEKDFGGRIGVSAIDTGANRT
FD
>gb|ADB56658_1|ARO_3004753|BIC-1__Pseudomonas_fluorescens__TM_#02
GAVLSHSQQQEGLLEKRIDYK
>gb|ADB56658_1|ARO_3004753|BIC-1__Pseudomonas_fluorescens__TM_#03
SAQHSSTGMTVAQLAA
>gb|ADB56658_1|ARO_3004753|BIC-1__Pseudomonas_fluorescens__TM_#04
NVLGGPAGMTTFM
>gb|ADB56658_1|ARO_3004753|BIC-1__Pseudomonas_fluorescens__TM_#05
HAIARSLQKIALGEALQTAPRQQ
>gb|ADB56658_1|ARO_3004753|BIC-1__Pseudomonas_fluorescens__TM_#06
TSAPIVLAIYTAKPNKEDKHSDAVIAEVTRAVLESFE
>gb|BAL43359_1|ARO_3003742|mphG__Photobacterium_damselae_subsp__damselae__TM_#01
MKNRDIQKLAERN
>gb|BAL43359_1|ARO_3003742|mphG__Photobacterium_damselae_subsp__damselae__TM_#02
RDGTKWLLRIPRRTTLGEQIANEK
>gb|BAL43359_1|ARO_3003742|mphG__Photobacterium_damselae_subsp__damselae__TM_#03
TEEVLRNNLKILTPEQVRNEISERLILVKSELGINAELEL
>gb|BAL43359_1|ARO_3003742|mphG__Photobacterium_damselae_subsp__damselae__TM_#04
LSSAKLQLGVE
>gb|CAA90683_1|ARO_3003012|dfrA13__Escherichia_coli__TM_#01
VRIYLVAAMGA
>gb|CAA90683_1|ARO_3003012|dfrA13__Escherichia_coli__TM_#02
KIFRRLTE
>gb|CAA90683_1|ARO_3003012|dfrA13__Escherichia_coli__TM_#03
TVVLSRQA
>gb|CAA90683_1|ARO_3003012|dfrA13__Escherichia_coli__TM_#04
GCAVVSTLS
>gb|CAA90683_1|ARO_3003012|dfrA13__Escherichia_coli__TM_#05
AEHGKELYVA
>gb|CAA90683_1|ARO_3003012|dfrA13__Escherichia_coli__TM_#06
GAEVYALALP
>gb|EGE18576_1|ARO_3004466|ICR-Mc__Moraxella_catarrhalis_BC1__TM_#01
PFIASLAVVLTGVLLLLLVLLGYRHTLKTVAICFILIAAFAGH
>gb|EGE18576_1|ARO_3004466|ICR-Mc__Moraxella_catarrhalis_BC1__TM_#02
KLANMSYKNATKPTETIMHANDAIQKTTASTR
>gb|EGE18576_1|ARO_3004466|ICR-Mc__Moraxella_catarrhalis_BC1__TM_#03
ASFNGYQRATFPHMDKLIGLGQVH
>gb|EGE18576_1|ARO_3004466|ICR-Mc__Moraxella_catarrhalis_BC1__TM_#04
KYDVDTADYHENVIDTLDRL
>gb|EGE18576_1|ARO_3004466|ICR-Mc__Moraxella_catarrhalis_BC1__TM_#05
MNRLPAKQYQDYKNSPLQGG
>gb|EGE18576_1|ARO_3004466|ICR-Mc__Moraxella_catarrhalis_BC1__TM_#06
DLDDHVKAHA
>gb|EGE18576_1|ARO_3004466|ICR-Mc__Moraxella_catarrhalis_BC1__TM_#07
DEFAQFLPVCTSSELAECERQT
>gb|EGE18576_1|ARO_3004466|ICR-Mc__Moraxella_catarrhalis_BC1__TM_#08
LSIPALLWLGADTPFAVANSPTAGFS
>gb|EGE18576_1|ARO_3004466|ICR-Mc__Moraxella_catarrhalis_BC1__TM_#09
STQATADKTAFVNPLD
>gb|CAL25116_3|ARO_3003854|ADC-8__Acinetobacter_baylyi_ADP1__TM_#01
MMKDILGNLDNVPFKIMTGCIAGLLSCGTVAQSTVQQSIQQSVDRHF

```
>gb|CAL25116_3|ARO_3003854|ADC-8__Acinetobacter_baylyi_ADP1__TM_#02
QHYYQNYGVASKQTEQN
>gb|CAL25116_3|ARO_3003854|ADC-8__Acinetobacter_baylyi_ADP1__TM_#03
KTTQDMTQ
>gb|CAL25116_3|ARO_3003854|ADC-8__Acinetobacter_baylyi_ADP1__TM_#04
NSYSNLIENTLFPA
>gb|CAL25116_3|ARO_3003854|ADC-8__Acinetobacter_baylyi_ADP1__TM_#05
SNYAWGYQADQ
>gb|CAL25116_3|ARO_3003854|ADC-8__Acinetobacter_baylyi_ADP1__TM_#06
KFLDAQINPQNLKPTLRKAIQTTQMGYFRVGQMR
>gb|CAL25116_3|ARO_3003854|ADC-8__Acinetobacter_baylyi_ADP1__TM_#07
AGNSAKMALQPQPVTGISKPITAPQQALLNKTGATNGFSAYVVV
>gb|CAL25116_3|ARO_3003854|ADC-8__Acinetobacter_baylyi_ADP1__TM_#08
ATLQQILNADIQK
>gb|BAH63251_1|ARO_3004580|Klebsiella_pneumoniae_KpnE__Klebsiella_pneumoniae_subsp__pneumoniae_NTUH-K2044__TM_#01
MFYWILLA
>gb|BAH63251_1|ARO_3004580|Klebsiella_pneumoniae_KpnE__Klebsiella_pneumoniae_subsp__pneumoniae_NTUH-K2044__TM_#02
IGIVLIKSGTQKKASSQQEVAHAAV
>gb|BAP68758_1|ARO_3001855|ACT-35__Enterobacter_cloacae__TM_#01
PGMAVAVIY
>gb|BAP68758_1|ARO_3001855|ACT-35__Enterobacter_cloacae__TM_#02
HTWINVPK
>gb|BAP68758_1|ARO_3001855|ACT-35__Enterobacter_cloacae__TM_#03
LAQSRYWR
>gb|WP_064190968_1|ARO_3002583|AAC(6_)-29a__Pseudomonas_aeruginosa__TM_#01
MSVSILPVKEQDAADWLALRNLLWLADDHASEIEQYFSGG
>gb|WP_064190968_1|ARO_3002583|AAC(6_)-29a__Pseudomonas_aeruginosa__TM_#02
EVLIARDATGAAVGHVELSIRHDLEELQ
>gb|WP_064190968_1|ARO_3002583|AAC(6_)-29a__Pseudomonas_aeruginosa__TM_#03
APSHRSTDLVRRFLRESEKWALEQGCSAFASDRSDRVITHRKFAGSAV
>gb|NP_862226_1|ARO_3004639|Corynebacterium_striatum_tetA__Corynebacterium_striatum__TM_#01
IQFIGEATGDGLATASVRRVTHNAQQHLSS
>gb|NP_862226_1|ARO_3004639|Corynebacterium_striatum_tetA__Corynebacterium_striatum__TM_#02
VCAMVAMWSVSPWISLAIPA
>gb|NP_862226_1|ARO_3004639|Corynebacterium_striatum_tetA__Corynebacterium_striatum__TM_#03
TRFHAETAKANG
>gb|NP_862226_1|ARO_3004639|Corynebacterium_striatum_tetA__Corynebacterium_striatum__TM_#04
QPALTMAGL
>gb|NP_862226_1|ARO_3004639|Corynebacterium_striatum_tetA__Corynebacterium_striatum__TM_#05
DDAAGPEPTDTPVPGAGLWILEPAERSYATAAAWAQRAD
>gb|AHA41505_1|ARO_3002930|vanRO__Rhodococcus_hoagii__TM_#01
DARGGGDGG
>gb|AAD12162_1|ARO_3001299|tlrB_conferring_tylosin_resistance__Streptomyces_fradiae__TM_#01
MRKNVVRYLRCPHCAAPLRSSDRT
>gb|AAD12162_1|ARO_3001299|tlrB_conferring_tylosin_resistance__Streptomyces_fradiae__TM_#02
RPTKLAADTTDMVAARAALLDSGHYAPLTERLAGTA
>gb|AAD12162_1|ARO_3001299|tlrB_conferring_tylosin_resistance__Streptomyces_fradiae__TM_#03
RAAGAGAPD
>gb|AAD12162_1|ARO_3001299|tlrB_conferring_tylosin_resistance__Streptomyces_fradiae__TM_#04
DTLPLRDGAAAMA
>gb|AAD12162_1|ARO_3001299|tlrB_conferring_tylosin_resistance__Streptomyces_fradiae__TM_#05
EAVGQERLRTTLRLDHDALGRVVAMGPSSWHQDPDELARRIAELPGIHEVTLSVTFTVCR
PLP
>gb|CAD53575_1|ARO_3002531|AAC(3)-Ic__Pseudomonas_aeruginosa__TM_#01
MISTQTKITRLNSQDVG
>gb|CAD53575_1|ARO_3002531|AAC(3)-Ic__Pseudomonas_aeruginosa__TM_#02
EDAENYCRAQPSDSYLQDLLCGSGFIAIAALQGQ
>gb|CAD53575_1|ARO_3002531|AAC(3)-Ic__Pseudomonas_aeruginosa__TM_#03
KEIYIYDLGVQGA
>gb|AAD22403_1|ARO_3002970|vanTC__Enterococcus_gallinarum__TM_#01
MKNKGIDQFRVIAAMMVVAIHCLPLHYLWPEGDI
>gb|AAD22403_1|ARO_3002970|vanTC__Enterococcus_gallinarum__TM_#02
VFAELAVANSYPSRQRVFNFIKKQLKVYLLATLMFLPLALYSQTIGFDLPVGTLVQV
>gb|AAD22403_1|ARO_3002970|vanTC__Enterococcus_gallinarum__TM_#03
LTSLLIHVSFKKVFWLAAGL
>gb|AAD22403_1|ARO_3002970|vanTC__Enterococcus_gallinarum__TM_#04
FGLIQQTPIEPFYTAVFHLLDGTRNGIFFTPLFLCLGVLVRKQSEKRSLSKTALFFLISL
IGLLIESAYLHG
>gb|AAD22403_1|ARO_3002970|vanTC__Enterococcus_gallinarum__TM_#05
RWHPHRTWKHP
>gb|AAD22403_1|ARO_3002970|vanTC__Enterococcus_gallinarum__TM_#06
GTHFLSQKISILQNNLINYLVVLILTIGFICLFLRQKHSWFRHKQTTPVKRAVKEFSKTA
LLHNLQEIQRIISPK
>gb|AAD22403_1|ARO_3002970|vanTC__Enterococcus_gallinarum__TM_#07
KEVAPVLEQA
>gb|AAD22403_1|ARO_3002970|vanTC__Enterococcus_gallinarum__TM_#08
NAVKSPILVLGYTSPKRIKEL
>gb|AAB53445_1|ARO_3002897|SAT-4__Campylobacter_coli__TM_#01
MITEMKAEHLKDIDKPSEPFEVIGKIIPRYENENWTFTELLYEAPYLKSYQDEEDEEDEE
ADCLEYIDNTDKIIYLYYQDDKCVGKVK
>gb|AAB53445_1|ARO_3002897|SAT-4__Campylobacter_coli__TM_#02
AVCKDFRGQGIGSALINISIEWAKHKNLHGLMLETQDNNLIACKFYHNCGFKIGSVDTML
YANFENNFEKAVFWYLRF
>gb|AAA71915_1|ARO_3000173|tet(E)__Escherichia_coli__TM_#01
MNRTVMMALVIIF
>gb|AAA71915_1|ARO_3000173|tet(E)__Escherichia_coli__TM_#02
GKANVAEN
>gb|AAA71915_1|ARO_3000173|tet(E)__Escherichia_coli__TM_#03
ALMATASVV
>gb|AAA71915_1|ARO_3000173|tet(E)__Escherichia_coli__TM_#04
PEESRTHWFGMMGACFGGGMIAGPVIGGFAGQLSVQAPFMFAAAINGLAFLVSLFILHET
HNANQVSDELKNETINETTSSIREMISPLSGLLVVFFII
>gb|AAA71915_1|ARO_3000173|tet(E)__Escherichia_coli__TM_#05
GEERFAWDGVMVGVSLAVFGLTHALFQGLAAGFIAKHLGERKAIAVGILADGCGLFLLAV
ITQSWMVWPVLLLLAC
>gb|AAA71915_1|ARO_3000173|tet(E)__Escherichia_coli__TM_#06
VRVGQVAQGQLQGVLTSLTHLTAVIGPLVFAFLYSATRETWNGWVWIIGCGLYVVALIIL
RFFHPGRVIHPINKSDVQQRI
>gb|AAB51421_1|ARO_3002686|catP__Clostridium_perfringens__TM_#01
MVFEKIDK
>gb|AAB51421_1|ARO_3002686|catP__Clostridium_perfringens__TM_#02
CKSDFKSFLADYESDTQRYGNNHRME
>gb|ATC67679_1|ARO_3000838|arlR__Staphylococcus_aureus_subsp__aureus_str__Newman__TM_#01
```

DGLDKALSHY
>gb|ATC67679_1|ARO_3000838|arlR__Staphylococcus_aureus_subsp__aureus_str__Newman__TM_#02
STPIIIITAKSDT
>gb|ATC67679_1|ARO_3000838|arlR__Staphylococcus_aureus_subsp__aureus_str__Newman__TM_#03
TIDKNAFKVTVNGAEI
>gb|AAA98349_1|ARO_3001307|VgbA__Staphylococcus_aureus__TM_#01
EEYPIQIKSAEPHGICF
>gb|CBY77552_1|ARO_3003106|Erm(42)__Pasteurella_multocida__TM_#01
MNKNTKIKNKNF
>gb|CBY77552_1|ARO_3003106|Erm(42)__Pasteurella_multocida__TM_#02
NTKLVEDLLFKSNITKEDF
>gb|CBY77552_1|ARO_3003106|Erm(42)__Pasteurella_multocida__TM_#03
KALSKICKAVNAIEFDSVLADKLSHEFKSSNVSIIEADFLKYNLPDHN
>gb|CBY77552_1|ARO_3003106|Erm(42)__Pasteurella_multocida__TM_#04
LNKLLDSENPPLDT
>gb|CBY77552_1|ARO_3003106|Erm(42)__Pasteurella_multocida__TM_#05
APSYKESYKSLLYKPFFKTNILHSFSKFDFKPAPNANIILGQFSYKDFTDINLEDRHAWK
DFLAFVFLEKGVTFKEKTKRIFSYKQQKIILKESRINDDSNISNWSYEFWLKMFKLYNSN
MVSKDKKVLVNNSYKRMLEHESSLEKIHRNRKQNNRK
>gb|ACC54440_2|ARO_3002799|QnrVC1__Vibrio_cholerae__TM_#01
HIFSNCTFIHCNFKR
>gb|ACC54440_2|ARO_3002799|QnrVC1__Vibrio_cholerae__TM_#02
FSEDCWEQF
>gb|ATM29809_1|ARO_3004688|MCR-1_11__Escherichia_coli__TM_#01
HTSVVWYR
>gb|ATM29809_1|ARO_3004688|MCR-1_11__Escherichia_coli__TM_#02
PVVAFSSHYASFFRV
>gb|ATM29809_1|ARO_3004688|MCR-1_11__Escherichia_coli__TM_#03
QSIQWLQT
>gb|ATM29809_1|ARO_3004688|MCR-1_11__Escherichia_coli__TM_#04
KQTGITPM
>gb|AAF61147_1|ARO_3004751|BES-1__Serratia_marcescens__TM_#01
MWQWLGKVRWLLVVIALLGGKVMAASELDSALARLEQQHHGRLGLAYIDSGSGESY
>gb|AAF61147_1|ARO_3004751|BES-1__Serratia_marcescens__TM_#02
SVSQPGLLDKRVHYAATDLLAYAPITKTHLDKGMRIGELA
>gb|AAF61147_1|ARO_3004751|BES-1__Serratia_marcescens__TM_#03
VQALNRFVQGLGDPAFRLDRIEPH
>gb|AAF61147_1|ARO_3004751|BES-1__Serratia_marcescens__TM_#04
QAMTLGKGLPQAQQA
>gb|AAF61147_1|ARO_3004751|BES-1__Serratia_marcescens__TM_#05
EQGAPKVLAIYFTQPAADAEANRAILAEATRLVLQDKSINKIK
>gb|AKO71461_1|ARO_3004608|EreD__Riemerella_anatipestifer__TM_#01
MKNIKPLTFPFNSENNTSIKQSLFRFREYFDSSTIVGLGENSHFIKEFFTFRHQVIEFLV
TECDFDTLAFEFGFSEGLEVDKWIKSQIPFDDLDKLLSHFYYPNEFKDTLLWLRRYNQDN
NNQITFLGVDIPKNGGSYFPNFRIVSDYLQRLSIVSSDVLQKILNLAEKFDFYSTSQLAL
NLSLFDEAEHNELKALLLKVYIRLVTLQPKLESLEFQSILHQVKGLIYMNYNADAMESFI
TEKGIEGDMGAKDQYMAESIDWFLKNSLGKKIILVAHNAHIQKTPVDFDGFISCYPMGQR
>gb|SIP52035_1|ARO_3004054|Pseudomonas_aeruginosa_CpxR__Pseudomonas_aeruginosa__TM_#01
AQPSAQMQLGDLSLNLTRGVAQIDG
>gb|AMY61250_1|ARO_3004092|HMB-1_beta-lactamase__Pseudomonas_aeruginosa__TM_#01
MKIHLWISLGLLLLTNIV
>gb|AMY61250_1|ARO_3004092|HMB-1_beta-lactamase__Pseudomonas_aeruginosa__TM_#02
NYPSWGLVASHGLVFVDGKDAYIIDTPATVKDTEVLVQWINDQGFKPKA
>gb|AMY61250_1|ARO_3004092|HMB-1_beta-lactamase__Pseudomonas_aeruginosa__TM_#03
QTNELLNKEVAAQATHSFK
>gb|AAF03531_1|ARO_3002569|AAC(6_)-Iy__Salmonella_enterica_subsp__enterica_serovar_Enteritidis__TM_#01
HWRGLRKQLWPGHPDDAHLADGEEILQADHLASFIAMADGVA
>gb|AAF03531_1|ARO_3002569|AAC(6_)-Iy__Salmonella_enterica_subsp__enterica_serovar_Enteritidis__TM_#02
LPSFRQRGVAKQLIAAVQRWGTNKGCREMASDTSPENTISQK
>gb|AAC41392_1|ARO_3002557|AAC(6_)-Ij__Acinetobacter_genomosp__13__TM_#01
MNIMPVSESLMADWLGLRKLLWPDHDEAHLQEMQRLLQQTQS
>gb|AAC41392_1|ARO_3002557|AAC(6_)-Ij__Acinetobacter_genomosp__13__TM_#02
HLVQQVEAWAKPFGCIEFASDAALDNR
>gb|AAL26797_1|ARO_3004448|HERA-1__Atlantibacter_hermannii__TM_#01
MKKITPLF
>gb|AAL26797_1|ARO_3004448|HERA-1__Atlantibacter_hermannii__TM_#02
IAFLTLIALLAPAQASVTPDMTDFLRQQEQ
>gb|AAL26797_1|ARO_3004448|HERA-1__Atlantibacter_hermannii__TM_#03
ALLERLQKNGGSLDEQVTIPPDALLDYAPVTKNYLAPATISLRMLCA
>gb|AAL26797_1|ARO_3004448|HERA-1__Atlantibacter_hermannii__TM_#04
GPDAVTQFMRGIG
>gb|AAL26797_1|ARO_3004448|HERA-1__Atlantibacter_hermannii__TM_#05
QKMAAGLQKILT
>gb|AAL26797_1|ARO_3004448|HERA-1__Atlantibacter_hermannii__TM_#06
EASMKMANE
>gb|AAL26797_1|ARO_3004448|HERA-1__Atlantibacter_hermannii__TM_#07
IGKQLFAGQP
>gb|AAB05624_1|ARO_3002956|vanYB__Enterococcus_faecalis__TM_#01
GEKRAFLW
>gb|AAB05624_1|ARO_3002956|vanYB__Enterococcus_faecalis__TM_#02
GVAENPEENTLATAKEQGDEQEWSLILVNRQNPIPAQYDVELEQLSNGERIDIRISPYLQ
D
>gb|AAB05624_1|ARO_3002956|vanYB__Enterococcus_faecalis__TM_#03
EIMDEKVAEYKAKGYTSA
>gb|AAB05624_1|ARO_3002956|vanYB__Enterococcus_faecalis__TM_#04
RWLDENSYR
>gb|AAB05624_1|ARO_3002956|vanYB__Enterococcus_faecalis__TM_#05
YHQGLCLEEYLNTEK
>gb|BAL43360_1|ARO_3003745|mefC__Photobacterium_damselae_subsp__damselae__TM_#01
MENRKWFKTYMFIWAGQFASMLTSYAVQFAIVIWLSLEY
>gb|BAL43360_1|ARO_3003745|mefC__Photobacterium_damselae_subsp__damselae__TM_#02
LIAGVYVDRLNRKYVMIFSDAFIALCALLLLVILQNENVNLIWIYILLGLRSVGNAFHAP
ALQAIAPLIVPQNELIKVAGINQVLHSVCRIGGPAIGTLAIAY
>gb|BAL43360_1|ARO_3003745|mefC__Photobacterium_damselae_subsp__damselae__TM_#03
LSLVMVKIPNVVAKSKSSAHSIATEFSEGFQTVSKNKGLRYLFLYAMAITFV
>gb|BAL43360_1|ARO_3003745|mefC__Photobacterium_damselae_subsp__damselae__TM_#04
IGIVEVVWGGGMLIGGVILSIFKLKGSKVVAVNVMYVLLGLTFILSGVLPASWFVGFVMV
TAIGGISLSVFNGCFTAIVQTEVS
>gb|BAL43360_1|ARO_3003745|mefC__Photobacterium_damselae_subsp__damselae__TM_#05
STRNLMQLGKIKNI
>gb|CAA68516_1|ARO_3002657|APH(6)-Ia__Streptomyces_griseus__TM_#01
MSSSDHIHVPDGLAESYSRSG
>gb|CAA68516_1|ARO_3002657|APH(6)-Ia__Streptomyces_griseus__TM_#02

RCVDRWELKRDGGVR
>gb|CAA68516_1|ARO_3002657|APH(6)-Ia__Streptomyces_griseus__TM_#03
NRLHSVPAPPGLRGLGEIAGAMVEEVPSAVDSLADPEDRS
>gb|CAA68516_1|ARO_3002657|APH(6)-Ia__Streptomyces_griseus__TM_#04
LTAIAPSQIAVAEALAKP
>gb|CAL30186_1|ARO_3002699|cmlB1__Bordetella_bronchiseptica__TM_#01
VVGALLLAVGET
>gb|CAL30186_1|ARO_3002699|cmlB1__Bordetella_bronchiseptica__TM_#02
NPNRHHPSDDEHDSLATQDIGRSQSGHGHD
>gb|AAL12234_1|ARO_3003552|fusB__Staphylococcus_aureus__TM_#01
MKTMIYPHQYNYIRSVILRLKNVYKTVNDKETVKVIQSETYNDINEIFGHIDDDIEESLK
VLMNIRLSNKEIEAILNKFLEYVVPFELPSPQKLQKVFKKVKKIKIPQFEEYDLKVSSFV
GWNELASNRKYIIYYDEKKQLKGLYGEISN
>gb|AAL12234_1|ARO_3003552|fusB__Staphylococcus_aureus__TM_#02
SKTNSDGQYVKKGDYICRDSIHCNKQLTDINQFYNFIDKLD
>gb|ACL82959_1|ARO_3002961|vanYM__Enterococcus_faecium__TM_#01
NLFKHDELTKGYELLNREIYLSEKVAREFSEMVDAAEKEGVRHFSINSGFRN
>gb|ACL82959_1|ARO_3002961|vanYM__Enterococcus_faecium__TM_#02
GTIHGENYEISYYPITEKTDIEM
>gb|WP_063978071_1|ARO_3000862|sgm__Micromonospora_zionensis__TM_#01
MTAPAADDRIDEIERAITKSRRYQTVAPATVRRLARAALVAARGDVPDAVKRTKRGLHEI
YGAFLPPSPPNYAALLRHLDSAVDAGDDEAVRAALL
>gb|WP_063978071_1|ARO_3000862|sgm__Micromonospora_zionensis__TM_#02
ETVYIASDIDARLVGFVDEALTRLNVPHRTNVADLLED
>gb|AAK07612_2|ARO_3000604|Erm(35)__Bacteroides_coprosuis_DSM_18011__TM_#01
VALSQHLRKKFIHAQNVQVVSCDYRNFVVPKV
>gb|AAK07612_2|ARO_3000604|Erm(35)__Bacteroides_coprosuis_DSM_18011__TM_#02
SSLMFENVEYFLCGSIILQSEPAKKLFSSKVYNPLTVLYHTYYDLKF
>gb|AAK07612_2|ARO_3000604|Erm(35)__Bacteroides_coprosuis_DSM_18011__TM_#03
RIERKQISLDIGLKVKYLNFVSYMLQKPDLTVKTAMKSIFRKKQVRSISEKFGVDLNSK
>gb|AGH88989_1|ARO_3001205|determinant_of_bleomycin_resistance__Acinetobacter_baumannii__TM_#01
KLGFATSWKDRG
>gb|AGH88989_1|ARO_3001205|determinant_of_bleomycin_resistance__Acinetobacter_baumannii__TM_#02
ALVNAAGAEEKSTGWPRFKAPQLEASGLRIGY
>gb|CCP45647_1|ARO_3003955|efpA__Mycobacterium_tuberculosis_H37Rv__TM_#01
MTALNDTERAVRNWTAGRPHRPAPMRPPRSE
>gb|CCP45647_1|ARO_3003955|efpA__Mycobacterium_tuberculosis_H37Rv__TM_#02
FGAMLYGSFFMH
>gb|CZT31773_1|ARO_3004094|Erm(48)__Staphylococcus_xylosus__TM_#01
YINEILQNTNIES
>gb|CZT31773_1|ARO_3004094|Erm(48)__Staphylococcus_xylosus__TM_#02
ALLKISHFVTGIEIDRNLYYKLKKDTDLYDNLKLINKDVLRFQFHQNEP
>gb|CZT31773_1|ARO_3004094|Erm(48)__Staphylococcus_xylosus__TM_#03
LNKKRALSLLLLPKMDVEILKIIPNVYFHPKPTVDSALILLKRHKPLVSEKDEKI
>gb|CZT31773_1|ARO_3004094|Erm(48)__Staphylococcus_xylosus__TM_#04
HANVQDINE
>gb|ADM92606_1|ARO_3000549|adeS__Acinetobacter_baumannii__TM_#01
SFQQEDWTS
>gb|ADM92606_1|ARO_3000549|adeS__Acinetobacter_baumannii__TM_#02
CIISLVIGMRLAKRFIV
>gb|ADM92606_1|ARO_3000549|adeS__Acinetobacter_baumannii__TM_#03
YDNRIHSAEMSELLY
>gb|ADM92606_1|ARO_3000549|adeS__Acinetobacter_baumannii__TM_#04
VADNWILKIEDEGPGIATEFRD
>gb|ANZ79241_1|ARO_3004036|tetB(60)__uncultured_bacterium__TM_#01
MRTMKRLLSYLRYEKKGVLIGLFCLLLSTGATLTGPLVAKHIIDNVITPMGQAHDFKAGG
LLLWVGIYVTVNLVGVAGAYLNRVY
>gb|ANZ79241_1|ARO_3004036|tetB(60)__uncultured_bacterium__TM_#02
FSTIVMLVGVYITIFLLNATLGLVLLFLVPVMILWQRTVATKQKKYYSENRELYSQLSGQ
LNESIQGAGIVQAFQQEEKIVAEYDATATSWVEVGRKELILESYFSWSLVGMLRNITHFG
VIYYFSMQFIGGTLGISAGLLYAFIDYINRIYEPIQTFMNVVSGFQQSMAAGDRVFELMD
TPSEESGEELFTFDEGCIEFKDVSFEYTAGVPVLK
>gb|ACH58998_1|ARO_3002486|LRA-7__uncultured_bacterium_BLR7__TM_#01
ADAPKRLPVNITN
>gb|ACH58998_1|ARO_3002486|LRA-7__uncultured_bacterium_BLR7__TM_#02
IMGSYPLMKASIES
>gb|ACH58998_1|ARO_3002486|LRA-7__uncultured_bacterium_BLR7__TM_#03
VYMSERDVESL
>gb|ACH58998_1|ARO_3002486|LRA-7__uncultured_bacterium_BLR7__TM_#04
APEGRGGLVYDPIH
>gb|ACH58998_1|ARO_3002486|LRA-7__uncultured_bacterium_BLR7__TM_#05
QQTDAGKT
>gb|ACH58998_1|ARO_3002486|LRA-7__uncultured_bacterium_BLR7__TM_#06
QVYKPGDAYNPARFGD
>gb|ACH58998_1|ARO_3002486|LRA-7__uncultured_bacterium_BLR7__TM_#07
KIATAKAN
>gb|AAA90937_1|ARO_3004635|AAC(6_)-Il__Klebsiella_aerogenes__Partial_TM_#01
MDSSPLVRPVETTDSASWLSMRCELWPDGTCQEHQSEIAEFLSGKVARPAAVLIAVAPDG
EALGFAELSIRPYAEECYSGNVAFLEGWYVVPSARRQGVGVALVKAAEHWARGRGCTEFA
SDTQLTNSASTSAHLAAGFTEVAQVRCFRKPL
>gb|ACL82958_1|ARO_3002939|vanSM__Enterococcus_faecium__TM_#01
MAKMRSSFRTKIILLFAV
>gb|ACL82958_1|ARO_3002939|vanSM__Enterococcus_faecium__TM_#02
MIHRGNPLAELRDFIESIGDFNFFFLLFILLSLSVFYILTKPYSAYFDEISTGIQYLAL
>gb|ACL82958_1|ARO_3002939|vanSM__Enterococcus_faecium__TM_#03
HDGQFVDINGFVDE
>gb|ACL82958_1|ARO_3002939|vanSM__Enterococcus_faecium__TM_#04
ESAPIDQV
>gb|AAG05913_1|ARO_3004072|OpmB__Pseudomonas_aeruginosa_PAO1__TM_#01
AVPAEFKEA
>gb|AAG05913_1|ARO_3004072|OpmB__Pseudomonas_aeruginosa_PAO1__TM_#02
VAQFRQAEALVRGARAAFFPSITGNVGKTRSGQGGGDSTVLLPGGSTVSSGGSGAISTSY
S
>gb|AAG05913_1|ARO_3004072|OpmB__Pseudomonas_aeruginosa_PAO1__TM_#03
TAYERSLKVAENK
>gb|AAG05913_1|ARO_3004072|OpmB__Pseudomonas_aeruginosa_PAO1__TM_#04
ERTDERLGRVEEGLPPSP
>gb|SBV31106_1|ARO_3004110|MCR-2__Escherichia_coli__TM_#01
ATANLTFF
>gb|SBV31106_1|ARO_3004110|MCR-2__Escherichia_coli__TM_#02
FVRIIGLGVLPS
>gb|SBV31106_1|ARO_3004110|MCR-2__Escherichia_coli__TM_#03
LIQRAMTWGV

```
>gb|SBV31106_1|ARO_3004110|MCR-2__Escherichia_coli__TM_#04
PIYSVGKLASIEYKKATAP
>gb|SBV31106_1|ARO_3004110|MCR-2__Escherichia_coli__TM_#05
DTIYHAKDAVQTT
>gb|SBV31106_1|ARO_3004110|MCR-2__Escherichia_coli__TM_#06
KSIDWLKTH
>gb|WP_102607457_1|ARO_3004361|sul4__Proteobacteria__TM_#01
VEEDVIAK
>gb|WP_102607457_1|ARO_3004361|sul4__Proteobacteria__TM_#02
LARVLPVVQAVRQAVDVV
>gb|WP_102607457_1|ARO_3004361|sul4__Proteobacteria__TM_#03
IAQEKKLGGRFIGVKYDDLITDVKRELQESIDIALK
>gb|WP_102607457_1|ARO_3004361|sul4__Proteobacteria__TM_#04
KAIVRVARMTDAIVRR
>gb|CAA30695_1|ARO_3002685|catIII__Plasmid_R387__TM_#01
MNYTKFDVKNWVRREHFEFYRHRLP
>gb|CAA30695_1|ARO_3002685|catIII__Plasmid_R387__TM_#02
FDELRMAIKDDELIVWDSVDPQFTVFHQETETFSALSCPYSSDIDQFMVNYLSVMERYKS
DTKLFPQGVTPENHLNISALPWVN
>gb|CAA30695_1|ARO_3002685|catIII__Plasmid_R387__TM_#03
RLQELCNSKLK
>gb|AAB05623_1|ARO_3002932|vanSB__Enterococcus_faecalis__TM_#01
AQQTVKSYQPLVEL
>gb|AAB05623_1|ARO_3002932|vanSB__Enterococcus_faecalis__TM_#02
IVAQSKAGVGLLYQGLTIRGIVMIAIMVVFSLLCAYIFARQMTTPIKALADSANKMANLK
EVPPPLERKDELGALAHDMHSMYIRLKETIARLEDEIA
>gb|AAB05623_1|ARO_3002932|vanSB__Enterococcus_faecalis__TM_#03
RQGKTISEILELVSLNDGRIVPIAEPLDIGRTVAELLPDFQTLAEANNQRFVTDIPAGQI
VLSDPKLI
>gb|AAB05623_1|ARO_3002932|vanSB__Enterococcus_faecalis__TM_#04
LFIPFYRIDQARSRKSGRSGLGLAIVQKTLDAMSLQYALENTSDGVLFWLDLPPTSTL
>gb|BAH66386_1|ARO_3002574|AAC(6_)-Iaf__Pseudomonas_aeruginosa__TM_#01
MDYSICDIAESNELILEAAK
>gb|BAH66386_1|ARO_3002574|AAC(6_)-Iaf__Pseudomonas_aeruginosa__TM_#02
CLGICLDDKLIGWTGLRP
>gb|BAH66386_1|ARO_3002574|AAC(6_)-Iaf__Pseudomonas_aeruginosa__TM_#03
QKTSLSMIDINERNIFDEI
>gb|BAH66386_1|ARO_3002574|AAC(6_)-Iaf__Pseudomonas_aeruginosa__TM_#04
RKNSPTIAST
>gb|AAC36915_1|ARO_3000593|ErmQ__Clostridium_perfringens__TM_#01
MKAKSNNYRGKVDISVSQNFITSKNTIYKLIKKTNISKNDFVIEIGPGKGHITEALCEKS
YWVTAIELDRSLYGNLINKFKSKNNVTLINKDFLNWKLPKKRE
>gb|AAC36915_1|ARO_3000593|ErmQ__Clostridium_perfringens__TM_#02
YITTKIIKKLLLEELNSPTDMWLVMEKGSAKRFMGIPRESKLSLLLKTKFDIKIVHYFNR
EDFHPMPSVDCVLVYFKRKYKYDISKDEWNEYTSFISKSINNLRDVFTKNQIHAVIKYLG
INLNNISEVSYNDWIQLFRYKQKID
>gb|ENV82218_1|ARO_3001754|OXA-299__Acinetobacter_bouvetii_DSM_14964___CIP_107468__TM_#01
MNPFTKYCAILCPIIFLGACTISPFSHDQAHSAHASQLTDAATIRNLFNQANVQGVILIK
SGNDLQAYGNAIQ
>gb|ENV82218_1|ARO_3001754|OXA-299__Acinetobacter_bouvetii_DSM_14964___CIP_107468__TM_#02
IEHNKTSPD
>gb|ENV82218_1|ARO_3001754|OXA-299__Acinetobacter_bouvetii_DSM_14964___CIP_107468__TM_#03
RQEVQFADQLSHLQLPFRKSTQQQVIQMLFIEQIGSKA
>gb|ENV82218_1|ARO_3001754|OXA-299__Acinetobacter_bouvetii_DSM_14964___CIP_107468__TM_#04
QGKTTAFSLNLEMDQ
>gb|CAA26235_1|ARO_3003061|viomycin_phosphotransferase__Streptomyces_vinaceus__TM_#01
MRIIETHRDLLSRLLPGDTVGGLAVHEGQFHHVVIGSH
>gb|CAA26235_1|ARO_3003061|viomycin_phosphotransferase__Streptomyces_vinaceus__TM_#02
DEEKVRAALPEAPANEWQEFATGVRTELFP
>gb|AAA22081_1|ARO_3004451|Agrobacterium_fabrum_chloramphenicol_acetyltransferase__Agrobacterium_fabrum_str__C58__TM_#01
ALYRFWRSDSL
>gb|CAB51471_1|ARO_3004359|ACI-1__Acidaminococcus_fermentans__TM_#01
MKKFCFLFLIICGLMVFCLQDCQARQKLNLADLENKYNAVIGVYAVDMENGKKICYKPDT
RFSYCSTHKVFTAAELLRQKNTSDLNEIRKFSAEDILSYAPITKDHVADGMTLAEICSAS
LRWSDNTAANLILQEIGGVENFKVALKNIGDKTTKPARNEPELNLFNPK
>gb|CAB51471_1|ARO_3004359|ACI-1__Acidaminococcus_fermentans__TM_#02
VKNLQVYIFGDILSDDKKKLLIDWMSDNSITDTLIKAETPQGWKVIDKSGSGDYGARNDI
AVIYP
>gb|CAB51471_1|ARO_3004359|ACI-1__Acidaminococcus_fermentans__TM_#03
RRTEKNAKS
>gb|AAB58447_1|ARO_3002662|APH(9)-Ia__Legionella_pneumophila_130b__TM_#01
MLKQPIQA
>gb|AAB58447_1|ARO_3002662|APH(9)-Ia__Legionella_pneumophila_130b__TM_#02
TAQFIQGGADTNAFAYQADSESKSYFIKLKYGY
>gb|AAB58447_1|ARO_3002662|APH(9)-Ia__Legionella_pneumophila_130b__TM_#03
AKLFQQLK
>gb|AAB58447_1|ARO_3002662|APH(9)-Ia__Legionella_pneumophila_130b__TM_#04
ISIQQQLRKEIYSPKWREIVRSFYNQIEFDNSDDKLTAAFKSFFNQNSAAIHRLVDTSEK
>gb|BAI83385_1|ARO_3003440|mecB__Macrococcus_caseolyticus__TM_#01
MKNKALAILIICICLLIAYNFVKKDEVDKIFDAIELRDSEYLNEHATFLSKSLYDKDQRY
KRMDKIDASLGIKEVKVSNVRLVQKKKNKRQYSANLNFRTKYGNFSREGNYSFEKDEITK
KWLLDWSPEVIIPGLTDRNQISIETLESFRGKILDRNGIDIAKDGIHYEVGIDIKNLNKK
NKKNISKLLSISESTLNKKLKQTWVKEGVFLPLKSYIELDDELKLGIQKYHLTVNQTKGR
VYPLR
>gb|BAI83385_1|ARO_3003440|mecB__Macrococcus_caseolyticus__TM_#02
EINAEELKNKKFKDYDEHSIVGKSGIELQYDKQLQNKDGYKVVITSDDALNNDED
>gb|ALV80601_1|ARO_3003801|bcr-1__Pseudomonas_aeruginosa__TM_#01
MPASASRIQVGSGER
>gb|ALV80601_1|ARO_3003801|bcr-1__Pseudomonas_aeruginosa__TM_#02
DAQVQRSISGFLV
>gb|ALV80601_1|ARO_3003801|bcr-1__Pseudomonas_aeruginosa__TM_#03
FSSLACALADSAGQLVL
>gb|ALV80601_1|ARO_3003801|bcr-1__Pseudomonas_aeruginosa__TM_#04
ESHPPERRGGSLAQAFLAYGRLLGDRRALGYVLCMGLAF
>gb|ALV80601_1|ARO_3003801|bcr-1__Pseudomonas_aeruginosa__TM_#05
HGPRPLLRAGSLLACVSGLFLLGYAALGERGGLWALVPG
>gb|ALV80601_1|ARO_3003801|bcr-1__Pseudomonas_aeruginosa__TM_#06
GQAGAASAVAVSGQFGLGCLASLAV
>gb|BAO79432_1|ARO_3000785|cmeC__Campylobacter_jejuni_subsp__jejuni__TM_#01
MNKIISISAIASFT
>gb|BAO79432_1|ARO_3000785|cmeC__Campylobacter_jejuni_subsp__jejuni__TM_#02
SSITKNWWKDFDDENLNK
>gb|BAO79432_1|ARO_3000785|cmeC__Campylobacter_jejuni_subsp__jejuni__TM_#03
```

FDGSASGSRAKTAINAPSNRTGEVS
>gb|AAA65959_1|ARO_3002962|vanZA__Enterococcus_faecium__TM_#01
HQRSLNLTPFTATGNF
>gb|AAA65959_1|ARO_3002962|vanZA__Enterococcus_faecium__TM_#02
EIGFLPKFAFVLV
>gb|AAA65959_1|ARO_3002962|vanZA__Enterococcus_faecium__TM_#03
KLYGLSNKHMNQ
>gb|AAA65959_1|ARO_3002962|vanZA__Enterococcus_faecium__TM_#04
VYRTHLRINYV
>gb|ACS83559_1|ARO_3004601|LnuP__Clostridium_perfringens__TM_#01
MIGINDACEILSWAYNNN
>gb|ACS83559_1|ARO_3004601|LnuP__Clostridium_perfringens__TM_#02
YNKFIEIIKNKGFNEIVVEYTSEVHTIWSDNKLRIIDLHMFKDNCDGTICYEGEVFQKNI
FDGVGKIGNIMVSCINAK
>gb|ACS83559_1|ARO_3004601|LnuP__Clostridium_perfringens__TM_#03
GESDIHDVKLLCKEFNIPIPKEYENF
>gb|AAL19305_1|ARO_3000790|mdsB__Salmonella_enterica_subsp__enterica_serovar_Typhimurium_str__LT2__TM_#01
IGADGEVTRLRDVARVTLGADAYTLRSLLNGEAAP
>gb|AAL19305_1|ARO_3000790|mdsB__Salmonella_enterica_subsp__enterica_serovar_Typhimurium_str__LT2__TM_#02
KMDELQQN
>gb|CAL64019_1|ARO_3003441|cfrA__Staphylococcus_warneri__TM_#01
MNFNNKTKYGKIQEFLRSNNEPDYRIKQITNAIFKQRISRFEDMKVLPKLLREDLINNFG
ETVLNIKLLAEQNSEQVTKVLFEVSK
>gb|CAL64019_1|ARO_3003441|cfrA__Staphylococcus_warneri__TM_#02
RQVFDALDSF
>gb|CAL64019_1|ARO_3003441|cfrA__Staphylococcus_warneri__TM_#03
QEYPQVNLTFSLHSPYSEERSKLMPINDRYPIDEVMNI
>gb|CAL64019_1|ARO_3003441|cfrA__Staphylococcus_warneri__TM_#04
NEVVSLLKSRYK
>gb|CAL64019_1|ARO_3003441|cfrA__Staphylococcus_warneri__TM_#05
SAPEMYGE
>gb|WP_010896559_1|ARO_3004643|Erm(K)__Bacillus_halodurans__TM_#01
QEIVDQAKVSKK
>gb|WP_010896559_1|ARO_3004643|Erm(K)__Bacillus_halodurans__TM_#02
KRVLAVEYDQTFIQVLNRKMAHAANTTIIHEDIMRIHLPKGE
>gb|WP_010896559_1|ARO_3004643|Erm(K)__Bacillus_halodurans__TM_#03
PDPIVPYKDRSAFFGLAEYALREPKAPADSLLRGIFTATQLKHVKRNAGIKHDVSIGALS
ERQWGVIFETMTQYVRRPLWPRPRKTTL
>gb|AAF74725_1|ARO_3004659|Mef(En2)__Bacteroides_fragilis__TM_#01
MNHWKSTLAVIGI
>gb|AAF74725_1|ARO_3004659|Mef(En2)__Bacteroides_fragilis__TM_#02
NEFKSPTALSLAILAGFLPQFVLGL
>gb|AAF74725_1|ARO_3004659|Mef(En2)__Bacteroides_fragilis__TM_#03
FYSDLFIAFCTLCLFIVITK
>gb|AAF74725_1|ARO_3004659|Mef(En2)__Bacteroides_fragilis__TM_#04
QSFSEVIAPVVGASLVVWLPIQYILLIDVIGAVAACLTLLCVQIPSLQKTKVLPDFKKEL
TEWLHTLRRTMGILPLFVCFTLVTFVLMPVFTLFPFMTLLHFNGNILQMGVVEMGWGSGA
LLGGLVLACKALKSKQTLVMHTAYVILGLYLISASYLPSSAFIGFVCLTFTGGIAYSIYH
ALFIAIIQQNLASDM
>gb|AAF74725_1|ARO_3004659|Mef(En2)__Bacteroides_fragilis__TM_#05
ASGYWVEAWGITSVFMISGWVIFLIGVGANFISSIKQLDNYA
>gb|AAG05881_1|ARO_3000803|MexE__Pseudomonas_aeruginosa_PAO1__TM_#01
MEQSSHFS
>gb|AAA98484_1|ARO_3000596|ErmX__Plasmid_pNG2__TM_#01
KIINSIIDLVKQTSGPIIEIGPGSGALTHPMAHLGRAITAVEVDAKLAAKITQETSSAAV
EVVHDDFLNFR
>gb|AAA98484_1|ARO_3000596|ErmX__Plasmid_pNG2__TM_#02
SAFRPQPNVDGGILVIRRVGDPKIPIEQRKAFQAMVHT
>gb|AAA98484_1|ARO_3000596|ErmX__Plasmid_pNG2__TM_#03
RRQGCFHHVQKHNHGCAREESTPRPYLPDCY
>gb|AAL68827_1|ARO_3000605|Erm(36)__Micrococcus_luteus__TM_#01
DRLSRLVGDSTGPIVEIGPGQGRLTRELQKLGRSLTAVEIDSRLADRLASASQFREQKHV
TVVNADFLHWPLPTTPY
>gb|AAL68827_1|ARO_3000605|Erm(36)__Micrococcus_luteus__TM_#02
HGRVPRSAFKPAPSVDGGLLEMTRRPDPLLSPDARESYRQFVHD
>gb|AAL68827_1|ARO_3000605|Erm(36)__Micrococcus_luteus__TM_#03
ANVSSSLGKRGALQLLKSEGIRSSS
>gb|AAL68827_1|ARO_3000605|Erm(36)__Micrococcus_luteus__TM_#04
RLFTSASPTKSAKT
>gb|AAC77293_1|ARO_3001214|mdtM__Escherichia_coli_str__K-12_substr__MG1655__TM_#01
MGFISFVGLLLAMPETVKRGAVPFSAKSVLRDFRNVFC
>gb|AAC77293_1|ARO_3001214|mdtM__Escherichia_coli_str__K-12_substr__MG1655__TM_#02
IVVFTLAGLLNRVRQHQAAELVEEQ
>gb|AAG03814_1|ARO_3000377|MexA__Pseudomonas_aeruginosa_PAO1__TM_#01
MQRTPAMRVLVPALLVAISALSGCGKSEAPPPAQTPEVGIVTLEAQT
>gb|AAG03814_1|ARO_3000377|MexA__Pseudomonas_aeruginosa_PAO1__TM_#02
YQSAQANLASTQEQAQRYKLLVADQAVSKQQYADANAAYLQSK
>gb|AAG03814_1|ARO_3000377|MexA__Pseudomonas_aeruginosa_PAO1__TM_#03
TEGLNAGDKIITEGLQFVQPGVEVKTVPAKNVASAQKADAAPAKTDSKG
>gb|CAG27800_1|ARO_3003203|TLA-2__uncultured_bacterium__TM_#01
MNIKYFKFAEKFILLVLIMSFSSLAFCKSDDSLEQRINSIISGKKASVGVAVAGIEDNFS
LSINGKKNFPMMSVYKLHIVLAVLNKVDGGSLKLDEKIPLNKKDLHPG
>gb|CAG27800_1|ARO_3003203|TLA-2__uncultured_bacterium__TM_#02
SIPLSEIIE
>gb|CAG27800_1|ARO_3003203|TLA-2__uncultured_bacterium__TM_#03
ALAGGTEAVKRYIISKGISDFDIRATEKECHESWNVQYSNWSTPVSAVALLKKFNDRKIL
SSVSTEYLMNVMIHTSTGNKRIKGLIPPSADVAHKTGTSGIRNGITPG
>gb|CAD70268_1|ARO_3002704|fexA__Staphylococcus_lentus__TM_#01
LVNPVLPLISKDLEASKSQVSWIVSGIA
>gb|CAD70268_1|ARO_3002704|fexA__Staphylococcus_lentus__TM_#02
ELRKLYIFAIMILAS
>gb|CAD70268_1|ARO_3002704|fexA__Staphylococcus_lentus__TM_#03
NALFWFTFLLAIMIVIGAYYALPTIKPAESVGSNKNFDFIGGLF
>gb|CAD70268_1|ARO_3002704|fexA__Staphylococcus_lentus__TM_#04
SGFSSFSSLTSLIGSVVALVGFIWRIVTAENPFVPPVLFNNKDYVNTVIIA
>gb|CAD70268_1|ARO_3002704|fexA__Staphylococcus_lentus__TM_#05
DKRLIITGMTLMGLSTLFLSTYASGASPLLVSVGVLGVGIAFAFTNSPANNAAVSALDAD
K
>gb|CAD70268_1|ARO_3002704|fexA__Staphylococcus_lentus__TM_#06
TEPINPLYILDAMSYSDAFLAATGAILIALIAGLGLKKRG
>gb|ABW05394_1|ARO_3004450|TRU-1__Aeromonas_enteropelogenes__TM_#01
MKQRIALSLLALGPLLLVPRVYAAADEPMANIVEKAVQPLLE
>gb|ABW05394_1|ARO_3004450|TRU-1__Aeromonas_enteropelogenes__TM_#02

NERMRQYYRQWSPL
>gb|ABW05394_1|ARO_3004450|TRU-1__Aeromonas_enteropelogenes__TM_#03
EVPDAAMV
>gb|ABW05394_1|ARO_3004450|TRU-1__Aeromonas_enteropelogenes__TM_#04
GANMTGSGDKAVQQ
>gb|ABW05394_1|ARO_3004450|TRU-1__Aeromonas_enteropelogenes__TM_#05
AIMRHLAP
>gb|AEM62948_1|ARO_3003196|tet(45)__Bhargavaea_cecembensis__TM_#01
PSTHLYSNLLLLFAGITIISWLVTIKMHKRSRGNI
>gb|CAE53424_1|ARO_3003016|dfrA20__Pasteurella_multocida__TM_#01
MGIKYSLIVAIGKHREMGA
>gb|CAE53424_1|ARO_3003016|dfrA20__Pasteurella_multocida__TM_#02
ESIPQKYRPLPNRLNFVLTRDKNYSAEGATVIYDLKEVAQHLEGKNLT
>gb|CAE53424_1|ARO_3003016|dfrA20__Pasteurella_multocida__TM_#03
ETGLLNEMYVTQVHNTFEEADTFFPFVNWGEWEEEDILEQDKDEKHLYSFNIKKFTR
>gb|AAQ08905_1|ARO_3001808|OXA-60__Ralstonia_pickettii__TM_#01
VRNDLKRVFD
>gb|AAQ08905_1|ARO_3001808|OXA-60__Ralstonia_pickettii__TM_#02
AGVSGTFVLMDI
>gb|AAQ08905_1|ARO_3001808|OXA-60__Ralstonia_pickettii__TM_#03
ERMQAYVDAFDYGNRQLG
>gb|AAQ08905_1|ARO_3001808|OXA-60__Ralstonia_pickettii__TM_#04
IDQFWLRGPLEISA
>gb|AAQ08905_1|ARO_3001808|OXA-60__Ralstonia_pickettii__TM_#05
EEARFTSRMALKQLPVKPRTWDMV
>gb|AAQ08905_1|ARO_3001808|OXA-60__Ralstonia_pickettii__TM_#06
RMLLIEQQGDAALYAKTGVATEYQPEIGWW
>gb|AAQ08905_1|ARO_3001808|OXA-60__Ralstonia_pickettii__TM_#07
REGDMAKRIPLGKQLM
>gb|ACR57831_1|ARO_3003022|dfrB3__Klebsiella_oxytoca__TM_#01
VAGQFALP
>gb|AAA65957_1|ARO_3002949|vanXA__Enterococcus_faecium__TM_#01
ESLLKAKELAAT
>gb|AAA65957_1|ARO_3002949|vanXA__Enterococcus_faecium__TM_#02
RAVNCFMQWAAQPENNLTKES
>gb|CAC80727_1|ARO_3000476|tet(31)__Aeromonas_salmonicida_subsp__salmonicida__TM_#01
MIGKLIMMNRYITIALLITFLDAT
>gb|CAC80727_1|ARO_3000476|tet(31)__Aeromonas_salmonicida_subsp__salmonicida__TM_#02
EEFSVKESIATHYGFI
>gb|CAC80727_1|ARO_3000476|tet(31)__Aeromonas_salmonicida_subsp__salmonicida__TM_#03
AVCDYTLLSFSSALWMLYLGRMIAGISAATGAVAASMVADHTKKA
>gb|CAC80727_1|ARO_3000476|tet(31)__Aeromonas_salmonicida_subsp__salmonicida__TM_#04
IALIMVIILFPKEQSRPKEIEQDQSKIHEKTTINAPLIHILKPVLLLLML
>gb|CAC80727_1|ARO_3000476|tet(31)__Aeromonas_salmonicida_subsp__salmonicida__TM_#05
WKNETVFILGFILDASAFLLLAFISQVWLVI
>gb|CAC80727_1|ARO_3000476|tet(31)__Aeromonas_salmonicida_subsp__salmonicida__TM_#06
IKTADEHQGKIQGIMVSLTNITGIIGPPIFAFSFAKTVTNWDGTLWLIGAVLYSILLGLY
FLYQKIRAYKQLKSQTA
>gb|AAR20595_1|ARO_3000090|BLA1__Bacillus_anthracis__TM_#01
CVGILGLSITSL
>gb|AAR20595_1|ARO_3000090|BLA1__Bacillus_anthracis__TM_#02
LQVEAKEKTGQVKHKNQATHKE
>gb|AAR20595_1|ARO_3000090|BLA1__Bacillus_anthracis__TM_#03
KIGGPKGYEKALR
>gb|APB03214_1|ARO_3003980|tetA(48)__Paenibacillus_sp__LC231__TM_#01
MEHAYKKKIEPKPGDLAVEAY
>gb|APB03214_1|ARO_3003980|tetA(48)__Paenibacillus_sp__LC231__TM_#02
TVEDSVDMDRACHTVEQVLGVHAN
>gb|APB03214_1|ARO_3003980|tetA(48)__Paenibacillus_sp__LC231__TM_#03
YESNDREVARV
>gb|AAL27444_1|ARO_3002971|vanTE__Enterococcus_faecalis__TM_#01
MKHRANGIDLFRIFAATMVVAIHTFPFQSIAPF
>gb|AAL27444_1|ARO_3002971|vanTE__Enterococcus_faecalis__TM_#02
GRLSLNFSYNNNQRVKKYLYKIGMIYLYSILLYFPLSLLNGTISLKMNILLLLKVF
>gb|AAL27444_1|ARO_3002971|vanTE__Enterococcus_faecalis__TM_#03
TILVTLLLRSIGFKLTVAFSTCLYLVGLGGDSWYGITNQVPLLNKLYTFIFSWSDYTRSG
VFFTPVFLCLGIFAYRVSKKLTASKILNLLFYVFIIGMTFESIFLHRFTNVKHDSMYLLL
PSCALILFLMLLNWQPKLKVKESADLTLLVYILHPLVIVIVHSISKYIPILKNSLLNFLL
VVVCSFILAQLLLNLKRKLRVSKQKIPFERA
>gb|AAD51345_1|ARO_3003052|smeB__Stenotrophomonas_maltophilia__TM_#01
SLLGLLAIL
>gb|AAD51345_1|ARO_3003052|smeB__Stenotrophomonas_maltophilia__TM_#02
WDIETALQAQNTDVSAGELGGQPAL
>gb|AAD51345_1|ARO_3003052|smeB__Stenotrophomonas_maltophilia__TM_#03
AQVVLKADANGSVVRLGDVAKIGLGPESYDS
>gb|AAD51345_1|ARO_3003052|smeB__Stenotrophomonas_maltophilia__TM_#04
GLRHRVDR
>gb|AAD51345_1|ARO_3003052|smeB__Stenotrophomonas_maltophilia__TM_#05
RAMASVPDAQVFITSPPAILGLGDAGGFTLELQDEGGAGHAAAVAARNTLLKEAAK
>gb|AAD51345_1|ARO_3003052|smeB__Stenotrophomonas_maltophilia__TM_#06
AKAQAMGVNPQDVND
>gb|AAD51345_1|ARO_3003052|smeB__Stenotrophomonas_maltophilia__TM_#07
EARMQPQDIERWSVRNQAGQMVPLSSLISTHWTSAPAAVQRYNGISAMEIT
>gb|CAM12479_1|ARO_3000567|tet(40)__uncultured_bacterium__TM_#01
MFLYGLGFLMEGALPWFAP
>gb|CAM12479_1|ARO_3000567|tet(40)__uncultured_bacterium__TM_#02
EDKPIDKVFLRGSQMGQIGGVLGVVLGTLLGNINLQMPLILGGSLCLLLGLVMVRIMPET
NFSPAIEERQGLLKDFVCLFKLNLGFVKGAPVLLALLAITL
>gb|CAM12479_1|ARO_3000567|tet(40)__uncultured_bacterium__TM_#03
DDTVIPVIGPLNSVTWFGVISLIGNGLGILASQLLIARMEKKGTVSRTSVVMSTSAGYIL
FLVLFAVGRSFWFMLLVFLLAGLMRTIKEPVLAAWMNDHVDEKMRATVFSTSGQLDSFGQ
IIGGPIVGLVAQQVSIPWGLVCTAFLLLPALFLVPVAGKKRD
>gb|ENX60508_1|ARO_3001743|OXA-288__Acinetobacter_sp__CIP_70_18__TM_#01
IYDGKKIQ
>gb|ENX60508_1|ARO_3001743|OXA-288__Acinetobacter_sp__CIP_70_18__TM_#02
YALAKQKLPFDQ
>gb|ABA71727_1|ARO_3002926|vanRG__Enterococcus_faecalis__TM_#01
CYNNAADIEKENVLVTEYDINGLVINKNTHKCTL
>gb|XP_753111_1|ARO_3003942|abcA__Aspergillus_fumigatus_Af293__TM_#01
MNESHEAGKNSSTNVEEREEEVLR
>gb|XP_753111_1|ARO_3003942|abcA__Aspergillus_fumigatus_Af293__TM_#02
KTLSLARIAF
>gb|XP_753111_1|ARO_3003942|abcA__Aspergillus_fumigatus_Af293__TM_#03

VGLNNRVCSTVR
>gb|XP_753111_1|ARO_3003942|abcA__Aspergillus_fumigatus_Af293__TM_#04
RHQTNLFRTTSGQKREDKDSYREFAAPFWAQLY
>gb|XP_753111_1|ARO_3003942|abcA__Aspergillus_fumigatus_Af293__TM_#05
SLYWWVRVPRDKKPVAKAE
>gb|CCE94500_1|ARO_3001502|OXA-256__Enterobacter_cloacae__TM_#01
MKTFAAYVI
>gb|CCE94500_1|ARO_3001502|OXA-256__Enterobacter_cloacae__TM_#02
ACLSSTALA
>gb|CCE94500_1|ARO_3001502|OXA-256__Enterobacter_cloacae__TM_#03
WNKEFSAEAVNGVFVLCKSSSKSACATN
>gb|CCE94500_1|ARO_3001502|OXA-256__Enterobacter_cloacae__TM_#04
LARASKEYLP
>gb|CCE94500_1|ARO_3001502|OXA-256__Enterobacter_cloacae__TM_#05
AIIGLETGVIKNEHQ
>gb|CCE94500_1|ARO_3001502|OXA-256__Enterobacter_cloacae__TM_#06
QLRISAVNQVEFLESL
>gb|CCE94500_1|ARO_3001502|OXA-256__Enterobacter_cloacae__TM_#07
LNKLSASKENQLIVKEALVTEAAPEYLVHSKTGFSGVGTESNPGVAWWVGWVEK
>gb|CCE94500_1|ARO_3001502|OXA-256__Enterobacter_cloacae__TM_#08
EVYFFAFNMDIDNE
>gb|AAA88675_1|ARO_3000498|ErmF__Bacteroides_fragilis__TM_#01
NVVAIENDTALVEHLRKLFSDARNVQVVGCDFRNFA
>gb|AAA88675_1|ARO_3000498|ErmF__Bacteroides_fragilis__TM_#02
IVCLSPSQWL
>gb|CAC35728_1|ARO_3001424|OXA-29__Fluoribacter_gormanii__TM_#01
MKKLSVLLWLTLFYCGTIW
>gb|CAC35728_1|ARO_3001424|OXA-29__Fluoribacter_gormanii__TM_#02
QFLQKIVNKKLSVN
>gb|CAC35728_1|ARO_3001424|OXA-29__Fluoribacter_gormanii__TM_#03
TKHIADSKKHV
>gb|ACH87088_1|ARO_3000196|tet32__Escherichia_coli__TM_#01
EDSLDIQELQYEKCKRTRC
>gb|AAA26742_1|ARO_3001304|ErmS__Streptomyces_fradiae__TM_#01
MARAPRSPHPARSRETSRAHPPYGTRADRAPGRGRDRDRSPDSPGNTSSRDGGRSPDRAR
RELSQNFLARRAVAERVARLVRPAPGG
>gb|AAA26742_1|ARO_3001304|ErmS__Streptomyces_fradiae__TM_#02
YCGRLVAHEIDPRLLPALRDRFGGPHHAHVRISGGDFLAAPVPREPFALAGNIPYSRTAG
IVDWALRART
>gb|AAA26742_1|ARO_3001304|ErmS__Streptomyces_fradiae__TM_#03
RSVVVAYVTPEQWLTVFRTLRPVRSRPAGR
>gb|CAA88265_1|ARO_3002898|SAT-3__Escherichia_coli__TM_#01
MTPQSMRELVICRASDADVLQLARCDFSFEVTAELEEPFDDMRSVPVKPPYLKNYGFDAD
ELVEHMNNSAGALFVARADNCLVGYLAVSQSWNEYAVIDDIAVDVPYRGSGVSRLLMDAA
VDWARNVPSAGVRLETQSVNLA
>gb|CAA88265_1|ARO_3002898|SAT-3__Escherichia_coli__TM_#02
RGLHPGSREVALFWYLSF
>gb|AAL51021_2|ARO_3002598|ANT(3__)-Ii-AAC(6_)-IId_fusion_protein__Serratia_marcescens__TM_#01
AIERLPDQHK
>gb|AAL51021_2|ARO_3002598|ANT(3__)-Ii-AAC(6_)-IId_fusion_protein__Serratia_marcescens__TM_#02
ELLFNDPE
>gb|TKW44661_1|ARO_3005040|YajC__Pseudomonas_aeruginosa__TM_#01
MSFLIPAAYADAAAPAAAGPAGT
>gb|TKW44661_1|ARO_3005040|YajC__Pseudomonas_aeruginosa__TM_#02
DEVVTSGGIAGKVTKVADDFVVVEVSDNVELKFQKAAIAATLPKGTLKAI
>gb|AAF40763_1|ARO_3003961|farA__Neisseria_meningitidis_MC58__TM_#01
KGGTVRKVLHDDTDAVKKGDVLAVLDDDNDVLAYERAKNELVQAVRQNRRQNAATSQAGA
QVALRRADLARAQDDLRRRSALAESGAVSAEELAHARAAVSQAQAAVKAALAEESSARAA
LGGQVSLREQPAVQTAIGR
>gb|AAF40763_1|ARO_3003961|farA__Neisseria_meningitidis_MC58__TM_#02
AELVSDLYGKQ
>gb|AAF40763_1|ARO_3003961|farA__Neisseria_meningitidis_MC58__TM_#03
TSAAGAPVSKTPGAALPEMESTDWSEVDRTVDEILGQSAP
>gb|AAA74437_1|ARO_3000378|MexB__Pseudomonas_aeruginosa__TM_#01
MLAAQNPALQR
>gb|AEA37904_1|ARO_3003112|lsaC__Streptococcus_agalactiae__TM_#01
KGKISKSVDFIKFPPNLSDTSKLGIELYRELISDEEE
>gb|AEA37904_1|ARO_3003112|lsaC__Streptococcus_agalactiae__TM_#02
CDLSSYYGKK
>gb|AEA37904_1|ARO_3003112|lsaC__Streptococcus_agalactiae__TM_#03
SFVEDIANKIIRL
>gb|AAL27442_1|ARO_3002907|vanE__Enterococcus_faecalis__TM_#01
MKTVAIIFGGVSSEYEVSLKSAVAIIKNMESIDYNVMKIGITEEGHWYLFEGTTDKIKKD
RWFLDESCEEIVVDFAKKSFVLKNSKKIIKP
>gb|AAL27442_1|ARO_3002907|vanE__Enterococcus_faecalis__TM_#02
AIGVKSTPSMIIEKGQDLQKVDAFAK
>gb|AAL27442_1|ARO_3002907|vanE__Enterococcus_faecalis__TM_#03
RKSDLYKAIDEASKYDSRILIQKEVK
>gb|AAL27442_1|ARO_3002907|vanE__Enterococcus_faecalis__TM_#04
NLVTAEILLPAKLSIDKKEDIQMKAKK
>gb|AAL27442_1|ARO_3002907|vanE__Enterococcus_faecalis__TM_#05
MMMNEIGMDYKEIIENLLVLAVENHEKKLSTID
>gb|BAC11911_1|ARO_3003551|emeA__Enterococcus_faecalis_ATCC_29212__TM_#01
IAGVLSDKIGRKKMIAT
>gb|BAC11911_1|ARO_3003551|emeA__Enterococcus_faecalis_ATCC_29212__TM_#02
LAAVEAKKG
>gb|BAC11911_1|ARO_3003551|emeA__Enterococcus_faecalis_ATCC_29212__TM_#03
CQLFFFDAIVQ
>gb|BAA34300_1|ARO_3003033|mexY__Pseudomonas_aeruginosa_PAO1__TM_#01
AHPRDGGAIRLRDVARVEFGQSEYGFVSRVNQMT
>gb|BAA34300_1|ARO_3003033|mexY__Pseudomonas_aeruginosa_PAO1__TM_#02
RATLDELSRYFPEGVSYNIPYDTSA
>gb|BAA34300_1|ARO_3003033|mexY__Pseudomonas_aeruginosa_PAO1__TM_#03
GRYRNAVAGILARPIRWMLVYTLVIG
>gb|BAA34300_1|ARO_3003033|mexY__Pseudomonas_aeruginosa_PAO1__TM_#04
HVGAIVERINQRFAGLPNR
>gb|BAA34300_1|ARO_3003033|mexY__Pseudomonas_aeruginosa_PAO1__TM_#05
VKARDQLLARAAEDPRLANVMFAGQGEAPQIRLDIDRRKAET
>gb|BAA34300_1|ARO_3003033|mexY__Pseudomonas_aeruginosa_PAO1__TM_#06
GAAGDVRQGRLDPRPAATDPLQRLSLVQPRGPGRAGLQQREAMQAMEQLMQGTARGIRPR
VVRPVLRRTPV
>gb|BAH45481_1|ARO_3002815|clbB__Brevibacillus_brevis_NBRC_100599__TM_#01
MKLTSKYETIRRILSEC

>gb|BAH45481_1|ARO_3002815|clbB__Brevibacillus_brevis_NBRC_100599__TM_#02
FLRDELNRILGPNVCSIAPVKELT
>gb|BAH45481_1|ARO_3002815|clbB__Brevibacillus_brevis_NBRC_100599__TM_#03
QSDPSRIKAFVRILKSR
>gb|BAB41874_1|ARO_3000815|mgrA__Staphylococcus_aureus_subsp__aureus_N315__TM_#01
ETIRPELSNASDKVASASSLSQDEVKELNRLLGKVIHAFDETKEK
>gb|BAM46120_1|ARO_3003677|AAC(6_)-Iaj__Pseudomonas_aeruginosa__TM_#01
MEYSIINIVEQNNYQIDAARILTNTFLDIGNKTWPTIQSAIDEVEECIDLPNICIGLIHN
NQ
>gb|BAM46120_1|ARO_3003677|AAC(6_)-Iaj__Pseudomonas_aeruginosa__TM_#02
TDYQSKGIGSVLLAEVEKRAREV
>gb|BAM46120_1|ARO_3003677|AAC(6_)-Iaj__Pseudomonas_aeruginosa__TM_#03
TIDENNIFDAIQ
>gb|AGW81558_1|ARO_3002631|spd__Staphylococcus_aureus__TM_#01
MEEPNKQIDNVLIELKRLFSKDLLGVYLYGSYVKGGLKKDSDVDFLVIINRDMTKEEKRI
LISKIMPISKEIGEDTSLKYIELTVLNYHENEN
>gb|AGW81558_1|ARO_3002631|spd__Staphylococcus_aureus__TM_#02
EDYLNYFIPEKNNNIDLTILLYQAKLSSISIYGENNINNLIPDVPFIDLQKAIKESSKEL
IKDFYGDETNVILTLCRMIVTYETGKFYSKDLAGSMIIENLSENLSIEENNLISLAISSY
KNGNSVDWELFPVKSVIKKLYAYLNYKL
>gb|CAC04522_1|ARO_3002514|OCH-1__Ochrobactrum_anthropi__TM_#01
STTLLIGFLTTAA
>gb|CAC04522_1|ARO_3002514|OCH-1__Ochrobactrum_anthropi__TM_#02
MLGGYGLATGAF
>gb|CAC04522_1|ARO_3002514|OCH-1__Ochrobactrum_anthropi__TM_#03
LSDPATKWAPELA
>gb|CAC04522_1|ARO_3002514|OCH-1__Ochrobactrum_anthropi__TM_#04
SSFDKITM
>gb|CAC04522_1|ARO_3002514|OCH-1__Ochrobactrum_anthropi__TM_#05
DDSSMLAYFK
>gb|CAC04522_1|ARO_3002514|OCH-1__Ochrobactrum_anthropi__TM_#06
ARFVELNIDSSSLE
>gb|CAC04522_1|ARO_3002514|OCH-1__Ochrobactrum_anthropi__TM_#07
DFQKAVAATHTGYY
>gb|CAC04522_1|ARO_3002514|OCH-1__Ochrobactrum_anthropi__TM_#08
ANNQGLGWEFYNYPTA
>gb|CAC04522_1|ARO_3002514|OCH-1__Ochrobactrum_anthropi__TM_#09
KAAYRILQALDNKQ
>gb|AAG04750_1|ARO_3004077|PmpM__Pseudomonas_aeruginosa_PAO1__TM_#01
MNSPALPLSRGLRIR
>gb|AAG04750_1|ARO_3004077|PmpM__Pseudomonas_aeruginosa_PAO1__TM_#02
PILGLMKVRPELIGPSLLYLKGIAL
>gb|AAG04750_1|ARO_3004077|PmpM__Pseudomonas_aeruginosa_PAO1__TM_#03
KMGGPGCGWATGSVMWFMFLGMLFWVNKASIYRASQLFSRWEWPDRAT
>gb|AAG04750_1|ARO_3004077|PmpM__Pseudomonas_aeruginosa_PAO1__TM_#04
GAAIMLCIRLARSARRFIRQHERLQREDAEAASVLGR
>gb|AJP77080|ARO_3003720|SPG-1__Sphingomonas_sp__ALS-13__TM_#01
MRPLILLASALAMTAPSGALAQSAEQRAKWNG
>gb|AJP77080|ARO_3003720|SPG-1__Sphingomonas_sp__ALS-13__TM_#02
SGLSAYLIAG
>gb|AJP77080|ARO_3003720|SPG-1__Sphingomonas_sp__ALS-13__TM_#03
KLGFKLSDVKYLLSNHSHVDHAGGLAELKRLTGAQMVANVADKPDLEAGTTIGRTDIADF
PAVKVDRVIGDGDRLTLGPIALVAILTPGHTKGATSWTTRIGSKNVIFTSSISVAGQNLI
NNINYPNAA
>gb|AJP77080|ARO_3003720|SPG-1__Sphingomonas_sp__ALS-13__TM_#04
LKADIFLSFHAEAFALDEKRARAQAGATDAFVDPNELSRQVDLAEKAFDATLAKQQAANA
DRR
>gb|CAA83855_1|ARO_3000380|FosC__Pseudomonas_syringae__TM_#01
MMTSIMFSMLSDITRIFVEQGLRVYPFQSSALLGVDEEGRVTLHARQLATAMASGYMPLL
TGDLLLRGEQEAQVFSSDNIAPLLAADFEVRRVLYYSDVAGVYDQGNALVPWVGNANAAC
MEACVGASSM
>gb|CAA83855_1|ARO_3000380|FosC__Pseudomonas_syringae__TM_#02
QLARLGVVSEVLSFECFDRVHLSLCGLRQFGTVFLSE
>gb|EPF73263|ARO_3004086|APH(3_)-VIII__Acinetobacter_rudis_CIP_110305__TM_#01
MKLPQKIRNFIGNNRLIV
>gb|EPF73263|ARO_3004086|APH(3_)-VIII__Acinetobacter_rudis_CIP_110305__TM_#02
CFERNRETFFLKVSSVQYATTTYSVAREAQMMLWLADKINVPELVFSEIDQNFEYMLSKS
IDAQPISDLSLAQSELIMLYQDVLSQLRSVPVQNCPFNSDINSRLQESQYFMEIGLLNQV
DDENIDIELWGEHQSYLELWTELNNHRVKENLVFTHGDITDSNIFVDQSNK
>gb|EPF73263|ARO_3004086|APH(3_)-VIII__Acinetobacter_rudis_CIP_110305__TM_#03
QKFLKQLSFDDLSKRQYFLKLDELN
>gb|CCP44758_1|ARO_3000392|Erm(37)__Mycobacterium_tuberculosis_H37Rv__TM_#01
MSALGRSRRAWGWHRLHDEWAARVVSAAAVRPGELVFDIGAGEGALTAH
>gb|CCP44758_1|ARO_3000392|Erm(37)__Mycobacterium_tuberculosis_H37Rv__TM_#02
RRVGVLRERFPGITVVHADAASI
>gb|CCP44758_1|ARO_3000392|Erm(37)__Mycobacterium_tuberculosis_H37Rv__TM_#03
SSRLLRTLLAPNSGLVAADLVLQRALVCKFASRNARRFTLTVGLMLPRRAFLPPPHVDSA
VLVVRRRKCGDWQGR
>gb|AFQ90085_1|ARO_3001610|OXA-243__Achromobacter_xylosoxidans__TM_#01
ALGAALSL
>gb|AFQ90085_1|ARO_3001610|OXA-243__Achromobacter_xylosoxidans__TM_#02
LCTVVADAADGRI
>gb|AFQ90085_1|ARO_3001610|OXA-243__Achromobacter_xylosoxidans__TM_#03
RYTPASTFKL
>gb|AFQ90085_1|ARO_3001610|OXA-243__Achromobacter_xylosoxidans__TM_#04
IALMGADAGIL
>gb|AFQ90085_1|ARO_3001610|OXA-243__Achromobacter_xylosoxidans__TM_#05
DVSGEPGKHNG
>gb|AFQ90085_1|ARO_3001610|OXA-243__Achromobacter_xylosoxidans__TM_#06
GKTGTGSPG
>gb|AFQ90085_1|ARO_3001610|OXA-243__Achromobacter_xylosoxidans__TM_#07
PNAGLRARD
>gb|BAD95494_1|ARO_3002895|SAT-1__Escherichia_coli__TM_#01
FDVHLSDQGFELSTRSVSPYRKDYISDDDSDEDSACYGAFIDQELVGKIELNSTWNDLAS
IEHIVVSHTHRGKGVAHSLIEFAKKWALSRQLL
>gb|BAD95494_1|ARO_3002895|SAT-1__Escherichia_coli__TM_#02
SNETAMYWYWFSGAQDDA
>gb|AAK60186_1|ARO_3003014|dfrA16__Escherichia_coli__TM_#01
NFSTNDEGVMVFSSIQDALINLE
>gb|AAK60186_1|ARO_3003014|dfrA16__Escherichia_coli__TM_#02
RDGDIVFPEIPDTFK
>gb|ABX54691_1|ARO_3002927|vanRL__Enterococcus_faecalis__TM_#01
MTDRIVVVDDEQ

>gb|ABX54691_1|ARO_3002927|vanRL__Enterococcus_faecalis__TM_#02
QVTTFYKGEDFLTYIARES
>gb|ABX54691_1|ARO_3002927|vanRL__Enterococcus_faecalis__TM_#03
RKVENESVIEFNKDGLT
>gb|ABX54691_1|ARO_3002927|vanRL__Enterococcus_faecalis__TM_#04
KEITVTPIEFNLLLYLFEHQGV
>gb|CAG41812_1|ARO_3003733|fusC__Staphylococcus_aureus_subsp__aureus_MSSA476__TM_#01
MNKIEVYKFVKVKQLVYQLIKLYRTNDMNSHKTQKDFLLNEINDIFKEKDIDISDFITSI
DDVKLTKKKAEHLLNELKVYIQDFEIPSSSQLEKIFRKVKKLKRPDINLIDTKEISYLGW
NDNSSNRKYIVYKNLDDKFEGIYGEISPN
>gb|CAG41812_1|ARO_3003733|fusC__Staphylococcus_aureus_subsp__aureus_MSSA476__TM_#02
TSLFLNKTKHNKSS
>gb|CAG41812_1|ARO_3003733|fusC__Staphylococcus_aureus_subsp__aureus_MSSA476__TM_#03
FKCNQNLDDINNLYEFIVKIK
>gb|ABA42119_2|ARO_3000865|oleD__Streptomyces_antibioticus__TM_#01
ATEEATADL
>gb|ABA42119_2|ARO_3000865|oleD__Streptomyces_antibioticus__TM_#02
ERQEPVGDRPNGG
>gb|AAC69327_1|ARO_3000598|Erm(31)__Streptomyces_venezuelae__TM_#01
HGRPITAVELDGRRAQRLGARTPGHVTVVHHDFLQYPLPRN
>gb|AAC69327_1|ARO_3000598|Erm(31)__Streptomyces_venezuelae__TM_#02
APWYEFDLHS
>gb|AAC69327_1|ARO_3000598|Erm(31)__Streptomyces_venezuelae__TM_#03
SAPLVGQVKTYQDFVRQVFTGKGNGLKEILRRTGRISQRDLATWLRRNE
>gb|AAC69327_1|ARO_3000598|Erm(31)__Streptomyces_venezuelae__TM_#04
TGGTADGSFDGTAGGGAAGSHGAARVGAGHPGGRVSASRRGVPQARRGRGHAVRSSTGTE
PRWGRGRAESA
>gb|ENV38192_1|ARO_3001722|OXA-266__Acinetobacter_venetianus_RAG-1___CIP_110063__TM_#01
MRKKFKVALLCSSLCLSLGLVACHSLNSELQIAEQQKQQQKISKLFVNAKTEGVFVTYDG
QKIHEYGNALNRAQ
>gb|ENV38192_1|ARO_3001722|OXA-266__Acinetobacter_venetianus_RAG-1___CIP_110063__TM_#02
TYALANEQLAFDIPVQQQVKQMLLVDQM
>gb|ENV38192_1|ARO_3001722|OXA-266__Acinetobacter_venetianus_RAG-1___CIP_110063__TM_#03
TEHVEARKTIVYEALQQLGLI
>gb|AAA25293_1|ARO_3000192|tetS__Listeria_monocytogenes__TM_#01
PDVYQDIKDKLSDDIIIKQTVNLNLKPYVIDYTEPEQWETVIVGNDYLLEKYTIGKTLNI
AELEKEENERIQSC
>gb|AAA25293_1|ARO_3000192|tetS__Listeria_monocytogenes__TM_#02
NIGIKQLIEVITSKLFSPTQLNSDKLCGNVFKVEYSDDGQRL
>gb|AAA25293_1|ARO_3000192|tetS__Listeria_monocytogenes__TM_#03
RQIDKAEPGEIIILKNELLKLNNVLGDKKRLPHREIL
>gb|AAA25293_1|ARO_3000192|tetS__Listeria_monocytogenes__TM_#04
EPCKSVQREKLLDALF
>gb|AAA25293_1|ARO_3000192|tetS__Listeria_monocytogenes__TM_#05
VTHEIVLSFLGEVQMEVTCTLIQEKYHIEIETRK
>gb|AAA25293_1|ARO_3000192|tetS__Listeria_monocytogenes__TM_#06
NILNTKLKGN
>gb|AXY65402_1|ARO_3005021|cfr(D)__Enterococcus_faecium__TM_#01
MLQKQLTKYQQIEKVLKECQQPNYRMKQILHCIFKEKKTDFNEMSVLPKNLRDTLTTEIG
TSLLTVEPLIEQKSQQVRKVLFNLSGDNRIETVN
>gb|AXY65402_1|ARO_3005021|cfr(D)__Enterococcus_faecium__TM_#02
KIGLRKNLTAEEITDQVLYFLLK
>gb|AXY65402_1|ARO_3005021|cfr(D)__Enterococcus_faecium__TM_#03
FFEALDIFMNPDMFQLSPRRLSVSTIGVIPKIKRLTA
>gb|AXY65402_1|ARO_3005021|cfr(D)__Enterococcus_faecium__TM_#04
PDVNDSLTHAVALADLLRSRYKK
>gb|AXY65402_1|ARO_3005021|cfr(D)__Enterococcus_faecium__TM_#05
RTFKQVDEKQVKLFYQTLISK
>gb|AXY65402_1|ARO_3005021|cfr(D)__Enterococcus_faecium__TM_#06
IKNKKTIQVND
>gb|ABB89122_1|ARO_3002861|dfrA5__Salmonella_enterica_subsp__enterica_serovar_Enteritidis__TM_#01
MKVSLMAA
>gb|ABB89122_1|ARO_3002861|dfrA5__Salmonella_enterica_subsp__enterica_serovar_Enteritidis__TM_#02
EPEGDVFFP
>gb|AAG07595_1|ARO_3000809|opmD__Pseudomonas_aeruginosa_PAO1__TM_#01
MKRSYPNLSRLALALAVGTG
>gb|AAG07595_1|ARO_3000809|opmD__Pseudomonas_aeruginosa_PAO1__TM_#02
PPRVASEHL
>gb|AAG07595_1|ARO_3000809|opmD__Pseudomonas_aeruginosa_PAO1__TM_#03
APLAKNLPLGDVDRLIL
>gb|AAG07595_1|ARO_3000809|opmD__Pseudomonas_aeruginosa_PAO1__TM_#04
AVEAQSDAALARYQRSLLLAQEDVGNALNQLAEHQRRLVALFQSATHGA
>gb|AAA25717_1|ARO_3002624|ANT(4_)-IIa__Pseudomonas_aeruginosa__TM_#01
MHLTITYWIDRLREAYPH
>gb|AAA25717_1|ARO_3002624|ANT(4_)-IIa__Pseudomonas_aeruginosa__TM_#02
DVLVSDEEVEEYRTWIEPVGERLVHISVAVEWVTGWERDSADPSSWSYGLPTQETTQLLW
AADENIRRRLDRPFKV
>gb|AAA25717_1|ARO_3002624|ANT(4_)-IIa__Pseudomonas_aeruginosa__TM_#03
MVRGDDLAVYQAAQVVGKLIPTLLVPINPPTYARFAREAIDRILAFPNVPEGFAADWLTC
MGLVDRRTHDPQPTRPNEWCAARSRFCRRMRTSSVRISRGCWKQDWYLRISART
>gb|AAA23215_1|ARO_3002687|catQ__Clostridium_perfringens__TM_#01
RHKEFRTCFDQK
>gb|AAA23215_1|ARO_3002687|catQ__Clostridium_perfringens__TM_#02
PRFYYNYLEDIRNYSDVLNFMPKTGEPANTIN
>gb|AAA23215_1|ARO_3002687|catQ__Clostridium_perfringens__TM_#03
IPIFTLGKYFQQD
>gb|AAA23215_1|ARO_3002687|catQ__Clostridium_perfringens__TM_#04
LASNYETWLGEK
>gb|AAB81957_1|ARO_3004784|FAR-1__Nocardia_farcinica__TM_#01
MAAAAAIALLGGCGADAGSEPATTAASTTAPSTATDAATAEFAALEQR
>gb|AAB81957_1|ARO_3004784|FAR-1__Nocardia_farcinica__TM_#02
AAVAANYRA
>gb|AAB81957_1|ARO_3004784|FAR-1__Nocardia_farcinica__TM_#03
NNDVAVAWTETGDPIVIALLSHRTDPA
>gb|ABA00479_1|ARO_3001859|IMI-2__Enterobacter_asburiae__TM_#01
GVYALDTGSGKSF
>gb|ABA00479_1|ARO_3001859|IMI-2__Enterobacter_asburiae__TM_#02
NGMSLGDMAA
>gb|ABA00479_1|ARO_3001859|IMI-2__Enterobacter_asburiae__TM_#03
KNRAPLIISVYTTK
>gb|CAA25854_1|ARO_3002659|APH(6)-Ic__Escherichia_coli__TM_#01
MERWRLLRDGELLTTH
>gb|CAA25854_1|ARO_3002659|APH(6)-Ic__Escherichia_coli__TM_#02

SGPPPDLHPLQEWFQPLFRLAAEHAALAPAASVARQLLAAPRE
>gb|CAA25854_1|ARO_3002659|APH(6)-Ic__Escherichia_coli__TM_#03
LEARLSIVVAT
>gb|CAA25854_1|ARO_3002659|APH(6)-Ic__Escherichia_coli__TM_#04
GEGEGAAIDLAVNAMARRLLD
>gb|AAG05661_1|ARO_3004107|Pseudomonas_aeruginosa_soxR__Pseudomonas_aeruginosa_PAO1__TM_#01
QRNAGNQRRFSRET
>gb|AAG05661_1|ARO_3004107|Pseudomonas_aeruginosa_soxR__Pseudomonas_aeruginosa_PAO1__TM_#02
QTLPAGRSPSAADWARLSAQWKEDLTERIDK
>gb|AAG05661_1|ARO_3004107|Pseudomonas_aeruginosa_soxR__Pseudomonas_aeruginosa_PAO1__TM_#03
SAEGPGAHWLDAEGREHDG
>gb|BAD00739_1|ARO_3002607|aadA7__Vibrio_fluvialis__TM_#01
AVRQALLVDLLEVSAS
>gb|BAD00739_1|ARO_3002607|aadA7__Vibrio_fluvialis__TM_#02
VVLAGSAAKDL
>gb|BAD00739_1|ARO_3002607|aadA7__Vibrio_fluvialis__TM_#03
TWAMARLPAQHQPILLNAKRAYLGQEEDYLPARADQVAALIKFVKYEAVKLLGASQ
>gb|AAS68233_1|ARO_3000391|norA__Staphylococcus_epidermidis__TM_#01
VAGTLGVVAFIMSVLLIHNPQKATTDGFHQYQPELF
>gb|AAS68233_1|ARO_3000391|norA__Staphylococcus_epidermidis__TM_#02
GLKSRRKEAN
>gb|ABX54687_1|ARO_3002910|vanL__Enterococcus_faecalis__TM_#01
MMKLKKIAIIFGGQSSEYEVSLKSTVSVLETLSTCNFEIIKIGIDLGGKWYLTTSNNKDI
EYDVWQTDPSLQEIIPCFNNRGFYNKTTNKYFRPDVLFPILHGGT
>gb|ABX54687_1|ARO_3002910|vanL__Enterococcus_faecalis__TM_#02
YLLHEFAQSVGVKSAPTLIIRTRNCKDEIDKFIEKND
>gb|ABX54687_1|ARO_3002910|vanL__Enterococcus_faecalis__TM_#03
NKVNEPDKLEDALTEAFKYSKSVIIQKAIIGREIGCAVLGNEK
>gb|ABX54687_1|ARO_3002910|vanL__Enterococcus_faecalis__TM_#04
NSDFFDYTEKYQMISAKVNIPASISVEFSNEMKKQAQLL
>gb|ABX54687_1|ARO_3002910|vanL__Enterococcus_faecalis__TM_#05
KMMEAVGVTYKEIITKLINLAEEKYYG
>gb|AGV10830_1|ARO_3002647|APH(3_)-IIIa__Campylobacter_coli_CVM_N29710__TM_#01
NLYLKMTDSRYK
>gb|AGV10830_1|ARO_3002647|APH(3_)-IIIa__Campylobacter_coli_CVM_N29710__TM_#02
HFERHDGWSNLLMSEADGVLCSEEYEDEQSPEKIIELY
>gb|AGV10830_1|ARO_3002647|APH(3_)-IIIa__Campylobacter_coli_CVM_N29710__TM_#03
NNDLADVDC
>gb|AAA26793|ARO_3003748|oleC__Streptomyces_antibioticus__JM_#01__[gb|AAA26793|ARO_3003748|oleC__Streptomyces_antibioticus__w=0.004,gb|APB03214_1|ARO_30
03980|tetA(48)__Paenibacillus_sp__LC231__w=0.001]
MTVRGLVKHYGETKALDGVDLDVREGTVMGVLGP
>gb|AAA26793|ARO_3003748|oleC__Streptomyces_antibioticus__JM_#02__[gb|AAA26793|ARO_3003748|oleC__Streptomyces_antibioticus__w=0.003,gb|AAR96051_1|ARO_3
002894|otrC__Streptomyces_rimosus__w=0.000]
LSRKEARARADELLERFSLTEAARRPAGTYSGGM
>gb|AAA26793|ARO_3003748|oleC__Streptomyces_antibioticus__JM_#03__[gb|AAA26793|ARO_3003748|oleC__Streptomyces_antibioticus__w=0.005,gb|AAR96051_1|ARO_3
002894|otrC__Streptomyces_rimosus__w=0.001,gb|AAR96051_1|ARO_3002894|otrC__Streptomyces_rimosus__w=0.003,gb|AAR96051_1|ARO_3002894|otrC__Streptomyces_ri
mosus__w=0.000]
NEVWDEVKAMVGDGVTVLLTTQYMEEAEQLASEL
>gb|AAA65953_1|ARO_3002919|vanRA__Enterococcus_faecium__JM_#01__[gb|AAA65953_1|ARO_3002919|vanRA__Enterococcus_faecium__w=0.179,gb|BAE85478_1|ARO_3
003728|vanRI__Desulfitobacterium_hafniense_Y51__w=0.053,gb|ABA71727_1|ARO_3002926|vanRG__Enterococcus_faecalis__w=0.053,gb|AAM09851_1|ARO_3002923|vanRD_
_Enterococcus_faecium__w=0.063,gb|AAL27445_1|ARO_3002924|vanRE__Enterococcus_faecalis__w=0.058]
NENYTVFKYYTAKEALECIDKSEIDLAILDIMLP
>gb|AAA65953_1|ARO_3002919|vanRA__Enterococcus_faecium__JM_#02__[gb|AAA65953_1|ARO_3002919|vanRA__Enterococcus_faecium__w=0.064,gb|BAE85478_1|ARO_3
003728|vanRI__Desulfitobacterium_hafniense_Y51__w=0.013,gb|AEP40503_1|ARO_3002929|vanRN__Enterococcus_faecium__w=0.009,gb|ABA71727_1|ARO_3002926|vanRG_
_Enterococcus_faecalis__w=0.002,gb|ABX54691_1|ARO_3002927|vanRL__Enterococcus_faecalis__w=0.002]
GTSGLTICQKIRDKHTYPIIMLTGKDTEVDKITG
>gb|AAA65953_1|ARO_3002919|vanRA__Enterococcus_faecium__JM_#03__[gb|AAA65953_1|ARO_3002919|vanRA__Enterococcus_faecium__w=0.021,gb|BAE85478_1|ARO_3
003728|vanRI__Desulfitobacterium_hafniense_Y51__w=0.003]
AQLRRYKKFSGVKEQNENVIVHSGLVINVNTHEC
>gb|AAB58161_1|ARO_3003010|ceoB__Burkholderia_cepacia__JM_#01__[gb|AAB58161_1|ARO_3003010|ceoB__Burkholderia_cepacia__w=0.016,gb|CAJ77856_1|ARO_300077
7|adeF__Acinetobacter_baumannii_AYE__w=0.007]
VRKTMAELKQDMPAGVDYKIVYDPTQFVRSSIKA
>gb|AAB58161_1|ARO_3003010|ceoB__Burkholderia_cepacia__JM_#02__[gb|AAB58161_1|ARO_3003010|ceoB__Burkholderia_cepacia__w=0.015,gb|CAJ77856_1|ARO_300077
7|adeF__Acinetobacter_baumannii_AYE__w=0.007]
TLMLGVYLVLVGATVLVSKVVPGGFVPAQDKEYL
>gb|AAB58161_1|ARO_3003010|ceoB__Burkholderia_cepacia__JM_#03__[gb|AAB58161_1|ARO_3003010|ceoB__Burkholderia_cepacia__w=0.034,gb|CAJ77856_1|ARO_300077
7|adeF__Acinetobacter_baumannii_AYE__w=0.005,gb|CAJ77856_1|ARO_3000777|adeF__Acinetobacter_baumannii_AYE__w=0.012]
VTLKPFAERHGKALSAGAIAGALNQKYGAMKDSF
>gb|AAC73564_1|ARO_3000216|acrB__Escherichia_coli_str__K-12_substr__MG1655__QM73_#01__[gb|AAC73564_1|ARO_3000216|acrB__Escherichia_coli_str__K-
12_substr__MG1655__w=0.038]
VINTDGTMTQEDISDYVAANMKDAISRTSGVGD
>gb|AAC74000_1|ARO_3003950|msbA__Escherichia_coli_str__K-12_substr__MG1655__JM_#01__[gb|AAC74000_1|ARO_3003950|msbA__Escherichia_coli_str__K-
12_substr__MG1655__w=0.031]
NRMRLQGMKMVSASSISDPIIQLIASLALAFVLY
>gb|AAC75089_1|ARO_3003577|ugd__Escherichia_coli_str__K-12_substr__MG1655__JM_#01__[gb|AAC75089_1|ARO_3003577|ugd__Escherichia_coli_str__K-
12_substr__MG1655__w=0.026]
PIVDKEIQQFLQSDKIHFNATLDKNEAYRDADYV
>gb|AAC75089_1|ARO_3003577|ugd__Escherichia_coli_str__K-12_substr__MG1655__JM_#02__[gb|AAC75089_1|ARO_3003577|ugd__Escherichia_coli_str__K-
12_substr__MG1655__w=0.001]
VPVGFTAAMHKKYRTENIIFSPEFLREGKALYDN
>gb|AAC75135_2|ARO_3000792|mdtA__Escherichia_coli_str__K-12_substr__MG1655__JM_#01__[gb|AAC75135_2|ARO_3000792|mdtA__Escherichia_coli_str__K-
12_substr__MG1655__w=0.032]
AWDRTNSKKLSEGTLLSLDNQIDATTGTIKVKAR
>gb|AAC75136_1|ARO_3000793|mdtB__Escherichia_coli_str__K-12_substr__MG1655__QM60_#01__[gb|AAC75136_1|ARO_3000793|mdtB__Escherichia_coli_str__K-
12_substr__MG1655__w=0.081]
SDGGVVPLSSIAKIEQRFAPLSINHLDQFPVTT
>gb|AAC75137_1|ARO_3000794|mdtC__Escherichia_coli_str__K-12_substr__MG1655__QM59_#01__[gb|AAC75137_1|ARO_3000794|mdtC__Escherichia_coli_str__K-
12_substr__MG1655__w=0.077]
WMLKASKPREQKRLRGFGRMLVALQQGYGKSLK
>gb|AAC75271_1|ARO_3003952|yojI__Escherichia_coli_str__K-12_substr__MG1655__JM_#01__[gb|AAC75271_1|ARO_3003952|yojI__Escherichia_coli_str__K-
12_substr__MG1655__w=0.018]
LNRERAEYVFNNLYIPDAQEYRHHIIRADTFHLS
>gb|AAC75271_1|ARO_3003952|yojI__Escherichia_coli_str__K-12_substr__MG1655__JM_#02__[gb|AAC75271_1|ARO_3003952|yojI__Escherichia_coli_str__K-
12_substr__MG1655__w=0.086]
LLGPEGKPANPQLVEKWLAQLKMAHKLELSNGRI
>gb|AAC75429_1|ARO_3000833|evgS__Escherichia_coli_str__K-12_substr__MG1655__QM51_#01__[gb|AAC75429_1|ARO_3000833|evgS__Escherichia_coli_str__K-
12_substr__MG1655__w=0.140]
AVKITTSLGHIDDNHAVIKMTIMDSGSGLSQEE
>gb|AAC75733_1|ARO_3000074|emrB__Escherichia_coli_str__K-12_substr__MG1655__JM_#01__[gb|AAC75733_1|ARO_3000074|emrB__Escherichia_coli_str__K-
12_substr__MG1655__w=0.044]
IIILTVVAVVAICFLIVWELTDDNPIVDLSLFKS
>gb|AAC76093_1|ARO_3002986|bacA__Escherichia_coli_str__K-12_substr__MG1655__JM_#01__[gb|AAC76093_1|ARO_3002986|bacA__Escherichia_coli_str__K-
12_substr__MG1655__w=0.002]

```
MSDMHSLLIAAILGVVEGLTEFLPVSSTGHMIIV
>gb|AAC76093_1|ARO_3002986|bacA__Escherichia_coli_str__K-12_substr__MG1655__JM_#02__[gb|AAC76093_1|ARO_3002986|bacA__Escherichia_coli_str__K-
12_substr__MG1655__w=0.012]
GHLLGFEGDTAKTFEVVIQLGSILAVVVMFWRRL
>gb|AAC76093_1|ARO_3002986|bacA__Escherichia_coli_str__K-12_substr__MG1655__JM_#03__[gb|AAC76093_1|ARO_3002986|bacA__Escherichia_coli_str__K-
12_substr__MG1655__w=0.049]
FGLIGIHFGRPLQHEGESKGRLTLIHILLGMIPA
>gb|AAC76297_1|ARO_3000499|acrE__Escherichia_coli_str__K-12_substr__MG1655__JM_#01__[gb|AAC76297_1|ARO_3000499|acrE__Escherichia_coli_str__K-
12_substr__MG1655__w=0.032]
VHIVKTAPLEVKTELPGRTNAYRIAEVRPQVSGI
>gb|AAC76297_1|ARO_3000499|acrE__Escherichia_coli_str__K-12_substr__MG1655__JM_#02__[gb|AAC76297_1|ARO_3000499|acrE__Escherichia_coli_str__K-
12_substr__MG1655__w=0.077]
ISQQEYDQAIADARQADAAVIAAKATVESARINL
>gb|AAC76297_1|ARO_3000499|acrE__Escherichia_coli_str__K-12_substr__MG1655__JM_#03__[gb|AAC76297_1|ARO_3000499|acrE__Escherichia_coli_str__K-
12_substr__MG1655__w=0.123]
IVNDKSQVEARPVVASQAIGDKWLISEGLKSGDQ
>gb|AAC76298_1|ARO_3000502|acrF__Escherichia_coli_str__K-12_substr__MG1655__JM_#01__[gb|AAC76298_1|ARO_3000502|acrF__Escherichia_coli_str__K-
12_substr__MG1655__w=0.027]
VAGFVSDNPGTTQDDISDYVASNVKDTLSRLNGV
>gb|AAC76298_1|ARO_3000502|acrF__Escherichia_coli_str__K-12_substr__MG1655__JM_#02__[gb|AAC76298_1|ARO_3000502|acrF__Escherichia_coli_str__K-
12_substr__MG1655__w=0.043,gb|AAC73564_1|ARO_3000216|acrB__Escherichia_coli_str__K-
12_substr__MG1655__w=0.013,gb|AAC76539_1|ARO_3000796|mdtF__Escherichia_coli_str__K-
12_substr__MG1655__w=0.013,gb|EGP45231_1|ARO_3004144|AxyY__Achromobacter_insuavis_AXX-A__w=0.003]
QLNASIIAQTRFKNPEEFGKVTLRVNSDGSVVRL
>gb|AAC76298_1|ARO_3000502|acrF__Escherichia_coli_str__K-12_substr__MG1655__JM_#03__[gb|AAC76298_1|ARO_3000502|acrF__Escherichia_coli_str__K-
12_substr__MG1655__w=0.031]
SLVSVRPNGLEDTAQFKLEVDQEKAQALGVSLSD
>gb|AAC76539_1|ARO_3000796|mdtF__Escherichia_coli_str__K-12_substr__MG1655__JM_#01__[gb|AAC76539_1|ARO_3000796|mdtF__Escherichia_coli_str__K-
12_substr__MG1655__w=0.086]
EEFGKILLKVQQDGSQVLLRDVARVELGAEDYST

>gb|AAC76539_1|ARO_3000796|mdtF__Escherichia_coli_str__K-12_substr__MG1655__JM_#02__[gb|AAC76539_1|ARO_3000796|mdtF__Escherichia_coli_str__K-
12_substr__MG1655__w=0.004,gb|AAC73564_1|ARO_3000216|acrB__Escherichia_coli_str__K-
12_substr__MG1655__w=0.003,gb|AAC76298_1|ARO_3000502|acrF__Escherichia_coli_str__K-12_substr__MG1655__w=0.002]
TPFIEISIQEVFKTLVEAIILVFLVMYLFLQNFR

>gb|AAC76539_1|ARO_3000796|mdtF__Escherichia_coli_str__K-12_substr__MG1655__JM_#03__[gb|AAC76539_1|ARO_3000796|mdtF__Escherichia_coli_str__K-
12_substr__MG1655__w=0.202,gb|AAL14440_1|ARO_3000775|adeB__Acinetobacter_baumannii__w=0.054]
TPALCATILKAAPEGGHKPNALFARFNTLFEKST

>gb|AAF42062_1|ARO_3000811|mtrD__Neisseria_meningitidis_MC58__JM_#01__[gb|AAF42062_1|ARO_3000811|mtrD__Neisseria_meningitidis_MC58__w=0.472]
ISAGSIGSLPAVRGQTVTATVTAQGQLGTAEEFG

>gb|AAF42062_1|ARO_3000811|mtrD__Neisseria_meningitidis_MC58__JM_#02__[gb|AAF42062_1|ARO_3000811|mtrD__Neisseria_meningitidis_MC58__w=0.138]
ATAKAVKERMATLEKYFPQGMSWKTPYDTSKFVE
>gb|AAF42062_1|ARO_3000811|mtrD__Neisseria_meningitidis_MC58__JM_#03__[gb|AAF42062_1|ARO_3000811|mtrD__Neisseria_meningitidis_MC58__w=0.144,gb|AAC7629
8_1|ARO_3000502|acrF__Escherichia_coli_str__K-12_substr__MG1655__w=0.051,gb|AAC73564_1|ARO_3000216|acrB__Escherichia_coli_str__K-
12_substr__MG1655__w=0.051]
MVSVQLPAGATQERTNATLAQVTQLAKSIPEIEN

>gb|AAG05882_1|ARO_3000804|MexF__Pseudomonas_aeruginosa_PAO1__JM_#01__[gb|AAG05882_1|ARO_3000804|MexF__Pseudomonas_aeruginosa_PAO1__w=0.047]
AAGTLGAPPAPSDTSFQLSINTQGRLVTEEEFEN
>gb|AAG05882_1|ARO_3000804|MexF__Pseudomonas_aeruginosa_PAO1__JM_#02__[gb|AAG05882_1|ARO_3000804|MexF__Pseudomonas_aeruginosa_PAO1__w=0.052,gb|A
AP43110_2|ARO_3003923|oqxB__Escherichia_coli__w=0.015]
GHHEPKDRFSVFLDKLLGSWLFRPFNRFFDRASH

>gb|AAG05882_1|ARO_3000804|MexF__Pseudomonas_aeruginosa_PAO1__JM_#03__[gb|AAG05882_1|ARO_3000804|MexF__Pseudomonas_aeruginosa_PAO1__w=0.061]
QIEDRGNQGYEELFKQTQNIITKARALPELEPSS

>gb|AAG05883_1|ARO_3000805|OprN__Pseudomonas_aeruginosa_PAO1__JM_#01__[gb|AAG05883_1|ARO_3000805|OprN__Pseudomonas_aeruginosa_PAO1__w=0.165,gb|A
AD51346_1|ARO_3003053|smeC__Stenotrophomonas_maltophilia__w=0.049,gb|AAG03816_1|ARO_3000379|OprM__Pseudomonas_aeruginosa_PAO1__w=0.044]
IGKGQQPGVTEDRVNSERYDLGLDSAWELDLFGR

>gb|AAG05883_1|ARO_3000805|OprN__Pseudomonas_aeruginosa_PAO1__JM_#02__[gb|AAG05883_1|ARO_3000805|OprN__Pseudomonas_aeruginosa_PAO1__w=0.019]
QLESSDALSEAAEADLQQLQVSLIAELVDAYGQL

>gb|AAG05883_1|ARO_3000805|OprN__Pseudomonas_aeruginosa_PAO1__JM_#03__[gb|AAG05883_1|ARO_3000805|OprN__Pseudomonas_aeruginosa_PAO1__w=0.128]
EKIALSNLENQKESRQLTEQLRDAGVGAELDVLR

>gb|AAG06942_1|ARO_3002985|arnA__Pseudomonas_aeruginosa_PAO1__JM_#01__[gb|AAG06942_1|ARO_3002985|arnA__Pseudomonas_aeruginosa_PAO1__w=0.113]
HPLWLERIRQLRPDFLFSFYYRRLLGAELLACAA

>gb|AAG06942_1|ARO_3002985|arnA__Pseudomonas_aeruginosa_PAO1__JM_#02__[gb|AAG06942_1|ARO_3002985|arnA__Pseudomonas_aeruginosa_PAO1__w=0.155]
ESQASHFGRRTPADGLLDWHRPARQLYDLVRAVT

>gb|AAG06942_1|ARO_3002985|arnA__Pseudomonas_aeruginosa_PAO1__JM_#03__[gb|AAG06942_1|ARO_3002985|arnA__Pseudomonas_aeruginosa_PAO1__w=0.008]
LVEGARLRGAACSPQRRTRVLILGVNGFIGNHLS
>gb|AAK37619_1|ARO_3000481|tet(35)__Vibrio_harveyi__JM_#01__[gb|AAK37619_1|ARO_3000481|tet(35)__Vibrio_harveyi__w=0.124]
ALLMVFAVAWFGLDIGKMREHEIAASQGRGFDKD
>gb|AAK37619_1|ARO_3000481|tet(35)__Vibrio_harveyi__JM_#02__[gb|AAK37619_1|ARO_3000481|tet(35)__Vibrio_harveyi__w=0.442]
DKENDSQEAHDLNEELDIRESEKGKVSDLILPIV
>gb|AAK37619_1|ARO_3000481|tet(35)__Vibrio_harveyi__JM_#03__[gb|AAK37619_1|ARO_3000481|tet(35)__Vibrio_harveyi__w=0.236]
VTLIVATIASMLYTGGQALAADGKEFVLLGAFEN
>gb|AAL14440_1|ARO_3000775|adeB__Acinetobacter_baumannii__JM_#01__[gb|AAL14440_1|ARO_3000775|adeB__Acinetobacter_baumannii__w=0.079]
SLKIDREKLSAFGVKFSDVSDIISTSMGSMYIND
>gb|AAL20862_1|ARO_3000826|sdiA__Salmonella_enterica_subsp__enterica_serovar_Typhimurium_str__LT2__JM_#01__[gb|AAL20862_1|ARO_3000826|sdiA__Salmonella_en
terica_subsp__enterica_serovar_Typhimurium_str__LT2__w=0.069]
AAAEDVYTELQYQTQRLEFDYYALCVRHPVPFTR
>gb|AAL20862_1|ARO_3000826|sdiA__Salmonella_enterica_subsp__enterica_serovar_Typhimurium_str__LT2__JM_#02__[gb|AAL20862_1|ARO_3000826|sdiA__Salmonella_en
terica_subsp__enterica_serovar_Typhimurium_str__LT2__w=0.071]
RPKISLRTTYPPAWVTHYQSENYFAIDPVLKPEN
>gb|AAL20862_1|ARO_3000826|sdiA__Salmonella_enterica_subsp__enterica_serovar_Typhimurium_str__LT2__JM_#03__[gb|AAL20862_1|ARO_3000826|sdiA__Salmonella_en
terica_subsp__enterica_serovar_Typhimurium_str__LT2__w=0.153]
LHWDDVLFHEAKAMWDAAQRFGLRRGVTQCVMLP
>gb|AAL73129_1|ARO_3000616|mel__Streptococcus_pyogenes__JM_#01__[gb|AAL73129_1|ARO_3000616|mel__Streptococcus_pyogenes__w=0.204]
IPKLGDRVQSLDPNFIREMQEGMAVLRQNKGLFA
>gb|AAL73129_1|ARO_3000616|mel__Streptococcus_pyogenes__JM_#02__[gb|AAL73129_1|ARO_3000616|mel__Streptococcus_pyogenes__w=0.044]
PQSGFFIFVVCCAIMGLSVPFYSGVQTALFQEKI

>gb|AAM92464_1|ARO_3002594|AAC(6_)-IIa__Salmonella_enterica_subsp__enterica_serovar_Typhi__JM_#01__[gb|AAM92464_1|ARO_3002594|AAC(6_)-
IIa__Salmonella_enterica_subsp__enterica_serovar_Typhi__w=0.128,gb|AAD46626_1|ARO_3002596|AAC(6_)-
IIc__Pseudomonas_aeruginosa__w=0.068,gb|ABR10839_1|ARO_3002586|AAC(6_)-32__Pseudomonas_aeruginosa__w=0.105,gb|ABR10839_1|ARO_3002586|AAC(6_)-
32__Pseudomonas_aeruginosa__w=0.060,gb|CAK55557_1|ARO_3002585|AAC(6_)-31__Pseudomonas_putida__w=0.083]
```

```
MSASTPPITLRLMTERDLPMLHDWLNRPHIVEWW

>gb|AAM92464_1|ARO_3002594|AAC(6_)-IIa__Salmonella_enterica_subsp__enterica_serovar_Typhi__JM_#02__[gb|AAM92464_1|ARO_3002594|AAC(6_)-
IIa__Salmonella_enterica_subsp__enterica_serovar_Typhi__w=0.215,gb|AAD46626_1|ARO_3002596|AAC(6_)-
IIc__Pseudomonas_aeruginosa__w=0.063,gb|ABR10839_1|ARO_3002586|AAC(6_)-32__Pseudomonas_aeruginosa__w=0.158,gb|ABR10839_1|ARO_3002586|AAC(6_)-
32__Pseudomonas_aeruginosa__w=0.019,gb|CAK55557_1|ARO_3002585|AAC(6_)-31__Pseudomonas_putida__w=0.032,gb|CAK55557_1|ARO_3002585|AAC(6_)-
31__Pseudomonas_putida__w=0.063]
RPTLDEVLEHYLPRAMAEESVTPYIAMLGEEPIG

>gb|AAM92464_1|ARO_3002594|AAC(6_)-IIa__Salmonella_enterica_subsp__enterica_serovar_Typhi__JM_#03__[gb|AAM92464_1|ARO_3002594|AAC(6_)-
IIa__Salmonella_enterica_subsp__enterica_serovar_Typhi__w=0.243,gb|ABR10839_1|ARO_3002586|AAC(6_)-
32__Pseudomonas_aeruginosa__w=0.014,gb|CAK55557_1|ARO_3002585|AAC(6_)-31__Pseudomonas_putida__w=0.071,gb|CAK55557_1|ARO_3002585|AAC(6_)-
31__Pseudomonas_putida__w=0.014,gb|AAA25680_1|ARO_3002595|AAC(6_)-IIb__Pseudomonas_fluorescens__w=0.043]
IDQSLADPTQLNKGLGTRLVRALVELLFSDPTVT
>gb|AAP43110_2|ARO_3003923|oqxB__Escherichia_coli__QM50_#01__[gb|AAP43110_2|ARO_3003923|oqxB__Escherichia_coli__w=0.149]
DQRKHTAAEINAEINAKIAQIQQGFGFSILPPP

>gb|AAP74961_2|ARO_3002854|dfrA1__Klebsiella_oxytoca__QM93_#01__[gb|AAP74961_2|ARO_3002854|dfrA1__Klebsiella_oxytoca__w=0.232,gb|ABB89122_1|ARO_300286
1|dfrA5__Salmonella_enterica_subsp__enterica_serovar_Enteritidis__w=0.162,gb|CAA90683_1|ARO_3003012|dfrA13__Escherichia_coli__w=0.070,gb|ADG84870_1|ARO_30028
58|dfrA12__Klebsiella_pneumoniae__w=0.070,gb|ADG84870_1|ARO_3002858|dfrA12__Klebsiella_pneumoniae__w=0.070,gb|WP_149100971_1|ARO_3005351|DfrA39__Pseud
omonas_aeruginosa__w=0.070,gb|WP_149100971_1|ARO_3005351|DfrA39__Pseudomonas_aeruginosa__w=0.070]
AISXNGVIGNGPDIPWSAKGEQLLFKAITYNQW
>gb|AAR84673_1|ARO_3002936|vanSF__Paenibacillus_popilliae_ATCC_14706__JM_#01__[gb|AAR84673_1|ARO_3002936|vanSF__Paenibacillus_popilliae_ATCC_14706__w=0.3
86]
KVLQLYYYTSVDYGDTLAYFRKIIQNIGDFNVFL
>gb|AAR84673_1|ARO_3002936|vanSF__Paenibacillus_popilliae_ATCC_14706__JM_#02__[gb|AAR84673_1|ARO_3002936|vanSF__Paenibacillus_popilliae_ATCC_14706__w=0.1
92]
LLLFILLSILFFFLLTKPYSAYFNEISKGIHYLA
>gb|AAR84673_1|ARO_3002936|vanSF__Paenibacillus_popilliae_ATCC_14706__JM_#03__[gb|AAR84673_1|ARO_3002936|vanSF__Paenibacillus_popilliae_ATCC_14706__w=0.0
16]
ILKDDNLTENQIRHYLTIAFTKSQRLERLIDELF
>gb|AAT45742_1|ARO_3000988|TEM-126__Escherichia_coli__QM57_#01__[gb|AAT45742_1|ARO_3000988|TEM-
126__Escherichia_coli__w=0.094,gb|ABV89601_1|ARO_3004826|LAP-2__Enterobacter_cloacae__w=0.014,gb|CAA33795_1|ARO_3003563|RCP-
1__Rhodobacter_capsulatus__w=0.011]
PAAMATTLRKLLTGELLTLASRQQLIDWMEADK

>gb|AAU10334_1|ARO_3002628|aad(6)__Streptococcus_oralis__Partial_JM_#01__[gb|AAU10334_1|ARO_3002628|aad(6)__Streptococcus_oralis__Partial_w=0.130,gb|AHE4055
7_1|ARO_3002626|ANT(6)-Ia__Exiguobacterium_sp__S3-2__w=0.038,gb|AHE40557_1|ARO_3002626|ANT(6)-Ia__Exiguobacterium_sp__S3-2__w=0.038]
TYRMDSYENIWEALFLCHQLFRAVSGEVAERLHY

>gb|AAU10334_1|ARO_3002628|aad(6)__Streptococcus_oralis__Partial_JM_#02__[gb|AAU10334_1|ARO_3002628|aad(6)__Streptococcus_oralis__Partial_w=0.486]
DRNITKYTRDMYKKYTGKTGCLDSTYAADIEERR
>gb|ABR10839_1|ARO_3002586|AAC(6_)-32__Pseudomonas_aeruginosa__JM_#01__[gb|ABR10839_1|ARO_3002586|AAC(6_)-
32__Pseudomonas_aeruginosa__w=0.020,gb|AAM92464_1|ARO_3002594|AAC(6_)-
IIa__Salmonella_enterica_subsp__enterica_serovar_Typhi__w=0.017,gb|AAM92464_1|ARO_3002594|AAC(6_)-
IIa__Salmonella_enterica_subsp__enterica_serovar_Typhi__w=0.010,gb|AAD46626_1|ARO_3002596|AAC(6_)-
IIc__Pseudomonas_aeruginosa__w=0.011,gb|CAK55557_1|ARO_3002585|AAC(6_)-31__Pseudomonas_putida__w=0.013,gb|CAK55557_1|ARO_3002585|AAC(6_)-
31__Pseudomonas_putida__w=0.006,gb|CAE48335_2|ARO_3002599|AAC(6_)-30_AAC(6_)-
Ib__fusion_protein__Pseudomonas_aeruginosa__w=0.007,gb|AAA25680_1|ARO_3002595|AAC(6_)-IIb__Pseudomonas_fluorescens__w=0.009]
MSPSKTPVTLRLMTERDLPMLHAWLNRPHIVEWW
>gb|ABR10839_1|ARO_3002586|AAC(6_)-32__Pseudomonas_aeruginosa__JM_#02__[gb|ABR10839_1|ARO_3002586|AAC(6_)-
32__Pseudomonas_aeruginosa__w=0.028,gb|AAM92464_1|ARO_3002594|AAC(6_)-
IIa__Salmonella_enterica_subsp__enterica_serovar_Typhi__w=0.016,gb|AAM92464_1|ARO_3002594|AAC(6_)-
IIa__Salmonella_enterica_subsp__enterica_serovar_Typhi__w=0.007,gb|AAD46626_1|ARO_3002596|AAC(6_)-
IIc__Pseudomonas_aeruginosa__w=0.007,gb|CAK55557_1|ARO_3002585|AAC(6_)-31__Pseudomonas_putida__w=0.003,gb|CAK55557_1|ARO_3002585|AAC(6_)-
31__Pseudomonas_putida__w=0.008,gb|CAE48335_2|ARO_3002599|AAC(6_)-30_AAC(6_)-
Ib__fusion_protein__Pseudomonas_aeruginosa__w=0.007,gb|AAA25680_1|ARO_3002595|AAC(6_)-IIb__Pseudomonas_fluorescens__w=0.002]
WGGEEERPTLHEVVKHYLPRVLAEEAVTPYIAML

>gb|ABR10839_1|ARO_3002586|AAC(6_)-32__Pseudomonas_aeruginosa__JM_#03__[gb|ABR10839_1|ARO_3002586|AAC(6_)-
32__Pseudomonas_aeruginosa__w=0.160,gb|AAM92464_1|ARO_3002594|AAC(6_)-
IIa__Salmonella_enterica_subsp__enterica_serovar_Typhi__w=0.028,gb|AAD46626_1|ARO_3002596|AAC(6_)-
IIc__Pseudomonas_aeruginosa__w=0.028,gb|CAK55557_1|ARO_3002585|AAC(6_)-31__Pseudomonas_putida__w=0.028,gb|AAL82588_1|ARO_3002600|AAC(3)-Ib_AAC(6_)-
Ib____Pseudomonas_aeruginosa__w=0.028,gb|AAA25680_1|ARO_3002595|AAC(6_)-IIb__Pseudomonas_fluorescens__w=0.028]
GVRGIDQFLSNHTQLNQGLGTKLVQALVELLFSD
>gb|ABV18113_1|ARO_3001329|mdtG__Escherichia_coli_O139_H28_str__E24377A__JM_#01__[gb|ABV18113_1|ARO_3001329|mdtG__Escherichia_coli_O139_H28_str__E2437
7A__w=0.193]
SVLILCFFVTLFCIREKFQPVSKKEMLHMREVVT
>gb|ABV18113_1|ARO_3001329|mdtG__Escherichia_coli_O139_H28_str__E24377A__JM_#02__[gb|ABV18113_1|ARO_3001329|mdtG__Escherichia_coli_O139_H28_str__E2437
7A__w=0.067]
LITALIFSVLLLIPMSYVQTPLQLGILRFLLGAA
>gb|ABV18113_1|ARO_3001329|mdtG__Escherichia_coli_O139_H28_str__E24377A__JM_#03__[gb|ABV18113_1|ARO_3001329|mdtG__Escherichia_coli_O139_H28_str__E2437
7A__w=0.026]
ISANYGFRAVFLVTAGVVLFNAVYSWNSLRRRRI

>gb|ACL19707_1|ARO_3003725|vanXI__Desulfitobacterium_hafniense_DCB-2__JM_#01__[gb|ACL19707_1|ARO_3003725|vanXI__Desulfitobacterium_hafniense_DCB-
2__w=0.011,gb|AAM09852_1|ARO_3003070|vanXD__Enterococcus_faecium__w=0.007,gb|AHA41501_1|ARO_3002954|vanXO__Rhodococcus_hoagii__w=0.007,gb|AAA65957_
1|ARO_3002949|vanXA__Enterococcus_faecium__w=0.007,gb|AAB05628_1|ARO_3002950|vanXB__Enterococcus_faecalis__w=0.007,gb|AAF36804_1|ARO_3002952|vanXF__Pa
enibacillus_popilliae_ATCC_14706__w=0.007,gb|ACL82962_1|ARO_3002953|vanXM__Enterococcus_faecium__w=0.006]
MKSDFVFVDELVSGIRWDAKYATWDNFTGKPVDG
>gb|ACL19707_1|ARO_3003725|vanXI__Desulfitobacterium_hafniense_DCB-2__JM_#02__[gb|ACL19707_1|ARO_3003725|vanXI__Desulfitobacterium_hafniense_DCB-
2__w=0.044]
HYPNIDRSEIIEKGYVAAKSGHSRGSAIDLTLYH


>gb|ACL19707_1|ARO_3003725|vanXI__Desulfitobacterium_hafniense_DCB-2__JM_#03__[gb|ACL19707_1|ARO_3003725|vanXI__Desulfitobacterium_hafniense_DCB-
2__w=0.147,gb|AAM09852_1|ARO_3003070|vanXD__Enterococcus_faecium__w=0.052,gb|AAM09852_1|ARO_3003070|vanXD__Enterococcus_faecium__w=0.039]
HLASGTLVPMGGDFDLMDSVSHHGAHGISQAEAR
>gb|ACL82957_1|ARO_3002928|vanRM__Enterococcus_faecium__JM_#01__[gb|ACL82957_1|ARO_3002928|vanRM__Enterococcus_faecium__w=0.150]
MRRISILIAEDEEEIADLLAIHLEKEGYDVIKVH
>gb|ACL82957_1|ARO_3002928|vanRM__Enterococcus_faecium__JM_#02__[gb|ACL82957_1|ARO_3002928|vanRM__Enterococcus_faecium__w=0.007]
HDGQEALHVIQAQSIDLIILDIMMPKMDGYEVTR
>gb|ACL82957_1|ARO_3002928|vanRM__Enterococcus_faecium__JM_#03__[gb|ACL82957_1|ARO_3002928|vanRM__Enterococcus_faecium__w=0.065,gb|AAR84672_1|ARO_3
002925|vanRF__Paenibacillus_popilliae_ATCC_14706__w=0.052,gb|AAA65953_1|ARO_3002919|vanRA__Enterococcus_faecium__w=0.011]
RQVRAQYNMPIIFLSAKTSDFDKVHGLVIGGDDY
>gb|AEJ33969_1|ARO_3000410|sul1__Vibrio_fluvialis__JM_#01__[gb|AEJ33969_1|ARO_3000410|sul1__Vibrio_fluvialis__w=0.246]
RVGSDVVDVGPAASHPDARPVSPADEIRRIAPLL
>gb|AEJ33969_1|ARO_3000410|sul1__Vibrio_fluvialis__JM_#02__[gb|AEJ33969_1|ARO_3000410|sul1__Vibrio_fluvialis__w=0.250]
DQMHRVSIDSFQPETQRYALKRGVGYLNDIQGFP
>gb|AEJ33969_1|ARO_3000410|sul1__Vibrio_fluvialis__JM_#03__[gb|AEJ33969_1|ARO_3000410|sul1__Vibrio_fluvialis__w=0.312]
LGPASLAAELHAIGNGADYVRTHAPGDLRSAITF
```

```
>gb|AFO09968_1|ARO_3001476|OXA-184__Campylobacter_jejuni__QM44_#01__[gb|AFO09968_1|ARO_3001476|OXA-
184__Campylobacter_jejuni__w=0.194,gb|AKI29908_1|ARO_3003603|OXA-447__Campylobacter_jejuni__w=0.194,gb|AAT01092_1|ARO_3001773|OXA-
61__Campylobacter_jejuni__w=0.012]
SAIKRSQVPAFKELARKIGLKTMQESLNKLSYG
>gb|AFP97028_1|ARO_3002228|IMP-37__Pseudomonas_aeruginosa__JM_#01__[gb|AFP97028_1|ARO_3002228|IMP-
37__Pseudomonas_aeruginosa__w=0.257,gb|BAB72072_1|ARO_3002202|IMP-11__Acinetobacter_baumannii__w=0.143,gb|BAB72072_1|ARO_3002202|IMP-
11__Acinetobacter_baumannii__w=0.095,gb|AIT76110_1|ARO_3002239|IMP-48__Pseudomonas_aeruginosa__w=0.095,gb|AEH41427_1|ARO_3002218|IMP-
27__Proteus_mirabilis__w=0.143,gb|AEH41427_1|ARO_3002218|IMP-27__Proteus_mirabilis__w=0.095,gb|BAM62793_1|ARO_3002233|IMP-
42__Acinetobacter_soli__w=0.095,gb|AFO59566_1|ARO_3002226|IMP-35__Pseudomonas_aeruginosa__w=0.038]
DAYLIDTPFTATDTEKLVNWFVERGYX
>gb|AGC51118_1|ARO_3000617|mecA__Staphylococcus_aureus__JM_#01__[gb|AGC51118_1|ARO_3000617|mecA__Staphylococcus_aureus__w=0.330]
INNTIDAIEDKNFKQVYKDSSYISKSDNGEVEMT
>gb|AGC51118_1|ARO_3000617|mecA__Staphylococcus_aureus__JM_#02__[gb|AGC51118_1|ARO_3000617|mecA__Staphylococcus_aureus__w=0.162,gb|CCC86795_1|ARO_3
001209|mecC__Staphylococcus_aureus_subsp__aureus_LGA251__w=0.052]
AKKFHLTTNETKSRNYPLEKATSHLLGYVGPINS
>gb|AGC51118_1|ARO_3000617|mecA__Staphylococcus_aureus__JM_#03__[gb|AGC51118_1|ARO_3000617|mecA__Staphylococcus_aureus__w=0.164]
DNSNTIAHTLIEKKKKDGKDIQLTIDAKVQKSIY
>gb|AHH83938_1|ARO_3000013|vanB__Enterococcus_faecium__JM_#01__[gb|AHH83938_1|ARO_3000013|vanB__Enterococcus_faecium__w=0.386,gb|AAA65956_1|ARO_300
0010|vanA__Enterococcus_faecium__w=0.114,gb|KTE89608_1|ARO_3003723|vanI__Desulfitobacterium_hafniense__w=0.114,gb|KTE89608_1|ARO_3003723|vanI__Desulfitobac
terium_hafniense__w=0.114]
KSAIEIAANIDTEKFDPHYIGITKNGVWKLCKKP
>gb|AHH83938_1|ARO_3000013|vanB__Enterococcus_faecium__JM_#02__[gb|AHH83938_1|ARO_3000013|vanB__Enterococcus_faecium__w=0.296]
TEWEADSLPAILSPDRKTHGLLVMKESEYETRRI

>gb|AHH83938_1|ARO_3000013|vanB__Enterococcus_faecium__JM_#03__[gb|AHH83938_1|ARO_3000013|vanB__Enterococcus_faecium__w=0.028,gb|AAM09849_1|ARO_30
00005|vanD__Enterococcus_faecium__w=0.012,gb|AAA65956_1|ARO_3000010|vanA__Enterococcus_faecium__w=0.012,gb|ACL82961_1|ARO_3002911|vanM__Enterococcus_f
aecium__w=0.018,gb|ACL82961_1|ARO_3002911|vanM__Enterococcus_faecium__w=0.008,gb|KTE89608_1|ARO_3003723|vanI__Desulfitobacterium_hafniense__w=0.009,gb|K
TE89608_1|ARO_3003723|vanI__Desulfitobacterium_hafniense__w=0.005,gb|AAF36803_1|ARO_3002908|vanF__Paenibacillus_popilliae_ATCC_14706__w=0.008]
CGEDGAIQGLFVLSGIPYVGCDIQSSAACMDKSL
>gb|AIA35096_1|ARO_3005059|LptD__Klebsiella_pneumoniae_subsp__pneumoniae_KPNIH10__JM_#01__[gb|AIA35096_1|ARO_3005059|LptD__Klebsiella_pneumoniae_subsp
__pneumoniae_KPNIH10__w=0.061]
LATHYQQSNLDKYNAANGTDYKESVSRVMPQFKV
>gb|AIA35096_1|ARO_3005059|LptD__Klebsiella_pneumoniae_subsp__pneumoniae_KPNIH10__JM_#02__[gb|AIA35096_1|ARO_3005059|LptD__Klebsiella_pneumoniae_subsp
__pneumoniae_KPNIH10__w=0.022]
GKMVFERDLQEGFTQTLEPRVQYLYVPYRDQSEI
>gb|AIS67611_1|ARO_3003163|CTX-M-155__Klebsiella_pneumoniae__QM52_#01__[gb|AIS67611_1|ARO_3003163|CTX-M-
155__Klebsiella_pneumoniae__w=0.123,gb|CAI43424_1|ARO_3002415|OXY-6-3__Klebsiella_oxytoca__w=0.056,gb|ABA00479_1|ARO_3001859|IMI-
2__Enterobacter_asburiae__w=0.041]
AQTLRNLTLGKALGDSQRAQLVTWMKGNTTGAA
>gb|AKD43328_1|ARO_3004757|BKC-1__Klebsiella_pneumoniae__JM_#01__[gb|AKD43328_1|ARO_3004757|BKC-1__Klebsiella_pneumoniae__w=0.337]
AVSTLAASAGALLAVPAVSTLAASAGAATGGPLE
>gb|AKD43328_1|ARO_3004757|BKC-1__Klebsiella_pneumoniae__JM_#02__[gb|AKD43328_1|ARO_3004757|BKC-
1__Klebsiella_pneumoniae__w=0.070,gb|AWB15813_1|ARO_3005096|GPC-1__Pseudomonas_aeruginosa__w=0.004]
EKRLAELEGRHKGRIGVAIHNLATGARIGHRADE
>gb|AKD43328_1|ARO_3004757|BKC-1__Klebsiella_pneumoniae__JM_#03__[gb|AKD43328_1|ARO_3004757|BKC-
1__Klebsiella_pneumoniae__w=0.347,gb|AWB15813_1|ARO_3005096|GPC-1__Pseudomonas_aeruginosa__w=0.031,gb|AWB15813_1|ARO_3005096|GPC-
1__Pseudomonas_aeruginosa__w=0.102,gb|AWB15813_1|ARO_3005096|GPC-1__Pseudomonas_aeruginosa__w=0.102]
KEETLDRRIVVGKSDLVDWSPVVETRVGGEGISI


>gb|AKN19287_1|ARO_3002578|AAC(6_)-Ib7__Salmonella_enterica_subsp__enterica_serovar_Corvallis__QM54_#01__[gb|AKN19287_1|ARO_3002578|AAC(6_)-
Ib7__Salmonella_enterica_subsp__enterica_serovar_Corvallis__w=0.108,gb|CAE48335_2|ARO_3002599|AAC(6_)-30_AAC(6_)-
Ib__fusion_protein__Pseudomonas_aeruginosa__w=0.108,gb|AAL82588_1|ARO_3002600|AAC(3)-Ib_AAC(6_)-
Ib____Pseudomonas_aeruginosa__w=0.108,gb|AAD46626_1|ARO_3002596|AAC(6_)-IIc__Pseudomonas_aeruginosa__w=0.039,gb|AAD46626_1|ARO_3002596|AAC(6_)-
IIc__Pseudomonas_aeruginosa__w=0.036,gb|AAM92464_1|ARO_3002594|AAC(6_)-
IIa__Salmonella_enterica_subsp__enterica_serovar_Typhi__w=0.036,gb|ABR10839_1|ARO_3002586|AAC(6_)-32__Pseudomonas_aeruginosa__w=0.036]
VTKIQTDPSPSNLRAIRCYEKAGFERQGTVTTP
>gb|APB03216_1|ARO_3003982|LlmA_23S_ribosomal_RNA_methyltransferase__Paenibacillus_sp__LC231__JM_#01__[gb|APB03216_1|ARO_3003982|LlmA_23S_ribosomal_RN
A_methyltransferase__Paenibacillus_sp__LC231__w=0.049]
QIIWPIEREDGQWRSVHGHYHRSSSGGGQWDMKK
>gb|APB03216_1|ARO_3003982|LlmA_23S_ribosomal_RNA_methyltransferase__Paenibacillus_sp__LC231__JM_#02__[gb|APB03216_1|ARO_3003982|LlmA_23S_ribosomal_RN
A_methyltransferase__Paenibacillus_sp__LC231__w=0.004]
RWTISYDQLKFHIRPTNFKHTGLFPEQAANWRWM
>gb|APB03216_1|ARO_3003982|LlmA_23S_ribosomal_RNA_methyltransferase__Paenibacillus_sp__LC231__JM_#03__[gb|APB03216_1|ARO_3003982|LlmA_23S_ribosomal_RN
A_methyltransferase__Paenibacillus_sp__LC231__w=0.006]
MDKIASAGRPVSVLNLFAYTGGATVAAASAGAQV
>gb|BAA15934_1|ARO_3000829|baeS__Escherichia_coli_str__K-12_substr__W3110__JM_#01__[gb|BAA15934_1|ARO_3000829|baeS__Escherichia_coli_str__K-
12_substr__W3110__w=0.044]
RSFEHDNSEDKPGPGMPPHGWRTQFWVVDQNNKV
>gb|BAA15934_1|ARO_3000829|baeS__Escherichia_coli_str__K-12_substr__W3110__JM_#02__[gb|BAA15934_1|ARO_3000829|baeS__Escherichia_coli_str__K-
12_substr__W3110__w=0.048]
YQKAPVDLIPLLEVAGGAFRERFASRGLKLQFSL
>gb|BAA15934_1|ARO_3000829|baeS__Escherichia_coli_str__K-12_substr__W3110__JM_#03__[gb|BAA15934_1|ARO_3000829|baeS__Escherichia_coli_str__K-
12_substr__W3110__w=0.059]
SGGSLQISAGQRDKTVRLTFADSAPGVSDDQLQK
>gb|BAA15935_1|ARO_3000828|baeR__Escherichia_coli_str__K-12_substr__W3110__JM_#01__[gb|BAA15935_1|ARO_3000828|baeR__Escherichia_coli_str__K-
12_substr__W3110__w=0.018,gb|AAD51348_1|ARO_3003066|smeR__Stenotrophomonas_maltophilia__w=0.006,gb|CCP46065_1|ARO_3000816|mtrA__Mycobacterium_tuberc
ulosis_H37Rv__w=0.007]
SYAPTLISHGDQVLPYVRQTPPDLILLDLMLPGT
>gb|BAA15935_1|ARO_3000828|baeR__Escherichia_coli_str__K-12_substr__W3110__JM_#02__[gb|BAA15935_1|ARO_3000828|baeR__Escherichia_coli_str__K-
12_substr__W3110__w=0.116]
LRRCKPQRELQQQDAESPLIIDEGRFQASWRGKM
>gb|BAA15935_1|ARO_3000828|baeR__Escherichia_coli_str__K-12_substr__W3110__JM_#03__[gb|BAA15935_1|ARO_3000828|baeR__Escherichia_coli_str__K-
12_substr__W3110__w=0.012]
GKVFSREQLLNHLYDDYRVVTDRTIDSHIKNLRR
>gb|BAA16344_1|ARO_3000491|acrD__Escherichia_coli_str__K-12_substr__W3110__QM67_#01__[gb|BAA16344_1|ARO_3000491|acrD__Escherichia_coli_str__K-
12_substr__W3110__w=0.052]
KGEHHGQKGFFAWFNQMFNRNAERYEKGVAKIL

>gb|BAA16547_1|ARO_3000027|emrA__Escherichia_coli_str__K-12_substr__W3110__JM_#01__[gb|BAA16547_1|ARO_3000027|emrA__Escherichia_coli_str__K-
12_substr__W3110__w=0.034,gb|EHL92831_1|ARO_3004588|Klebsiella_pneumoniae_KpnG__Klebsiella_sp__4_1_44FAA__w=0.007]
LFIIIAVAIGIYWFLVLRHFEETDDAYVAGNQIQ

>gb|BAA16547_1|ARO_3000027|emrA__Escherichia_coli_str__K-12_substr__W3110__JM_#02__[gb|BAA16547_1|ARO_3000027|emrA__Escherichia_coli_str__K-
12_substr__W3110__w=0.039,gb|EHL92831_1|ARO_3004588|Klebsiella_pneumoniae_KpnG__Klebsiella_sp__4_1_44FAA__w=0.014]
VKEGDVLVTLDPTDARQAFEKAKTALASSVRQTH

>gb|BAA16547_1|ARO_3000027|emrA__Escherichia_coli_str__K-12_substr__W3110__JM_#03__[gb|BAA16547_1|ARO_3000027|emrA__Escherichia_coli_str__K-
12_substr__W3110__w=0.010,gb|EHL92831_1|ARO_3004588|Klebsiella_pneumoniae_KpnG__Klebsiella_sp__4_1_44FAA__w=0.002,gb|AAF40763_1|ARO_3003961|farA__Neiss
eria_meningitidis_MC58__w=0.000]
RIELDQKQLEQYPLRIGLSTLVSVNTTNRDGQVL
>gb|BAA78619_1|ARO_3002544|AAC(3)-Xa__Streptomyces_griseus__JM_#01__[gb|BAA78619_1|ARO_3002544|AAC(3)-
Xa__Streptomyces_griseus__w=0.137,gb|AAA88552_1|ARO_3002541|AAC(3)-VIIa__Streptomyces_rimosus__w=0.024]
```

SDGPVTRDRIRHDLAALGLVPGDTVMFHTRLSAI
>gb|BAA78619_1|ARO_3002544|AAC(3)-Xa__Streptomyces_griseus__JM_#02__[gb|BAA78619_1|ARO_3002544|AAC(3)-
Xa__Streptomyces_griseus__w=0.032,gb|AAA88552_1|ARO_3002541|AAC(3)-VIIa__Streptomyces_rimosus__w=0.015,gb|AAA88552_1|ARO_3002541|AAC(3)-
VIIa__Streptomyces_rimosus__w=0.015,gb|AAA25334_1|ARO_3002543|AAC(3)-IXa__Micromonospora_chalcea__w=0.009,gb|AAA26685_1|ARO_3002542|AAC(3)-
VIIIa__Streptomyces_fradiae__w=0.009]
SGGPQTVIDALLDVVGPTGTLLVTCGWNDAPPYD
>gb|BAA78619_1|ARO_3002544|AAC(3)-Xa__Streptomyces_griseus__JM_#03__[gb|BAA78619_1|ARO_3002544|AAC(3)-
Xa__Streptomyces_griseus__w=0.064,gb|AAA88552_1|ARO_3002541|AAC(3)-VIIa__Streptomyces_rimosus__w=0.009,gb|AAA88552_1|ARO_3002541|AAC(3)-
VIIa__Streptomyces_rimosus__w=0.009,gb|AAA88552_1|ARO_3002541|AAC(3)-VIIa__Streptomyces_rimosus__w=0.008,gb|AAA25334_1|ARO_3002543|AAC(3)-
IXa__Micromonospora_chalcea__w=0.002]
DFTDWPPAWQEAVRAHHPAFDPRTSEAEHANGRL
>gb|BAB38260_1|ARO_3000830|cpxA__Escherichia_coli_O157_H7_str__Sakai__JM_#01__[gb|BAB38260_1|ARO_3000830|cpxA__Escherichia_coli_O157_H7_str__Sakai__w=0.0
40]
AFEAEQMGKSLTVNFPPGPWPLYGNPNALESALE
>gb|BAB82500_1|ARO_3000186|tetM__Erysipelothrix_rhusiopathiae__JM_#01__[gb|BAB82500_1|ARO_3000186|tetM__Erysipelothrix_rhusiopathiae__w=0.062,gb|CAA10975_
1|ARO_3000194|tetW__Butyrivibrio_fibrisolvens__w=0.002]
IEGNDDLLEKYTSGKLLEALELEQEESIRFHNCS

>gb|BAE06008_1|ARO_3003699|mexQ__Pseudomonas_aeruginosa__JM_#01__[gb|BAE06008_1|ARO_3003699|mexQ__Pseudomonas_aeruginosa__w=0.081,gb|AAL19305_1|A
RO_3000790|mdsB__Salmonella_enterica_subsp__enterica_serovar_Typhimurium_str__LT2__w=0.007,gb|AAL19305_1|ARO_3000790|mdsB__Salmonella_enterica_subsp__ente
rica_serovar_Typhimurium_str__LT2__w=0.019,gb|AAX14802_1|ARO_3000781|adeJ__Acinetobacter_baumannii__w=0.026]
QSPGANALDTAEAVRATVARLEGNFPAGLSARIA
>gb|BAE06008_1|ARO_3003699|mexQ__Pseudomonas_aeruginosa__JM_#02__[gb|BAE06008_1|ARO_3003699|mexQ__Pseudomonas_aeruginosa__w=0.093,gb|CAJ77856_1|A
RO_3000777|adeF__Acinetobacter_baumannii_AYE__w=0.011]
AGLLLRPRPAGGAVAGRFQRLLQVLGRPLRNAPE
>gb|BAE06008_1|ARO_3003699|mexQ__Pseudomonas_aeruginosa__JM_#03__[gb|BAE06008_1|ARO_3003699|mexQ__Pseudomonas_aeruginosa__w=0.106]
PGLGTIGGFKMQVEDRGGAGLEALARQTQVLMMK
>gb|BAE15963_1|ARO_3002868|dfrG__Staphylococcus_aureus__JM_#01__[gb|BAE15963_1|ARO_3002868|dfrG__Staphylococcus_aureus__w=0.254,gb|AAA85213_1|ARO_300
2866|dfrD__Listeria_monocytogenes__w=0.239]
PIILGRKNLESIGRALPX
>gb|BAE77933_1|ARO_3000518|CRP__Escherichia_coli_str__K-12_substr__W3110__JM_#01__[gb|BAE77933_1|ARO_3000518|CRP__Escherichia_coli_str__K-
12_substr__W3110__w=0.060]
HCHIHKYPSKSKLIHQGEKAETLYYIVKGSVAVL
>gb|BAE77933_1|ARO_3000518|CRP__Escherichia_coli_str__K-12_substr__W3110__JM_#02__[gb|BAE77933_1|ARO_3000518|CRP__Escherichia_coli_str__K-
12_substr__W3110__w=0.062]
DILMRLSAQMARRLQVTSEKVGNLAFLDVTGRIA
>gb|BAE77933_1|ARO_3000518|CRP__Escherichia_coli_str__K-12_substr__W3110__JM_#03__[gb|BAE77933_1|ARO_3000518|CRP__Escherichia_coli_str__K-
12_substr__W3110__w=0.006]
AQTLLNLAKQPDAMTHPDGMQIKITRQEIGQIVG
>gb|BAE78084_1|ARO_3003548|mdtN__Escherichia_coli_str__K-12_substr__W3110__JM_#01__[gb|BAE78084_1|ARO_3003548|mdtN__Escherichia_coli_str__K-
12_substr__W3110__w=0.028]
RIVELAVTDNQAVKQGDLLFRIDPRPYEANLAKA
>gb|BAE78084_1|ARO_3003548|mdtN__Escherichia_coli_str__K-12_substr__W3110__JM_#02__[gb|BAE78084_1|ARO_3003548|mdtN__Escherichia_coli_str__K-
12_substr__W3110__w=0.241]
VDAQQFGADSVNATVEKARAAAKQATDTLRRTEP
>gb|BAE78084_1|ARO_3003548|mdtN__Escherichia_coli_str__K-12_substr__W3110__JM_#03__[gb|BAE78084_1|ARO_3003548|mdtN__Escherichia_coli_str__K-
12_substr__W3110__w=0.075]
PLLKEGFVSAEDVDRARTAQRAAEADLNAVLLQA
>gb|BAJ42218_1|ARO_3004612|Escherichia_coli_ampH__Escherichia_coli_DH1__JM_#01__[gb|BAJ42218_1|ARO_3004612|Escherichia_coli_ampH__Escherichia_coli_DH1__w=0
.052]
REQRWKYLSTAKLKAAPGSQAAYSNLAFDLLADA
>gb|BAJ42218_1|ARO_3004612|Escherichia_coli_ampH__Escherichia_coli_DH1__JM_#02__[gb|BAJ42218_1|ARO_3004612|Escherichia_coli_ampH__Escherichia_coli_DH1__w=0
.058]
SGKPYTQLFEEQITRPLGMKDTTYTPSPDQCRRL
>gb|BAJ42218_1|ARO_3004612|Escherichia_coli_ampH__Escherichia_coli_DH1__JM_#03__[gb|BAJ42218_1|ARO_3004612|Escherichia_coli_ampH__Escherichia_coli_DH1__w=0
.076]
VTRSPLTRFKNMSDGINDLVTELSGNKPLVIPAS
>gb|CAA23899_1|ARO_3002683|catI__Escherichia_coli__JM_#01__[gb|CAA23899_1|ARO_3002683|catI__Escherichia_coli__w=0.330,gb|AAA25655_1|ARO_3004657|catA4__Pr
oteus_mirabilis__w=0.097]
MEKKITGYTTVDISQWHRKEHFEAFQSVAQCTYN
>gb|CAA23899_1|ARO_3002683|catI__Escherichia_coli__JM_#02__[gb|CAA23899_1|ARO_3002683|catI__Escherichia_coli__w=0.248,gb|AAA25655_1|ARO_3004657|catA4__Pr
oteus_mirabilis__w=0.117]
KHKFYPAFIHILARLMNAHPEFRMAMKDGELVIW
>gb|CAA23899_1|ARO_3002683|catI__Escherichia_coli__JM_#03__[gb|CAA23899_1|ARO_3002683|catI__Escherichia_coli__w=0.288,gb|AAA25655_1|ARO_3004657|catA4__Pr
oteus_mirabilis__w=0.042,gb|AAB51421_1|ARO_3002686|catP__Clostridium_perfringens__w=0.085]
WDSVHPCYTVFHEQTETFSSLWSEYHDDFRQFLH
>gb|CAA79727_1|ARO_3000191|tetQ__Bacteroides_fragilis__JM_#01__[gb|CAA79727_1|ARO_3000191|tetQ__Bacteroides_fragilis__w=0.187]
MEDFRIGDYLGTKPCLIQGLSHQHPALKSSVRPD
>gb|CAA79727_1|ARO_3000191|tetQ__Bacteroides_fragilis__JM_#02__[gb|CAA79727_1|ARO_3000191|tetQ__Bacteroides_fragilis__w=0.166]
DEIKTIYKERPVKKVNKIIQIEVPPNPYWATIGL
>gb|CAA79727_1|ARO_3000191|tetQ__Bacteroides_fragilis__JM_#03__[gb|CAA79727_1|ARO_3000191|tetQ__Bacteroides_fragilis__w=0.126]
VKPCGYQITKGDYSDNIRMNEKDKLLFMFQKSMS

>gb|CAB45459_1|ARO_3002914|vanJ__Streptomyces_coelicolor_A3(2)__JM_#01__[gb|CAB45459_1|ARO_3002914|vanJ__Streptomyces_coelicolor_A3(2)__w=0.081]
KRAPVLTVSALLLGLVMMLHAEIPNRFGSVGSLV

>gb|CAB45459_1|ARO_3002914|vanJ__Streptomyces_coelicolor_A3(2)__JM_#02__[gb|CAB45459_1|ARO_3002914|vanJ__Streptomyces_coelicolor_A3(2)__w=0.063]
SAAAVTALVLPVTVWLSLFGGLLGDKSGAGGEFT

>gb|CAB45459_1|ARO_3002914|vanJ__Streptomyces_coelicolor_A3(2)__JM_#03__[gb|CAB45459_1|ARO_3002914|vanJ__Streptomyces_coelicolor_A3(2)__w=0.017]
DVLALEEITAQDREVYEKGLAREYPYHTVQGTVG
>gb|CAC86410_1|ARO_3000599|Erm(33)__Staphylococcus_sciuri__JM_#01__[gb|CAC86410_1|ARO_3000599|Erm(33)__Staphylococcus_sciuri__w=0.206,gb|AAA98296_1|ARO_
3000250|ErmC__Staphylococcus_epidermidis__w=0.042,gb|AAA98296_1|ARO_3000250|ErmC__Staphylococcus_epidermidis__w=0.109,gb|AAA98096_1|ARO_3000595|ErmT__
Plasmid_pGT633__w=0.018,gb|AAC37034_1|ARO_3000522|ErmG__Bacteroides_thetaiotaomicron__w=0.018,gb|AAC37034_1|ARO_3000522|ErmG__Bacteroides_thetaiotaomic
ron__w=0.073]
MNKKNIKDSQNFITSKRNIDKIMTNISLNEHDNI
>gb|CAE18141_1|ARO_3003111|lsaB__Staphylococcus_sciuri__JM_#01__[gb|CAE18141_1|ARO_3003111|lsaB__Staphylococcus_sciuri__w=0.029,gb|AAO43110_1|ARO_300030
0|lsaA__Enterococcus_faecalis_ATCC_29212__w=0.004]
RGKTTLFNLLLDKFEYRGKIISSVDFNYFPYPVE
>gb|CAE18141_1|ARO_3003111|lsaB__Staphylococcus_sciuri__JM_#02__[gb|CAE18141_1|ARO_3003111|lsaB__Staphylococcus_sciuri__w=0.078,gb|AEA37904_1|ARO_3003112
|lsaC__Streptococcus_agalactiae__w=0.025,gb|AXF35727_1|ARO_3005121|lsa(D)__Lactococcus_garvieae__w=0.025]
TVSNYLRKKKGNILISHDRNFLDGSVDHILSINR
>gb|CAE18141_1|ARO_3003111|lsaB__Staphylococcus_sciuri__JM_#03__[gb|CAE18141_1|ARO_3003111|lsaB__Staphylococcus_sciuri__w=0.109]
RQQGHEQATNERLQKDIGRLEQSTKRSAGWSNRV

>gb|CAJ77856_1|ARO_3000777|adeF__Acinetobacter_baumannii_AYE__JM_#01__[gb|CAJ77856_1|ARO_3000777|adeF__Acinetobacter_baumannii_AYE__w=0.167]
LFNRVFSRASDRYSQGVSRVISHKASAMGVYAAL

>gb|CAJ77856_1|ARO_3000777|adeF__Acinetobacter_baumannii_AYE__JM_#02__[gb|CAJ77856_1|ARO_3000777|adeF__Acinetobacter_baumannii_AYE__w=0.037]
MVPLSSLVNVTQTYGPEMVVRYNGYTSADINGGP
>gb|CCP46065_1|ARO_3000816|mtrA__Mycobacterium_tuberculosis_H37Rv__JM_#01__[gb|CCP46065_1|ARO_3000816|mtrA__Mycobacterium_tuberculosis_H37Rv__w=0.029
]
MDTMRQRILVVDDDASLAEMLTIVLRGEGFDTAV

>gb|CCP46065_1|ARO_3000816|mtrA__Mycobacterium_tuberculosis_H37Rv__JM_#02__[gb|CCP46065_1|ARO_3000816|mtrA__Mycobacterium_tuberculosis_H37Rv__w=0.010
,gb|BAA15935_1|ARO_3000828|baeR__Escherichia_coli_str__K-
12_substr__W3110__w=0.004,gb|AAD51348_1|ARO_3003066|smeR__Stenotrophomonas_maltophilia__w=0.003]
VIGDGTQALTAVRELRPDLVLLDLMLPGMNGIDV
>gb|CCP46065_1|ARO_3000816|mtrA__Mycobacterium_tuberculosis_H37Rv__JM_#03__[gb|CCP46065_1|ARO_3000816|mtrA__Mycobacterium_tuberculosis_H37Rv__w=0.013
]
VCRVLRADSGVPIVMLTAKTDTVDVVLGLESGAD
>gb|CCP46182_1|ARO_3003035|mfpA__Mycobacterium_tuberculosis_H37Rv__JM_#01__[gb|CCP46182_1|ARO_3003035|mfpA__Mycobacterium_tuberculosis_H37Rv__w=0.07
3]
EDLSRLHTERAMFSECDFSGVNLAESQHRGSAFR
>gb|CCP46182_1|ARO_3003035|mfpA__Mycobacterium_tuberculosis_H37Rv__JM_#02__[gb|CCP46182_1|ARO_3003035|mfpA__Mycobacterium_tuberculosis_H37Rv__w=0.15
5]
VFVACRLRPLTLDDVDFTLAVLGGNDLRGLNLTG
>gb|CCP46182_1|ARO_3003035|mfpA__Mycobacterium_tuberculosis_H37Rv__JM_#03__[gb|CCP46182_1|ARO_3003035|mfpA__Mycobacterium_tuberculosis_H37Rv__w=0.10
8]
ETSLVDTDLRKCVLRGADLSGARTTGARLDDADL
>gb|EIM09515_1|ARO_3004747|BAT-1__Bacillus_atrophaeus_C89__JM_#01__[gb|EIM09515_1|ARO_3004747|BAT-1__Bacillus_atrophaeus_C89__w=0.515]
FTTSVQPISEASAAEKNLNIKKLDVDEFFADHNG
>gb|EIM09515_1|ARO_3004747|BAT-1__Bacillus_atrophaeus_C89__JM_#02__[gb|EIM09515_1|ARO_3004747|BAT-1__Bacillus_atrophaeus_C89__w=0.124]
GTFILRDMNKGKTFIYNHERANQRFAPQSTFKVP
>gb|EIM09515_1|ARO_3004747|BAT-1__Bacillus_atrophaeus_C89__JM_#03__[gb|EIM09515_1|ARO_3004747|BAT-
1__Bacillus_atrophaeus_C89__w=0.094,gb|ABV63006_2|ARO_3004759|BPU-1__Bacillus_pumilus_SAFR-032__w=0.061,gb|ABV63006_2|ARO_3004759|BPU-
1__Bacillus_pumilus_SAFR-032__w=0.033]
GLQVGAVEDEYSIKYWDGVKREIDIWNQDHTLGS
>gb|KTE89608_1|ARO_3003723|vanI__Desulfitobacterium_hafniense__JM_#01__[gb|KTE89608_1|ARO_3003723|vanI__Desulfitobacterium_hafniense__w=0.054,gb|AHA41500
_1|ARO_3002913|vanO__Rhodococcus_hoagii__w=0.016,gb|AHH83938_1|ARO_3000013|vanB__Enterococcus_faecium__w=0.005,gb|AHH83938_1|ARO_3000013|vanB__Enter
ococcus_faecium__w=0.013,gb|AHH83938_1|ARO_3000013|vanB__Enterococcus_faecium__w=0.013,gb|AAM09849_1|ARO_3000005|vanD__Enterococcus_faecium__w=0.019,g
b|AAF36803_1|ARO_3002908|vanF__Paenibacillus_popilliae_ATCC_14706__w=0.018]
SEEHPVSVKSAGEVAKNLDPEKYEPFYIGITKDG
>gb|KTE89608_1|ARO_3003723|vanI__Desulfitobacterium_hafniense__JM_#02__[gb|KTE89608_1|ARO_3003723|vanI__Desulfitobacterium_hafniense__w=0.121,gb|AHH83938
_1|ARO_3000013|vanB__Enterococcus_faecium__w=0.007,gb|AHH83938_1|ARO_3000013|vanB__Enterococcus_faecium__w=0.007]
GVWQLCHYPEVNWEKGSCRPAILSPDRSVQGLLV
>gb|KTE89608_1|ARO_3003723|vanI__Desulfitobacterium_hafniense__JM_#03__[gb|KTE89608_1|ARO_3003723|vanI__Desulfitobacterium_hafniense__w=0.013,gb|AHA41500
_1|ARO_3002913|vanO__Rhodococcus_hoagii__w=0.004,gb|AHA41500_1|ARO_3002913|vanO__Rhodococcus_hoagii__w=0.004,gb|AAA65956_1|ARO_3000010|vanA__Enteroc
occus_faecium__w=0.000,gb|ACL82961_1|ARO_3002911|vanM__Enterococcus_faecium__w=0.000,gb|ABA71731_1|ARO_3002909|vanG__Enterococcus_faecalis__w=0.004,gb|
AAA24786_1|ARO_3000368|vanC__Enterococcus_gallinarum__w=0.005]
VLEQGQYQRIPLDLVFPVLHGKFGEDGAMQGLLE
>gb|WP_041777404_1|ARO_3004480|Bifidobacterium_adolescentis_rpoB_conferring_resistance_to_rifampicin__Bifidobacterium_adolescentis__JM_#01__[gb|WP_041777404_
1|ARO_3004480|Bifidobacterium_adolescentis_rpoB_conferring_resistance_to_rifampicin__Bifidobacterium_adolescentis__w=0.128]
MAEATTNTTTIIARADQHDIDLHKASDRVNFGSI
>gb|WP_041777404_1|ARO_3004480|Bifidobacterium_adolescentis_rpoB_conferring_resistance_to_rifampicin__Bifidobacterium_adolescentis__JM_#02__[gb|WP_041777404_
1|ARO_3004480|Bifidobacterium_adolescentis_rpoB_conferring_resistance_to_rifampicin__Bifidobacterium_adolescentis__w=0.101]
ERWQKRVQEDLENGTNTVPHTSGLDEVFQEISPI
>gb|WP_041777404_1|ARO_3004480|Bifidobacterium_adolescentis_rpoB_conferring_resistance_to_rifampicin__Bifidobacterium_adolescentis__JM_#03__[gb|WP_041777404_
1|ARO_3004480|Bifidobacterium_adolescentis_rpoB_conferring_resistance_to_rifampicin__Bifidobacterium_adolescentis__w=0.029]
KINRKLGLEKDVNDRSLSREDIIATIKYLVTLHA
>gb|WP_064483990_1|ARO_3005335|OXA-496__Acinetobacter_lwoffii__QM98_#01__[gb|WP_064483990_1|ARO_3005335|OXA-
496__Acinetobacter_lwoffii__w=0.206,gb|ENV18923_1|ARO_3001731|OXA-275__Acinetobacter_guillouiae_NIPH_991__w=0.069,gb|AGW16411_1|ARO_3001517|OXA-
329__Acinetobacter_calcoaceticus__w=0.031]
NALIGLQHGKATTNEIFKWDGKKRSFXAWEKDM
